# Neurofilament accumulation disrupts autophagy in giant axonal neuropathy

**DOI:** 10.1101/2024.03.29.587353

**Authors:** Jean-Michel Paumier, James Zewe, Melissa R Pergande, Meghana Venkatesan, Eitan Israeli, Chiranjit Panja, Natasha Snider, Jeffrey Savas, Puneet Opal

## Abstract

Neurofilament accumulation is a marker of several neurodegenerative diseases, but it is the primary pathology in Giant Axonal Neuropathy (GAN). This childhood onset autosomal recessive disease is caused by loss-of-function mutations in gigaxonin, the E3 adaptor protein that is essential for neurofilament degradation. Using a combination of genetic and RNA interference (RNAi) approaches, we found that dorsal root ganglia from mice lacking gigaxonin have impaired autophagy and lysosomal degradation through two mechanisms. First, neurofilament accumulations interfere with the distribution of autophagic organelles, impairing their maturation and fusion with lysosomes. Second, the accumulations sequester the chaperone 14-3-3, a protein responsible for the localization of the transcription factor EB (TFEB), a key regulator of autophagy. This dual disruption of autophagy likely contributes to the pathogenesis of other neurodegenerative diseases with neurofilament accumulations.

## INTRODUCTION

Giant axonal neuropathy (GAN) affects virtually every facet of the peripheral and central nervous systems (1, 2). Patients initially present with weakness, wasting, and sensory deficits in a glove and stocking distribution. As the disease progresses, patients suffer from cognitive decline, cranial nerve dysfunction, seizures, spasticity, and cerebellar incoordination. GAN has no treatment, and most patients do not live beyond the third decade of life. This distal-to-proximal pattern of deterioration reflects the physical impediment caused by mislocalization and dysfunction of certain cytoskeletal proteins, which manifests first in those neurons with the longest axons.

GAN is caused by biallelic loss-of-function mutations of the *GAN* gene (1, 3), which encodes gigaxonin, a member of the BTB (Bric-a-brac, Tramtrack and Broad)-Kelch family of E3 ligase adaptors that recruit substrates for ubiquitin-mediated proteasomal degradation (4). The N-terminal BTB domain binds Cullin3, which serves as a bridge to the rest of the ubiquitination machinery (5); the C terminal Kelch domain binds select substrates. The best characterized GAN substrates are several intermediate filament proteins (IFs), which belong to the larger family of cytoskeletal proteins (6–8). IFs are so named because their 10 nm diameter places them intermediate between the sizes of the two other major cytoskeletal proteins, actin and microtubules (9). They are classified into six major types (I-VI) based on their primary structure and tissue of expression (9, 10). All IFs share a tripartite structure, including variable globular N- and C-terminal domains and a central conserved rod domain. It is this central rod domain to which gigaxonin binds.

Neurons have the most complex repertoire of IFs of any cell type, and they bear the brunt of GAN pathology. All neurons express three of the type IV proteins: neurofilament triplet proteins neurofilament heavy (NFH), middle (NFM), and light (NFL), so named based on their molecular weights. Some neurons also express alpha internexin (a type IV IF), while yet others, particularly those in the peripheral nervous system, express peripherin, a type III IF (11). The highly polarized morphology of neurons creates special metabolic and cytoskeletal demands. Neurons from the spinal cord, for instance, send long axons to skin and muscle at vast distances from the perikaryon. When neurofilament degradation is stalled, neurofilament homeostasis is dysregulated, compromising the mechanical properties, and signaling events that they regulate (11, 12). In GAN, neurofilaments accumulate throughout the cytoplasmic space but also in discrete foci that are remarkably stable (6). Recent research using photoactivatable IFs suggests that individual filaments cannot escape from these IF bundles because of a disruption in their kinesin-based transport along microtubules (13).

We have been interested in the effects of NF accumulation downstream from the obvious mechanical obstacles because subcellular localization and transport are crucial for many cellular functions. Indeed, it has remained unclear whether other dysregulated proteins are involved in GAN pathogenesis. Previously, we found that NF accumulations interfere with mitochondrial movement, thereby impairing mitochondrial function (6). Here we report that NF accumulations also alter the spatial distribution of lysosomes, which impedes autophagic processes that are spatially orchestrated in neurons (normally, substrates are engulfed by autophagic vesicles more distally, then are transported retrogradely from the neurites to the soma, where they are delivered to the lysosomes for degradation) (14). We also show that the NF foci sequester 14-3-3 proteins, a family of chaperone proteins known to bind IFs, along with TFEB (the master transcriptional regulator transcription factor EB), thereby preventing TFEB from shuttling to the nucleus to perform its transcriptional functions. The result is a progressive loss of quality control of both proteins and organelles, causing cellular deterioration.

## RESULTS

### Proteomic analysis of mouse dorsal root ganglia with gigaxonin silenced

*GAN* null mice recapitulate the neurofilament aggregates in neurons and the IF aggregates in other cell types, but because of their small size, they do not manifest overt signs of the disease until they are very old (6, 15–17). We therefore developed a primary neuronal culture model using dorsal root ganglia (DRG) neurons (6), which are affected early in the disease course and display severe neuropathology. DRG neurons isolated from *GAN* null mice demonstrate progressive neurofilament accumulation starting from as early as two days *in vitro*, both in the cell soma and neurites **(Fig 1A)**. The GAN phenotype is equally well reproduced in wild-type neurons using small hairpin (sh) RNA based RNAi to reduce gigaxonin expression. The degree of knockdown achieved is approximately 90% (6). Neurons lacking gigaxonin degenerate, as evidenced by axonal fragmentation after an additional 8-9 days in culture **(Fig 1B)**.

**Fig. 1.**
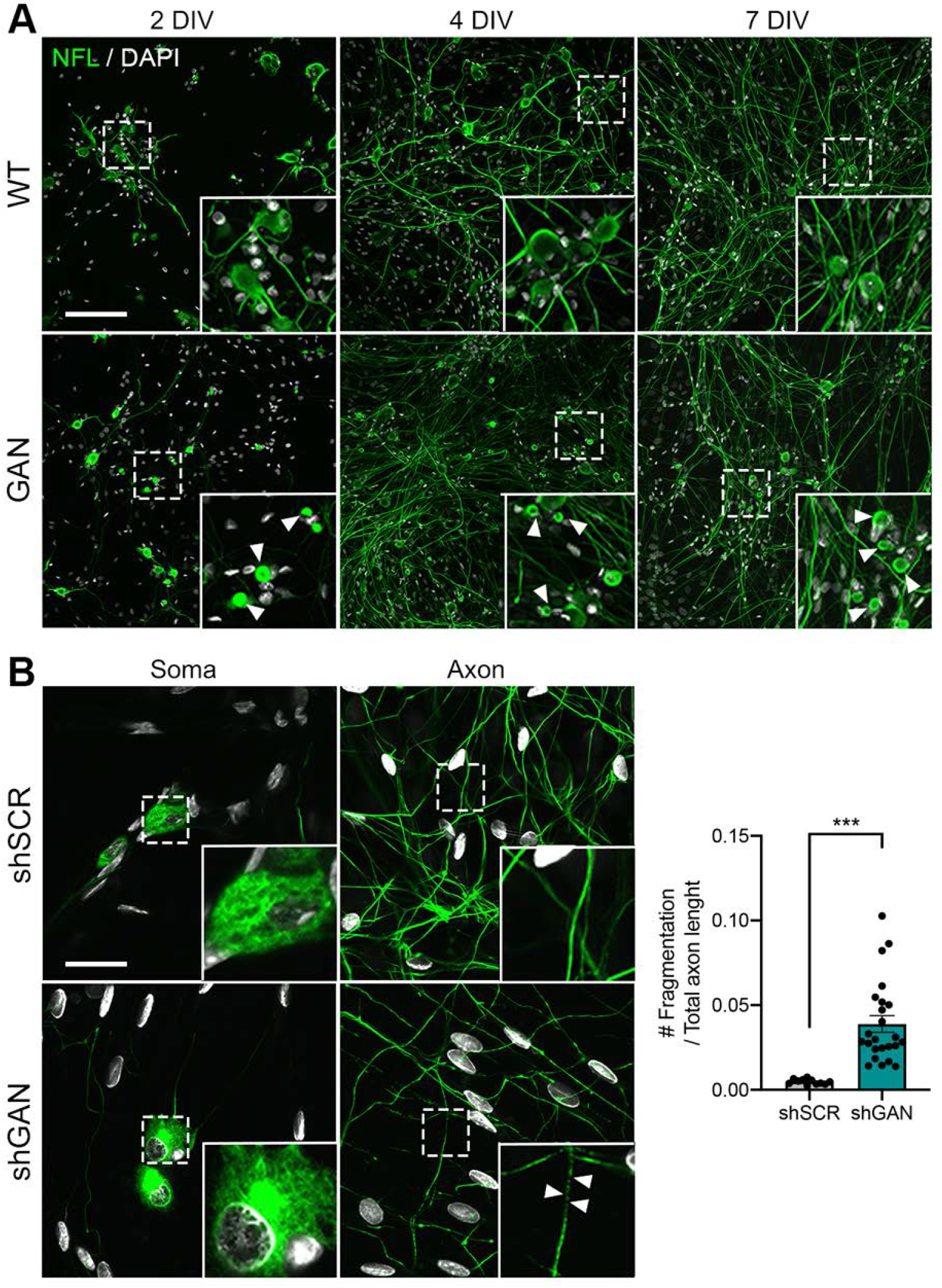
Dorsal root ganglia (DRG) neurons model the hallmark pathology of giant axonal neuropathy (GAN). (A) Representative fluorescence microscopy images of dorsal root ganglia (DRGs) from WT and *GAN* knockout mice stained for neurofilament light (NFL) after 2, 4, and 7 days *in vitro* (DIV). Arrowheads denote NFL aggregation in the soma of GAN DRGs. Note that NFL aggregates are already present after 2 DIV. Scale bar, 30 µm. (B) The GAN phenotype can be recapitulated by lentiviral delivery of shRNA targeting the *Gan* gene. Scale bar, 30 µm. After 12 DIV, axonal fragmentation, a sign of neurodegeneration, occurs in GAN-silenced DRGs, as denoted by arrowheads; large aggregate shown in zoom. (C) Quantification of axonal fragmentation was accomplished using Fiji’s measurement tool, reported here as the means of 3 independent experiments +/-SEM. ***p<0.001 by a two-tailed unpaired t-test.

To gain insight into the molecular consequences of loss of gigaxonin function, we performed proteomics with stable isotope labeling using amino acids in cell culture (SILAC) (18). We quantified 3,507 proteins in DRG cultures (shSCR) and gigaxonin-knockdown cells (shGAN) under the same experimental conditions **(Fig 2A)**. Of these, 149 proteins had significantly altered fold change with a false discovery rate (FDR) adjusted with a p value of <0.05 (62 elevated and 87 reduced; see **Table S1**). As expected, the IF proteins (NFL, NFM, and peripherin) were among those that were most significantly elevated.

**Fig. 2.**
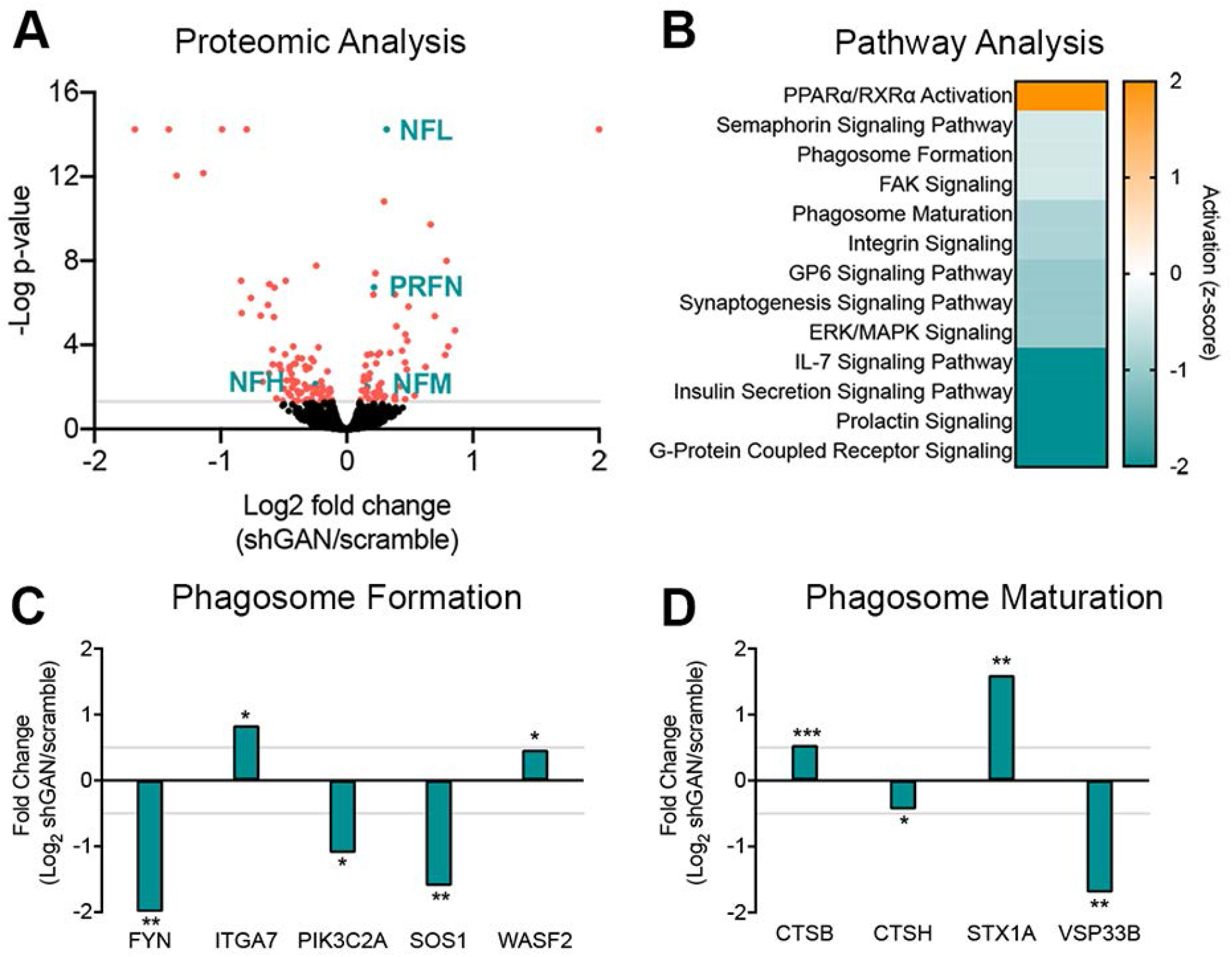
Mass spectrometry-based proteomic analysis of dorsal root ganglia (DRG) cultures silenced for gigaxonin. (A) Volcano plot showing the distribution of measured proteins extracted from DRG cultures in which *GAN* was silenced using shRNA (shGAN). (B) Plot showing top-ranked altered pathways in GAN where pathway activation (change in z-score) is shown (orange represents up-regulation and blue represents down-regulation). (C-D) Plots showing altered proteins for the phagosome formation and phagosome maturation pathways. Adjusted p-values denoted by *p<0.05, **p<0.001, and ***p<0.0001. NFL = neurofilament light, NFM= neurofilament medium, and PRFN = peripherin.

Ingenuity Pathway Analysis (IPA) identified altered biological functions in GAN-silenced DRG cultures (**Fig 2B**). Several signaling pathways were altered, but PPARa/RXRa, which regulates cell growth, differentiation, and metabolism, was the only pathway that was hyperactivated. Several other signaling pathways were downregulated. Interestingly, phagosome formation and maturation, two key aspects of autophagy (19), were downregulated, along with IL-7 signaling (which upregulates the expression of anti-apoptotic genes) and insulin signaling (which inhibits autophagy), which were strongly suppressed. The overall impression is of a very dysregulated autophagic system. For example, FYN, which is known to play a role in autophagy by AMPK phosphorylation (19), was strongly downregulated (**Fig 1C**), as were PIK3C2A, whose knockdown has been described to decrease autophagy and the maturation of endocytic vesicles (21); and SOS1, whose deletion has been related to accumulation of phagosomes and lysosomal bodies (31). ITGA7, which participates in phagocytosis (30), and WASF2, which is involved in autophagosome and trafficking to lysosome in human immune cells (24), were moderately upregulated compared to controls. In the phagosome maturation category (**Fig 1D**), we had upregulation of cathepsin B (CTSB), whose deletion impairs autophagy and lysosomal recycling (32), and Syntaxin 1A (STX1A), which regulates vesicle fusion and trafficking (26); among the downregulated proteins were cathepsin H (CTSH) (33), which is also involved in vesicle fusion and trafficking, and VPS33B, which is involved in endosomal recycling and late endosomal-lysosomal fusion events (27, 34). These results are supported by a recent publication describing the role of gigaxonin in autophagosome production through Atg16L1 turnover regulation (35).

### Neurofilament aggregates alter the spatial distribution, abundance, and morphology of autophagic organelles

The autophagic process involves the formation of vesicles around substrates; these vesicles mature into autophagosomes that fuse with lysosomes, where the substrate is ultimately degraded (20). These steps require the free movement of autophagic organelles, a process that we hypothesized would be particularly compromised in neurons by the space-occupying neurofilament aggregates. Therefore, we stained cells for LC3, a small polypeptide that recruits substrates and is a specific marker for autophagosomes (21). Neurofilaments were delineated by staining for NFL, a protein that forms the backbone of the neurofilament heteropolymer (12). In *GAN* knockout DRG neurons, the autophagosomes were not uniformly distributed throughout the cytoplasm, as they would normally be, but were now more abundant at the perimeters of the aggregates **(Fig 3A)**.

**Fig. 3.**
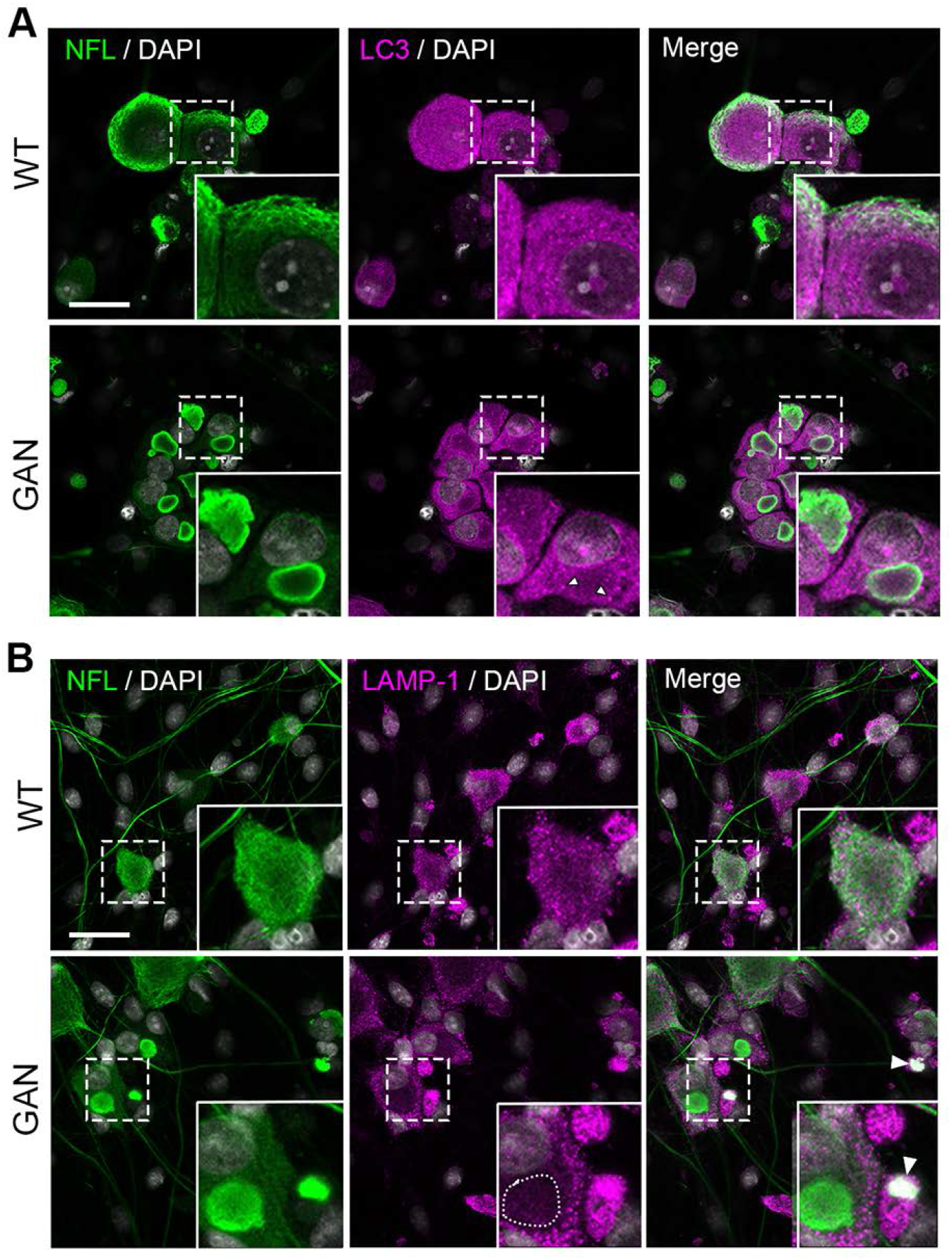
Neurofilament aggregates alter autophagosome spatial distribution. (A) Representative fluorescence images of DRG neurons from WT or *GAN* knockout mice co-stained for the cytoskeleton marker neurofilament light (NFL) and the autophagosome marker LC3. Autophagosomes are excluded from neurofilament aggregates, and LC3 puncta are found at the periphery of the aggregates (white arrowhead). (B) WT or *GAN* knockout mouse DRGs co-stained for the cytoskeletal marker NFL and the lysosomal marker LAMP-1. Two phenotypes are observed in GAN DRG neurons with neurofilament aggregates: lysosomes are either excluded (circular dotted line) or co-localized with neurofilament aggregates (arrowhead). Scale bar, 30 µm. Representative images from three independent experiments.

Next, we determined the location of lysosomes using immunofluorescence microscopy. Neurofilaments were visualized as before by staining for NFL, while lysosomes were visualized by staining for LAMP-1, which constitutes approximately 50% of the lysosomal membrane (12, 22) **(Fig. 3B)**. Neurons showed either exclusion of LAMP-1 staining from neurofilament aggregates, or they showed a colocalization. We surmised that these two patterns reflect the status of different lysosomal populations or their membranous fragments.

To confirm this observation, we stained additional lysosomal components, both across the membrane and within the lumen. For the former, we evaluated the distribution of mucolipin 1, a calcium channel protein and member of the transient receptor potential cation channel mucolipin subfamily (23, 24), and vacuolar ATPase, which is a protein essential for lysosomal acidification that pumps protons into the lumen (25, 26). Both of these co-localized with neurofilament aggregates **(Fig 4A)**. To study intraluminal proteins, we stained for two lysosomal proteases: cathepsin B and cathepsin D (27). Cathepsin B was typically sequestered in aggregates, while cathepsin D was typically excluded from them, reminiscent of the two staining patterns of LAMP-1 **(Fig 4B)**.

**Fig. 4.**
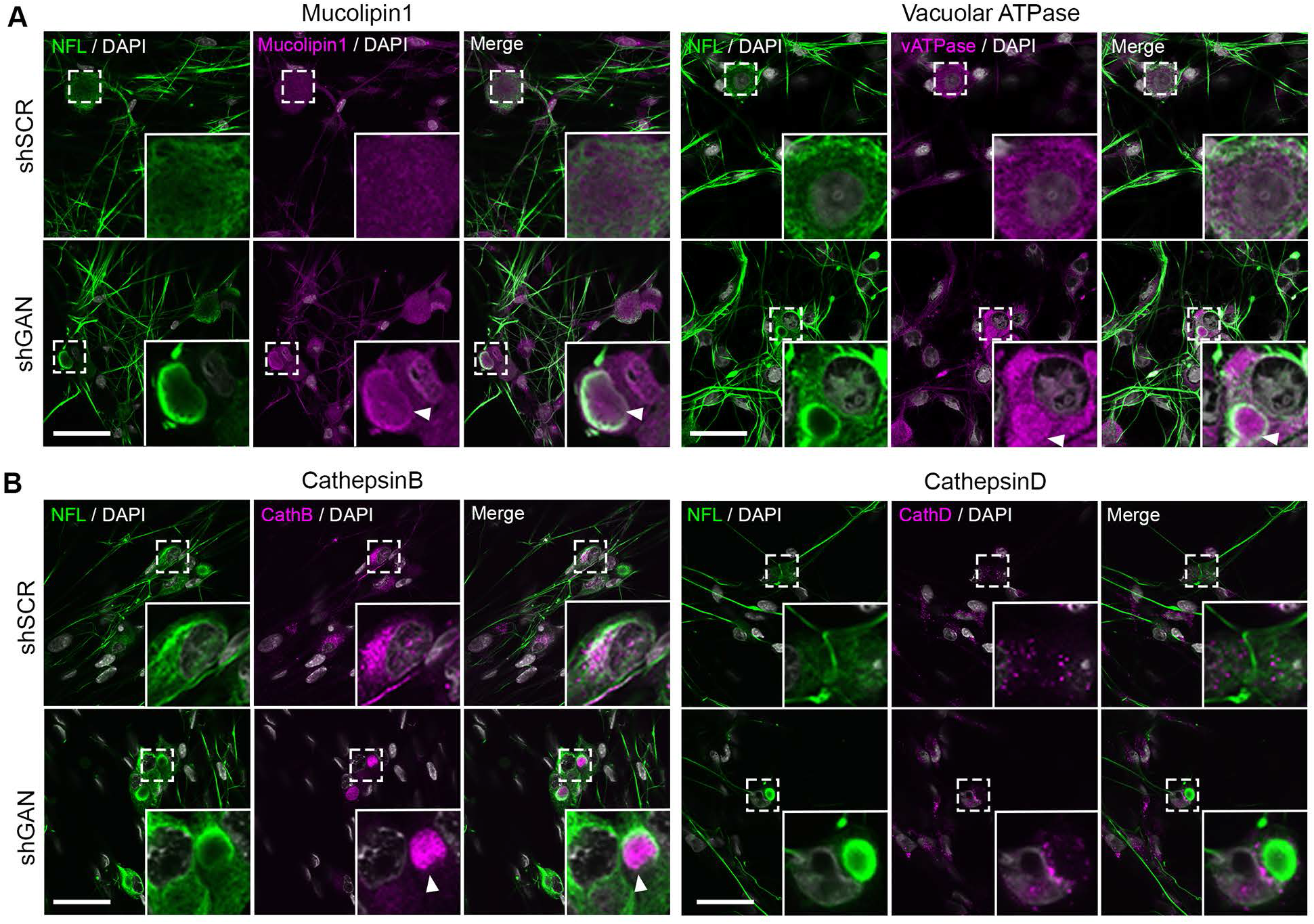
Spatial distribution of lysosomes is altered by neurofilament aggregates in GAN DRG neurons. Representative fluorescence images of DRG neurons silenced for gigaxonin (shGAN) and control neurons (shSCR) co-stained for NFL and (A) mucolipin-1 and vacuolar ATPase or (B) cathepsin B and cathepsin D. While cathepsin D is excluded from neurofilament aggregates, the three other lysosomal proteins are sequestered in aggregates in shGAN DRG neurons. Scale bar, 30 µm. Arrowheads highlight lysosomal proteins clumped in NFL aggregates in the shGAN condition.

We next tracked intact lysosomes in living DRG neurons with lysotracker, a cell-permeable dye that stains the acidic compartment of lysosomes (28). Since we wished to correlate lysosomal distribution with neurofilament aggregates, we also infected cells with a lentivirus expressing GFP-tagged NFL. Lysotracker staining tended to be absent from regions with aggregates, suggesting that intact lysosomes are spatially excluded from neurofilament accumulations **(Fig 5A)**. The amount and intensity of lysosomal staining with lysotracker was greater in GAN DRG cultures **(Fig 5A, B)**, likely because the lysosomes also tended to be larger (Fig 5C).

**Fig. 5.**
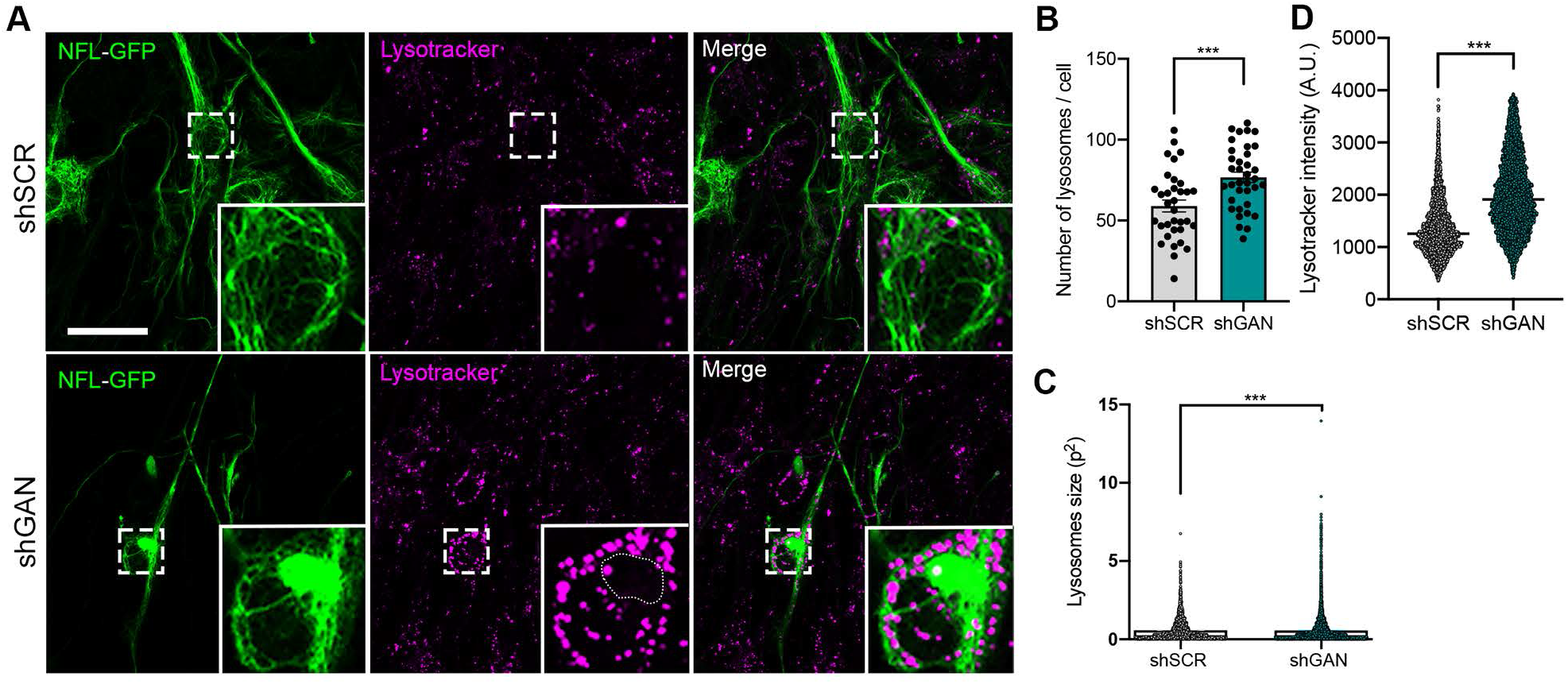
Lysosome alterations in GAN DRG neurons. (A) Representative live imaging fluorescence images of control and shGAN DRGs transduced with NFL-GFP tagged lentivirus and treated with red lysotracker to visualize lysosomes. Expression of the NFL-GFP construct in shGAN DRG neurons allowed neurofilament aggregate visualization in living cells. shGAN induces an increase in the number of lysosomes (B) and the surface area covered by these organelles (C). Lysotracker mean intensity is also increased in shGAN cultures (D). Note that as observed after NFL and LAMP-1 co-staining, lysotracker dye is also mainly excluded from neurofilament aggregates (circular dotted line). Scale bar, 30 µm. Quantitative data are presented as means +/-SEM, with ***p<0.001 by a two-tailed unpaired t-test. Representative images from three independent experiments.

Transmission electron microscopy revealed a greater abundance of autophagic organelles at different stages of maturation in GAN DRGs: these included large autophagic vacuoles, multilamellar bodies, and immature autophagosomes **(Fig 6)**. The electron-dense lysosomes tended to decorate the perimeter of the aggregates. These data demonstrate that NF aggregates influence the distribution of autophagic organelles.

**Fig. 6.**
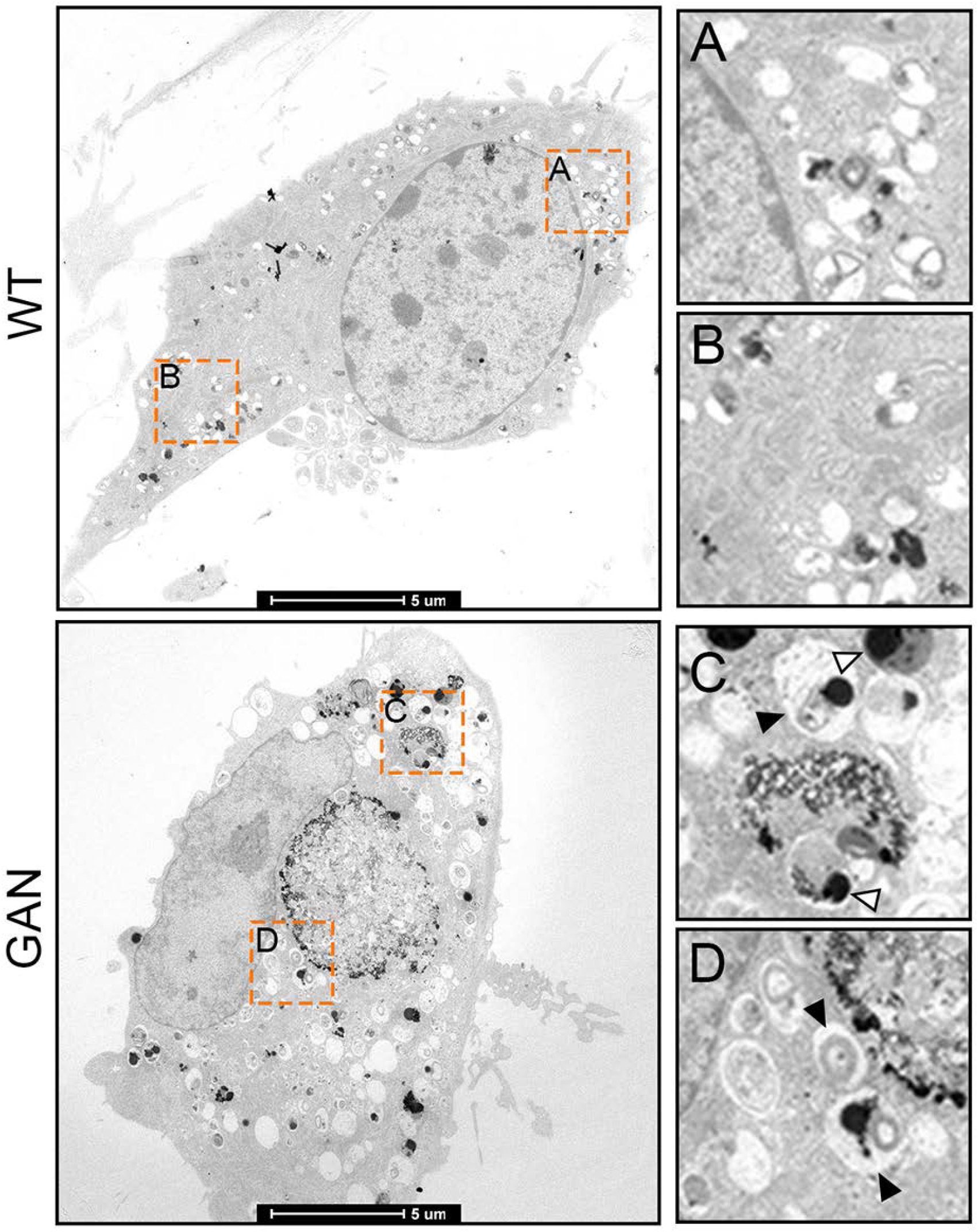
GAN is associated with abnormal autophagic organelles. Transmission electronic microscopy (TEM) microphotographs of control and GAN DRGs showing the presence of a variety of autophagic organelles including large autophagic vacuoles, dark dense lysosomes (open arrowheads), multilamellar bodies, and immature autophagosomes (filled arrowheads).

### Lysosomal acidity and autophagic flux are dysregulated in GAN

To address lysosomal function, we evaluated lysosomal pH, which is crucial to the ability to degrade substrates. We used lysosensor, a sensitive dye designed specifically for this purpose (20). *GAN-*silenced DRG cultures displayed a significant reduction in lysosensor signal intensity within lysotracker defined compartments **(Fig 7A)**. These results suggest that lysosomes in GAN are defective at maintaining a robust acidic internal environment, which would translate into a reduction in autophagic flux. To test this possibility, we performed live-cell imaging using an mRFP-GFP tandem-tagged LC3. This protein incorporates into the membrane of autophagic vacuoles, exhibiting a punctate signal within cells. It fluoresces from fluorophores as autophagosomes mature before fusing with the lysosome; after fusion, the GFP fluorescence (which is pH-sensitive) is lost in the acidic lysosomal milieu, while the mRFP fluorescence (not pH-sensitive) persists until LC3 is fully degraded, providing a quantifiable readout for fusion delay or lysosomal dysfunction (21). *GAN*-silenced neurons showed a significantly greater GFP fluorescence signal within RFP-positive vehicles, suggesting either a greater quantity of autophagosomes yet to fuse with lysosomes or a loss of lysosomal acidity **(Fig 7B)**. These autophagic organelles were larger in GAN DRGs than in controls.

**Fig. 7.**
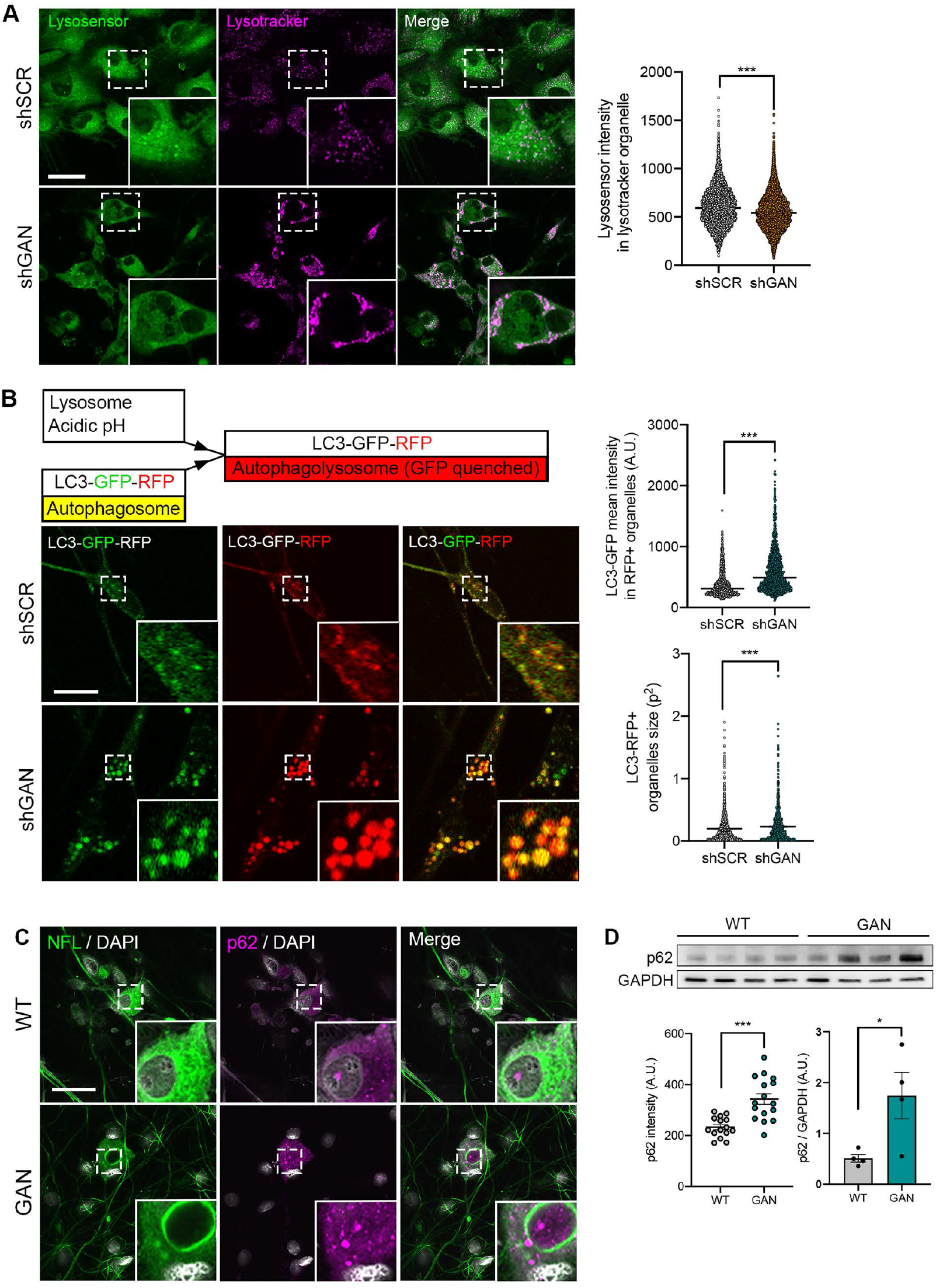
Autophagic flux is altered in GAN DRG cultures. Lysosomal pH is dysregulated in GAN DRG cultures. (A) Representative live imaging microphotographs of control and GAN DRG cultures treated with a combination of red lysotracker to visualize lysosomes and green lysosensor to evaluate pH alterations. Mean intensity of lysosensor (pH sensitive probe) is decreased in lysosomes from GAN cultures, suggesting lysosomal milieu is less acidic in the GAN condition. (B) Representative live imaging microphotographs of control or GAN DRG cells transduced with the sensor LC3-GFP-RFP. Individual panels are presented for GFP, RFP, and merged signals. The construct is composed of an RFP pH-resistant tag, a GFP pH-sensitive tag, and LC3 that targets the tags to nascent autophagosomes. After fusion with lysosomes, autophagolysosomes are formed and the GFP signal is quenched due to the acidic milieu provided by lysosomes, converting the fluorescent signal from yellow to red. The GAN condition is associated with an increase in autophagic organelles positive for both GFP and RFP signals. Autophagic organelle size is also increased in GAN. (C) Representative fluorescent images of control and GAN DRG neurons co-stained for NFL and p62. (D) Immunoblot of p62 in control and GAN DRG cultures. Both p62 mean intensity and protein levels are increased in GAN DRG cultures. Quantitative data are presented as means +/-SEM, with *p<0.05 or ***p<0.001 the two-tailed P values from an unpaired t-test. Scale bar, 30 µm.

To further assess the functional status of autophagy, we evaluated the levels of p62 (also known as SQSTM1), an autophagic receptor that recruits cargo to be degraded and is itself degraded by autophagy (21). By both immunostaining and western blotting, p62 levels were significantly greater in *GAN*-silenced DRGs, consistent with the inability of lysosomes to degrade substrates **(Fig 7C, D)**.

### TFEB is sequestered in neurofilament aggregates in GAN

Autophagy is dependent on the activity of the transcription factor TFEB (29–31), whose activity is highly dependent on its subcellular location. Under conditions of cellular stress, TFEB translocate to the nucleus and drive the CLEAR (Coordinated Lysosomal Expression and Regulation) network of genes responsible for autophagy and lysosomal biogenesis (32). When phosphorylated, TFEB is bound to the cytoplasmic chaperone 14-3-3 proteins, a family of acidic phosphoproteins (28-33 kDa in size) that serve as adapters regulating a number of signaling pathways. Intriguingly, IFs, including neurofilaments, are known to recruit these proteins in a phosphorylation-dependent manner (33). For these reasons we decided to look for both TFEB and 14-3-3 localization in *GAN* silenced DRG neurons.

After co-staining TFEB and NFL, we found that TFEB was enriched in cytoplasmic neurofilament aggregates when *GAN* was silenced **(Fig 8A, B)**. Moreover, we found significantly less TFEB in the nucleus of *GAN* null neurons (∼33% less than wild type). We also found that 14-3-3 proteins co-aggregate with NFL in *GAN*-silenced DRGs **(Fig 8C)**.

**Fig. 8.**
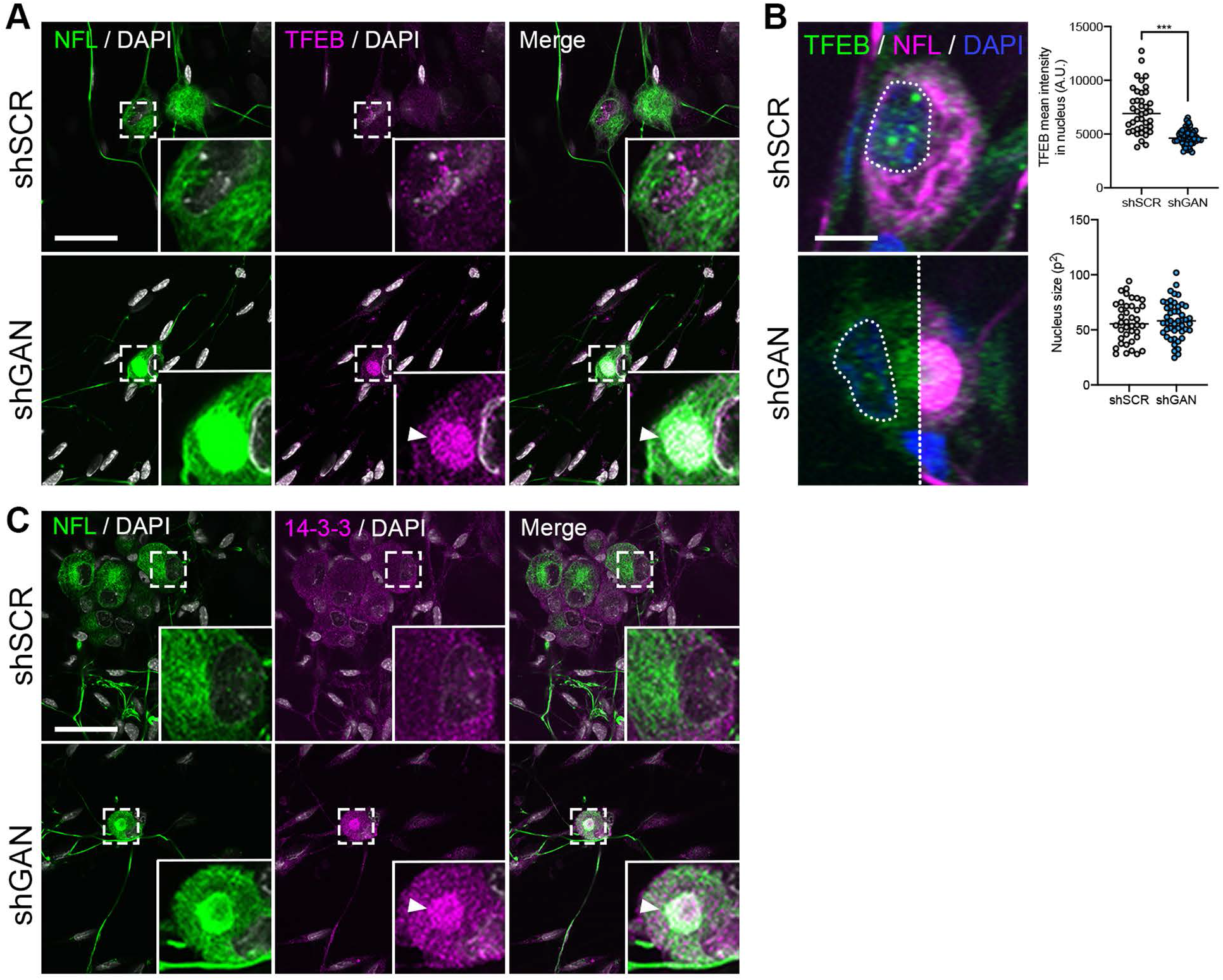
Neurofilament aggregates sequester TFEB and impairs its nuclear translocation. (A) Representative fluorescence images of control and shGAN DRGs co-stained for NFL and TFEB showing sequestering of the transcription factor (arrowhead). Scale bar, 30 µm. (B) Higher magnification pictures showing decreased TFEB localization in the nuclear compartment in shGAN DRG neurons. Accompanying plots show that TFEB has decreased intensity, and that nuclear size is similar between control and shGAN DRG neurons. (C) Representative fluorescence images of control and shGAN DRG neurons co-stained for NFL and 14-3-3. Neurofilament aggregates sequester 14-3-3 (arrowhead). Scale bar, 30 µm.

## DISCUSSION

Neurofilaments accumulate in several neurodegenerative syndromes—Alzheimer disease, Parkinson disease, polyglutamine diseases, and amyotrophic lateral sclerosis (ALS), to name just a few (12). Yet the role of neurofilament aggregation in disease has been largely overlooked in favor of disease-specific features. Rare diseases, such as GAN, provides a unique opportunity to understand the effects of neurofilament aggregation (12). Here we show that neurofilament accumulations in GAN impair autophagy by disrupting the spatial distribution and transport of autophagic organelles and the master transcription factor TFEB, which is required for lysosome biogenesis and autophagic flux.

The first hint that impaired autophagy contributes to the pathogenesis of human degenerative syndromes came from genetically engineered mice. Mice lacking ATG5 or ATG7 (autophagy-related 5 or 7, respectively) in their nervous system, for example, demonstrate progressive neurological deficits and protein aggregation. Patients have also been described with mutations in genes directly linked with autophagic processes (34, 35). These diseases are rare, but there has been a growing appreciation for the role of autophagy in the common neurodegenerative proteopathies such as Parkinson disease. In GAN, autophagy’s role is more complex: clearance of disease-specific proteins that resist ubiquitin-proteasome degradation is attempted via autophagy, while autophagy itself is compromised by a range of cellular events triggered by the misfolded proteins themselves in pathways yet to be completely elucidated (34, 36, 37). In the normal physiological state, neurofilaments undergo autophagic clearance, and the autophagic vesicles surrounding NF aggregates in GAN suggest that autophagy might even be recruited as a salvage pathway to clear neurofilaments (38). But as we show here, neurofilament accumulations themselves hinder autophagy. It is intriguing that in many of the common proteopathies, disease proteins such as alpha synuclein, mutant Huntingtin, and TDP43 are surrounded by neurofilament caps in structures called aggresomes. It is tempting to speculate that the neurofilament caps contribute to the autophagic impairments in these disorders as well. Gigaxonin has recently been shown to degrade ATG16L, a protein involved in autophagosome maturation, and thus might play an independent role as an autophagy regulator (39). In DRGs we did not observe accumulation of this protein, but we cannot exclude that such a mechanism might compound the neurofilament-induced autophagic deficits that we observe.

The mechanisms by which neurofilaments interfere with autophagy—altering the localization of autophagic vesicles and sequestering TFEB—are in essence distortions of the normal role of neurofilaments to serve as a docking platform. When aggregated, neurofilaments have very distinct biophysical properties from well-distributed wild-type polymers: they are tightly packed, display a lack of dynamic behavior, and even appear to undergo solid-to-liquid phase transitions (11, 40). Phosphorylation could further affect these properties (11). The mislocalization of autophagic organelles is reminiscent of what has been previously observed with mitochondria in GAN (6). It will now be important to determine the full complement of proteins and organelles affected by neurofilament aggregates in the disease state. Abnormal interactions are likely to be further compounded by inter-organellar co-dependence, exacerbating the pathology. For instance, mitochondrial deficits could limit cellular energy supplies, impairing lysosomal acidification, while abnormal autophagy could affect mitochondrial quality control through mitophagy. Mitochondrial and lysosomal contact sites could serve as additional points of cross-talk between these two dynamic organelles (41). Future studies will be required to tease out the complex interactions between signaling pathways, organelle dysfunction, and neurofilament aggregation to determine strategies to best treat GAN and other neurofilament proteopathies.

## MATERIAL AND METHODS

### Mice

The generation of *GAN* null mice has been previously described (15). The animals were housed in a specific pathogen free facility at Northwestern University, and experiments were performed in accordance with the National Institutes of Health’s Guide for the Care and Use of Laboratory Animals (with protocols approved by Northwestern University’s Institutional Animal Care and Use Committee).

### Dorsal root ganglia cultures

DRG neurons were isolated from post-natal day 5 mice using a published protocol with few modifications (6). Briefly, mouse DRG were dissected from their paraspinal location; they were then placed in cold dissection medium in a microcentrifuge tube (97.5% HBSS Ca^2+^ and Mg^2+^ free, 1X sodium pyruvate, 0.1% glucose, 10 mM HEPES), pelleted by a 10 s centrifugation pulse on a table-top centrifuge, and then washed in the same dissection media and finally harvested by pelleting. Washed ganglia were dissociated at 37°C for 10 min, first in 1 mL of neurobasal media (Gibco) containing 35 U papain/mL followed by a centrifugation pulse and then in 1 mL of hibernate medium containing 4 mg/mL collagenase type II and 4.6 mg/mL dispase II. This was followed by another pulsed centrifugation and wash. The resuspended cells were triturated by pipetting through a P1000 tip in plating medium (Neurobasal (Gibco) containing penicillin-streptomycin 100 U/mL-100 µg/mL and 1X GlutaMax). The dissociated cells were separated from any clumps by filtering them through a 100-micron cell strainer (Corning). They were then harvested for plating by a low-speed centrifugation for 5 min at 230 x g.

The cells were then cultured on plating medium in 35 mm microwells with a 14 mm glass bottom (MatTek). Each mouse provided sufficient DRGs to plate 5 dishes at approximately 80,000 cells per dish. After allowing the plated cells to settle, the plating media was gently aspirated and replaced with 2 mL of pre-warmed maintenance medium (plating media supplemented with 1% nerve growth factor). Primary neurons were maintained at 37 °C in a humidified 5% CO_2_ atmosphere with addition of 0.5 mL of fresh maintenance media every 3 days. On day 3 *in vitro,* DRG cultures were transduced with lentivirus encoding shRNA against gigaxonin (shGAN) or a scrambled control (shSCR) using 10 µL of concentrated lentivirus per microwell, prepared as described below.

### Lentiviral constructs

EGFP derived from pEGFP-C1 (Clontech) was cloned in-frame to the N-terminal domain of the mouse *NFL* gene (derived from pmNFL; Addgene ID 83127) using the NEBuilder HiFi DNA Assembly cloning kit (New England Biolabs E5520S). The fusion construct was then amplified by PCR and inserted into the multiple-cloning site of the lentiviral vector pLEX using the same NEBuilder cloning kit to generate pLEX-mNFL-GFP. Constructs were validated with Sanger sequencing using primers to sequence over insertion sites.

Autophagic flux was measured by an mRFP-GFP-LC3 construct as described previously (42). Lentiviruses expressing shGAN and shSCR have been described previously and were obtained from SIGMA MISSION shRNA systems: shGAN (MISSION^®^ vector TRC # TRCN0000251146, Sigma) and shSCR (#SHC002, Sigma).

### Lentivirus production and transduction

HEK293T cells were used to produce all the lentiviruses. HEK293T cells were plated in T75 flasks at a density of 60-80% in a culture medium of high-glucose DMEM (Gibco 11965092) supplemented with 10% fetal bovine serum (Gibco 16140089) and penicillin/streptomycin (100 U/mL, 100 µg/mL, Gibco 15140122). The lentiviral construct of interest was co-transfected with the packaging constructs pCMV-VSV-G and pCMV Gag/Pol at a ratio of 10 µg: 4 µg: 6 µg respectively using Lipofectamine 2000 (60 µL, Invitrogen 11668027). After 4 h, the growth media of the transfected HEK293T cells was replaced with 10 mL of fresh complete DMEM.

Media was collected 48 h after transfection and clarified by syringe filtration through a 0.45 µm polyvinylidene difluoride membrane (Millex-HV) before concentrating the virus with a Lenti-X concentrator (Clontech 631231). The concentrated virus was resuspended in PBS and aliquoted into single-use tubes stored at −80 °C. The concentrated lentivirus in a volume of 10 µL was delivered to DRG cultures cultured on glass-bottomed microwells. Each new lot of lentiviruses was first tested for at least 50% transduction efficacy in wild-type DRG cultures before experimental use (as measured by fluorescence of tagged constructs or knockdown of shRNA constructs as determined by qPCR).

### SILAC sample preparation and data analysis

DRG neurons from p3 wild-type mice were isolated and plated as previously described (6). In brief, the dissociated neurons were plated on poly-D-lysine coverslips and maintained in Neurobasal media for 13 days. Gigaxonin was silenced using lentivirus expressing shRNA to gigaxonin (shGAN) or control (scrambled sequence; shSCR) after 3 days in culture. In total, 8 dishes of DRG neurons were cultured for SILAC analysis where a standard label swapping approach was carried out. Here, two cultures treated with shSCR and two cultures treated with shGAN were incubated with media containing heavy isotope-enriched arginine and lysine (^13^C^15^N). In parallel, two cultures treated with shSCR and two cultures treated with shGAN were incubated with normal culture media. On DIV13, cells were lysed with 1% SDS containing 100 mM Tris-Cl and boiled for 10 min. Protein concentrations were determined via a bicinchoninic assay (BCA, Pierce). Equal amounts of protein (50 µg) from pairs of heavy and light cultures were mixed at a 1:1 ratio. An in-solution trypsin digestion of proteins was carried out after reduction with 5 mM dithiothreitol at 55°C for 15 min and alkylation with 15 mM iodoacetamide for 20 min at room temperature. The resulting peptides were fractionated via HyperSep Strong Cation Exchange using 25 mM, 50 mM, 500 mM, 1 M, 2 M, and 4 M KCl, and each fraction was subsequently desalted by C18 ZipTip and dried *in vacuo*. Each of the samples was resuspended in 0.1% formic acid and analyzed via nano-electrospray ionization on a Thermo Orbitrap Fusion mass spectrometer, where auto MS/MS data was acquired in positive ion mode. Here, solvent A was 0.1% formic acid whereas solvent B was acetonitrile with 0.1% formic acid. Peptides were resolved using a 60 min linear gradient where a single mass spectrometry analysis was done for each fraction. Data analysis was performed using the Thermo Proteome Discoverer software. “Light” samples were Lys0, Arg0 and “heavy” samples were Lys8, Arg10. The database search included fixed modifications, such as carbaminomidomethyl (C), and variable modifications such as oxidation (M) and deamination (N,Q). Precursor quantification of pairwise ratios (matched median peptide abundance) and t-test analysis using no missing channels were used to calculate relative quantification ratios. Significance of shGAN/shSCR ratios was determined at a threshold of p ≤ 0.05 with Benjamini-Hochberg correction applied to reduce the false discovery rate. Pathway and biofunction analysis of altered proteins were performed in Ingenuity Pathway Analysis.

### Immunocytochemistry

DRG neurons on glass-bottomed microwells were fixed in −20 °C methanol for 7 min, a method which is ideal for fixation of neurofilaments (43). After fixation the cells were incubated for 1 h with blocking solution (5% normal goat serum in PBS) and then incubated overnight at 4 °C with the relevant primary antibodies diluted in blocking solution. The microwells were then washed twice with PBS containing 0.05% Tween 20 followed by a similar wash twice with PBS (each wash 5 min). The cells were then incubated for an hour at room temperature with Alexa fluorophore-conjugated secondary antibodies and DAPI (1:500 dilution) diluted in blocking solution (see antibodies table) and washed again as described above. A drop of mounting media (Prolong Diamond; Life Technologies) was added to the microwell and a coverslip was placed.

### Live imaging of lysosomes and autophagic flux

The conditioned media of DRG cultures that would be subjected to lysosomal staining was first removed and kept aside. The cells were then treated with 50 mM lysotracker (Red DND-99; L7528 Invitrogen) either alone or in combination with 1 µM lysosensor (Green DND-189; L7535 Invitrogen) in plating media for 30 min, after which the media was replaced with the previously stored media (to avoid lysotracker cytotoxicity). When we wished to visualize neurofilaments, we also transduced the cells 3 days before treatment with pLEX-mNFL-GFP. To measure autophagic flux in living cells, we transduced DRG cultures with mRFP-GFP-LC3 lentivirus 3 days prior to imaging.

### Light microscopy

Resonant scanning confocal microscopy was performed using a Nikon A1R+ platform equipped with a 100× oil-immersion objective and PerfectFocus focal drift compensation mechanism with automated XY stage. The green fluorophores were excited using laser lines set at 488 nm with emission filters set at 525-550 nm; the red fluorophores were excited using a laser line set at 561 nm with emission filters set at 600-650 nm. The confocal pinhole size was fixed at 1.2× the size of the Airy disc of the red channel. For live cell microscopy, images were captured at a single XYZ position every second for 3 min using 8 frame averages to improve the signal-to-noise ratio. For fixed cell microscopy, images were acquired in Galvano mode. Image resolution was optimized using the Nyquist criterion by the Nikon Elements software.

### Transmission electronic microscopy

DRG cultures were fixed in 0.1 M sodium cacodylate buffer (pH 7.3) containing 2% paraformaldehyde and 2.5% glutaraldehyde. They were then processed and embedded in resin blocks which were then sectioned, stained, and imaged as previously described (6).

### Antibodies

The following primary antibodies were used: Neurofilament NFL chicken polyclonal / CPCA-NF-L / Encor LC3A/B monoclonal / #12741/ Cell Signaling Technology LAMP-1 rabbit polyclonal / ab24170 / abcam GAPDH rabbit monoclonal / #2118 / Cell Signaling Technology TFEB rabbit polyclonal / SAB2108453 / Labome TFEB rabbit polyclonal / A303-773A / Bethyl Laboratories 14-3-3 rabbit polyclonal / #51-0700 / Cell Signaling Technology Cathepsin B mouse monoclonal / ab58802 / abcam Cathepsin D mouse monoclonal / ab75852 / abcam.

### Western blotting

The DRG media of each microwell culture was aspirated and replaced with 70 µL RIPA buffer containing 1% protease inhibitor. The cells were gently scraped with the pipette tip and the lysates were collected in microcentrifuge tubes. To ensure adequate lysates for western blotting, typically 5 dishes of each experimental sample were pooled. Protein concentrations were determined using the Pierce BCA assay (Thermo Fisher Scientific). Protein extracts were prepared with Laemmli buffer, warmed at 95 °C for 5 min, and separated on gradient SDS-polyacrylamide gels by electrophoresis. The proteins were then electrophoretically transferred to nitrocellulose membranes. Membranes were blocked at room temperature for 1 h using 5% blocking solution and blotted sequentially with the appropriate primary antibody (overnight 4 °C) and the horseradish peroxidase-conjugated secondary antibodies (2 h RT). Between primary and secondary antibody incubation, membranes were washed 3×10 min with PBS with 0.1%.

Tween 20. Similarly, membranes were washed after secondary antibody incubation before protein detection using SuperSignal West Pico Chemiluminescence Detection kit (Thermo Fisher Scientific). Images were acquired with a Bio-Rad Chemidoc XRS+ gel imaging system and analyzed with ImageJ/Fiji software (National Institutes of Health). Relative protein quantification was carried out by measuring the area under the curve (AUC) normalized to values of loading controls (GAPDH) as indicated in the figure legends.

### Image analysis

All images were quantified with ImageJ/Fiji software (NIH). For particle analysis (defining lysosomal compartments and measuring within specific regions of interest (ROIs)), images were prepared by applying general background subtraction on independent channels before applying a trous wavelet transformation (Ihor Smal, A Trous Wavelet Filter) on 3 scales. The output was then converted to a binary mask and Fiji’s built-in particle analysis feature was used to define ROIs. These ROIs served as measurements of lysosome number and size within a cell and were used as bounds for measurements of lysosensor intensity within lysotracker-defined compartments (intensity was determined using the multi-measure plugin).

### Statistics

All values represent the means ± SEM of the number of replicates indicated in the figure legends. All analyses were performed using GraphPad Prism software. Student’s *t* test was used to compare two groups, ANOVA was used to compare more than two groups, and statistical significance was defined as p<0.05. See figures legends for more details.

## FUNDING

P.O. receives grant support from NINDS (1R01NS127204-01 and R01NS082351-09), the Giddan foundation, and prior seed funding from Hannah’s Hope Fund. J.S receives support from R01AG078796 and R21AG080705. Research reported in this publication was also supported by the National Institutes of Health’s National Center for Advancing Translational Sciences, Grant Number TL1TR001423 (M.R.P).

## ACKNOWLEDGMENTS

We thank Abigail Brown and Morgan Pooler for their assistance with mouse husbandry and genotyping. Morgan Pooler also assisted with preparation of samples for electron microscopy. Light and electron microscopy imaging was performed at the Northwestern University Center for Advanced Microscopy, supported by NCI CCSG P30 CA060553 awarded to the Robert H. Lurie Comprehensive Cancer Center.

## AUTHOR CONTRIBUTIONS

JMP assisted by JZ and MV performed most of the cell biological experiments; JMP wrote the first draft of the manuscript; EI, MRP and JS performed the proteomic experiments and data analysis; CP and NS helped in the conceptual analysis and writing of the manuscript; PO supervised the entire work, analyzed and interpreted the data and wrote the manuscript with input from all the authors.

## POTENTIAL CONFLICTS OF INTEREST

The authors have declared that no conflict of interest exists.

## DATA AVAILABILITY

The list of proteins identified in mass spectrometry analysis from Database search is available in Supplementary Table S1 and S2. Any other data will be made available with request in accordance with Northwestern University data sharing policy.

## Notes

### Competing Interest Statement

The authors have declared no competing interest.

