## Supplementary material for "Neurofilament accumulation disrupts autophagy in giant axonal neuropathy": Table S1

| Description | Accession | Exp. q-value | Sum PEP Score | Coverage [%] | # Peptides | # PSMs | # Unique Peptides | # AAs | MW [kDa] | calc. pI | Score Sequest HT: Sequest HT | Abundance Ratio: (shGAN) / (Scramble) | Log Fold Change: (shGAN) / (Scramble) | Abundance Ratio Adj. P-Value: (shGAN) / (Scramble) | Abundance Ratio Variability [%]: (shGAN) / (Scramble) | Abundances (Grouped): shGAN | Abundances (Grouped): Scramble | Abundances (Grouped) CV [%]: shGAN | Abundances (Grouped) CV [%]: Scramble | shGAN DRG1124 (Light) | ShGAN DRG1207 (Heavy) | shGAN DRG1204 (Light) | ShGAN DRG1211 (Heavy) | Scramble DRG1124 (Heavy) | Scramble DRG1207 (Light) | Scramble DRG1204 (Heavy) | Scramble DRG1211 (Light) | Modifications |
|---|---|---|---|---|---|---|---|---|---|---|---|---|---|---|---|---|---|---|---|---|---|---|---|---|---|---|---|---|
| (E3-independent) E2 ubiquitin-conjugating enzyme UBE2O OS=Mus musculus OX=10090 GN=Ube2o PE=1 SV=3 | Q6ZPJ3 | 0 | 21.036 | 5 | 4 | 12 | 4 | 1288 | 140.7 | 5.06 | 22.96 | 1.393 | 0.478195258 | 0.602986469 | 83.05 | 127.8 | 72.2 | 52.58 | 93.92 | Peak Found | High | Peak Found | High | Peak Found | High | Peak Found | High | |
| [F-actin]-monooxygenase MICAL1 OS=Mus musculus OX=10090 GN=Mical1 PE=1 SV=1 | Q8VDP3 | 0 | 9.315 | 4 | 3 | 6 | 3 | 1048 | 116.7 | 6.05 | 9.77 | 1.272 | 0.347098671 | 0.791619886 | 79.95 | 109.9 | 90.1 | 42.17 | 46.25 | Not Found | High | High | High | Not Found | Peak Found | Peak Found | Peak Found | |
| [F-actin]-monooxygenase MICAL3 OS=Mus musculus OX=10090 GN=Mical3 PE=1 SV=2 | Q8CJ19 | 0 | 19.83 | 2 | 3 | 35 | 3 | 1993 | 223.6 | 5.47 | 82.52 | 0.866 | -0.20756107 | 0.893249611 | 41.1 | 96.3 | 103.7 | 53.62 | 48.55 | High | High | High | High | High | High | High | High | |
| [Protein ADP-ribosylarginine] hydrolase OS=Mus musculus OX=10090 GN=Adprh PE=1 SV=1 | P54923 | 0 | 63.697 | 23 | 4 | 56 | 4 | 362 | 40 | 5.76 | 135.16 | 0.821 | -0.284545873 | 0.372107078 | 72.71 | 90 | 110 | 39.98 | 50.49 | High | High | High | High | High | High | High | High | |
| [Pyruvate dehydrogenase (acetyl-transferring)] kinase isozyme 3, mitochondrial OS=Mus musculus OX=10090 GN=Pdk3 PE=1 SV=1 | Q922H2 | 0 | 7.965 | 8 | 2 | 4 | 2 | 415 | 47.9 | 8.82 | 11.58 | 0.914 | -0.12973393 | 0.991028699 | 0 | 95.5 | 104.5 | | | Not Found | Not Found | High | Not Found | Not Found | Not Found | High | High | |
| [Pyruvate dehydrogenase [acetyl-transferring]]-phosphatase 1, mitochondrial OS=Mus musculus OX=10090 GN=Pdp1 PE=1 SV=1 | Q3UV70 | 0 | 6.447 | 5 | 2 | 3 | 2 | 538 | 61.1 | 6.67 | 5.62 | 1.229 | 0.297484916 | 0.764830559 | 76.45 | 123.1 | 76.9 | 50.54 | 76.06 | Peak Found | Peak Found | Peak Found | High | Not Found | Peak Found | High | Peak Found | |
| 1,4-alpha-glucan-branching enzyme OS=Mus musculus OX=10090 GN=Gbe1 PE=1 SV=1 | Q9D6Y9 | 0 | 13.893 | 5 | 3 | 11 | 3 | 702 | 80.3 | 6.43 | 21.54 | 1.241 | 0.311503115 | 0.877723533 | 69.18 | 81 | 119 | 124.46 | 118.76 | High | Peak Found | High | Peak Found | High | Not Found | Not Found | Peak Found | |
| 1,5-anhydro-D-fructose reductase OS=Mus musculus OX=10090 GN=Akr1e2 PE=1 SV=1 | Q9DCT1 | 0 | 10.322 | 8 | 2 | 13 | 2 | 301 | 34.4 | 7.33 | 23.56 | 0.912 | -0.13289427 | 0.976368403 | 74.67 | 95.4 | 104.6 | 55.55 | 44.62 | High | High | High | Peak Found | High | High | Peak Found | High | Acetyl [N-Term] |
| 10 kDa heat shock protein, mitochondrial OS=Mus musculus OX=10090 GN=Hspe1 PE=1 SV=2 | Q64433 | 0 | 39.601 | 25 | 2 | 110 | 2 | 102 | 11 | 8.35 | 205.01 | 1.031 | 0.044044333 | 0.950529591 | 50.81 | 93.8 | 106.2 | 68.21 | 65.02 | High | High | High | High | High | High | High | High | |
| 116 kDa U5 small nuclear ribonucleoprotein component OS=Mus musculus OX=10090 GN=Eftud2 PE=1 SV=1 | O08810 | 0 | 185.221 | 33 | 19 | 172 | 19 | 971 | 109.3 | 5 | 431.94 | 1.069 | 0.096261853 | 0.882906899 | 68.92 | 102.8 | 97.2 | 35.97 | 41.79 | High | High | High | High | High | High | High | High | |
| 14 kDa phosphohistidine phosphatase OS=Mus musculus OX=10090 GN=Phpt1 PE=1 SV=1 | Q9DAK9 | 0 | 20.242 | 22 | 3 | 91 | 3 | 124 | 14 | 5.53 | 200.91 | 0.918 | -0.123433941 | 0.893249611 | 75.91 | 109 | 91 | 57.43 | 53.82 | High | High | High | High | High | High | High | High | |
| 14-3-3 protein beta/alpha OS=Mus musculus OX=10090 GN=Ywhab PE=1 SV=3 | Q9CQV8 | 0 | 232.799 | 54 | 13 | 376 | 9 | 246 | 28.1 | 4.83 | 893.17 | 1.007 | 0.010063683 | 0.991162618 | 82.16 | 94.8 | 105.2 | 43.24 | 38.53 | High | High | High | High | High | High | High | High | |
| 14-3-3 protein epsilon OS=Mus musculus OX=10090 GN=Ywhae PE=1 SV=1 | P62259 | 0 | 274.086 | 68 | 15 | 668 | 13 | 255 | 29.2 | 4.74 | 1557.58 | 1.051 | 0.071762669 | 0.893249611 | 93.25 | 97.5 | 102.5 | 43.05 | 46.13 | High | High | High | High | High | High | High | High | Acetyl [N-Term] |
| 14-3-3 protein eta OS=Mus musculus OX=10090 GN=Ywhah PE=1 SV=2 | P68510 | 0 | 264.528 | 53 | 12 | 371 | 9 | 246 | 28.2 | 4.89 | 876.33 | 1.079 | 0.109694865 | 0.78846753 | 76.59 | 101 | 99 | 34.55 | 39.22 | High | High | High | High | High | High | High | High | |
| 14-3-3 protein gamma OS=Mus musculus OX=10090 GN=Ywhag PE=1 SV=2 | P61982 | 0 | 173.21 | 53 | 11 | 262 | 7 | 247 | 28.3 | 4.89 | 566.4 | 1.038 | 0.053806444 | 0.947492375 | 68.86 | 109.3 | 90.7 | 17.61 | 35.96 | High | High | High | High | High | High | High | High | |
| 14-3-3 protein theta OS=Mus musculus OX=10090 GN=Ywhaq PE=1 SV=1 | P68254 | 0 | 237.11 | 55 | 14 | 371 | 12 | 245 | 27.8 | 4.78 | 871.3 | 0.929 | -0.106249498 | 0.868660687 | 79.5 | 104.6 | 95.4 | 24.25 | 25.57 | High | High | High | High | High | High | High | High | |
| 14-3-3 protein zeta/delta OS=Mus musculus OX=10090 GN=Ywhaz PE=1 SV=1 | P63101 | 0 | 323.673 | 60 | 18 | 627 | 15 | 245 | 27.8 | 4.79 | 1507.53 | 0.914 | -0.12973393 | 0.805818532 | 73.85 | 97.2 | 102.8 | 18.39 | 24.76 | High | High | High | High | High | High | High | High | |
| 182 kDa tankyrase-1-binding protein OS=Mus musculus OX=10090 GN=Tnks1bp1 PE=1 SV=2 | P58871 | 0 | 153.245 | 19 | 19 | 185 | 19 | 1720 | 181.7 | 4.88 | 435.07 | 0.957 | -0.06340917 | 0.95979474 | 89.4 | 92.9 | 107.1 | 42.2 | 42.86 | High | High | High | High | High | High | High | High | |
| 1-phosphatidylinositol 4,5-bisphosphate phosphodiesterase beta-3 OS=Mus musculus OX=10090 GN=Plcb3 PE=1 SV=2 | P51432 | 0 | 57.765 | 9 | 7 | 42 | 7 | 1234 | 139.4 | 5.94 | 91.89 | 0.706 | -0.502259911 | 0.806428765 | 44.21 | 130.1 | 69.9 | 94.1 | 118.65 | High | High | High | High | High | High | High | High | |
| 1-phosphatidylinositol 4,5-bisphosphate phosphodiesterase delta-1 OS=Mus musculus OX=10090 GN=Plcd1 PE=1 SV=2 | Q8R3B1 | 0 | 138.023 | 15 | 7 | 189 | 7 | 756 | 85.8 | 6.21 | 522.95 | 1.065 | 0.09085343 | 0.857108928 | 62.63 | 100.4 | 99.6 | 25.2 | 11.52 | High | High | High | High | High | High | High | High | |
| 1-phosphatidylinositol 4,5-bisphosphate phosphodiesterase delta-3 OS=Mus musculus OX=10090 GN=Plcd3 PE=1 SV=2 | Q8K2J0 | 0 | 12.055 | 6 | 3 | 7 | 3 | 785 | 88.6 | 6.83 | 20.14 | | | | | | | | | Not Found | High | High | High | Peak Found | Not Found | Peak Found | High | |
| 1-phosphatidylinositol 4,5-bisphosphate phosphodiesterase gamma-1 OS=Mus musculus OX=10090 GN=Plcg1 PE=1 SV=2 | Q62077 | 0 | 50.254 | 9 | 7 | 31 | 7 | 1302 | 149.6 | 5.8 | 60.76 | 0.92 | -0.120294234 | 0.968806194 | 79.8 | 98.6 | 101.4 | 59.8 | 68.84 | High | High | High | High | High | High | High | High | |
| 1-phosphatidylinositol 4,5-bisphosphate phosphodiesterase gamma-2 OS=Mus musculus OX=10090 GN=Plcg2 PE=1 SV=1 | Q8CIH5 | 0 | 29.581 | 6 | 4 | 19 | 4 | 1265 | 147.5 | 6.77 | 55.88 | 1.32 | 0.40053793 | 0.859686786 | 108.77 | 113.8 | 86.2 | 76.81 | 135.65 | High | High | High | High | High | High | Peak Found | Peak Found | |
| 2',3'-cyclic-nucleotide 3'-phosphodiesterase OS=Mus musculus OX=10090 GN=Cnp PE=1 SV=3 | P16330 | 0 | 35.904 | 8 | 3 | 53 | 3 | 420 | 47.1 | 8.97 | 58.15 | 2.909 | 1.540523296 | 0.03788856 | 99.42 | 56.9 | 143.1 | 138.22 | | High | Not Found | High | Peak Found | Peak Found | Not Found | Peak Found | Peak Found | |
| 2,4-dienoyl-CoA reductase, mitochondrial OS=Mus musculus OX=10090 GN=Decr1 PE=1 SV=1 | Q9CQ62 | 0 | 108.361 | 27 | 9 | 75 | 9 | 335 | 36.2 | 8.95 | 189.84 | 0.821 | -0.284545873 | 0.540619792 | 59.42 | 99.5 | 100.5 | 78.27 | 88.85 | High | High | High | High | High | High | High | High | |
| 2',5'-phosphodiesterase 12 OS=Mus musculus OX=10090 GN=Pde12 PE=1 SV=2 | Q3TIU4 | 0 | 17.967 | 8 | 3 | 6 | 3 | 608 | 67.5 | 6.92 | 10.64 | | | | | | | | | High | High | High | Peak Found | High | High | Not Found | Not Found | |
| 26S proteasome non-ATPase regulatory subunit 1 OS=Mus musculus OX=10090 GN=Psmd1 PE=1 SV=1 | Q3TXS7 | 0 | 161.616 | 28 | 17 | 162 | 17 | 953 | 105.7 | 5.39 | 310.65 | 1.003 | 0.004321606 | 0.995586491 | 77.11 | 95.4 | 104.6 | 55.64 | 60.75 | High | High | High | High | High | High | High | High | Acetyl [N-Term] |
| 26S proteasome non-ATPase regulatory subunit 11 OS=Mus musculus OX=10090 GN=Psmd11 PE=1 SV=3 | Q8BG32 | 0 | 103.272 | 26 | 8 | 158 | 8 | 422 | 47.4 | 6.48 | 437.67 | 0.92 | -0.120294234 | 0.835500847 | 56.14 | 102.2 | 97.8 | 52.39 | 53.77 | High | High | High | High | High | High | High | High | |
| 26S proteasome non-ATPase regulatory subunit 12 OS=Mus musculus OX=10090 GN=Psmd12 PE=1 SV=4 | Q9D8W5 | 0 | 146.448 | 23 | 9 | 209 | 9 | 456 | 52.9 | 7.06 | 485.55 | 0.995 | -0.007231569 | 0.995586491 | 90.71 | 107.9 | 92.1 | 68.59 | 54.64 | High | High | High | High | High | High | High | High | |
| 26S proteasome non-ATPase regulatory subunit 13 OS=Mus musculus OX=10090 GN=Psmd13 PE=1 SV=1 | Q9WVJ2 | 0 | 39.666 | 27 | 6 | 42 | 6 | 376 | 42.8 | 5.71 | 60.48 | 0.998 | -0.002888279 | 0.995586491 | 99.71 | 110.3 | 89.7 | 71.49 | 83.01 | High | High | Peak Found | High | High | High | Peak Found | High | |
| 26S proteasome non-ATPase regulatory subunit 14 OS=Mus musculus OX=10090 GN=Psmd14 PE=1 SV=2 | O35593 | 0 | 21.067 | 26 | 3 | 13 | 3 | 310 | 34.6 | 6.52 | 35.72 | 1.329 | 0.410341105 | 0.851237473 | 87.2 | 127.2 | 72.8 | 125.39 | 125.34 | Not Found | High | High | High | Not Found | High | High | High | |
| 26S proteasome non-ATPase regulatory subunit 2 OS=Mus musculus OX=10090 GN=Psmd2 PE=1 SV=1 | Q8VDM4 | 0 | 113.827 | 18 | 10 | 46 | 10 | 908 | 100.1 | 5.17 | 139.76 | 1.048 | 0.067638717 | 0.948607132 | 72.03 | 103.3 | 96.7 | 41.95 | 43.29 | High | High | High | High | Peak Found | High | High | High | |
| 26S proteasome non-ATPase regulatory subunit 3 OS=Mus musculus OX=10090 GN=Psmd3 PE=1 SV=3 | P14685 | 0 | 110.119 | 22 | 9 | 106 | 9 | 530 | 60.7 | 8.44 | 233.3 | 1.16 | 0.214124805 | 0.38722733 | 82.89 | 108.4 | 91.6 | 29.1 | 31.6 | High | High | High | High | High | High | High | High | |
| 26S proteasome non-ATPase regulatory subunit 4 OS=Mus musculus OX=10090 GN=Psmd4 PE=1 SV=1 | O35226 | 0 | 116.629 | 28 | 7 | 142 | 2 | 376 | 40.7 | 4.79 | 357.47 | 1.014 | 0.020057652 | 0.976368403 | 85.1 | 104.2 | 95.8 | 16.91 | 20.45 | High | High | High | High | High | High | High | High | |
| 26S proteasome non-ATPase regulatory subunit 5 OS=Mus musculus OX=10090 GN=Psmd5 PE=1 SV=4 | Q8BJY1 | 0 | 98.35 | 25 | 7 | 134 | 7 | 504 | 55.9 | 5.21 | 379.02 | 0.881 | -0.182786076 | 0.756834249 | 52.43 | 91.7 | 108.3 | 30.49 | 42.83 | High | High | High | High | High | High | High | High | |
| 26S proteasome non-ATPase regulatory subunit 6 OS=Mus musculus OX=10090 GN=Psmd6 PE=1 SV=1 | Q99JI4 | 0 | 122.92 | 30 | 10 | 97 | 10 | 389 | 45.5 | 5.52 | 234 | 1.16 | 0.214124805 | 0.625673794 | 68.33 | 109.1 | 90.9 | 61.84 | 64.84 | High | High | High | High | Peak Found | High | High | High | |
| 26S proteasome non-ATPase regulatory subunit 7 OS=Mus musculus OX=10090 GN=Psmd7 PE=1 SV=2 | P26516 | 0 | 59.809 | 20 | 3 | 42 | 3 | 321 | 36.5 | 6.77 | 116.09 | 1.006 | 0.008630305 | 0.991386436 | 109.8 | 108 | 92 | 65.46 | 13.06 | Peak Found | High | High | High | Not Found | High | High | High | |
| 26S proteasome non-ATPase regulatory subunit 8 OS=Mus musculus OX=10090 GN=Psmd8 PE=1 SV=2 | Q9CX56 | 0 | 23.667 | 7 | 2 | 13 | 2 | 353 | 39.9 | 9.58 | 26.75 | 1.151 | 0.202887833 | 0.910610405 | 130.65 | 129.7 | 70.3 | 90.12 | 100.04 | Peak Found | High | Peak Found | High | Peak Found | High | Peak Found | High | |
| 26S proteasome non-ATPase regulatory subunit 9 OS=Mus musculus OX=10090 GN=Psmd9 PE=1 SV=1 | Q9CR00 | 0 | 25.248 | 13 | 2 | 35 | 2 | 222 | 24.7 | 6.43 | 89.02 | 0.823 | -0.281035664 | 0.820983453 | 82.55 | 84.9 | 115.1 | 92.26 | 80.69 | High | High | High | High | High | High | High | High | |
| 26S proteasome regulatory subunit 10B OS=Mus musculus OX=10090 GN=Psmc6 PE=1 SV=1 | P62334 | 0 | 130.885 | 45 | 12 | 192 | 12 | 389 | 44.1 | 7.49 | 497.39 | 1.015 | 0.021479727 | 0.976368403 | 62.66 | 98.6 | 101.4 | 40.51 | 36.78 | High | High | High | High | High | High | High | High | |
| 26S proteasome regulatory subunit 4 OS=Mus musculus OX=10090 GN=Psmc1 PE=1 SV=1 | P62192 | 0 | 186.127 | 26 | 10 | 194 | 10 | 440 | 49.2 | 6.21 | 564.4 | 1.085 | 0.117695043 | 0.768272619 | 75.67 | 104.5 | 95.5 | 18.12 | 15.17 | High | High | High | High | High | High | High | High | |
| 26S proteasome regulatory subunit 6A OS=Mus musculus OX=10090 GN=Psmc3 PE=1 SV=2 | O88685 | 0 | 271.192 | 52 | 17 | 331 | 17 | 442 | 49.5 | 5.19 | 850.53 | 1.01 | 0.014355293 | 0.987640341 | 92.9 | 103.6 | 96.4 | 22.06 | 33.42 | High | High | High | High | High | High | High | High | Acetyl [N-Term] |
| 26S proteasome regulatory subunit 6B OS=Mus musculus OX=10090 GN=Psmc4 PE=1 SV=2 | P54775 | 0 | 232.756 | 37 | 10 | 333 | 10 | 418 | 47.4 | 5.21 | 904.34 | 0.946 | -0.080087911 | 0.929173852 | 67.58 | 97.9 | 102.1 | 9.06 | 15.71 | High | High | High | High | High | High | High | High | Acetyl [N-Term] |
| 26S proteasome regulatory subunit 7 OS=Mus musculus OX=10090 GN=Psmc2 PE=1 SV=5 | P46471 | 0 | 154.618 | 37 | 13 | 212 | 13 | 433 | 48.6 | 5.95 | 562.22 | 1.027 | 0.038436182 | 0.95979474 | 83.28 | 101.7 | 98.3 | 47.57 | 43.5 | High | High | High | High | High | High | High | High | |
| 26S proteasome regulatory subunit 8 OS=Mus musculus OX=10090 GN=Psmc5 PE=1 SV=1 | P62196 | 0 | 177.402 | 35 | 11 | 207 | 11 | 406 | 45.6 | 7.55 | 609.1 | 0.956 | -0.064917477 | 0.95979474 | 52.42 | 100.6 | 99.4 | 33.41 | 44.18 | High | High | High | High | High | High | High | High | |
| 28 kDa heat- and acid-stable phosphoprotein OS=Mus musculus OX=10090 GN=Pdap1 PE=1 SV=1 | Q3UHX2 | 0 | 62.262 | 22 | 6 | 118 | 6 | 181 | 20.6 | 7.39 | 254.64 | 0.905 | -0.144010303 | 0.923493786 | 80.69 | 96.2 | 103.8 | 69.9 | 63.68 | High | High | High | High | High | High | High | High | |
| 28S ribosomal protein S23, mitochondrial OS=Mus musculus OX=10090 GN=Mrps23 PE=1 SV=1 | Q8VE22 | 0 | 10.062 | 20 | 2 | 5 | 2 | 177 | 20.3 | 8.59 | 7.7 | 0.851 | -0.232768963 | 0.930533245 | 42.83 | 92 | 108 | 52.74 | 25.17 | High | High | Peak Found | High | Not Found | Peak Found | Peak Found | High | |
| 28S ribosomal protein S26, mitochondrial OS=Mus musculus OX=10090 GN=Mrps26 PE=1 SV=1 | Q80ZS3 | 0 | 10.97 | 10 | 2 | 7 | 2 | 200 | 23.4 | 9.96 | 4.88 | 1.067 | 0.093560176 | 0.983876985 | 56.43 | 100.4 | 99.6 | 22.1 | 13.06 | Not Found | High | Peak Found | Peak Found | Not Found | Peak Found | Not Found | High | |
| 28S ribosomal protein S27, mitochondrial OS=Mus musculus OX=10090 GN=Mrps27 PE=1 SV=2 | Q8BK72 | 0 | 39.5 | 15 | 5 | 62 | 5 | 415 | 47.7 | 5.5 | 123.87 | 0.833 | -0.263611599 | 0.643154579 | 47.39 | 96.2 | 103.8 | 26.72 | 38.05 | High | High | High | High | High | High | High | High | |
| 28S ribosomal protein S30, mitochondrial OS=Mus musculus OX=10090 GN=Mrps30 PE=1 SV=1 | Q9D0G0 | 0 | 8.295 | 5 | 2 | 5 | 2 | 442 | 49.9 | 9.38 | 10.87 | | | | | | | | | Not Found | High | Not Found | High | Not Found | Not Found | Not Found | Not Found | |
| 28S ribosomal protein S9, mitochondrial OS=Mus musculus OX=10090 GN=Mrps9 PE=1 SV=3 | Q9D7N3 | 0 | 17.681 | 7 | 2 | 13 | 2 | 390 | 44.9 | 8.81 | 27.68 | | | | | | | | | Not Found | High | Not Found | High | High | High | Not Found | High | |
| 2-amino-3-ketobutyrate coenzyme A ligase, mitochondrial OS=Mus musculus OX=10090 GN=Gcat PE=1 SV=2 | O88986 | 0 | 81.528 | 17 | 4 | 73 | 4 | 416 | 44.9 | 7.33 | 210.88 | 0.845 | -0.242976753 | 0.858963147 | 44.52 | 95.3 | 104.7 | 40.34 | 31.54 | High | High | High | High | Not Found | High | High | High | |
| 2-oxoglutarate dehydrogenase, mitochondrial OS=Mus musculus OX=10090 GN=Ogdh PE=1 SV=3 | Q60597 | 0 | 112.757 | 19 | 14 | 129 | 14 | 1023 | 116.4 | 6.83 | 275.23 | 1.235 | 0.304511042 | 0.146471874 | 74.96 | 100.8 | 99.2 | 70.07 | 54.37 | High | High | High | High | High | High | High | High | |
| 2-oxoisovalerate dehydrogenase subunit alpha, mitochondrial OS=Mus musculus OX=10090 GN=Bckdha PE=1 SV=1 | P50136 | 0 | 20.948 | 12 | 3 | 26 | 3 | 442 | 50.3 | 8.06 | 49.99 | 1.319 | 0.399444565 | 0.70515503 | 102.42 | 119.7 | 80.3 | 117.97 | 115.13 | High | High | High | High | High | High | Peak Found | High | |
| 3'(2'),5'-bisphosphate nucleotidase 1 OS=Mus musculus OX=10090 GN=Bpnt1 PE=1 SV=2 | Q9Z0S1 | 0 | 23.82 | 20 | 4 | 8 | 4 | 308 | 33.2 | 5.82 | 20.46 | 1.218 | 0.284514133 | 0.911239848 | 133.1 | 133.2 | 66.8 | 86.06 | 73.65 | Peak Found | High | High | High | Peak Found | High | High | Peak Found | |
| 3'-5' RNA helicase YTHDC2 OS=Mus musculus OX=10090 GN=Ythdc2 PE=1 SV=1 | B2RR83 | 0.001 | 4.494 | 2 | 2 | 3 | 2 | 1445 | 161 | 8.51 | 7.87 | 2.06 | 1.042644337 | 0.162051981 | 87.49 | 134.6 | 65.4 | | 71.87 | Not Found | High | High | Peak Found | Not Found | Peak Found | Not Found | Peak Found | |
| 39S ribosomal protein L13, mitochondrial OS=Mus musculus OX=10090 GN=Mrpl13 PE=1 SV=1 | Q9D1P0 | 0 | 5.716 | 17 | 2 | 7 | 2 | 178 | 20.7 | 9.35 | 11.84 | | | | | | | | | Not Found | High | High | Peak Found | Not Found | High | Peak Found | High | |
| 39S ribosomal protein L14, mitochondrial OS=Mus musculus OX=10090 GN=Mrpl14 PE=1 SV=1 | Q9D1I6 | 0 | 16.456 | 19 | 2 | 13 | 2 | 145 | 15.9 | 10.99 | 26.03 | | | | | | | | | Not Found | High | Not Found | High | High | Not Found | Peak Found | High | |
| 39S ribosomal protein L15, mitochondrial OS=Mus musculus OX=10090 GN=Mrpl15 PE=1 SV=1 | Q9CPR5 | 0 | 22.247 | 11 | 2 | 10 | 2 | 295 | 33.5 | 10.07 | 20.41 | | | | | | | | | High | High | Peak Found | High | Not Found | High | Not Found | High | |
| 39S ribosomal protein L24, mitochondrial OS=Mus musculus OX=10090 GN=Mrpl24 PE=1 SV=1 | Q9CQ06 | 0 | 30.327 | 9 | 3 | 19 | 3 | 216 | 24.9 | 9.5 | 48.18 | 0.551 | -0.859875776 | 0.064920879 | 35.09 | 98 | 102 | 77.23 | 67.61 | Peak Found | High | Peak Found | High | Not Found | High | High | High | |
| 3-hydroxyacyl-CoA dehydrogenase type-2 OS=Mus musculus OX=10090 GN=Hsd17b10 PE=1 SV=4 | O08756 | 0 | 73.511 | 42 | 7 | 42 | 7 | 261 | 27.4 | 8.41 | 89.05 | 0.775 | -0.367731785 | 0.657543235 | 40 | 94.9 | 105.1 | 12.83 | 29.4 | High | High | High | High | High | High | Peak Found | High | |
| 3-hydroxyisobutyrate dehydrogenase, mitochondrial OS=Mus musculus OX=10090 GN=Hibadh PE=1 SV=1 | Q99L13 | 0 | 14.785 | 9 | 3 | 22 | 3 | 335 | 35.4 | 8.13 | 47.6 | 1.04 | 0.056583528 | 0.968806194 | 32.99 | 101.7 | 98.3 | 21.73 | 32.3 | High | High | High | High | High | High | High | High | |
| 3-hydroxyisobutyryl-CoA hydrolase, mitochondrial OS=Mus musculus OX=10090 GN=Hibch PE=1 SV=1 | Q8QZS1 | 0 | 36.412 | 19 | 7 | 47 | 7 | 385 | 43 | 8.06 | 101.13 | 0.971 | -0.042456799 | 0.995717381 | 59.54 | 101.6 | 98.4 | 49.94 | 62.01 | High | High | High | High | High | High | High | High | |
| 3-ketoacyl-CoA thiolase A, peroxisomal OS=Mus musculus OX=10090 GN=Acaa1a PE=1 SV=1 | Q921H8 | 0 | 143.243 | 51 | 12 | 133 | 12 | 424 | 43.9 | 8.44 | 366.81 | 0.938 | -0.092340172 | 0.932720092 | 87.98 | 104.9 | 95.1 | 40.91 | 63.22 | High | High | High | High | High | High | High | High | |
| 3-ketoacyl-CoA thiolase, mitochondrial OS=Mus musculus OX=10090 GN=Acaa2 PE=1 SV=3 | Q8BWT1 | 0 | 332.346 | 52 | 13 | 411 | 13 | 397 | 41.8 | 8.09 | 1145.15 | 0.964 | -0.052894948 | 0.968806194 | 70.43 | 99.5 | 100.5 | 20.3 | 15.95 | High | High | High | High | High | High | High | High | |
| 3-keto-steroid reductase OS=Mus musculus OX=10090 GN=Hsd17b7 PE=1 SV=1 | O88736 | 0 | 14.074 | 9 | 2 | 23 | 2 | 334 | 37.3 | 6.73 | 45.52 | 1.109 | 0.149259365 | 0.89438774 | 101.67 | 104.1 | 95.9 | 75.6 | 70.7 | High | High | High | High | High | High | High | High | |
| 3-mercaptopyruvate sulfurtransferase OS=Mus musculus OX=10090 GN=Mpst PE=1 SV=4 | Q99J99 | 0 | 45.633 | 19 | 4 | 36 | 4 | 297 | 33.1 | 6.47 | 78.12 | 1 | 0 | 0.995586491 | 101.51 | 94.6 | 105.4 | 26.63 | 17.06 | High | High | High | High | Peak Found | High | High | High | |
| 40S ribosomal protein S10 OS=Mus musculus OX=10090 GN=Rps10 PE=1 SV=1 | P63325 | 0 | 50.259 | 43 | 7 | 81 | 7 | 165 | 18.9 | 10.15 | 159.34 | 1.033 | 0.046840254 | 0.941777527 | 106.29 | 94.6 | 105.4 | 90.85 | 89.7 | High | High | High | High | High | High | High | High | |
| 40S ribosomal protein S12 OS=Mus musculus OX=10090 GN=Rps12 PE=1 SV=2 | P63323 | 0 | 75.052 | 42 | 7 | 119 | 7 | 132 | 14.5 | 7.24 | 219.18 | 0.978 | -0.03209363 | 0.995586491 | 73.48 | 101.4 | 98.6 | 38.42 | 51.79 | High | High | High | High | High | High | High | High | |
| 40S ribosomal protein S13 OS=Mus musculus OX=10090 GN=Rps13 PE=1 SV=2 | P62301 | 0 | 59.37 | 28 | 6 | 37 | 6 | 151 | 17.2 | 10.54 | 75.73 | 1.01 | 0.014355293 | 0.992199867 | 96.11 | 109.1 | 90.9 | 81.33 | 80.31 | Peak Found | High | High | High | Peak Found | High | High | High | |
| 40S ribosomal protein S14 OS=Mus musculus OX=10090 GN=Rps14 PE=1 SV=3 | P62264 | 0 | 68.209 | 15 | 3 | 89 | 3 | 151 | 16.3 | 10.05 | 206.15 | 0.925 | -0.112474729 | 0.895647774 | 47.69 | 96.8 | 103.2 | 27.32 | 44.07 | High | High | High | High | High | High | High | High | |
| 40S ribosomal protein S15 OS=Mus musculus OX=10090 GN=Rps15 PE=1 SV=2 | P62843 | 0 | 74.541 | 28 | 2 | 145 | 2 | 145 | 17 | 10.39 | 349.62 | 0.806 | -0.311148256 | 0.59241686 | 50.25 | 104 | 96 | 46.33 | 81.65 | Not Found | High | High | High | Peak Found | High | High | High | |
| 40S ribosomal protein S15a OS=Mus musculus OX=10090 GN=Rps15a PE=1 SV=2 | P62245 | 0 | 11.686 | 18 | 2 | 4 | 2 | 130 | 14.8 | 10.13 | 8.74 | 1.051 | 0.071762669 | 0.966742388 | 20.03 | 106.9 | 93.1 | 73.08 | 80.37 | Not Found | High | Peak Found | Peak Found | Not Found | High | High | Peak Found | |
| 40S ribosomal protein S16 OS=Mus musculus OX=10090 GN=Rps16 PE=1 SV=4 | P14131 | 0 | 21.788 | 14 | 3 | 34 | 3 | 146 | 16.4 | 10.21 | 26.71 | 0.713 | -0.488026018 | 0.283906223 | 54.09 | 91.1 | 108.9 | 67.32 | 37.4 | High | High | High | High | High | High | High | High | |
| 40S ribosomal protein S17 OS=Mus musculus OX=10090 GN=Rps17 PE=1 SV=2 | P63276 | 0 | 163.088 | 36 | 5 | 207 | 5 | 135 | 15.5 | 9.85 | 540.41 | 1.017 | 0.024319679 | 0.971905195 | 73.72 | 100.4 | 99.6 | 63.61 | 60.84 | High | High | High | High | High | High | High | High | |
| 40S ribosomal protein S18 OS=Mus musculus OX=10090 GN=Rps18 PE=1 SV=3 | P62270 | 0 | 35.623 | 39 | 6 | 45 | 6 | 152 | 17.7 | 10.99 | 72.87 | 0.924 | -0.114035243 | 0.955075379 | 51.7 | 98.3 | 101.7 | 49.89 | 52.06 | High | High | High | High | High | High | High | High | |
| 40S ribosomal protein S19 OS=Mus musculus OX=10090 GN=Rps19 PE=1 SV=3 | Q9CZX8 | 0 | 38.069 | 24 | 4 | 48 | 4 | 145 | 16.1 | 10.4 | 88.68 | 0.955 | -0.066427362 | 0.962674291 | 66.45 | 96.2 | 103.8 | 26.11 | 35.78 | High | High | High | High | High | High | High | High | |
| 40S ribosomal protein S2 OS=Mus musculus OX=10090 GN=Rps2 PE=1 SV=3 | P25444 | 0 | 79.649 | 24 | 7 | 180 | 7 | 293 | 31.2 | 10.24 | 429 | 1.001 | 0.001441974 | 0.995586491 | 93.08 | 92.8 | 107.2 | 19.78 | 32.63 | High | High | High | High | High | High | High | High | |
| 40S ribosomal protein S21 OS=Mus musculus OX=10090 GN=Rps21 PE=1 SV=1 | Q9CQR2 | 0 | 116.031 | 47 | 5 | 298 | 5 | 83 | 9.1 | 8.51 | 668.2 | 1.061 | 0.085424656 | 0.857108928 | 69.43 | 99.1 | 100.9 | 36.72 | 30.71 | High | High | High | High | High | High | High | High | Acetyl [N-Term] |
| 40S ribosomal protein S23 OS=Mus musculus OX=10090 GN=Rps23 PE=1 SV=3 | P62267 | 0 | 19.433 | 27 | 2 | 21 | 2 | 143 | 15.8 | 10.49 | 55 | 1.01 | 0.014355293 | 0.994561598 | 37.56 | 97.7 | 102.3 | 38.6 | 42.87 | High | High | High | High | High | High | High | High | |
| 40S ribosomal protein S24 OS=Mus musculus OX=10090 GN=Rps24 PE=1 SV=1 | P62849 | 0 | 20.781 | 20 | 2 | 39 | 2 | 133 | 15.4 | 10.78 | 84.36 | 1.07 | 0.097610797 | 0.930533245 | 88.34 | 101.4 | 98.6 | 106.51 | 106.31 | High | High | High | High | Peak Found | High | High | High | |
| 40S ribosomal protein S25 OS=Mus musculus OX=10090 GN=Rps25 PE=1 SV=1 | P62852 | 0 | 15.497 | 20 | 3 | 9 | 3 | 125 | 13.7 | 10.11 | 15.97 | 0.867 | -0.205896101 | 0.950529591 | 13.81 | 39 | 161 | 147.74 | 107.99 | High | Peak Found | High | Peak Found | Not Found | Not Found | High | Peak Found | |
| 40S ribosomal protein S28 OS=Mus musculus OX=10090 GN=Rps28 PE=1 SV=1 | P62858 | 0 | 60.106 | 61 | 6 | 105 | 6 | 69 | 7.8 | 10.7 | 219.03 | 0.936 | -0.095419565 | 0.893756428 | 88.71 | 99.4 | 100.6 | 52.43 | 50.2 | High | High | High | High | High | High | High | High | |
| 40S ribosomal protein S3 OS=Mus musculus OX=10090 GN=Rps3 PE=1 SV=1 | P62908 | 0 | 84.419 | 38 | 8 | 114 | 8 | 243 | 26.7 | 9.66 | 228.53 | 1.08 | 0.111031312 | 0.784755111 | 102.63 | 101.1 | 98.9 | 61.21 | 67.14 | High | High | High | High | High | High | High | High | |
| 40S ribosomal protein S3a OS=Mus musculus OX=10090 GN=Rps3a PE=1 SV=3 | P97351 | 0 | 84.632 | 27 | 6 | 93 | 6 | 264 | 29.9 | 9.73 | 265.02 | 0.82 | -0.286304185 | 0.212415078 | 61.17 | 102.5 | 97.5 | 81.4 | 85.56 | High | High | High | High | High | High | High | High | |
| 40S ribosomal protein S4, X isoform OS=Mus musculus OX=10090 GN=Rps4x PE=1 SV=2 | P62702 | 0 | 128.848 | 31 | 9 | 166 | 9 | 263 | 29.6 | 10.15 | 439.11 | 0.963 | -0.054392297 | 0.963070482 | 81.34 | 99.5 | 100.5 | 46.53 | 58.66 | High | High | High | High | High | High | High | High | |
| 40S ribosomal protein S5 OS=Mus musculus OX=10090 GN=Rps5 PE=1 SV=3 | P97461 | 0 | 79.886 | 28 | 5 | 107 | 5 | 204 | 22.9 | 9.72 | 324.07 | 0.898 | -0.15521265 | 0.745941542 | 69.91 | 97.6 | 102.4 | 20.15 | 19.66 | High | High | High | High | High | High | High | High | Acetyl [N-Term] |
| 40S ribosomal protein S6 OS=Mus musculus OX=10090 GN=Rps6 PE=1 SV=1 | P62754 | 0 | 93.923 | 19 | 5 | 115 | 5 | 249 | 28.7 | 10.84 | 296.68 | 1.006 | 0.008630305 | 0.991386436 | 51.96 | 98.7 | 101.3 | 18.28 | 19.59 | High | High | High | High | High | High | High | High | |
| 40S ribosomal protein S7 OS=Mus musculus OX=10090 GN=Rps7 PE=2 SV=1 | P62082 | 0 | 83.347 | 51 | 8 | 95 | 8 | 194 | 22.1 | 10.1 | 190.43 | 1.081 | 0.112366523 | 0.815985547 | 106.7 | 91.5 | 108.5 | 70.84 | 61.81 | High | High | High | High | Peak Found | High | High | High | |
| 40S ribosomal protein S8 OS=Mus musculus OX=10090 GN=Rps8 PE=1 SV=2 | P62242 | 0 | 98.455 | 47 | 8 | 151 | 8 | 208 | 24.2 | 10.32 | 420.28 | 1.043 | 0.060739158 | 0.90864342 | 74.04 | 99.3 | 100.7 | 24.68 | 23.09 | High | High | High | High | High | High | High | High | |
| 40S ribosomal protein S9 OS=Mus musculus OX=10090 GN=Rps9 PE=1 SV=3 | Q6ZWN5 | 0 | 20.965 | 11 | 2 | 15 | 2 | 194 | 22.6 | 10.65 | 20.98 | 0.979 | -0.030619235 | 0.991386436 | 126.34 | 106.5 | 93.5 | 86.85 | 85.18 | Peak Found | High | Peak Found | High | High | High | Peak Found | High | |
| 40S ribosomal protein SA OS=Mus musculus OX=10090 GN=Rpsa PE=1 SV=4 | P14206 | 0 | 93.375 | 30 | 5 | 151 | 5 | 295 | 32.8 | 4.87 | 435.83 | 0.991 | -0.013043037 | 0.999900066 | 81.37 | 99.6 | 100.4 | 41.98 | 43.32 | High | High | High | High | High | High | High | High | |

| | | | | | | | | | | | | | | | | | | | | | | | | | | | |
|---|---|---|---|---|---|---|---|---|---|---|---|---|---|---|---|---|---|---|---|---|---|---|---|---|---|---|---|
| ATP-dependent DNA/RNA helicase DHX36 OS=Mus musculus OX=10090 GN=Dhx36 PE=1 SV=2 | Q8VHK9 | 0 | 22.576 | 3 | 2 | 23 | 2 | 1001 | 113.8 | 8.29 | 66.64 | 1.03 | 0.042644337 | 0.982111074 | 57.42 | 99.4 | 100.6 | 110.61 | 115.9 | High | High | High | High | High | High | High | High |
| ATP-dependent RNA helicase DDX1 OS=Mus musculus OX=10090 GN=Ddx1 PE=1 SV=1 | Q91VR5 | 0 | 129.723 | 31 | 15 | 93 | 15 | 740 | 82.4 | 7.21 | 182.62 | 1.091 | 0.125651102 | 0.858963147 | 111.88 | 95.2 | 104.8 | 87.77 | 85.13 | High | High | High | High | High | High | High | High |
| ATP-dependent RNA helicase DDX19A OS=Mus musculus OX=10090 GN=Ddx19a PE=1 SV=2 | Q61655 | 0 | 85.786 | 24 | 10 | 74 | 10 | 478 | 53.9 | 6.67 | 168.54 | 1.047 | 0.066261442 | 0.947492375 | 46.34 | 104 | 96 | 33.09 | 42.44 | High | High | High | High | High | High | High | High |
| ATP-dependent RNA helicase DDX39A OS=Mus musculus OX=10090 GN=Ddx39a PE=1 SV=1 | Q8VDW0 | 0 | 38.584 | 16 | 5 | 25 | 3 | 427 | 49 | 5.68 | 51.87 | 1.384 | 0.468843943 | 0.764830559 | 77.01 | 116.5 | 83.5 | 54.96 | 46.29 | High | High | High | Peak Found | High | High | High | Peak Found |
| ATP-dependent RNA helicase DDX3X OS=Mus musculus OX=10090 GN=Ddx3x PE=1 SV=3 | Q62167 | 0 | 136.415 | 32 | 16 | 191 | 4 | 662 | 73.1 | 7.18 | 522.16 | 0.939 | -0.090802937 | 0.895647774 | 70.12 | 89.7 | 110.3 | 24.71 | 32.68 | Not Found | High | High | High | Not Found | High | High | High |
| ATP-dependent RNA helicase DDX3Y OS=Mus musculus OX=10090 GN=Ddx3y PE=1 SV=2 | Q62095 | 0 | 139.802 | 25 | 15 | 193 | 4 | 658 | 73.4 | 7.53 | 530.97 | 0.787 | -0.345564459 | 0.791619886 | 41.07 | 118.6 | 81.4 | 74.17 | 101.28 | Not Found | High | High | High | Not Found | High | Peak Found | High |
| ATP-dependent RNA helicase DDX42 OS=Mus musculus OX=10090 GN=Ddx42 PE=1 SV=3 | Q810A7 | 0 | 70.714 | 10 | 6 | 57 | 6 | 929 | 101.9 | 6.98 | 117.51 | 1.143 | 0.192825404 | 0.827521467 | 98.24 | 96.9 | 103.1 | 48.21 | 61.48 | High | High | High | High | Peak Found | High | High | High |
| ATP-dependent RNA helicase DDX50 OS=Mus musculus OX=10090 GN=Ddx50 PE=2 SV=1 | Q99MJ9 | 0 | 18.928 | 5 | 2 | 20 | 2 | 734 | 82.1 | 9.25 | 45.01 | 1.158 | 0.211635253 | 0.839204031 | 54.29 | 99 | 101 | 44.66 | 53.76 | High | High | High | High | Peak Found | High | Peak Found | High |
| ATP-dependent RNA helicase DHX29 OS=Mus musculus OX=10090 GN=Dhx29 PE=1 SV=1 | Q6PGC1 | 0 | 9.314 | 5 | 3 | 3 | 3 | 1365 | 153.9 | 7.94 | 5.39 | | | | | | | | | Not Found | High | High | Not Found | Not Found | Not Found | Not Found | Peak Found |
| ATP-dependent RNA helicase DHX30 OS=Mus musculus OX=10090 GN=Dhx30 PE=1 SV=1 | Q99PU8 | 0 | 37.874 | 5 | 5 | 20 | 5 | 1217 | 136.6 | 8.75 | 42.75 | 0.986 | -0.020340448 | 0.999900066 | 69.55 | 98.2 | 101.8 | 54.47 | 31.86 | High | High | High | High | Peak Found | High | High | High |
| ATP-dependent zinc metalloprotease YME1L1 OS=Mus musculus OX=10090 GN=Yme1l1 PE=1 SV=1 | O88967 | 0 | 63.087 | 8 | 3 | 50 | 3 | 715 | 80 | 8.97 | 162.71 | 0.54 | -0.888968688 | 0.020444886 | 20.32 | 56.1 | 143.9 | 66.69 | 77.72 | High | High | High | High | High | High | High | High |
| Atrial natriuretic peptide receptor 2 OS=Mus musculus OX=10090 GN=Npr2 PE=1 SV=2 | Q6VVW5 | 0.002 | 4.372 | 3 | 2 | 2 | 2 | 1047 | 117 | 6.87 | 5.03 | 1.466 | 0.551885103 | 0.759140135 | 29.95 | 118.9 | 81.1 | | 28 | Not Found | Peak Found | Not Found | High | Peak Found | High | Peak Found | Peak Found |
| Atrial natriuretic peptide receptor 3 OS=Mus musculus OX=10090 GN=Npr3 PE=1 SV=3 | P70180 | 0 | 16.283 | 4 | 2 | 26 | 2 | 536 | 59.8 | 7.09 | 46.86 | 0.757 | -0.401634795 | 0.656438059 | 69.34 | 94.3 | 105.7 | 162.15 | 128.73 | High | High | High | High | High | Peak Found | High | Peak Found |
| AT-rich interactive domain-containing protein 1A OS=Mus musculus OX=10090 GN=Arid1a PE=1 SV=1 | A2BH40 | 0 | 22.807 | 1 | 2 | 21 | 2 | 2283 | 241.9 | 6.68 | 46.79 | 4.997 | 2.321062218 | 4.30071E-06 | 115.34 | 169.1 | 30.9 | 5.49 | 112.92 | High | High | High | High | High | High | High | High |
| Attractin-like protein 1 OS=Mus musculus OX=10090 GN=Atrnl1 PE=1 SV=2 | Q6A051 | 0 | 8.018 | 2 | 2 | 2 | 2 | 1378 | 152.4 | 7.3 | 5.1 | | | | | | | | | High | Not Found | Not Found | Not Found | Not Found | High | Not Found | Not Found |
| Autophagy-related protein 16-1 OS=Mus musculus OX=10090 GN=Atg16l1 PE=1 SV=1 | Q8C0J2 | 0 | 33.757 | 16 | 4 | 19 | 4 | 607 | 68.1 | 6.38 | 60.63 | 0.743 | -0.428565884 | 0.548498005 | 65.11 | 49.3 | 150.7 | 127.51 | 68.12 | Peak Found | High | High | High | Not Found | High | High | High |
| Axin interactor, dorsalization-associated protein OS=Mus musculus OX=10090 GN=Aida PE=1 SV=1 | Q8C4Q6 | 0 | 18.631 | 27 | 4 | 15 | 4 | 305 | 34.9 | 6.74 | 44.02 | 0.832 | -0.265344567 | 0.962674291 | 83.38 | 131.8 | 68.2 | 82.67 | 126.7 | Peak Found | High | High | High | High | High | Peak Found | High |
| Baculoviral IAP repeat-containing protein 6 OS=Mus musculus OX=10090 GN=Birc6 PE=1 SV=2 | O88738 | 0 | 47.048 | 3 | 8 | 29 | 8 | 4882 | 531.8 | 6.07 | 61.42 | 0.761 | -0.394031641 | 0.792829069 | 37.05 | 100.4 | 99.6 | 64.68 | 74.31 | High | High | High | High | High | High | Peak Found | High |
| BAG family molecular chaperone regulator 3 OS=Mus musculus OX=10090 GN=Bag3 PE=1 SV=2 | Q9JLV1 | 0 | 38.025 | 7 | 4 | 88 | 4 | 577 | 61.8 | 7.27 | 228.68 | 1.001 | 0.001441974 | 0.995586491 | 73.64 | 93.7 | 106.3 | 33.21 | 40.8 | High | High | High | High | High | High | High | High |
| BAG family molecular chaperone regulator 4 OS=Mus musculus OX=10090 GN=Bag4 PE=1 SV=2 | Q8CI61 | 0 | 23.386 | 9 | 3 | 16 | 3 | 457 | 49.1 | 5.34 | 35.9 | 0.781 | -0.356605547 | 0.893249611 | 19.13 | 89.5 | 110.5 | 51.9 | 33.41 | High | High | Peak Found | High | High | High | Not Found | High |
| BAG family molecular chaperone regulator 5 OS=Mus musculus OX=10090 GN=Bag5 PE=1 SV=1 | Q8CI32 | 0 | 40.99 | 14 | 5 | 22 | 5 | 447 | 50.9 | 6.05 | 69.76 | 0.819 | -0.288064643 | 0.825631756 | 55.05 | 89.2 | 110.8 | 28.71 | 40.66 | Not Found | High | High | High | Peak Found | High | High | High |
| Band 4.1-like protein 1 OS=Mus musculus OX=10090 GN=Epb41l1 PE=1 SV=2 | Q9Z2H5 | 0 | 39.29 | 9 | 4 | 22 | 4 | 879 | 98.3 | 5.62 | 65.3 | | | | | | | | | High | High | High | High | Not Found | High | High | Peak Found |
| Band 4.1-like protein 2 OS=Mus musculus OX=10090 GN=Epb41l2 PE=1 SV=2 | O70318 | 0 | 206.359 | 31 | 19 | 213 | 19 | 988 | 109.9 | 5.43 | 498.57 | 0.952 | -0.070966521 | 0.950529591 | 80.53 | 92.4 | 107.6 | 60.54 | 59.8 | High | High | High | High | Peak Found | High | High | High |
| Barrier-to-autointegration factor OS=Mus musculus OX=10090 GN=Banf1 PE=1 SV=1 | O54962 | 0 | 15.829 | 28 | 2 | 17 | 2 | 89 | 10.1 | 6.09 | 42.96 | 0.867 | -0.205896101 | 0.792829069 | 83.1 | 118.7 | 81.3 | 57.63 | 93.73 | High | High | High | High | Peak Found | High | High | High |
| Basal cell adhesion molecule OS=Mus musculus OX=10090 GN=Bcam PE=1 SV=1 | Q9R069 | 0 | 25.277 | 5 | 2 | 24 | 2 | 622 | 67.6 | 6.25 | 72.89 | 1.85 | 0.887525271 | 0.509468536 | 0 | 129.8 | 70.2 | | | High | High | High | High | Peak Found | High | High | High |
| Basement membrane-specific heparan sulfate proteoglycan core protein OS=Mus musculus OX=10090 GN=Hspg2 PE=1 SV=1 | Q05793 | 0 | 534.216 | 23 | 49 | 618 | 49 | 3707 | 398 | 6.32 | 1202.33 | 1.038 | 0.053806444 | 0.929173852 | 77.95 | 110.5 | 89.5 | 19.99 | 18.07 | High | High | High | High | High | High | High | High |
| Basic leucine zipper and W2 domain-containing protein 1 OS=Mus musculus OX=10090 GN=Bzw1 PE=1 SV=1 | Q9CQC6 | 0 | 78.557 | 16 | 5 | 51 | 5 | 419 | 48 | 5.92 | 89.73 | 1.061 | 0.085424656 | 0.95979474 | 105.91 | 80.7 | 119.3 | 92.49 | 93.57 | Peak Found | High | High | High | Peak Found | High | Peak Found | High |
| Basic leucine zipper and W2 domain-containing protein 2 OS=Mus musculus OX=10090 GN=Bzw2 PE=1 SV=1 | Q91VK1 | 0 | 37.395 | 8 | 2 | 30 | 2 | 419 | 48 | 6.68 | 86.59 | 0.824 | -0.279283757 | 0.651382403 | 68.6 | 109.2 | 90.8 | 60.89 | 71.46 | High | High | High | High | High | High | High | High |
| Basigin OS=Mus musculus OX=10090 GN=Bsg PE=1 SV=2 | P18572 | 0 | 25.481 | 10 | 3 | 62 | 3 | 389 | 42.4 | 5.85 | 106.42 | 1.106 | 0.145351386 | 0.893249611 | 46.38 | 98.8 | 101.2 | 27.51 | 48.3 | Not Found | High | High | High | Peak Found | High | High | High |
| Bcl-2-associated transcription factor 1 OS=Mus musculus OX=10090 GN=Bclaf1 PE=1 SV=2 | Q8K019 | 0 | 18.889 | 7 | 3 | 8 | 3 | 919 | 105.9 | 9.99 | 26.92 | 0.819 | -0.288064643 | 0.858963147 | 59.22 | 97.7 | 102.3 | 111.23 | 82.5 | Peak Found | Peak Found | High | High | Peak Found | Peak Found | High | High |
| Beclin-1 OS=Mus musculus OX=10090 GN=Becn1 PE=1 SV=3 | O88597 | 0 | 11.339 | 8 | 2 | 4 | 2 | 448 | 51.6 | 4.93 | 4.24 | 0.702 | -0.510457064 | 0.745941542 | 7.12 | 82.5 | 117.5 | 16.42 | 9.54 | Not Found | Peak Found | High | High | Not Found | Not Found | Peak Found | Peak Found |
| Beta-2-syntrophin OS=Mus musculus OX=10090 GN=Sntb2 PE=1 SV=2 | Q61235 | 0 | 20.263 | 4 | 2 | 32 | 2 | 520 | 56.3 | 8.69 | 76.79 | 1.126 | 0.171206827 | 0.911276781 | 102.02 | 103.5 | 96.5 | 102.6 | 87.73 | High | High | High | High | High | High | High | High |
| Beta-actin-like protein 2 OS=Mus musculus OX=10090 GN=Actbl2 PE=1 SV=1 | Q8BFZ3 | 0 | 186.631 | 21 | 6 | 998 | 1 | 376 | 42 | 5.49 | 2370.13 | | | | | | | | | High | High | Peak Found | High | High | High | High | High |
| Beta-adrenergic receptor kinase 1 OS=Mus musculus OX=10090 GN=Grk2 PE=1 SV=2 | Q99MK8 | 0 | 27.942 | 12 | 5 | 13 | 5 | 689 | 79.6 | 7.28 | 34.76 | | | | | | | | | High | High | High | High | Not Found | High | Not Found | Not Found |
| Beta-arrestin-1 OS=Mus musculus OX=10090 GN=Arrb1 PE=1 SV=1 | Q8BWG8 | 0 | 12.676 | 11 | 2 | 6 | 2 | 418 | 46.9 | 6.28 | 18.53 | | | | | | | | | Peak Found | High | High | Not Found | Not Found | Peak Found | High | Peak Found |

[illegible]

| | | | | | | | | | |
|---|---|---|---|---|---|---|---|---|---|
| Cytoskeleton-associated protein 5 OS=Mus musculus OX=10090 GN=Ckap5 PE=1 SV=1 | A2AGT5 | 0 | 171.612 | 16 | 23 | 157 | 23 | 2032 | 225.5 |
| Cytosol aminopeptidase OS=Mus musculus OX=10090 GN=Lap3 PE=1 SV=3 | Q9CPY7 | 0 | 178.604 | 39 | 15 | 223 | 15 | 519 | 56.1 |
| Cytosolic 10-formyltetrahydrofolate dehydrogenase OS=Mus musculus OX=10090 GN=Aldh1l1 PE=1 SV=1 | Q8R0Y6 | 0 | 72.643 | 11 | 7 | 94 | 6 | 902 | 98.6 |
| Cytosolic 5'-nucleotidase 3A OS=Mus musculus OX=10090 GN=Nt5c3a PE=1 SV=4 | Q9D020 | 0 | 13.725 | 8 | 2 | 8 | 2 | 331 | 37.2 |
| Cytosolic acyl coenzyme A thioester hydrolase OS=Mus musculus OX=10090 GN=Acot7 PE=1 SV=2 | Q91V12 | 0 | 35.341 | 17 | 5 | 51 | 5 | 381 | 42.5 |
| Cytosolic Fe-S cluster assembly factor NUBP2 OS=Mus musculus OX=10090 GN=Nubp2 PE=1 SV=1 | Q9R061 | 0 | 10.365 | 10 | 2 | 7 | 2 | 275 | 29.5 |
| Cytosolic iron-sulfur assembly component 3 OS=Mus musculus OX=10090 GN=Ciao3 PE=1 SV=2 | Q7TMW6 | 0 | 12.793 | 8 | 2 | 10 | 2 | 476 | 53.1 |
| Cytosolic non-specific dipeptidase OS=Mus musculus OX=10090 GN=Cndp2 PE=1 SV=1 | Q9D1A2 | 0 | 235.138 | 41 | 17 | 338 | 17 | 475 | 52.7 |
| Cytosolic phospholipase A2 OS=Mus musculus OX=10090 GN=Pla2g4a PE=1 SV=1 | P47713 | 0 | 53.932 | 10 | 5 | 76 | 5 | 748 | 85.2 |
| Cytospin-A OS=Mus musculus OX=10090 GN=Specc1l PE=1 SV=1 | Q2KN98 | 0 | 24.394 | 4 | 3 | 10 | 3 | 1118 | 124.4 |
| Cytospin-B OS=Mus musculus OX=10090 GN=Specc1 PE=1 SV=2 | Q5SXY1 | 0 | 41.69 | 9 | 4 | 21 | 4 | 1067 | 118 |
| D-3-phosphoglycerate dehydrogenase OS=Mus musculus OX=10090 GN=Phgdh PE=1 SV=3 | Q61753 | 0 | 129.234 | 30 | 11 | 138 | 11 | 533 | 56.5 |
| DAZ-associated protein 1 OS=Mus musculus OX=10090 GN=Dazap1 PE=1 SV=2 | Q9JII5 | 0 | 36.936 | 14 | 4 | 28 | 4 | 406 | 43.2 |
| D-beta-hydroxybutyrate dehydrogenase, mitochondrial OS=Mus musculus OX=10090 GN=Bdh1 PE=1 SV=2 | Q80XN0 | 0 | 12.957 | 10 | 2 | 15 | 2 | 343 | 38.3 |
| DBIRD complex subunit ZNF326 OS=Mus musculus OX=10090 GN=Znf326 PE=1 SV=1 | O88291 | 0 | 25.993 | 9 | 3 | 38 | 3 | 580 | 65.2 |
| DCC-interacting protein 13-alpha OS=Mus musculus OX=10090 GN=Appl1 PE=1 SV=1 | Q8K3H0 | 0 | 53.961 | 20 | 8 | 38 | 8 | 707 | 79.3 |
| DCC-interacting protein 13-beta OS=Mus musculus OX=10090 GN=Appl2 PE=1 SV=1 | Q8K3G9 | 0 | 32.129 | 18 | 7 | 14 | 7 | 662 | 73.8 |
| DCN1-like protein 1 OS=Mus musculus OX=10090 GN=Dcun1d1 PE=2 SV=1 | Q9QZ73 | 0 | 32.446 | 13 | 2 | 49 | 2 | 259 | 30.1 |
| DDB1- and CUL4-associated factor 7 OS=Mus musculus OX=10090 GN=Dcaf7 PE=1 SV=1 | P61963 | 0 | 44.858 | 18 | 4 | 34 | 4 | 342 | 38.9 |
| D-dopachrome decarboxylase OS=Mus musculus OX=10090 GN=Ddt PE=1 SV=3 | O35215 | 0 | 14.37 | 39 | 3 | 10 | 3 | 118 | 13.1 |
| DDRGK domain-containing protein 1 OS=Mus musculus OX=10090 GN=Ddrgk1 PE=1 SV=2 | Q80WW9 | 0 | 38.83 | 16 | 2 | 33 | 2 | 315 | 36 |
| Deaminated glutathione amidase OS=Mus musculus OX=10090 GN=Nit1 PE=1 SV=2 | Q8VDK1 | 0 | 14.893 | 9 | 2 | 9 | 2 | 323 | 35.7 |
| Death domain-containing membrane protein NRADD OS=Mus musculus OX=10090 GN=Nradd PE=1 SV=1 | Q8CJ26 | 0 | 11.164 | 12 | 3 | 7 | 3 | 228 | 24.7 |
| Death-inducer obliterator 1 OS=Mus musculus OX=10090 GN=Dido1 PE=1 SV=4 | Q8C9B9 | 0 | 7.921 | 2 | 2 | 4 | 2 | 2256 | 247 |
| Decorin OS=Mus musculus OX=10090 GN=Dcn PE=1 SV=1 | P28654 | 0 | 10.064 | 8 | 2 | 4 | 2 | 354 | 39.8 |
| Dedicator of cytokinesis protein 1 OS=Mus musculus OX=10090 GN=Dock1 PE=1 SV=3 | Q8BUR4 | 0 | 37.291 | 6 | 7 | 25 | 7 | 1865 | 214.9 |
| Dedicator of cytokinesis protein 11 OS=Mus musculus OX=10090 GN=Dock11 PE=1 SV=1 | A2AF47 | 0 | 9.507 | 2 | 3 | 3 | 3 | 2073 | 237.6 |
| Dedicator of cytokinesis protein 2 OS=Mus musculus OX=10090 GN=Dock2 PE=1 SV=3 | Q8C3J5 | 0 | 8.032 | 2 | 3 | 5 | 3 | 1828 | 211.6 |
| Dedicator of cytokinesis protein 5 OS=Mus musculus OX=10090 GN=Dock5 PE=1 SV=2 | B2RY04 | 0 | 34.877 | 3 | 5 | 26 | 5 | 1868 | 214.3 |
| Dedicator of cytokinesis protein 7 OS=Mus musculus OX=10090 GN=Dock7 PE=1 SV=3 | Q8R1A4 | 0 | 93.462 | 8 | 14 | 82 | 13 | 2130 | 241.3 |
| Dedicator of cytokinesis protein 8 OS=Mus musculus OX=10090 GN=Dock8 PE=1 SV=4 | Q8C147 | 0 | 21.532 | 2 | 2 | 25 | 1 | 2100 | 238.8 |
| Dehydrogenase/reductase SDR family member 1 OS=Mus musculus OX=10090 GN=Dhrs1 PE=1 SV=1 | Q99L04 | 0 | 29.438 | 12 | 2 | 23 | 2 | 313 | 34 |
| Dehydrogenase/reductase SDR family member 7B OS=Mus musculus OX=10090 GN=Dhrs7b PE=1 SV=1 | Q99J47 | 0 | 20.948 | 5 | 2 | 27 | 2 | 323 | 35 |
| Delta(3,5)-Delta(2,4)-dienoyl-CoA isomerase, mitochondrial OS=Mus musculus OX=10090 GN=Ech1 PE=1 SV=1 | O35459 | 0 | 54.227 | 38 | 8 | 36 | 8 | 327 | 36.1 |
| Delta-1-pyrroline-5-carboxylate dehydrogenase, mitochondrial OS=Mus musculus OX=10090 GN=Aldh4a1 PE=1 SV=3 | Q8CHT0 | 0 | 57.576 | 15 | 5 | 61 | 5 | 562 | 61.8 |
[illegible]

| | | | | | | | | | | | | |
|---|---|---|---|---|---|---|---|---|---|---|---|---|
| Four and a half LIM domains protein 1 OS=Mus musculus OX=10090 GN=Fhl1 PE=1 SV=3 | P97447 | 0 | 39.768 | 29 | 5 | 43 | 5 | 280 | 31.9 | 8.37 | 118.66 | 0.946 |
| Four and a half LIM domains protein 2 OS=Mus musculus OX=10090 GN=Fhl2 PE=1 SV=1 | O70433 | 0 | 56.608 | 40 | 6 | 56 | 6 | 279 | 32.1 | 7.3 | 100.74 | 1.068 |
| Fragile X mental retardation syndrome-related protein 2 OS=Mus musculus OX=10090 GN=Fxr2 PE=1 SV=1 | Q9WVR4 | 0 | 42.637 | 14 | 6 | 40 | 5 | 673 | 73.7 | 6.23 | 94.39 | 0.893 |
| Fructose-bisphosphate aldolase A OS=Mus musculus OX=10090 GN=Aldoa PE=1 SV=2 | P05064 | 0 | 432.345 | 57 | 16 | 954 | 14 | 364 | 39.3 | 8.09 | 2521.58 | 1.204 |
| Fructose-bisphosphate aldolase C OS=Mus musculus OX=10090 GN=Aldoc PE=1 SV=4 | P05063 | 0 | 111.934 | 34 | 9 | 115 | 7 | 363 | 39.4 | 7.12 | 336.2 | 1.273 |
| Fucose mutarotase OS=Mus musculus OX=10090 GN=Fuom PE=1 SV=1 | Q8R2K1 | 0 | 9.777 | 18 | 2 | 3 | 2 | 153 | 16.8 | 5.36 | 6.51 | |
| Fumarate hydratase, mitochondrial OS=Mus musculus OX=10090 GN=Fh PE=1 SV=3 | P97807 | 0 | 118.616 | 30 | 8 | 97 | 8 | 507 | 54.3 | 9.04 | 256.15 | 0.898 |
| Fumarylacetoacetase OS=Mus musculus OX=10090 GN=Fah PE=1 SV=2 | P35505 | 0 | 7.42 | 7 | 2 | 4 | 2 | 419 | 46.1 | 7.18 | 8.5 | 0.857 |
| FYN-binding protein 1 OS=Mus musculus OX=10090 GN=Fyb1 PE=1 SV=2 | O35601 | 0 | 18.448 | 6 | 3 | 14 | 3 | 819 | 90 | 6.98 | 24.84 | 1.274 |
| FYVE and coiled-coil domain-containing protein 1 OS=Mus musculus OX=10090 GN=Fyco1 PE=1 SV=1 | Q8VDC1 | 0 | 70.655 | 6 | 6 | 55 | 6 | 1437 | 162.2 | 4.98 | 197.71 | 1.535 |
| G1/S-specific cyclin-D1 OS=Mus musculus OX=10090 GN=Ccnd1 PE=1 SV=1 | P25322 | 0 | 36.964 | 13 | 2 | 32 | 2 | 295 | 33.4 | 5.11 | 82.43 | 0.877 |
| G1/S-specific cyclin-D3 OS=Mus musculus OX=10090 GN=Ccnd3 PE=1 SV=1 | P30282 | 0 | 7.611 | 11 | 2 | 4 | 2 | 292 | 32.4 | 6.52 | 8.26 | |
| Galactocerebrosidase OS=Mus musculus OX=10090 GN=Galc PE=1 SV=2 | P54818 | 0 | 7.009 | 10 | 3 | 4 | 3 | 684 | 77.2 | 6.74 | 4.71 | 1.451 |
| Galactokinase OS=Mus musculus OX=10090 GN=Galk1 PE=1 SV=2 | Q9R0N0 | 0 | 57.264 | 13 | 5 | 54 | 5 | 392 | 42.3 | 5.26 | 143.21 | 1.359 |
| Galectin-1 OS=Mus musculus OX=10090 GN=Lgals1 PE=1 SV=3 | P16045 | 0 | 239.685 | 50 | 8 | 880 | 8 | 135 | 14.9 | 5.49 | 2068.32 | 0.959 |
| Galectin-3-binding protein OS=Mus musculus OX=10090 GN=Lgals3bp PE=1 SV=1 | Q07797 | 0 | 174.631 | 25 | 9 | 216 | 9 | 577 | 64.5 | 5.14 | 568.06 | 0.826 |
| Gamma-adducin OS=Mus musculus OX=10090 GN=Add3 PE=1 SV=2 | Q9QYB5 | 0 | 132.445 | 29 | 16 | 133 | 16 | 706 | 78.7 | 5.95 | 357.82 | 0.871 |
| Gamma-enolase OS=Mus musculus OX=10090 GN=Eno2 PE=1 SV=2 | P17183 | 0 | 169.337 | 30 | 7 | 435 | 4 | 434 | 47.3 | 5.11 | 971.69 | 0.657 |
| Gamma-glutamyl hydrolase OS=Mus musculus OX=10090 GN=Ggh PE=1 SV=2 | Q9Z0L8 | 0 | 10.332 | 10 | 2 | 4 | 2 | 317 | 35.4 | 7.5 | 7.75 | 1.216 |
| Gamma-glutamylaminecyclotransferase OS=Mus musculus OX=10090 GN=Ggact PE=1 SV=1 | Q923B0 | 0 | 8.362 | 21 | 2 | 4 | 2 | 149 | 17.1 | 5.29 | 4.52 | 0.909 |
| Gamma-interferon-inducible lysosomal thiol reductase OS=Mus musculus OX=10090 GN=Ifi30 PE=1 SV=3 | Q9ESY9 | 0 | 64.726 | 19 | 5 | 58 | 5 | 248 | 27.8 | 5.01 | 153.36 | 1.138 |
| Gamma-soluble NSF attachment protein OS=Mus musculus OX=10090 GN=Napg PE=1 SV=1 | Q9CWZ7 | 0 | 58.784 | 20 | 4 | 72 | 4 | 312 | 34.7 | 5.41 | 214.99 | 0.932 |
| Gamma-synuclein OS=Mus musculus OX=10090 GN=Sncg PE=1 SV=1 | Q9Z0F7 | 0 | 89.113 | 31 | 3 | 135 | 3 | 123 | 13.2 | 4.65 | 357.32 | 1.209 |
| Gamma-tubulin complex component 2 OS=Mus musculus OX=10090 GN=Tubgcp2 PE=1 SV=2 | Q921G8 | 0 | 19.892 | 5 | 3 | 14 | 3 | 905 | 103.2 | 6.77 | 23.58 | 1.058 |
| Gamma-tubulin complex component 3 OS=Mus musculus OX=10090 GN=Tubgcp3 PE=1 SV=2 | P58854 | 0 | 57.027 | 9 | 5 | 43 | 5 | 905 | 103.4 | 8.32 | 112.4 | 0.866 |
| Ganglioside GM2 activator OS=Mus musculus OX=10090 GN=Gm2a PE=1 SV=2 | Q60648 | 0 | 44.586 | 20 | 2 | 46 | 2 | 193 | 20.8 | 5.9 | 150.67 | 0.872 |
| Ganglioside-induced differentiation-associated protein 1-like 1 OS=Mus musculus OX=10090 GN=Gdap1l1 PE=1 SV=1 | Q8VE33 | 0 | 16.628 | 9 | 2 | 18 | 2 | 370 | 42.3 | 6.6 | 41.33 | 1.796 |
| Gap junction alpha-1 protein OS=Mus musculus OX=10090 GN=Gja1 PE=1 SV=2 | P23242 | 0 | 7.258 | 10 | 2 | 5 | 2 | 382 | 43 | 8.76 | 5.76 | 1.304 |
| Gasdermin-D OS=Mus musculus OX=10090 GN=Gsdmdc1 PE=1 SV=1 | Q9D8T2 | 0 | 16.793 | 10 | 3 | 18 | 3 | 487 | 53.2 | 5.1 | 47.6 | 0.852 |
| GATOR complex protein MIOS OS=Mus musculus OX=10090 GN=Mios PE=1 SV=2 | Q8VE19 | 0 | 30.813 | 9 | 5 | 19 | 5 | 875 | 98.3 | 6.71 | 48.32 | 0.535 |
| GDH/6PGL endoplasmic bifunctional protein OS=Mus musculus OX=10090 GN=H6pd PE=1 SV=2 | Q8CFX1 | 0 | 115.008 | 14 | 9 | 165 | 9 | 789 | 88.9 | 6.93 | 443.44 | 1.262 |
| GDP-fucose protein O-fucosyltransferase 1 OS=Mus musculus OX=10090 GN=Pofut1 PE=1 SV=1 | Q91ZW2 | 0 | 26.989 | 19 | 5 | 18 | 5 | 393 | 44.7 | 8.41 | 36.51 | 0.82 |
| GDP-fucose protein O-fucosyltransferase 2 OS=Mus musculus OX=10090 GN=Pofut2 PE=1 SV=1 | Q8VHI3 | 0 | 10.467 | 6 | 2 | 5 | 2 | 429 | 49.4 | 6.48 | 10.7 | 1.359 |
| GDP-L-fucose synthase OS=Mus musculus OX=10090 GN=Tsta3 PE=1 SV=3 | P23591 | 0 | 44.607 | 15 | 3 | 60 | 3 | 321 | 35.9 | 6.74 | 136.09 | 1.101 |
| GDP-Man:Man(3)GlcNAc(2)-PP-Dol alpha-1,2-mannosyltransferase OS=Mus musculus OX=10090 GN=Alg11 PE=2 SV=1 | Q3TZM9 | 0 | 6.67 | 6 | 2 | 2 | 2 | 492 | 55.2 | 8.29 | 3.7 | |
| GDP-mannose 4,6 dehydratase OS=Mus musculus OX=10090 GN=Gmds PE=1 SV=1 | Q8K0C9 | 0 | 76.544 | 28 | 7 | 82 | 7 | 372 | 42 | 7.03 | 188.02 | 0.911 |
| Gelsolin OS=Mus musculus OX=10090 GN=Gsn PE=1 SV=3 | P13020 | 0 | 262.486 | 26 | 11 | 369 | 11 | 780 | 85.9 | 6.18 | 988.47 | 0.987 |
| Gem-associated protein 5 OS=Mus musculus OX=10090 GN=Gemin5 PE=1 SV=2 | Q8BX17 | 0 | 21.454 | 2 | 2 | 21 | 2 | 1502 | 166.5 | 6.71 | 60.9 | 1.084 |
| GEM-interacting protein OS=Mus musculus OX=10090 GN=Gmip PE=1 SV=1 | Q6PGG2 | 0 | 13.191 | 4 | 2 | 4 | 2 | 971 | 107.5 | 5.77 | 11.9 | |
| General transcription and DNA repair factor IIH helicase subunit XPB OS=Mus musculus OX=10090 GN=Ercc3 PE=2 SV=1 | P49135 | 0 | 31.793 | 5 | 2 | 32 | 2 | 783 | 89.1 | 7.18 | 85.44 | 1.153 |
| General transcription factor 3C polypeptide 1 OS=Mus musculus OX=10090 GN=Gtf3c1 PE=1 SV=2 | Q8K284 | 0 | 17.617 | 1 | 2 | 15 | 2 | 2101 | 237.3 | 7.25 | 40.37 | 0.964 |
| General transcription factor IIE subunit 1 OS=Mus musculus OX=10090 GN=Gtf2e1 PE=1 SV=1 | Q9D0D5 | 0 | 20.191 | 9 | 2 | 11 | 2 | 440 | 49.6 | 4.86 | 37.58 | |
| General transcription factor II-I OS=Mus musculus OX=10090 GN=Gtf2i PE=1 SV=3 | Q9ESZ8 | 0 | 18.38 | 5 | 3 | 7 | 3 | 998 | 112.2 | 6.55 | 18.79 | 1.034 |
| General vesicular transport factor p115 OS=Mus musculus OX=10090 GN=Uso1 PE=1 SV=2 | Q9Z1Z0 | 0 | 145.25 | 19 | 13 | 89 | 13 | 959 | 106.9 | 4.93 | 219.12 | 0.97 |
| Gephyrin OS=Mus musculus OX=10090 GN=Gphn PE=1 SV=2 | Q8BUV3 | 0 | 25.411 | 11 | 4 | 22 | 4 | 769 | 83.2 | 5.6 | 62.11 | 1.124 |
| Geranylgeranyl transferase type-2 subunit alpha OS=Mus musculus OX=10090 GN=Rabggta PE=1 SV=1 | Q9JHK4 | 0 | 6.703 | 4 | 2 | 9 | 2 | 567 | 64.9 | 5.8 | 21.19 | 1.15 |
| Geranylgeranyl transferase type-2 subunit beta OS=Mus musculus OX=10090 GN=Rabggtb PE=1 SV=2 | P53612 | 0 | 8.494 | 9 | 2 | 7 | 2 | 339 | 37.8 | 5.16 | 8.24 | 1.101 |
| GH3 domain-containing protein OS=Mus musculus OX=10090 GN=Ghdc PE=2 SV=2 | Q99J23 | 0 | 4.729 | 7 | 2 | 2 | 2 | 532 | 58.5 | 7.56 | 2.08 | 1.201 |
| Girdin OS=Mus musculus OX=10090 GN=Ccdc88a PE=1 SV=2 | Q5SNZ0 | 0 | 15.546 | 3 | 3 | 8 | 3 | 1873 | 215.8 | 6.24 | 13.04 | 0.852 |
| Glia maturation factor beta OS=Mus musculus OX=10090 GN=Gmfb PE=1 SV=3 | Q9CQI3 | 0 | 33.1 | 25 | 4 | 56 | 4 | 142 | 16.7 | 5.16 | 152.78 | 0.989 |
| Glia maturation factor gamma OS=Mus musculus OX=10090 GN=Gmfg PE=1 SV=1 | Q9ERL7 | 0 | 24.429 | 20 | 2 | 25 | 2 | 142 | 16.7 | 5.71 | 80.01 | 1.903 |
| Glia-derived nexin OS=Mus musculus OX=10090 GN=Serpine2 PE=1 SV=2 | Q07235 | 0 | 10.967 | 7 | 3 | 4 | 3 | 397 | 44.2 | 9.85 | 9.7 | 0.692 |
| Glial fibrillary acidic protein OS=Mus musculus OX=10090 GN=Gfap PE=1 SV=4 | P03995 | 0 | 214.227 | 45 | 20 | 221 | 19 | 430 | 49.9 | 5.34 | 620.2 | 0.874 |
| Glucocorticoid receptor OS=Mus musculus OX=10090 GN=Nr3c1 PE=1 SV=1 | P06537 | 0 | 5.722 | 5 | 2 | 5 | 2 | 783 | 86 | 6.35 | 5.37 | 0.917 |
| Glucosamine 6-phosphate N-acetyltransferase OS=Mus musculus OX=10090 GN=Gnpnat1 PE=1 SV=1 | Q9JK38 | 0 | 41.597 | 32 | 5 | 30 | 5 | 184 | 20.8 | 7.69 | 73.41 | 1.018 |
| Glucosamine-6-phosphate isomerase 1 OS=Mus musculus OX=10090 GN=Gnpda1 PE=1 SV=3 | O88958 | 0 | 39.631 | 23 | 4 | 46 | 3 | 289 | 32.5 | 6.6 | 107.62 | 0.937 |
| Glucosamine-6-phosphate isomerase 2 OS=Mus musculus OX=10090 GN=Gnpda2 PE=1 SV=1 | Q9CRC9 | 0 | 20.259 | 25 | 4 | 35 | 3 | 276 | 31.1 | 6.9 | 88.63 | 0.655 |
| Glucose-6-phosphate 1-dehydrogenase X OS=Mus musculus OX=10090 GN=G6pdx PE=1 SV=3 | Q00612 | 0 | 37.586 | 18 | 6 | 41 | 6 | 515 | 59.2 | 6.49 | 94.56 | 1.139 |
| Glucose-6-phosphate isomerase OS=Mus musculus OX=10090 GN=Gpi PE=1 SV=4 | P06745 | 0 | 255.985 | 25 | 12 | 309 | 12 | 558 | 62.7 | 8.13 | 773.89 | 1.178 |
| Glucose-induced degradation protein 8 homolog OS=Mus musculus OX=10090 GN=Gid8 PE=1 SV=1 | Q9D7M1 | 0 | 32.727 | 33 | 5 | 21 | 5 | 228 | 26.8 | 4.97 | 42.18 | 1.035 |
| Glutamate dehydrogenase 1, mitochondrial OS=Mus musculus OX=10090 GN=Glud1 PE=1 SV=1 | P26443 | 0 | 191.999 | 38 | 17 | 290 | 17 | 558 | 61.3 | 8 | 709.2 | 0.965 |
| Glutamate receptor 2 OS=Mus musculus OX=10090 GN=Gria2 PE=1 SV=3 | P23819 | 0 | 21.905 | 7 | 4 | 13 | 4 | 883 | 98.6 | 7.39 | 10.8 | |
| Glutamate--cysteine ligase regulatory subunit OS=Mus musculus OX=10090 GN=Gclm PE=1 SV=1 | O09172 | 0 | 15.63 | 11 | 2 | 11 | 2 | 274 | 30.5 | 5.52 | 38.67 | 0.921 |
| Glutamate-rich WD repeat-containing protein 1 OS=Mus musculus OX=10090 GN=Grwd1 PE=1 SV=2 | Q810D6 | 0 | 9.518 | 8 | 2 | 3 | 2 | 446 | 49.2 | 4.81 | 6.63 | |
| Glutaminase kidney isoform, mitochondrial OS=Mus musculus OX=10090 GN=Gls PE=1 SV=1 | D3Z7P3 | 0 | 79.144 | 15 | 7 | 76 | 7 | 674 | 73.9 | 7.99 | 187.18 | 0.766 |
| Glutamine amidotransferase-like class 1 domain-containing protein 1 OS=Mus musculus OX=10090 GN=Gatd1 PE=1 SV=1 | Q8BFQ8 | 0 | 13.807 | 14 | 2 | 10 | 2 | 220 | 23.3 | 7.05 | 19 | 0.776 |
| Glutamine synthetase OS=Mus musculus OX=10090 GN=Glul PE=1 SV=6 | P15105 | 0 | 42.471 | 18 | 4 | 55 | 4 | 373 | 42.1 | 7.08 | 155.77 | 1.056 |
| Glutamine--fructose-6-phosphate aminotransferase [isomerizing] 1 OS=Mus musculus OX=10090 GN=Gfpt1 PE=1 SV=3 | P47856 | 0 | 103.87 | 22 | 9 | 129 | 8 | 697 | 78.5 | 6.84 | 319.64 | 1.036 |
| Glutamine--fructose-6-phosphate aminotransferase [isomerizing] 2 OS=Mus musculus OX=10090 GN=Gfpt2 PE=1 SV=3 | Q9Z2Z9 | 0 | 73.419 | 15 | 7 | 83 | 6 | 682 | 77 | 7.15 | 215.52 | 0.982 |
| Glutamine--tRNA ligase OS=Mus musculus OX=10090 GN=Qars PE=1 SV=1 | Q8BML9 | 0 | 89.891 | 15 | 9 | 83 | 9 | 775 | 87.6 | 7.31 | 239.23 | 1.058 |
| Glutamyl aminopeptidase OS=Mus musculus OX=10090 GN=Enpep PE=1 SV=1 | P16406 | 0 | 50.966 | 10 | 6 | 57 | 6 | 945 | 107.9 | 5.44 | 116.17 | 0.59 |
| Glutaredoxin-1 OS=Mus musculus OX=10090 GN=Glrx PE=1 SV=3 | Q9QUH0 | 0 | 44.681 | 31 | 3 | 47 | 3 | 107 | 11.9 | 8.37 | 153.4 | 1.746 |
| Glutaredoxin-3 OS=Mus musculus OX=10090 GN=Glrx3 PE=1 SV=1 | Q9CQM9 | 0 | 46.494 | 24 | 6 | 45 | 6 | 337 | 37.8 | 5.59 | 108.15 | 1.016 |
| Glutaredoxin-related protein 5, mitochondrial OS=Mus musculus OX=10090 GN=Glrx5 PE=1 SV=2 | Q80Y14 | 0 | 28.532 | 22 | 2 | 47 | 2 | 152 | 16.3 | 6.55 | 119.08 | 1.042 |
| Glutaryl-CoA dehydrogenase, mitochondrial OS=Mus musculus OX=10090 GN=Gcdh PE=1 SV=2 | Q60759 | 0 | 11.502 | 13 | 3 | 4 | 3 | 438 | 48.6 | 8.73 | 4.89 | 1.147 |
| Glutathione hydrolase 7 OS=Mus musculus OX=10090 GN=Ggt7 PE=1 SV=2 | Q99JP7 | 0 | 24.833 | 10 | 4 | 12 | 4 | 662 | 70.2 | 5.06 | 26.18 | 0.536 |
| Glutathione peroxidase 1 OS=Mus musculus OX=10090 GN=Gpx1 PE=1 SV=2 | P11352 | 0 | 41.456 | 32 | 5 | 67 | 5 | 201 | 22.3 | 7.21 | 151.97 | 0.749 |
| Glutathione peroxidase 7 OS=Mus musculus OX=10090 GN=Gpx7 PE=1 SV=1 | Q99LJ6 | 0 | 16.08 | 20 | 4 | 22 | 4 | 186 | 21 | 8.27 | 38.09 | 0.754 |
| Glutathione reductase, mitochondrial OS=Mus musculus OX=10090 GN=Gsr PE=1 SV=3 | P47791 | 0 | 63.632 | 13 | 3 | 39 | 3 | 500 | 53.6 | 7.99 | 107.86 | 0.967 |
| Glutathione S-transferase A4 OS=Mus musculus OX=10090 GN=Gsta4 PE=1 SV=3 | P24472 | 0.001 | 4.648 | 5 | 2 | 2 | 2 | 222 | 25.5 | 7.39 | 1.88 | 7.244 |
| Glutathione S-transferase kappa 1 OS=Mus musculus OX=10090 GN=Gstk1 PE=1 SV=3 | Q9DCM2 | 0 | 19.29 | 14 | 2 | 15 | 2 | 226 | 25.7 | 8.88 | 32.43 | 0.838 |
| Glutathione S-transferase Mu 1 OS=Mus musculus OX=10090 GN=Gstm1 PE=1 SV=2 | P10649 | 0 | 72.979 | 43 | 8 | 56 | 6 | 218 | 26 | 7.94 | 120.55 | 0.896 |
| Glutathione S-transferase Mu 2 OS=Mus musculus OX=10090 GN=Gstm2 PE=1 SV=2 | P15626 | 0 | 77.351 | 49 | 9 | 87 | 8 | 218 | 25.7 | 7.39 | 195.01 | 0.9 |
| Glutathione S-transferase Mu 3 OS=Mus musculus OX=10090 GN=Gstm3 PE=1 SV=2 | P19639 | 0 | 29.742 | 13 | 2 | 21 | 1 | 218 | 25.7 | 7.77 | 38.58 | |
| Glutathione S-transferase Mu 5 OS=Mus musculus OX=10090 GN=Gstm5 PE=1 SV=1 | P48774 | 0 | 44.169 | 16 | 4 | 48 | 4 | 224 | 26.6 | 7.21 | 125.87 | 0.831 |
| Glutathione S-transferase omega-1 OS=Mus musculus OX=10090 GN=Gsto1 PE=1 SV=2 | O09131 | 0 | 20.232 | 14 | 2 | 20 | 2 | 240 | 27.5 | 7.36 | 56.78 | 0.971 |
| Glutathione S-transferase P 1 OS=Mus musculus OX=10090 GN=Gstp1 PE=1 SV=2 | P19157 | 0 | 238.12 | 34 | 7 | 395 | 7 | 210 | 23.6 | 7.87 | 1037.81 | 0.899 |
| Glutathione S-transferase theta-3 OS=Mus musculus OX=10090 GN=Gstt3 PE=1 SV=1 | Q99L20 | 0 | 38.322 | 21 | 4 | 17 | 4 | 241 | 27.4 | 7.74 | 51.09 | 0.672 |
| Glutathione synthetase OS=Mus musculus OX=10090 GN=Gss PE=1 SV=1 | P51855 | 0 | 51.155 | 20 | 6 | 38 | 6 | 474 | 52.2 | 5.8 | 94.86 | 0.807 |
| Glyceraldehyde-3-phosphate dehydrogenase OS=Mus musculus OX=10090 GN=Gapdh PE=1 SV=2 | P16858 | 0 | 380.688 | 69 | 20 | 5424 | 20 | 333 | 35.8 | 8.25 | 11458.29 | 1.141 |
| Glycerol-3-phosphate dehydrogenase [NAD(+)], cytoplasmic OS=Mus musculus OX=10090 GN=Gpd1 PE=1 SV=3 | P13707 | 0 | 10.463 | 14 | 3 | 6 | 3 | 349 | 37.5 | 7.17 | 11.94 | 0.103 |
| Glycerol-3-phosphate dehydrogenase 1-like protein OS=Mus musculus OX=10090 GN=Gpd1l PE=1 SV=2 | Q3ULJ0 | 0 | 36.488 | 14 | 3 | 28 | 3 | 351 | 38.2 | 6.77 | 62.84 | 0.796 |
| Glycerol-3-phosphate dehydrogenase, mitochondrial OS=Mus musculus OX=10090 GN=Gpd2 PE=1 SV=2 | Q64521 | 0 | 69.114 | 19 | 8 | 51 | 8 | 727 | 80.9 | 6.61 | 119.39 | 0.981 |
| Glycerol-3-phosphate phosphatase OS=Mus musculus OX=10090 GN=Pgp PE=1 SV=1 | Q8CHP8 | 0 | 49.865 | 34 | 7 | 41 | 7 | 321 | 34.5 | 5.35 | 107.56 | 1.14 |
| Glycine amidinotransferase, mitochondrial OS=Mus musculus OX=10090 GN=Gatm PE=1 SV=1 | Q9D964 | 0 | 62.368 | 17 | 4 | 29 | 4 | 423 | 48.3 | 7.88 | 90.34 | 1.155 |
| Glycine--tRNA ligase OS=Mus musculus OX=10090 GN=Gars PE=1 SV=1 | Q9CZD3 | 0 | 107.121 | 17 | 8 | 148 | 8 | 729 | 81.8 | 6.65 | 409.81 | 1.04 |
| Glycogen [starch] synthase, muscle OS=Mus musculus OX=10090 GN=Gys1 PE=1 SV=2 | Q9Z1E4 | 0 | 43.332 | 9 | 6 | 44 | 6 | 738 | 83.9 | 6.11 | 90.69 | 1.579 |
| Glycogen phosphorylase, brain form OS=Mus musculus OX=10090 GN=Pygb PE=1 SV=3 | Q8CI94 | 0 | 433.416 | 44 | 30 | 464 | 24 | 843 | 96.7 | 6.73 | 1142.51 | 0.962 |
| Glycogen phosphorylase, liver form OS=Mus musculus OX=10090 GN=Pygl PE=1 SV=4 | Q9ET01 | 0 | 122.773 | 19 | 14 | 126 | 10 | 850 | 97.4 | 7.09 | 269.31 | 1.501 |
| [illegible] | [illegible] | [illegible] | [illegible] | [illegible] | [illegible] | [illegible] | [illegible] | [illegible] | [illegible] | [illegible] | [illegible] | [illegible] |

| | | | | | | | | | | | | |
|---|---|---|---|---|---|---|---|---|---|---|---|---|
| Isoform 3 of G-protein-signaling modulator 1 OS=Mus musculus OX=10090 GN=Gpsm1 | Q6IR34-3 | 0 | 44.695 | 15 | 6 | 37 | 6 | 650 | 71.9 | 6.15 | 94.29 | 1.061 |
| Isoform 3 of Kinesin-like protein KIF1B OS=Mus musculus OX=10090 GN=Kif1b | Q60575-3 | 0 | 40.412 | 9 | 7 | 23 | 1 | 1150 | 130 | 8.34 | 60.5 | 1.241 |
| Isoform 3 of Microtubule-actin cross-linking factor 1 OS=Mus musculus OX=10090 GN=Macf1 | Q9QXZ0-3 | 0 | 459.654 | 15 | 51 | 370 | 1 | 5270 | 601.4 | 5.45 | 947.99 | 0.917 |
| Isoform 3 of Microtubule-associated protein 6 OS=Mus musculus OX=10090 GN=Map6 | Q7TSJ2-3 | 0 | 7.225 | 12 | 2 | 3 | 2 | 306 | 32.8 | 9.48 | 2.44 | |
| Isoform 3 of PDZ and LIM domain protein 5 OS=Mus musculus OX=10090 GN=Pdlim5 | Q8CI51-3 | 0 | 64.218 | 40 | 6 | 61 | 1 | 239 | 26.3 | 9.67 | 120.91 | 0.57 |
| Isoform 3 of Serine protease HTRA3 OS=Mus musculus OX=10090 GN=Htra3 | Q9D236-1 | 0 | 26.894 | 10 | 3 | 15 | 3 | 459 | 49.3 | 6.98 | 50.71 | 0.951 |
| Isoform 3 of YTH domain-containing family protein 3 OS=Mus musculus OX=10090 GN=Ythdf3 | Q8BYK6-3 | 0 | 23.132 | 6 | 2 | 14 | 2 | 596 | 65.4 | 9.23 | 17.76 | 0.465 |
| Isoform 4 of Active breakpoint cluster region-related protein OS=Mus musculus OX=10090 GN=Abr | Q5SSL4-4 | 0 | 22.51 | 6 | 4 | 29 | 4 | 822 | 93.7 | 6.44 | 50.56 | 0.992 |
| Isoform 4 of Ankyrin-2 OS=Mus musculus OX=10090 GN=Ank2 | Q8C8R3-4 | 0 | 233.011 | 13 | 28 | 235 | 24 | 3959 | 432.6 | 5.16 | 489.43 | 0.963 |
| Isoform 4 of Kalirin OS=Mus musculus OX=10090 GN=Kalrn | A2CG49-4 | 0 | 5.923 | 5 | 2 | 2 | 1 | 673 | 75.7 | 6.84 | 2.32 | |
| Isoform 4 of Mitochondrial fission factor OS=Mus musculus OX=10090 GN=Mff | Q6PCP5-4 | 0 | 34.197 | 33 | 3 | 44 | 1 | 218 | 25.1 | 5.4 | 98.66 | |
| Isoform 5 of Formin-1 OS=Mus musculus OX=10090 GN=Fmn1 | Q05860-5 | 0 | 5.322 | 3 | 2 | 2 | 2 | 1204 | 133.1 | 5.71 | 2.39 | |
| Isoform 5 of Phosphatidylinositol-binding clathrin assembly protein OS=Mus musculus OX=10090 GN=Picalm | Q7M6Y3-5 | 0 | 106.42 | 15 | 5 | 93 | 1 | 655 | 70.9 | 8.37 | 256.51 | 1.068 |
| Isoform 6 of CUGBP Elav-like family member 2 OS=Mus musculus OX=10090 GN=Celf2 | Q9Z0H4-6 | 0 | 34.499 | 5 | 2 | 20 | 2 | 524 | 55.8 | 8.6 | 60.92 | 1.154 |
| Isoform 6 of Palladin OS=Mus musculus OX=10090 GN=Palld | Q9ET54-6 | 0 | 165.155 | 12 | 11 | 162 | 11 | 1391 | 150.5 | 6.24 | 425.71 | 0.939 |
| Isoform 7 of Band 4.1-like protein 3 OS=Mus musculus OX=10090 GN=Epb41l3 | Q9WV92-7 | 0 | 219.918 | 37 | 20 | 207 | 4 | 823 | 91.8 | 5.5 | 488.83 | 1.033 |
| Isoform 9 of Kinectin OS=Mus musculus OX=10090 GN=Ktn1 | Q61595-9 | 0 | 238.961 | 30 | 25 | 208 | 25 | 1252 | 143.6 | 6.2 | 526.86 | 0.893 |
| Isoform A1-I of V-type proton ATPase 116 kDa subunit a isoform 1 OS=Mus musculus OX=10090 GN=Atp6v0a1 | Q9Z1G4-2 | 0 | 84.568 | 13 | 9 | 96 | 9 | 838 | 96.2 | 6.57 | 156.52 | 1.168 |
| Isoform Alpha-7X2B of Integrin alpha-7 OS=Mus musculus OX=10090 GN=Itga7 | Q61738-5 | 0 | 32.349 | 4 | 3 | 28 | 3 | 1135 | 124.6 | 5.91 | 78.96 | 1.795 |
| Isoform B of Fragile X mental retardation syndrome-related protein 1 OS=Mus musculus OX=10090 GN=Fxr1 | Q61584-3 | 0 | 57.244 | 11 | 4 | 57 | 4 | 677 | 76.2 | 7.53 | 143.19 | 1.076 |
| Isoform B of Reticulon-4 OS=Mus musculus OX=10090 GN=Rtn4 | Q99P72-5 | 0 | 80.211 | 26 | 5 | 169 | 1 | 356 | 38.4 | 4.78 | 421.28 | |
| Isoform C1 of Heterogeneous nuclear ribonucleoproteins C1/C2 OS=Mus musculus OX=10090 GN=Hnrnpc | Q9Z204-2 | 0 | 132.245 | 30 | 9 | 153 | 9 | 300 | 33 | 5.03 | 444.51 | 0.991 |
| Isoform Ib of Synapsin-1 OS=Mus musculus OX=10090 GN=Syn1 | O88935-1 | 0 | 12.661 | 7 | 2 | 8 | 2 | 670 | 70 | 9.83 | 7.94 | |
| Isoform J of Receptor-type tyrosine-protein phosphatase delta OS=Mus musculus OX=10090 GN=Ptprd | Q64487-10 | 0 | 21.172 | 4 | 4 | 20 | 2 | 1903 | 213.4 | 6.55 | 39.09 | 1.35 |
| Isoform M1 of Pyruvate kinase PKM OS=Mus musculus OX=10090 GN=Pkm | P52480-2 | 0 | 1019.007 | 57 | 35 | 2269 | 2 | 531 | 57.9 | 7.14 | 6589.86 | 1.259 |
| Isoform NFIX1 of Nuclear factor 1 X-type OS=Mus musculus OX=10090 GN=Nfix | P70257-1 | 0 | 17.985 | 11 | 3 | 14 | 3 | 441 | 48.8 | 8.47 | 21.21 | 1.245 |
| Isoform PLEC-1A of Plectin OS=Mus musculus OX=10090 GN=Plec | Q9QXS1-3 | 0 | 2102.562 | 38 | 147 | 2675 | 1 | 4543 | 517 | 5.8 | 6505.73 | 1.14 |
| Isoform Rpn10B of 26S proteasome non-ATPase regulatory subunit 4 OS=Mus musculus OX=10090 GN=Psmd4 | O35226-2 | 0 | 98.414 | 28 | 6 | 107 | 1 | 379 | 41 | 4.79 | 294.8 | 1.225 |
| Isoform Smooth muscle of Myosin light polypeptide 6 OS=Mus musculus OX=10090 GN=Myl6 | Q60605-2 | 0 | 168.049 | 55 | 7 | 345 | 1 | 151 | 17 | 4.55 | 898.48 | 1.259 |
| Isoleucine--tRNA ligase, cytoplasmic OS=Mus musculus OX=10090 GN=Iars PE=1 SV=2 | Q8BU30 | 0 | 115.858 | 16 | 14 | 104 | 14 | 1262 | 144.2 | 6.55 | 200.11 | 0.954 |
| Isoleucine--tRNA ligase, mitochondrial OS=Mus musculus OX=10090 GN=Iars2 PE=1 SV=1 | Q8BIJ6 | 0 | 84.097 | 10 | 7 | 48 | 7 | 1012 | 112.7 | 6.81 | 106.14 | 0.942 |
| Isopentenyl-diphosphate Delta-isomerase 1 OS=Mus musculus OX=10090 GN=Idi1 PE=1 SV=1 | P58044 | 0 | 51.104 | 23 | 4 | 44 | 4 | 227 | 26.3 | 6.16 | 111.6 | 0.762 |
| IST1 homolog OS=Mus musculus OX=10090 GN=Ist1 PE=1 SV=1 | Q9CX00 | 0 | 25.224 | 15 | 2 | 14 | 2 | 362 | 39.4 | 5.44 | 33.75 | 1.095 |
| Junction plakoglobin OS=Mus musculus OX=10090 GN=Jup PE=1 SV=3 | Q02257 | 0 | 58.567 | 19 | 9 | 43 | 7 | 745 | 81.7 | 6.14 | 78.31 | 0.767 |
| Junctional protein associated with coronary artery disease OS=Mus musculus OX=10090 GN=Jcad PE=1 SV=2 | Q5DTX6 | 0 | 6.648 | 3 | 2 | 2 | 2 | 1320 | 144.7 | 6.52 | 2.65 | 1.004 |
| Kelch repeat and BTB domain-containing protein 11 OS=Mus musculus OX=10090 GN=Kbtbd11 PE=1 SV=3 | Q8BNW9 | 0 | 5.889 | 6 | 2 | 2 | 2 | 633 | 67.9 | 5.39 | 6.13 | |
| Keratin, type I cytoskeletal 42 OS=Mus musculus OX=10090 GN=Krt42 PE=1 SV=1 | Q6IFX2 | 0 | 5.336 | 7 | 2 | 5 | 2 | 452 | 50.1 | 5.16 | 4.09 | |
| Keratin, type II cuticular Hb5 OS=Mus musculus OX=10090 GN=Krt85 PE=1 SV=2 | Q9Z2T6 | 0 | 6.73 | 6 | 2 | 9 | 1 | 507 | 55.7 | 6.42 | 14.16 | 3.028 |
| Keratin, type II cytoskeletal 5 OS=Mus musculus OX=10090 GN=Krt5 PE=1 SV=1 | Q922U2 | 0 | 12.843 | 4 | 2 | 32 | 1 | 580 | 61.7 | 7.75 | 74.1 | 0.039 |
| Keratin, type II cytoskeletal 6A OS=Mus musculus OX=10090 GN=Krt6a PE=1 SV=3 | P50446 | 0 | 11.546 | 5 | 2 | 31 | 1 | 553 | 59.3 | 7.94 | 69.38 | |
| Keratin, type II cytoskeletal 7 OS=Mus musculus OX=10090 GN=Krt7 PE=1 SV=1 | Q9DCV7 | 0 | 20.618 | 11 | 4 | 33 | 3 | 457 | 50.7 | 5.87 | 80.7 | 1.225 |
| Keratin, type II cytoskeletal 75 OS=Mus musculus OX=10090 GN=Krt75 PE=1 SV=1 | Q8BGZ7 | 0 | 14.578 | 7 | 2 | 32 | 1 | 551 | 59.7 | 8.31 | 71.97 | |
| Ketimine reductase mu-crystallin OS=Mus musculus OX=10090 GN=Crym PE=1 SV=1 | O54983 | 0 | 42.358 | 15 | 4 | 39 | 4 | 313 | 33.5 | 5.67 | 116.35 | 0.806 |
| KH domain-containing, RNA-binding, signal transduction-associated protein 1 OS=Mus musculus OX=10090 GN=Khdrbs1 PE=1 SV=2 | Q60749 | 0 | 37.556 | 10 | 4 | 38 | 4 | 443 | 48.3 | 8.72 | 85.83 | 0.908 |
| KH domain-containing, RNA-binding, signal transduction-associated protein 3 OS=Mus musculus OX=10090 GN=Khdrbs3 PE=1 SV=1 | Q9R226 | 0 | 8.054 | 12 | 2 | 9 | 2 | 346 | 38.8 | 8.1 | 25.69 | 0.966 |
| KIF1-binding protein OS=Mus musculus OX=10090 GN=Kif1bp PE=1 SV=2 | Q6ZPU9 | 0 | 52.643 | 6 | 2 | 35 | 2 | 617 | 71 | 5.48 | 112.3 | 0.999 |
| Kin of IRRE-like protein 1 OS=Mus musculus OX=10090 GN=Kirrel1 PE=1 SV=1 | Q80W68 | 0 | 23.603 | 14 | 5 | 11 | 5 | 789 | 87.1 | 5.92 | 29.35 | 1.162 |
| Kinesin heavy chain isoform 5C OS=Mus musculus OX=10090 GN=Kif5c PE=1 SV=3 | P28738 | 0 | 85.021 | 8 | 6 | 87 | 1 | 956 | 109.2 | 6.19 | 256.01 | |
| Kinesin light chain 1 OS=Mus musculus OX=10090 GN=Klc1 PE=1 SV=3 | O88447 | 0 | 95.821 | 25 | 12 | 132 | 11 | 541 | 61.4 | 5.68 | 354.99 | 1.037 |
| Kinesin light chain 4 OS=Mus musculus OX=10090 GN=Klc4 PE=1 SV=1 | Q9DBS5 | 0 | 23.24 | 8 | 3 | 13 | 2 | 619 | 68.6 | 6.09 | 29.59 | 0.429 |
| Kinesin-1 heavy chain OS=Mus musculus OX=10090 GN=Kif5b PE=1 SV=3 | Q61768 | 0 | 313.986 | 28 | 23 | 376 | 18 | 963 | 109.5 | 6.44 | 1060.12 | 1.083 |
| Kinesin-associated protein 3 OS=Mus musculus OX=10090 GN=Kifap3 PE=1 SV=1 | P70188 | 0 | 26.436 | 6 | 3 | 11 | 3 | 793 | 91.2 | 5.11 | 28.3 | 1.388 |
| Kinesin-like protein KIF13A OS=Mus musculus OX=10090 GN=Kif13a PE=1 SV=1 | Q9EQW7 | 0 | 9.586 | 1 | 2 | 13 | 1 | 1749 | 195.7 | 5.41 | 31.19 | 0.361 |
| Kinesin-like protein KIF16B OS=Mus musculus OX=10090 GN=Kif16b PE=1 SV=1 | B1AVY7 | 0 | 16.248 | 3 | 4 | 17 | 3 | 1312 | 150 | 6.44 | 37.73 | |
| Kinesin-like protein KIF1B OS=Mus musculus OX=10090 GN=Kif1b PE=1 SV=2 | Q60575 | 0 | 47.039 | 6 | 7 | 28 | 1 | 1816 | 204 | 5.62 | 69.85 | 0.685 |
| Kinesin-like protein KIF1C OS=Mus musculus OX=10090 GN=Kif1c PE=1 SV=2 | O35071 | 0 | 60.598 | 13 | 10 | 44 | 7 | 1100 | 122.4 | 7.03 | 105.95 | 1.068 |
| Kinesin-like protein KIF21A OS=Mus musculus OX=10090 GN=Kif21a PE=1 SV=2 | Q9QXL2 | 0 | 45.699 | 7 | 6 | 37 | 6 | 1672 | 186.4 | 6.27 | 93.15 | 1.234 |
| Kinesin-like protein KIF3A OS=Mus musculus OX=10090 GN=Kif3a PE=1 SV=2 | P28741 | 0 | 43.279 | 8 | 5 | 43 | 5 | 701 | 80.1 | 6.54 | 99 | 0.83 |
| Kinesin-like protein KIF3B OS=Mus musculus OX=10090 GN=Kif3b PE=1 SV=1 | Q61771 | 0 | 17.447 | 7 | 3 | 9 | 3 | 747 | 85.2 | 7.69 | 20.59 | |
| Kinetochore protein Spc24 OS=Mus musculus OX=10090 GN=Spc24 PE=1 SV=1 | Q9D083 | 0 | 5.935 | 16 | 2 | 3 | 2 | 201 | 23.4 | 4.64 | 5 | |
| KN motif and ankyrin repeat domain-containing protein 2 OS=Mus musculus OX=10090 GN=Kank2 PE=1 SV=1 | Q8BX02 | 0 | 84.89 | 16 | 10 | 84 | 10 | 843 | 90.2 | 5.55 | 209.75 | 1.02 |
| Lactoylglutathione lyase OS=Mus musculus OX=10090 GN=Glo1 PE=1 SV=3 | Q9CPU0 | 0 | 43.464 | 26 | 2 | 47 | 2 | 184 | 20.8 | 5.47 | 104.81 | 0.829 |
| Lambda-crystallin homolog OS=Mus musculus OX=10090 GN=Cryl1 PE=1 SV=3 | Q99KP3 | 0 | 18.483 | 16 | 3 | 15 | 3 | 319 | 35.2 | 5.86 | 20.93 | 1.126 |
| Lamina-associated polypeptide 2, isoforms alpha/zeta OS=Mus musculus OX=10090 GN=Tmpo PE=1 SV=4 | Q61033 | 0 | 113.408 | 14 | 6 | 118 | 4 | 693 | 75.1 | 8.05 | 348.22 | 0.992 |
| Lamina-associated polypeptide 2, isoforms beta/delta/epsilon/gamma OS=Mus musculus OX=10090 GN=Tmpo PE=1 SV=4 | Q61029 | 0 | 92.277 | 16 | 5 | 137 | 3 | 452 | 50.3 | 9.45 | 399.22 | 1.008 |
| Lamin-B1 OS=Mus musculus OX=10090 GN=Lmnb1 PE=1 SV=3 | P14733 | 0 | 292.896 | 46 | 22 | 334 | 21 | 588 | 66.7 | 5.16 | 808.68 | 1.017 |
| Lamin-B2 OS=Mus musculus OX=10090 GN=Lmnb2 PE=1 SV=2 | P21619 | 0 | 175.666 | 27 | 10 | 167 | 9 | 596 | 67.3 | 5.5 | 462.54 | 1.022 |
| Laminin subunit alpha-1 OS=Mus musculus OX=10090 GN=Lama1 PE=1 SV=2 | P19137 | 0 | 9.015 | 1 | 2 | 4 | 2 | 3083 | 337.9 | 6.71 | 10.53 | 0.767 |
| Laminin subunit alpha-2 OS=Mus musculus OX=10090 GN=Lama2 PE=1 SV=2 | Q60675 | 0 | 173.595 | 8 | 18 | 142 | 18 | 3118 | 343.6 | 6.09 | 408.03 | 1.102 |
| Laminin subunit alpha-4 OS=Mus musculus OX=10090 GN=Lama4 PE=1 SV=2 | P97927 | 0 | 201.736 | 13 | 12 | 228 | 12 | 1816 | 201.7 | 6.21 | 657.1 | 1.279 |
| Laminin subunit alpha-5 OS=Mus musculus OX=10090 GN=Lama5 PE=1 SV=4 | Q61001 | 0 | 16.583 | 1 | 3 | 10 | 3 | 3718 | 403.8 | 6.73 | 30.41 | 1.011 |
| Laminin subunit beta-1 OS=Mus musculus OX=10090 GN=Lamb1 PE=1 SV=3 | P02469 | 0 | 293.111 | 15 | 20 | 347 | 20 | 1786 | 197 | 4.94 | 923.37 | 1.167 |
| Laminin subunit beta-2 OS=Mus musculus OX=10090 GN=Lamb2 PE=1 SV=2 | Q61292 | 0 | 192.16 | 16 | 21 | 195 | 21 | 1799 | 196.5 | 6.67 | 469.77 | 0.96 |
| Laminin subunit gamma-1 OS=Mus musculus OX=10090 GN=Lamc1 PE=1 SV=2 | P02468 | 0 | 314.615 | 23 | 24 | 298 | 24 | 1607 | 177.2 | 5.21 | 838.05 | 1.107 |
| LanC-like protein 2 OS=Mus musculus OX=10090 GN=Lancl2 PE=1 SV=1 | Q9JJK2 | 0 | 26.96 | 14 | 3 | 19 | 3 | 450 | 50.7 | 7.28 | 35.69 | 0.931 |
| Lanosterol 14-alpha demethylase OS=Mus musculus OX=10090 GN=Cyp51a1 PE=1 SV=1 | Q8K0C4 | 0 | 16.847 | 10 | 4 | 12 | 4 | 503 | 56.7 | 8.41 | 23.15 | 0.68 |
| Lanosterol synthase OS=Mus musculus OX=10090 GN=Lss PE=1 SV=2 | Q8BLN5 | 0 | 23.624 | 8 | 4 | 21 | 4 | 733 | 83.1 | 6.4 | 55.31 | 0.737 |
| La-related protein 1 OS=Mus musculus OX=10090 GN=Larp1 PE=1 SV=3 | Q6ZQ58 | 0 | 43.384 | 5 | 4 | 32 | 4 | 1072 | 121.1 | 8.79 | 66.82 | 1.127 |
| La-related protein 4B OS=Mus musculus OX=10090 GN=Larp4b PE=1 SV=2 | Q6A0A2 | 0 | 13.746 | 3 | 2 | 10 | 2 | 741 | 81.6 | 7.49 | 23.55 | 0.662 |
| Large proline-rich protein BAG6 OS=Mus musculus OX=10090 GN=Bag6 PE=1 SV=1 | Q9Z1R2 | 0 | 79.615 | 11 | 8 | 64 | 8 | 1154 | 121 | 5.71 | 162.22 | 0.787 |
| Latent-transforming growth factor beta-binding protein 1 OS=Mus musculus OX=10090 GN=Ltbp1 PE=1 SV=2 | Q8CG19 | 0 | 52.321 | 6 | 6 | 28 | 6 | 1712 | 186.6 | 6.02 | 67.64 | |
| Latent-transforming growth factor beta-binding protein 2 OS=Mus musculus OX=10090 GN=Ltbp2 PE=1 SV=2 | O08999 | 0 | 17.36 | 4 | 4 | 6 | 4 | 1813 | 195.7 | 5.29 | 10.42 | 1.163 |
| Latent-transforming growth factor beta-binding protein 4 OS=Mus musculus OX=10090 GN=Ltbp4 PE=1 SV=2 | Q8K4G1 | 0 | 139.26 | 15 | 14 | 141 | 14 | 1666 | 178.5 | 5.47 | 321.9 | 1.061 |
| Latexin OS=Mus musculus OX=10090 GN=Lxn PE=1 SV=2 | P70202 | 0 | 24.109 | 20 | 3 | 27 | 3 | 222 | 25.5 | 5.74 | 42.26 | 0.824 |
| Legumain OS=Mus musculus OX=10090 GN=Lgmn PE=1 SV=1 | O89017 | 0 | 111.901 | 20 | 8 | 154 | 8 | 435 | 49.3 | 6.39 | 380.25 | 1.377 |
| LEM domain-containing protein 2 OS=Mus musculus OX=10090 GN=Lemd2 PE=1 SV=1 | Q6DVA0 | 0 | 10.122 | 5 | 2 | 7 | 2 | 511 | 57.5 | 9.01 | 14.33 | 0.642 |
| Lethal(2) giant larvae protein homolog 1 OS=Mus musculus OX=10090 GN=Llgl1 PE=1 SV=1 | Q80Y17 | 0 | 23.054 | 5 | 2 | 17 | 2 | 1036 | 112.5 | 6.48 | 28.75 | |
| Leucine zipper protein 1 OS=Mus musculus OX=10090 GN=Luzp1 PE=1 SV=2 | Q8R4U7 | 0 | 55.894 | 13 | 7 | 31 | 7 | 1068 | 119.2 | 7.99 | 58.81 | 1.126 |
| Leucine zipper transcription factor-like protein 1 OS=Mus musculus OX=10090 GN=Lztfl1 PE=1 SV=1 | Q9JHQ5 | 0 | 50.305 | 19 | 5 | 53 | 5 | 299 | 34.8 | 5.17 | 159.19 | 1.215 |
| Leucine-rich glioma-inactivated protein 1 OS=Mus musculus OX=10090 GN=Lgi1 PE=1 SV=1 | Q9JIA1 | 0 | 8.769 | 8 | 2 | 5 | 2 | 557 | 63.6 | 8.02 | 5.3 | |
| Leucine-rich PPR motif-containing protein, mitochondrial OS=Mus musculus OX=10090 GN=Lrpprc PE=1 SV=2 | Q6PB66 | 0 | 140.549 | 17 | 16 | 164 | 16 | 1392 | 156.5 | 6.83 | 413.05 | 0.964 |
| Leucine-rich repeat and calponin homology domain-containing protein 3 OS=Mus musculus OX=10090 GN=Lrch3 PE=1 SV=3 | Q8BVU0 | 0 | 38.619 | 13 | 5 | 16 | 5 | 778 | 86.3 | 7.03 | 30.87 | 0.802 |
| Leucine-rich repeat and calponin homology domain-containing protein 4 OS=Mus musculus OX=10090 GN=Lrch4 PE=1 SV=1 | Q921G6 | 0 | 22.254 | 8 | 3 | 11 | 3 | 680 | 73.1 | 7.53 | 30 | 0.897 |
| Leucine-rich repeat flightless-interacting protein 1 OS=Mus musculus OX=10090 GN=Lrrfip1 PE=1 SV=2 | Q3UZ39 | 0 | 126.285 | 27 | 9 | 98 | 5 | 729 | 79.2 | 4.82 | 206.5 | 0.716 |
| Leucine-rich repeat flightless-interacting protein 2 OS=Mus musculus OX=10090 GN=Lrrfip2 PE=1 SV=1 | Q91WK0 | 0 | 40.592 | 15 | 5 | 35 | 5 | 415 | 47.1 | 5.68 | 66.21 | 1.013 |
| Leucine-rich repeat protein SHOC-2 OS=Mus musculus OX=10090 GN=Shoc2 PE=1 SV=2 | O88520 | 0 | 19.234 | 6 | 2 | 10 | 2 | 582 | 64.9 | 8.46 | 30.3 | 0.696 |
| Leucine-rich repeat-containing protein 1 OS=Mus musculus OX=10090 GN=Lrrc1 PE=1 SV=2 | Q80VQ1 | 0 | 26.173 | 7 | 2 | 29 | 1 | 524 | 59.4 | 5.16 | 86.12 | |
| Leucine-rich repeat-containing protein 25 OS=Mus musculus OX=10090 GN=Lrrc25 PE=1 SV=1 | Q8K1T1 | 0 | 23.184 | 11 | 2 | 23 | 2 | 297 | 32.7 | 8.24 | 71.09 | 1.38 |
| Leucine-rich repeat-containing protein 40 OS=Mus musculus OX=10090 GN=Lrrc40 PE=1 SV=2 | Q9CRC8 | 0 | 26.247 | 6 | 2 | 17 | 2 | 602 | 68 | 6.89 | 44.17 | 1.046 |
| Leucine-rich repeat-containing protein 47 OS=Mus musculus OX=10090 GN=Lrrc47 PE=1 SV=1 | Q505F5 | 0 | 54.152 | 15 | 5 | 65 | 5 | 581 | 63.6 | 8.1 | 158.71 | 1.087 |
| Leucine-rich repeat-containing protein 58 OS=Mus musculus OX=10090 GN=Lrrc58 PE=2 SV=1 | Q3UGP9 | 0 | 14.149 | 8 | 2 | 10 | 2 | 366 | 40.1 | 6.62 | 27.19 | |
| Leucine-rich repeat-containing protein 59 OS=Mus musculus OX=10090 GN=Lrrc59 PE=1 SV=1 | Q922Q8 | 0 | 156.207 | 40 | 9 | 139 | 9 | 307 | 34.9 | 9.52 | 341.56 | 0.817 |
| Leucine--tRNA ligase, cytoplasmic OS=Mus musculus OX=10090 GN=Lars PE=1 SV=2 | Q8BMJ2 | 0 | 273.012 | 26 | 22 | 359 | 22 | 1178 | 134.1 | 7.05 | 813.81 | 0.984 |
| Leucyl-cystinyl aminopeptidase OS=Mus musculus OX=10090 GN=Lnpep PE=1 SV=1 | Q8C129 | 0 | 59.441 | 10 | 7 | 36 | 7 | 1025 | 117.2 | 5.96 | 78.64 | 1.085 |
| Leukemia inhibitory factor receptor OS=Mus musculus OX=10090 GN=Lifr PE=1 SV=1 | P42703 | 0 | 8.977 | 3 | 2 | 6 | 2 | 1092 | 122.5 | 6.04 | 12.27 | |
| Leukocyte elastase inhibitor A OS=Mus musculus OX=10090 GN=Serpinb1a PE=1 SV=1 | Q9D154 | 0 | 136.044 | 35 | 12 | 146 | 12 | 379 | 42.5 | 6.21 | 313.63 | 0.898 |
| Leukocyte surface antigen CD47 OS=Mus musculus OX=10090 GN=Cd47 PE=1 SV=2 | Q61735 | 0 | 42.59 | 9 | 2 | 72 | 2 | 303 | 33.1 | 8.63 | 151.95 | 1.017 |
| Leukotriene A-4 hydrolase OS=Mus musculus OX=10090 GN=Lta4h PE=1 SV=4 | P24527 | 0 | 134.58 | 27 | 13 | 130 | 13 | 611 | 69 | 6.42 | 301.92 | 0.819 |
| Leupaxin OS=Mus musculus OX=10090 GN=Lpxn PE=1 SV=2 | Q99N69 | 0 | 20.372 | 9 | 2 | 8 | 2 | 386 | 43.5 | 6.43 | 27.65 | 1.155 |
| LIM and SH3 domain protein 1 OS=Mus musculus OX=10090 GN=Lasp1 PE=1 SV=1 | Q61792 | 0 | 45.856 | 23 | 5 | 69 | 5 | 263 | 30 | 7.05 | 126.3 | 0.823 |
| LIM domain and actin-binding protein 1 OS=Mus musculus OX=10090 GN=Lima1 PE=1 SV=3 | Q9ERG0 | 0 | 81.71 | 8 | 4 | 129 | 4 | 753 | 84 | 6.6 | 338.57 | 0.876 |
| LIM domain-containing protein 1 OS=Mus musculus OX=10090 GN=Limd1 PE=1 SV=2 | Q9QXD8 | 0 | 23.697 | 8 | 3 | 13 | 3 | 668 | 71.4 | 6.32 | 15.2 | 0.882 |
| Lipoamide acyltransferase component of branched-chain alpha-keto acid dehydrogenase complex, mitochondrial OS=Mus musculus OX=10090 GN=Dbt PE=1 SV=2 | P53395 | 0 | 33.081 | 9 | 4 | 34 | 4 | 482 | 53.2 | 8.6 | 92.76 | 0.873 |
| Lipoma-preferred partner homolog OS=Mus musculus OX=10090 GN=Lpp PE=1 SV=1 | Q8BFW7 | 0 | 53.473 | 13 | 5 | 91 | 5 | 613 | 65.8 | 7.37 | 179.68 | 1.076 |
| Lipopolysaccharide-responsive and beige-like anchor protein OS=Mus musculus OX=10090 GN=Lrba PE=1 SV=1 | Q9ESE1 | 0 | 12.134 | 2 | 3 | 5 | 3 | 2856 | 316.9 | 5.69 | 11.14 | |
| Liprin-alpha-2 OS=Mus musculus OX=10090 GN=Ppfia2 PE=1 SV=2 | Q8BSS9 | 0 | 8.841 | 1 | 2 | 5 | 2 | 1257 | 143.1 | 6.11 | 9.58 | 0.473 |
| Liprin-beta-1 OS=Mus musculus OX=10090 GN=Ppfibp1 PE=1 SV=3 | Q8C8U0 | 0 | 150.479 | 16 | 12 | 185 | 10 | 969 | 108.5 | 5.49 | 459 | 1.026 |
| Liprin-beta-2 OS=Mus musculus OX=10090 GN=Ppfibp2 PE=1 SV=3 | O35711 | 0 | 29.854 | 10 | 6 | 25 | 4 | 882 | 98.7 | 6.33 | 48.51 | 1.786 |
| L-lactate dehydrogenase A chain OS=Mus musculus OX=10090 GN=Ldha PE=1 SV=3 | P06151 | 0 | 199.943 | 51 | 16 | 258 | 16 | 332 | 36.5 | 7.74 | 598.53 | 1.04 |
| L-lactate dehydrogenase B chain OS=Mus musculus OX=10090 GN=Ldhb PE=1 SV=2 | P16125 | 0 | 75.253 | 17 | 7 | 68 | 7 | 334 | 36.5 | 6.05 | 168.46 | 1.235 |
| LMBR1 domain-containing protein 2 OS=Mus musculus OX=10090 GN=Lmbrd2 PE=1 SV=1 | Q8C561 | 0 | 29.229 | 7 | 3 | 19 | 3 | 694 | 81 | 7.34 | 46.16 | 0.767 |
| Lon protease homolog 2, peroxisomal OS=Mus musculus OX=10090 GN=Lonp2 PE=1 SV=1 | Q9DBN5 | 0.001 | 4.621 | 3 | 2 | 3 | 2 | 852 | 94.5 | 7.23 | 5.99 | |
| Lon protease homolog, mitochondrial OS=Mus musculus OX=10090 GN=Lonp1 PE=1 SV=2 | Q8CGK3 | 0 | 120.521 | 29 | 16 | 122 | 16 | 949 | 105.8 | 6.57 | 282.05 | 1.103 |
| Long-chain fatty acid transport protein 1 OS=Mus musculus OX=10090 GN=Slc27a1 PE=1 SV=1 | Q60714 | 0 | 22.907 | 6 | 2 | 16 | 2 | 646 | 71.2 | 8.38 | 31.52 | 0.896 |
| Long-chain fatty acid transport protein 4 OS=Mus musculus OX=10090 GN=Slc27a4 PE=1 SV=1 | Q91VE0 | 0 | 24.511 | 15 | 5 | 18 | 5 | 643 | 72.3 | 8.59 | 37.65 | 0.793 |
| Long-chain specific acyl-CoA dehydrogenase, mitochondrial OS=Mus musculus OX=10090 GN=Acadl PE=1 SV=2 | P51174 | 0 | 52.936 | 10 | 3 | 66 | 3 | 430 | 47.9 | 8.31 | 229.81 | 1.175 |
| Long-chain-fatty-acid--CoA ligase 1 OS=Mus musculus OX=10090 GN=Acsl1 PE=1 SV=2 | P41216 | 0 | 34.783 | 9 | 6 | 20 | 6 | 699 | 77.9 | 7.15 | 35.72 | 0.816 |
| Long-chain-fatty-acid--CoA ligase 3 OS=Mus musculus OX=10090 GN=Acsl3 PE=1 SV=2 | Q9CZW4 | 0 | 48.779 | 11 | 5 | 54 | 4 | 720 | 80.4 | 8.54 | 138.71 | 0.929 |
| Long-chain-fatty-acid--CoA ligase 4 OS=Mus musculus OX=10090 GN=Acsl4 PE=1 SV=2 | Q9QUJ7 | 0 | 33.273 | 9 | 4 | 24 | 3 | 711 | 79 | 8.28 | 66.56 | 1.965 |
| Long-chain-fatty-acid--CoA ligase 5 OS=Mus musculus OX=10090 GN=Acsl5 PE=1 SV=1 | Q8JZR0 | 0 | 18.545 | 4 | 2 | 20 | 2 | 683 | 76.2 | 7.09 | 50.13 | 0.917 |
| Low molecular weight phosphotyrosine protein phosphatase OS=Mus musculus OX=10090 GN=Acp1 PE=1 SV=3 | Q9D358 | 0 | 45.597 | 49 | 5 | 32 | 5 | 158 | 18.2 | 6.74 | 64.61 | 1.052 |
| Low-density lipoprotein receptor-related protein 4 OS=Mus musculus OX=10090 GN=Lrp4 PE=1 SV=3 | Q8VI56 | 0 | 16.142 | 3 | 3 | 10 | 3 | 1905 | 211.8 | 5.39 | 31.12 | 1.919 |
| Low-density lipoprotein receptor-related protein 6 OS=Mus musculus OX=10090 GN=Lrp6 PE=1 SV=1 | O88572 | 0 | 9.577 | 3 | 3 | 5 | 3 | 1613 | 180.1 | 5.33 | 10.13 | 0.628 |
| LRP chaperone MESD OS=Mus musculus OX=10090 GN=Mesd PE=1 SV=1 | Q9ERE7 | 0 | 47.059 | 33 | 4 | 49 | 4 | 224 | 25.2 | 6.34 | 92.73 | 0.864 |
| Luc7-like protein 3 OS=Mus musculus OX=10090 GN=Luc7l3 PE=1 SV=1 | Q5SUF2 | 0 | 33.779 | 9 | 4 | 28 | 4 | 432 | 51.4 | 9.77 | 52.82 | 0.885 |
| Lupus La protein homolog OS=Mus musculus OX=10090 GN=Ssb PE=1 SV=1 | P32067 | 0 | 69.953 | 19 | 7 | 82 | 7 | 415 | 47.7 | 9.77 | 225.32 | 0.906 |
| L-xylulose reductase OS=Mus musculus OX=10090 GN=Dcxr PE=1 SV=2 | Q91X52 | 0.002 | 4.464 | 12 | 2 | 5 | 2 | 244 | 25.7 | 7.31 | 9.84 | 2.219 |
| Lymphocyte-specific protein 1 OS=Mus musculus OX=10090 GN=Lsp1 PE=1 SV=2 | P19973 | 0 | 66.251 | 40 | 7 | 35 | 7 | 330 | 36.7 | 4.82 | 81.26 | 0.902 |
| Lys-63-specific deubiquitinase BRCC36 OS=Mus musculus OX=10090 GN=Brcc3 PE=1 SV=1 | P46737 | 0 | 19.087 | 15 | 3 | 16 | 3 | 291 | 33.3 | 5.83 | 22.32 | |
| Lysine-specific demethylase 3B OS=Mus musculus OX=10090 GN=Kdm3b PE=1 SV=1 | Q6ZPY7 | 0 | 24.822 | 2 | 2 | 19 | 2 | 1562 | 170.8 | 6.92 | 56.24 | 1.577 |
| Lysine-specific histone demethylase 1A OS=Mus musculus OX=10090 GN=Kdm1a PE=1 SV=2 | Q6ZQ88 | 0 | 10.279 | 4 | 3 | 10 | 3 | 853 | 92.8 | 6.52 | 10.44 | 2.033 |
| Lysine--tRNA ligase OS=Mus musculus OX=10090 GN=Kars PE=1 SV=1 | Q99MN1 | 0 | 52.726 | 12 | 4 | 128 | 4 | 595 | 67.8 | 5.94 | 248.55 | 0.883 |
| Lysophosphatidylcholine acyltransferase 1 OS=Mus musculus OX=10090 GN=Lpcat1 PE=1 SV=1 | Q3TFD2 | 0 | 24.548 | 12 | 4 | 11 | 4 | 534 | 59.7 | 6.34 | 32.24 | 0.996 |
| Lysophosphatidylserine lipase ABHD12 OS=Mus musculus OX=10090 GN=Abhd12 PE=1 SV=2 | Q99LR1 | 0 | 35.412 | 16 | 4 | 23 | 4 | 398 | 45.2 | 8.72 | 61.84 | 0.842 |
| Lysophospholipase D GDPD1 OS=Mus musculus OX=10090 GN=Gdpd1 PE=1 SV=1 | Q9CRY7 | 0 | 21.118 | 15 | 3 | 15 | 3 | 314 | 35.8 | 8.31 | 28.96 | 1.072 |
| Lysosomal acid glucosylceramidase OS=Mus musculus OX=10090 GN=Gba PE=1 SV=1 | P17439 | 0 | 24.503 | 12 | 4 | 30 | 4 | 515 | 57.6 | 7.75 | 71.78 | 0.773 |
| Lysosomal alpha-glucosidase OS=Mus musculus OX=10090 GN=Gaa PE=1 SV=2 | P70699 | 0 | 116.425 | 9 | 6 | 177 | 6 | 953 | 106.2 | 5.83 | 464.43 | 0.933 |
| Lysosomal alpha-mannosidase OS=Mus musculus OX=10090 GN=Man2b1 PE=1 SV=4 | O09159 | 0 | 95.247 | 15 | 10 | 88 | 10 | 1013 | 114.6 | 8.13 | 238.74 | 0.804 |
| Lysosomal protective protein OS=Mus musculus OX=10090 GN=Ctsa PE=1 SV=1 | P16675 | 0 | 86.766 | 17 | 5 | 74 | 5 | 474 | 53.8 | 5.86 | 204.31 | 1.206 |
| Lysosomal thioesterase PPT2 OS=Mus musculus OX=10090 GN=Ppt2 PE=1 SV=1 | O35448 | 0 | 5.868 | 9 | 2 | 7 | 2 | 302 | 34.3 | 6.39 | 10.93 | |
| Lysosome membrane protein 2 OS=Mus musculus OX=10090 GN=Scarb2 PE=1 SV=3 | O35114 | 0 | 18.395 | 9 | 3 | 32 | 3 | 478 | 54 | 5.1 | 71.84 | 0.839 |
| Lysosome-associated membrane glycoprotein 1 OS=Mus musculus OX=10090 GN=Lamp1 PE=1 SV=2 | P11438 | 0 | 96.028 | 8 | 4 | 130 | 4 | 406 | 43.8 | 8.4 | 270.93 | 1.222 |
| Lysosome-associated membrane glycoprotein 2 OS=Mus musculus OX=10090 GN=Lamp2 PE=1 SV=2 | P17047 | 0 | 22.382 | 12 | 5 | 20 | 5 | 415 | 45.7 | 7.39 | 35.3 | 0.977 |
| Lysozyme C-2 OS=Mus musculus OX=10090 GN=Lyz2 PE=1 SV=2 | P08905 | 0 | 104.657 | 50 | 5 | 155 | 5 | 148 | 16.7 | 8.81 | 437.64 | 1.628 |
| Lysyl oxidase homolog 1 OS=Mus musculus OX=10090 GN=Loxl1 PE=2 SV=3 | P97873 | 0 | 47.83 | 23 | 7 | 58 | 7 | 607 | 66.5 | 7.33 | 169.74 | 1.409 |
| Lysyl oxidase homolog 2 OS=Mus musculus OX=10090 GN=Loxl2 PE=1 SV=2 | P58022 | 0 | 22.3 | 5 | 2 | 17 | 2 | 776 | 86.9 | 6.57 | 24.1 | 1.312 |
| m7GpppX diphosphatase OS=Mus musculus OX=10090 GN=Dcps PE=1 SV=1 | Q9DAR7 | 0 | 23.576 | 14 | 3 | 50 | 3 | 338 | 39 | 6.48 | 115.84 | 1.043 |
| Macrophage mannose receptor 1 OS=Mus musculus OX=10090 GN=Mrc1 PE=1 SV=2 | Q61830 | 0 | 117.902 | 14 | 13 | 141 | 13 | 1456 | 164.9 | 6.83 | 380.65 | 1.215 |
| Macrophage migration inhibitory factor OS=Mus musculus OX=10090 GN=Mif PE=1 SV=2 | P34884 | 0 | 24.545 | 26 | 2 | 64 | 2 | 115 | 12.5 | 7.34 | 144.63 | 1.1 |
| Macrophage receptor MARCO OS=Mus musculus OX=10090 GN=Marco PE=1 SV=1 | Q60754 | 0 | 33.646 | 14 | 3 | 21 | 3 | 518 | 52.7 | 7.84 | 61.34 | 1.326 |
| Macrophage scavenger receptor types I and II OS=Mus musculus OX=10090 GN=Msr1 PE=1 SV=3 | P30204 | 0 | 20.614 | 9 | 2 | 10 | 2 | 458 | 50.1 | 6.54 | 24.33 | 1.644 |
| Macrophage-capping protein OS=Mus musculus OX=10090 GN=Capg PE=1 SV=2 | P24452 | 0 | 101.501 | 20 | 4 | 279 | 4 | 352 | 39.2 | 7.2 | 707.05 | 0.963 |
| Macrophage-expressed gene 1 protein OS=Mus musculus OX=10090 GN=Mpeg1 PE=1 SV=1 | A1L314 | 0 | 19.217 | 6 | 3 | 24 | 3 | 713 | 78.3 | 5.92 | 56.67 | 0.427 |
| Magnesium-dependent phosphatase 1 OS=Mus musculus OX=10090 GN=Mdp1 PE=1 SV=1 | Q9D967 | 0 | 59.458 | 31 | 3 | 56 | 3 | 164 | 18.6 | 6.8 | 166.18 | 0.83 |
| MAGUK p55 subfamily member 2 OS=Mus musculus OX=10090 GN=Mpp2 PE=1 SV=1 | Q9WV34 | 0 | 9.102 | 7 | 2 | 3 | 2 | 552 | 61.5 | 6.44 | 4.84 | |
| MAGUK p55 subfamily member 6 OS=Mus musculus OX=10090 GN=Mpp6 PE=1 SV=1 | Q9JLB0 | 0 | 42.613 | 10 | 4 | 60 | 4 | 553 | 62.6 | 6.4 | 161.34 | 1 |
| Major vault protein OS=Mus musculus OX=10090 GN=Mvp PE=1 SV=4 | Q9EQK5 | 0 | 319.354 | 32 | 19 | 434 | 19 | 861 | 95.9 | 5.59 | 1290.67 | 1.024 |
| Malate dehydrogenase, cytoplasmic OS=Mus musculus OX=10090 GN=Mdh1 PE=1 SV=3 | P14152 | 0 | 89.621 | 35 | 9 | 170 | 9 | 334 | 36.5 | 6.58 | 397.45 | 0.991 |
| Malate dehydrogenase, mitochondrial OS=Mus musculus OX=10090 GN=Mdh2 PE=1 SV=3 | P08249 | 0 | 170.961 | 50 | 14 | 436 | 14 | 338 | 35.6 | 8.68 | 1134.72 | 0.971 |
| Malectin OS=Mus musculus OX=10090 GN=Mlec PE=1 SV=2 | Q6ZQI3 | 0 | 37.523 | 22 | 4 | 29 | 4 | 291 | 32.3 | 6.05 | 52.07 | 0.812 |
| Maleylacetoacetate isomerase OS=Mus musculus OX=10090 GN=Gstz1 PE=1 SV=1 | Q9WVL0 | 0 | 13.971 | 19 | 2 | 10 | 2 | 216 | 24.3 | 7.85 | 21.19 | |
| Malignant T-cell-amplified sequence 1 OS=Mus musculus OX=10090 GN=Mcts1 PE=1 SV=1 | Q9DB27 | 0 | 28.724 | 27 | 3 | 42 | 3 | 181 | 20.5 | 8.82 | 58.34 | 0.941 |
| Mammalian ependymin-related protein 1 OS=Mus musculus OX=10090 GN=Epdr1 PE=1 SV=1 | Q99M71 | 0 | 16.372 | 9 | 2 | 27 | 2 | 224 | 25.5 | 7.58 | 78.51 | 0.733 |
| Manganese-transporting ATPase 13A1 OS=Mus musculus OX=10090 GN=Atp13a1 PE=1 SV=2 | Q9EPE9 | 0 | 34.509 | 10 | 7 | 20 | 7 | 1200 | 132.3 | 8.03 | 51.07 | 0.711 |
| Mannose-1-phosphate guanyltransferase beta OS=Mus musculus OX=10090 GN=Gmppb PE=1 SV=1 | Q8BTZ7 | 0 | 24.898 | 16 | 3 | 19 | 3 | 360 | 39.9 | 6.74 | 50.05 | 0.84 |
| Mannose-6-phosphate isomerase OS=Mus musculus OX=10090 GN=Mpi PE=1 SV=1 | Q924M7 | 0 | 31.324 | 14 | 5 | 22 | 5 | 423 | 46.5 | 5.95 | 49.74 | |
| Mannosyl-oligosaccharide 1,2-alpha-mannosidase IA OS=Mus musculus OX=10090 GN=Man1a1 PE=1 SV=1 | P45700 | 0 | 22.276 | 5 | 2 | 12 | 2 | 655 | 73.2 | 6.81 | 25.49 | 2.926 |
| Mannosyl-oligosaccharide 1,2-alpha-mannosidase IB OS=Mus musculus OX=10090 GN=Man1a2 PE=1 SV=1 | P39098 | 0 | 18.591 | 8 | 3 | 9 | 3 | 641 | 72.8 | 8.28 | 16.24 | 0.834 |
| Mannosyl-oligosaccharide glucosidase OS=Mus musculus OX=10090 GN=Mogs PE=1 SV=1 | Q80UM7 | 0 | 15.376 | 6 | 3 | 9 | 3 | 834 | 91.8 | 9 | 16.89 | 1.049 |
| MAP7 domain-containing protein 1 OS=Mus musculus OX=10090 GN=Map7d1 PE=1 SV=1 | A2AJI0 | 0 | 31.449 | 6 | 4 | 35 | 4 | 846 | 93.2 | 10.15 | 101.17 | 1.033 |
| Mapk-regulated corepressor-interacting protein 1 OS=Mus musculus OX=10090 GN=Mcrip1 PE=1 SV=1 | Q3UGS4 | 0 | 10.012 | 35 | 2 | 15 | 2 | 97 | 11.1 | 9.14 | 39.34 | 0.875 |
| MARCKS-related protein OS=Mus musculus OX=10090 GN=Marcksl1 PE=1 SV=2 | P28667 | 0 | 53.127 | 59 | 4 | 65 | 4 | 200 | 20.2 | 4.61 | 61.49 | 1.445 |
| Maspardin OS=Mus musculus OX=10090 GN=Spg21 PE=1 SV=1 | Q9CQC8 | 0 | 74.666 | 23 | 5 | 54 | 5 | 308 | 34.9 | 6.28 | 125.18 | 1.32 |
| Matrin-3 OS=Mus musculus OX=10090 GN=Matr3 PE=1 SV=1 | Q8K310 | 0 | 90.555 | 15 | 7 | 71 | 7 | 846 | 94.6 | 6.25 | 190.54 | 0.954 |
| Matrix metalloproteinase-14 OS=Mus musculus OX=10090 GN=Mmp14 PE=2 SV=3 | P53690 | 0 | 25.049 | 7 | 3 | 19 | 3 | 582 | 65.9 | 8.03 | 62.76 | 0.998 |
| Matrix-remodeling-associated protein 7 OS=Mus musculus OX=10090 GN=Mxra7 PE=1 SV=2 | Q9CZH7 | 0 | 9.282 | 12 | 2 | 5 | 2 | 178 | 19.4 | 4.26 | 17.57 | 0.564 |
| Mediator of RNA polymerase II transcription subunit 4 OS=Mus musculus OX=10090 GN=Med4 PE=2 SV=1 | Q9CQA5 | 0 | 7.984 | 10 | 2 | 3 | 2 | 270 | 29.8 | 5.05 | 8.74 | |
| Medium-chain specific acyl-CoA dehydrogenase, mitochondrial OS=Mus musculus OX=10090 GN=Acadm PE=1 SV=1 | P45952 | 0 | 33.177 | 19 | 4 | 31 | 4 | 421 | 46.5 | 8.37 | 69.76 | 1.025 |
| Melanoma inhibitory activity protein 2 OS=Mus musculus OX=10090 GN=Mia2 PE=1 SV=3 | Q91ZV0 | 0 | 27.71 | 4 | 6 | 25 | 6 | 1396 | 156.4 | 4.55 | 67.33 | 0.664 |
| Melanoma-associated antigen B4 OS=Mus musculus OX=10090 GN=Mageb4 PE=1 SV=1 | A2A9R3 | 0.004 | 3.457 | 12 | 2 | 2 | 2 | 556 | 60.9 | 9.16 | 2.16 | |
| Melanoma-associated antigen D1 OS=Mus musculus OX=10090 GN=Maged1 PE=1 SV=1 | Q9QYH6 | 0 | 33.37 | 6 | 2 | 67 | 2 | 775 | 85.6 | 7.5 | 108.94 | 0.945 |
| Membrane primary amine oxidase OS=Mus musculus OX=10090 GN=Aoc3 PE=1 SV=3 | O70423 | 0 | 10.714 | 4 | 2 | 5 | 2 | 765 | 84.5 | 6.42 | 13.11 | 0.881 |
| Membrane-associated progesterone receptor component 1 OS=Mus musculus OX=10090 GN=Pgrmc1 PE=1 SV=4 | O55022 | 0 | 59.285 | 24 | 4 | 78 | 4 | 195 | 21.7 | 4.7 | 224.13 | 0.937 |
| Membrane-associated progesterone receptor component 2 OS=Mus musculus OX=10090 GN=Pgrmc2 PE=1 SV=2 | Q80UU9 | 0 | 49.455 | 21 | 5 | 37 | 5 | 217 | 23.3 | 5.15 | 103.77 | 1.427 |
| Merlin OS=Mus musculus OX=10090 GN=Nf2 PE=1 SV=2 | P46662 | 0 | 15.534 | 4 | 2 | 7 | 2 | 596 | 69.7 | 6.35 | 20.52 | |
| Mesencephalic astrocyte-derived neurotrophic factor OS=Mus musculus OX=10090 GN=Manf PE=1 SV=1 | Q9CXI5 | 0 | 65.388 | 39 | 6 | 125 | 6 | 179 | 20.4 | 8.07 | 342.04 | 0.871 |
| Mesoderm-specific transcript protein OS=Mus musculus OX=10090 GN=Mest PE=2 SV=1 | Q07646 | 0 | 13.619 | 11 | 2 | 4 | 2 | 335 | 38.9 | 9.74 | 6.96 | |
| Metalloreductase STEAP3 OS=Mus musculus OX=10090 GN=Steap3 PE=1 SV=1 | Q8CI59 | 0 | 30.687 | 10 | 3 | 18 | 3 | 488 | 54.7 | 9.22 | 37.92 | 1.05 |
| Metalloreductase STEAP4 OS=Mus musculus OX=10090 GN=Steap4 PE=1 SV=1 | Q923B6 | 0 | 34.493 | 12 | 3 | 27 | 3 | 470 | 53 | 9.17 | 51.47 | 0.477 |
| Metastasis-associated protein MTA1 OS=Mus musculus OX=10090 GN=Mta1 PE=1 SV=1 | Q8K4B0 | 0 | 75.09 | 12 | 6 | 97 | 4 | 715 | 80.7 | 9.29 | 253.05 | 0.806 |
| Metastasis-associated protein MTA2 OS=Mus musculus OX=10090 GN=Mta2 PE=1 SV=1 | Q9R190 | 0 | 59.781 | 11 | 5 | 67 | 4 | 668 | 75 | 9.67 | 157.82 | 1.244 |
| Metastasis-associated protein MTA3 OS=Mus musculus OX=10090 GN=Mta3 PE=1 SV=1 | Q924K8 | 0 | 34.962 | 8 | 3 | 40 | 1 | 591 | 67 | 8.88 | 94.79 | 0.684 |
| Metaxin-1 OS=Mus musculus OX=10090 GN=Mtx1 PE=1 SV=1 | P47802 | 0 | 24.095 | 10 | 2 | 25 | 2 | 317 | 35.6 | 6.18 | 53.22 | 0.921 |
| Methanethiol oxidase OS=Mus musculus OX=10090 GN=Selenbp1 PE=1 SV=2 | P17563 | 0 | 22.871 | 7 | 2 | 18 | 2 | 472 | 52.5 | 6.29 | 40.62 | 0.477 |
| Methionine adenosyltransferase 2 subunit beta OS=Mus musculus OX=10090 GN=Mat2b PE=1 SV=1 | Q99LB6 | 0 | 7 | 11 | 2 | 3 | 2 | 334 | 37.4 | 6.95 | 7.47 | 0.855 |
| Methionine aminopeptidase 1 OS=Mus musculus OX=10090 GN=Metap1 PE=1 SV=1 | Q8BP48 | 0 | 77.085 | 23 | 5 | 73 | 5 | 386 | 43.2 | 7.17 | 153.94 | 0.98 |
| Methionine aminopeptidase 2 OS=Mus musculus OX=10090 GN=Metap2 PE=1 SV=1 | O08663 | 0 | 115.307 | 16 | 4 | 139 | 4 | 478 | 52.9 | 5.82 | 392.88 | 0.827 |
| Methionine synthase OS=Mus musculus OX=10090 GN=Mtr PE=1 SV=1 | A6H5Y3 | 0 | 27.582 | 4 | 3 | 27 | 3 | 1253 | 139 | 5.57 | 74.3 | 0.852 |
| Methionine--tRNA ligase, cytoplasmic OS=Mus musculus OX=10090 GN=Mars PE=1 SV=1 | Q68FL6 | 0 | 45.237 | 7 | 4 | 54 | 4 | 902 | 101.4 | 7.14 | 103.75 | 1.144 |
| Methyl-CpG-binding domain protein 3 OS=Mus musculus OX=10090 GN=Mbd3 PE=1 SV=1 | Q9Z2D8 | 0 | 21.394 | 21 | 4 | 25 | 4 | 285 | 32.1 | 5.82 | 77.58 | 1.499 |
| Methyl-CpG-binding protein 2 OS=Mus musculus OX=10090 GN=Mecp2 PE=1 SV=1 | Q9Z2D6 | 0 | 23.16 | 7 | 3 | 36 | 3 | 484 | 52.3 | 9.96 | 102.41 | 0.964 |
| Methylcrotonoyl-CoA carboxylase beta chain, mitochondrial OS=Mus musculus OX=10090 GN=Mccc2 PE=1 SV=1 | Q3ULD5 | 0 | 46.13 | 13 | 4 | 35 | 4 | 563 | 61.3 | 8 | 82.3 | 1.05 |
| Methylcrotonoyl-CoA carboxylase subunit alpha, mitochondrial OS=Mus musculus OX=10090 GN=Mccc1 PE=1 SV=2 | Q99MR8 | 0 | 15.286 | 5 | 2 | 9 | 2 | 717 | 79.3 | 7.83 | 21.53 | 3.097 |
| Methylmalonate-semialdehyde dehydrogenase [acylating], mitochondrial OS=Mus musculus OX=10090 GN=Aldh6a1 PE=1 SV=1 | Q9EQ20 | 0 | 37.958 | 15 | 5 | 29 | 5 | 535 | 57.9 | 8.07 | 61.79 | 1.019 |
| Methylmalonyl-CoA mutase, mitochondrial OS=Mus musculus OX=10090 GN=Mmut PE=1 SV=2 | P16332 | 0 | 50.299 | 16 | 7 | 30 | 7 | 748 | 82.8 | 6.89 | 78.37 | 0.556 |
| Methylosome protein 50 OS=Mus musculus OX=10090 GN=Wdr77 PE=1 SV=1 | Q99J09 | 0 | 13.845 | 11 | 2 | 6 | 2 | 342 | 36.9 | 5.27 | 19.82 | 0.634 |
| Methylosome subunit pICln OS=Mus musculus OX=10090 GN=Clns1a PE=1 SV=1 | Q61189 | 0 | 56.096 | 27 | 4 | 65 | 4 | 236 | 26 | 4.12 | 146.98 | 0.719 |
| Methylthioribose-1-phosphate isomerase OS=Mus musculus OX=10090 GN=Mri1 PE=1 SV=1 | Q9CQT1 | 0 | 39.473 | 23 | 4 | 34 | 4 | 369 | 39.4 | 5.91 | 70.22 | 0.874 |
| Methylthioribulose-1-phosphate dehydratase OS=Mus musculus OX=10090 GN=Apip PE=1 SV=1 | Q9WVQ5 | 0 | 31.352 | 29 | 4 | 26 | 4 | 241 | 26.9 | 6.9 | 41.54 | 0.764 |
| Methyltransferase-like 26 OS=Mus musculus OX=10090 GN=Mettl26 PE=1 SV=1 | Q9DCS2 | 0 | 37.223 | 16 | 2 | 29 | 2 | 204 | 22.7 | 6.52 | 72.24 | 0.912 |
| Mevalonate kinase OS=Mus musculus OX=10090 GN=Mvk PE=1 SV=1 | Q9R008 | 0 | 20.382 | 15 | 3 | 9 | 3 | 395 | 41.9 | 6.68 | 22.41 | |
| MICAL-like protein 1 OS=Mus musculus OX=10090 GN=Micall1 PE=1 SV=3 | Q8BGT6 | 0 | 16.975 | 8 | 4 | 12 | 4 | 870 | 94 | 6.68 | 21.58 | 0.822 |
| MICAL-like protein 2 OS=Mus musculus OX=10090 GN=Micall2 PE=1 SV=1 | Q3TN34 | 0 | 27.598 | 3 | 2 | 26 | 2 | 1009 | 108.2 | 9.55 | 76.11 | 1.274 |
| MICOS complex subunit MIC13 OS=Mus musculus OX=10090 GN=Micos13 PE=1 SV=1 | Q8R404 | 0 | 23.594 | 29 | 2 | 25 | 2 | 119 | 13.4 | 8.63 | 52.43 | 1.871 |
| MICOS complex subunit Mic19 OS=Mus musculus OX=10090 GN=Chchd3 PE=1 SV=1 | Q9CRB9 | 0 | 45.245 | 31 | 5 | 58 | 5 | 227 | 26.3 | 8.37 | 106.18 | 0.99 |
| MICOS complex subunit Mic27 OS=Mus musculus OX=10090 GN=Apool PE=1 SV=1 | Q78IK4 | 0 | 17.537 | 11 | 2 | 10 | 2 | 265 | 29.2 | 9.31 | 24.19 | 0.918 |
| MICOS complex subunit Mic60 OS=Mus musculus OX=10090 GN=Immt PE=1 SV=1 | Q8CAQ8 | 0 | 174.933 | 24 | 12 | 209 | 12 | 757 | 83.8 | 6.61 | 510.38 | 1.109 |
| Microfibrillar-associated protein 1B OS=Mus musculus OX=10090 GN=Mfap1b PE=1 SV=1 | C0HKD9 | 0 | 27.228 | 9 | 2 | 18 | 2 | 439 | 51.9 | 4.98 | 43.24 | 0.856 |
| Microtubule-actin cross-linking factor 1 OS=Mus musculus OX=10090 GN=Macf1 PE=1 SV=2 | Q9QXZ0 | 0 | 510.459 | 13 | 60 | 382 | 10 | 7354 | 831.4 | 5.43 | 973.64 | 0.905 |
| Microtubule-associated protein 1A OS=Mus musculus OX=10090 GN=Map1a PE=1 SV=2 | Q9QYR6 | 0 | 77.93 | 4 | 8 | 70 | 7 | 2776 | 300 | 5 | 165.55 | 1.079 |
| Microtubule-associated protein 1B OS=Mus musculus OX=10090 GN=Map1b PE=1 SV=2 | P14873 | 0 | 657.287 | 35 | 52 | 723 | 51 | 2464 | 270.1 | 4.83 | 1604.65 | 0.948 |
| Microtubule-associated protein 1S OS=Mus musculus OX=10090 GN=Map1s PE=1 SV=2 | Q8C052 | 0 | 32.837 | 5 | 3 | 32 | 3 | 973 | 102.9 | 7.02 | 86.81 | 0.825 |
| Microtubule-associated protein 2 OS=Mus musculus OX=10090 GN=Map2 PE=1 SV=2 | P20357 | 0 | 39.36 | 6 | 6 | 31 | 6 | 1828 | 199 | 4.91 | 76.14 | 0.727 |
| Microtubule-associated protein 4 OS=Mus musculus OX=10090 GN=Map4 PE=1 SV=3 | P27546 | 0 | 263.506 | 23 | 16 | 401 | 16 | 1125 | 117.4 | 4.98 | 1038.08 | 1.059 |
| Microtubule-associated protein RP/EB family member 1 OS=Mus musculus OX=10090 GN=Mapre1 PE=1 SV=3 | Q61166 | 0 | 138.282 | 37 | 8 | 210 | 8 | 268 | 30 | 5.22 | 489.72 | 0.968 |
| Microtubule-associated protein RP/EB family member 2 OS=Mus musculus OX=10090 GN=Mapre2 PE=1 SV=1 | Q8R001 | 0 | 62.141 | 32 | 5 | 46 | 5 | 326 | 36.9 | 5.38 | 118.55 | 0.792 |
| MIF4G domain-containing protein OS=Mus musculus OX=10090 GN=Mif4gd PE=1 SV=1 | Q3UBZ5 | 0 | 5.863 | 12 | 2 | 2 | 2 | 222 | 25.5 | 5.12 | 2.87 | 1.151 |
| Mimecan OS=Mus musculus OX=10090 GN=Ogn PE=1 SV=1 | Q62000 | 0 | 37.809 | 12 | 3 | 18 | 3 | 298 | 34 | 5.74 | 42.32 | 0.803 |
| Minor histocompatibility antigen H13 OS=Mus musculus OX=10090 GN=Hm13 PE=1 SV=1 | Q9D8V0 | 0 | 16.952 | 9 | 2 | 16 | 2 | 378 | 41.7 | 6.04 | 28.19 | 0.787 |
| Misshapen-like kinase 1 OS=Mus musculus OX=10090 GN=Mink1 PE=1 SV=3 | Q9JM52 | 0 | 21.252 | 5 | 4 | 12 | 1 | 1308 | 147.2 | 7.44 | 29.22 | 0.756 |
| Mitochondrial 10-formyltetrahydrofolate dehydrogenase OS=Mus musculus OX=10090 GN=Aldh1l2 PE=1 SV=2 | Q8K009 | 0 | 281.479 | 33 | 20 | 273 | 19 | 923 | 101.5 | 6.29 | 719.25 | 0.824 |
| Mitochondrial 2-oxoglutarate/malate carrier protein OS=Mus musculus OX=10090 GN=Slc25a11 PE=1 SV=3 | Q9CR62 | 0 | 25.611 | 9 | 2 | 20 | 2 | 314 | 34.1 | 9.94 | 51.27 | 1.11 |
| Mitochondrial amidoxime reducing component 2 OS=Mus musculus OX=10090 GN=Marc2 PE=1 SV=1 | Q922Q1 | 0 | 64.2 | 19 | 5 | 86 | 5 | 338 | 38.2 | 8.68 | 185.23 | 1.221 |
| Mitochondrial carnitine/acylcarnitine carrier protein OS=Mus musculus OX=10090 GN=Slc25a20 PE=1 SV=1 | Q9Z2Z6 | 0 | 13.438 | 13 | 2 | 16 | 2 | 301 | 33 | 9.11 | 18.55 | 0.519 |
| Mitochondrial fission 1 protein OS=Mus musculus OX=10090 GN=Fis1 PE=1 SV=1 | Q9CQ92 | 0 | 77.868 | 28 | 4 | 103 | 4 | 152 | 17 | 8.53 | 246.09 | 0.897 |
| Mitochondrial fission factor OS=Mus musculus OX=10090 GN=Mff PE=1 SV=1 | Q6PCP5 | 0 | 44.813 | 23 | 3 | 49 | 1 | 291 | 32.9 | 6.83 | 108.86 | 0.846 |
| Mitochondrial fission regulator 1-like OS=Mus musculus OX=10090 GN=Mtfr1l PE=1 SV=1 | Q9CWE0 | 0 | 19.956 | 22 | 4 | 11 | 4 | 289 | 31.7 | 6.1 | 22.53 | 0.992 |
| Mitochondrial import inner membrane translocase subunit Tim10 OS=Mus musculus OX=10090 GN=Timm10 PE=1 SV=1 | P62073 | 0 | 46.262 | 42 | 3 | 69 | 3 | 90 | 10.3 | 6.29 | 84.81 | 1.118 |
| Mitochondrial import inner membrane translocase subunit Tim13 OS=Mus musculus OX=10090 GN=Timm13 PE=1 SV=1 | P62075 | 0 | 15.316 | 43 | 2 | 38 | 2 | 95 | 10.5 | 8.18 | 126.69 | 1.07 |
| Mitochondrial import inner membrane translocase subunit Tim23 OS=Mus musculus OX=10090 GN=Timm23 PE=1 SV=1 | Q9WTQ8 | 0 | 24.453 | 18 | 2 | 20 | 2 | 209 | 22 | 9.13 | 48.2 | 0.972 |
| Mitochondrial import inner membrane translocase subunit TIM44 OS=Mus musculus OX=10090 GN=Timm44 PE=1 SV=2 | O35857 | 0 | 35.181 | 18 | 6 | 31 | 6 | 452 | 51.1 | 8.13 | 82.94 | 1.511 |
| Mitochondrial import inner membrane translocase subunit TIM50 OS=Mus musculus OX=10090 GN=Timm50 PE=1 SV=1 | Q9D880 | 0 | 16.147 | 11 | 3 | 21 | 3 | 353 | 39.8 | 8.13 | 41.96 | 0.746 |
| Mitochondrial import inner membrane translocase subunit Tim8 A OS=Mus musculus OX=10090 GN=Timm8a1 PE=1 SV=1 | Q9WVA2 | 0 | 14.901 | 41 | 2 | 9 | 2 | 97 | 11 | 5.16 | 31.8 | 1.064 |
| Mitochondrial import receptor subunit TOM22 homolog OS=Mus musculus OX=10090 GN=Tomm22 PE=1 SV=3 | Q9CPQ3 | 0 | 41.062 | 25 | 3 | 78 | 3 | 142 | 15.5 | 4.34 | 173.08 | 1.001 |
| Mitochondrial import receptor subunit TOM34 OS=Mus musculus OX=10090 GN=Tomm34 PE=1 SV=1 | Q9CYG7 | 0 | 71.518 | 17 | 3 | 88 | 3 | 309 | 34.3 | 9.14 | 288.06 | 1.044 |
| Mitochondrial import receptor subunit TOM40 homolog OS=Mus musculus OX=10090 GN=Tomm40 PE=1 SV=3 | Q9QYA2 | 0 | 44.301 | 12 | 3 | 49 | 3 | 361 | 37.9 | 7.74 | 136.93 | 1.083 |
| Mitochondrial import receptor subunit TOM70 OS=Mus musculus OX=10090 GN=Tomm70 PE=1 SV=2 | Q9CZW5 | 0 | 145.37 | 26 | 12 | 115 | 12 | 611 | 67.5 | 7.53 | 274.46 | 0.97 |
| Mitochondrial intermembrane space import and assembly protein 40 OS=Mus musculus OX=10090 GN=Chchd4 PE=1 SV=1 | Q8VEA4 | 0 | 6.436 | 26 | 2 | 3 | 2 | 139 | 15.5 | 4.32 | 4.34 | |
| Mitochondrial proton/calcium exchanger protein OS=Mus musculus OX=10090 GN=Letm1 PE=1 SV=1 | Q9Z2I0 | 0 | 65.664 | 18 | 7 | 61 | 7 | 738 | 82.9 | 6.52 | 148.61 | 0.824 |
| Mitochondrial Rho GTPase 1 OS=Mus musculus OX=10090 GN=Rhot1 PE=1 SV=1 | Q8BG51 | 0 | 26.935 | 5 | 2 | 35 | 2 | 631 | 72.2 | 6.49 | 90.07 | 1.264 |
| Mitochondrial-processing peptidase subunit alpha OS=Mus musculus OX=10090 GN=Pmpca PE=1 SV=1 | Q9DC61 | 0 | 12.451 | 9 | 3 | 15 | 3 | 524 | 58.2 | 6.83 | 40.54 | 1.05 |
| Mitochondrial-processing peptidase subunit beta OS=Mus musculus OX=10090 GN=Pmpcb PE=1 SV=1 | Q9CXT8 | 0 | 23.341 | 8 | 2 | 15 | 2 | 489 | 54.6 | 6.99 | 40.6 | 1.024 |
| Mitofusin-1 OS=Mus musculus OX=10090 GN=Mfn1 PE=1 SV=3 | Q811U4 | 0 | 17.731 | 4 | 2 | 10 | 2 | 741 | 83.7 | 6.51 | 18.73 | 0.756 |
| Mitogen-activated protein kinase 1 OS=Mus musculus OX=10090 GN=Mapk1 PE=1 SV=3 | P63085 | 0 | 149.948 | 26 | 6 | 130 | 4 | 358 | 41.2 | 6.98 | 320.61 | 0.991 |
| Mitogen-activated protein kinase 14 OS=Mus musculus OX=10090 GN=Mapk14 PE=1 SV=3 | P47811 | 0 | 8.284 | 15 | 3 | 7 | 3 | 360 | 41.3 | 5.88 | 6.8 | 0.673 |
| Mitogen-activated protein kinase 3 OS=Mus musculus OX=10090 GN=Mapk3 PE=1 SV=5 | Q63844 | 0 | 127.991 | 29 | 8 | 90 | 6 | 380 | 43 | 6.61 | 230.79 | 1.146 |
| Mitogen-activated protein kinase 9 OS=Mus musculus OX=10090 GN=Mapk9 PE=1 SV=2 | Q9WTU6 | 0 | 6.94 | 7 | 2 | 4 | 2 | 423 | 48.2 | 5.97 | 5.62 | |
| Mitogen-activated protein kinase kinase kinase 2 OS=Mus musculus OX=10090 GN=Map3k2 PE=1 SV=2 | Q61083 | 0 | 12.85 | 5 | 2 | 5 | 1 | 619 | 69.5 | 8.46 | 11.78 | 2.781 |
| Mitogen-activated protein kinase kinase kinase 3 OS=Mus musculus OX=10090 GN=Map3k3 PE=1 SV=1 | Q61084 | 0 | 22.122 | 6 | 3 | 9 | 2 | 626 | 70.7 | 8.87 | 20.99 | 0.985 |
| Mitogen-activated protein kinase kinase kinase kinase 2 OS=Mus musculus OX=10090 GN=Map4k2 PE=1 SV=1 | Q61161 | 0 | 12.045 | 4 | 2 | 4 | 2 | 821 | 91.2 | 6.46 | 8.29 | |
| Mitogen-activated protein kinase kinase kinase kinase 4 OS=Mus musculus OX=10090 GN=Map4k4 PE=1 SV=1 | P97820 | 0 | 22.598 | 6 | 4 | 12 | 1 | 1233 | 140.5 | 7.47 | 28.93 | |
| Mitotic spindle assembly checkpoint protein MAD1 OS=Mus musculus OX=10090 GN=Mad1l1 PE=1 SV=1 | Q9WTX8 | 0 | 40.912 | 14 | 6 | 35 | 6 | 717 | 83.5 | 5.66 | 59.05 | 0.934 |
| MKI67 FHA domain-interacting nucleolar phosphoprotein OS=Mus musculus OX=10090 GN=Nifk PE=1 SV=1 | Q91VE6 | 0 | 9.167 | 9 | 2 | 6 | 2 | 317 | 36.2 | 10.24 | 14.47 | 0.795 |
| MLV-related proviral Env polyprotein OS=Mus musculus OX=10090 PE=1 SV=3 | P10404 | 0 | 25.481 | 6 | 3 | 23 | 3 | 641 | 69.6 | 7.96 | 56.15 | 0.714 |
| MMS19 nucleotide excision repair protein homolog OS=Mus musculus OX=10090 GN=Mms19 PE=1 SV=1 | Q9D071 | 0 | 36.998 | 7 | 4 | 18 | 4 | 1031 | 113 | 6.21 | 35.91 | 1.452 |
| MOB-like protein phocein OS=Mus musculus OX=10090 GN=Mob4 PE=1 SV=1 | Q6PEB6 | 0 | 43.264 | 23 | 2 | 58 | 2 | 225 | 26 | 5.78 | 120.56 | 1.457 |
| Moesin OS=Mus musculus OX=10090 GN=Msn PE=1 SV=3 | P26041 | 0 | 540.603 | 37 | 18 | 990 | 13 | 577 | 67.7 | 6.6 | 2214.65 | 1.003 |
| Monofunctional C1-tetrahydrofolate synthase, mitochondrial OS=Mus musculus OX=10090 GN=Mthfd1l PE=1 SV=2 | Q3V3R1 | 0 | 115.84 | 22 | 13 | 134 | 13 | 977 | 105.7 | 7.02 | 339.12 | 1.003 |
| MORC family CW-type zinc finger protein 3 OS=Mus musculus OX=10090 GN=Morc3 PE=1 SV=1 | F7BJB9 | 0 | 29.328 | 6 | 3 | 13 | 3 | 942 | 106.5 | 5.68 | 41.89 | 0.943 |
| MORF4 family-associated protein 1 OS=Mus musculus OX=10090 GN=Mrfap1 PE=1 SV=1 | Q9CQL7 | 0 | 24.277 | 26 | 2 | 19 | 2 | 125 | 14.2 | 4.73 | 31.35 | 0.963 |
| Motile sperm domain-containing protein 2 OS=Mus musculus OX=10090 GN=Mospd2 PE=1 SV=2 | Q9CWP6 | 0 | 13.777 | 6 | 2 | 5 | 2 | 518 | 59.8 | 5.67 | 9.5 | 0.757 |
| mRNA cap guanine-N7 methyltransferase OS=Mus musculus OX=10090 GN=Rnmt PE=1 SV=1 | Q9D0L8 | 0 | 75.261 | 16 | 6 | 60 | 6 | 465 | 53.3 | 6.48 | 121.32 | 1.021 |
| mRNA export factor OS=Mus musculus OX=10090 GN=Rae1 PE=1 SV=1 | Q8C570 | 0 | 6.043 | 10 | 2 | 2 | 2 | 368 | 40.9 | 7.83 | 4.55 | |
| Multidrug resistance protein 1A OS=Mus musculus OX=10090 GN=Abcb1a PE=1 SV=3 | P21447 | 0 | 110.019 | 11 | 9 | 121 | 9 | 1276 | 140.6 | 8.85 | 291.28 | 0.638 |
| Multidrug resistance-associated protein 1 OS=Mus musculus OX=10090 GN=Abcc1 PE=1 SV=1 | O35379 | 0 | 15.67 | 4 | 4 | 6 | 4 | 1528 | 171.1 | 7.36 | 15.92 | 1.179 |
| Multifunctional methyltransferase subunit TRM112-like protein OS=Mus musculus OX=10090 GN=Trmt112 PE=1 SV=1 | Q9DCG9 | 0 | 20.038 | 22 | 2 | 14 | 2 | 125 | 14.1 | 5.27 | 24.83 | 0.539 |
| Multifunctional procollagen lysine hydroxylase and glycosyltransferase LH3 OS=Mus musculus OX=10090 GN=Plod3 PE=1 SV=1 | Q9R0E1 | 0 | 112.99 | 21 | 11 | 118 | 11 | 741 | 84.9 | 6.23 | 293.53 | 0.991 |
| Multifunctional protein ADE2 OS=Mus musculus OX=10090 GN=Paics PE=1 SV=4 | Q9DCL9 | 0 | 89.468 | 31 | 10 | 95 | 10 | 425 | 47 | 7.23 | 233.12 | 0.868 |
| Multiple coagulation factor deficiency protein 2 homolog OS=Mus musculus OX=10090 GN=Mcfd2 PE=1 SV=1 | Q8K5B2 | 0 | 46.335 | 52 | 3 | 36 | 3 | 145 | 16.2 | 4.72 | 60.63 | 1.032 |
| Multiple PDZ domain protein OS=Mus musculus OX=10090 GN=Mpdz PE=1 SV=2 | Q8VBX6 | 0 | 40.478 | 5 | 5 | 28 | 5 | 2055 | 218.6 | 5.02 | 65.34 | 1.168 |
| Multivesicular body subunit 12A OS=Mus musculus OX=10090 GN=Mvb12a PE=1 SV=1 | Q78HU3 | 0 | 20.689 | 12 | 2 | 18 | 2 | 271 | 28.7 | 9.11 | 45.98 | 0.861 |
| Muscleblind-like protein 2 OS=Mus musculus OX=10090 GN=Mbnl2 PE=2 SV=2 | Q8C181 | 0 | 13.498 | 9 | 2 | 5 | 2 | 373 | 40.1 | 9 | 12.3 | 0.79 |
| Myb-binding protein 1A OS=Mus musculus OX=10090 GN=Mybbp1a PE=1 SV=2 | Q7TPV4 | 0 | 88.421 | 12 | 10 | 82 | 10 | 1344 | 151.9 | 8.95 | 203.73 | 0.972 |
| Myc box-dependent-interacting protein 1 OS=Mus musculus OX=10090 GN=Bin1 PE=1 SV=1 | O08539 | 0 | 85.272 | 26 | 8 | 97 | 8 | 588 | 64.4 | 5.03 | 232.17 | 0.875 |
| Myelin expression factor 2 OS=Mus musculus OX=10090 GN=Myef2 PE=1 SV=1 | Q8C854 | 0 | 30.158 | 10 | 6 | 30 | 6 | 591 | 63.3 | 8.87 | 55.31 | 1.182 |
| Myeloid cell nuclear differentiation antigen-like protein OS=Mus musculus OX=10090 GN=Mndal PE=1 SV=1 | D0QMC3 | 0 | 10.474 | 6 | 2 | 11 | 2 | 538 | 60.5 | 9.28 | 22.52 | 1.025 |
| Myeloid-derived growth factor OS=Mus musculus OX=10090 GN=Mydgf PE=1 SV=1 | Q9CPT4 | 0 | 21.412 | 15 | 3 | 22 | 3 | 166 | 18 | 6.79 | 60.03 | 0.937 |
| Myoferlin OS=Mus musculus OX=10090 GN=Myof PE=1 SV=2 | Q69ZN7 | 0 | 200.91 | 17 | 26 | 188 | 26 | 2048 | 233.2 | 6.16 | 428.49 | 1.135 |
| Myomegalin OS=Mus musculus OX=10090 GN=Pde4dip PE=1 SV=2 | Q80YT7 | 0 | 7.357 | 1 | 2 | 7 | 1 | 2224 | 250.5 | 5.53 | 11.01 | |
| Myosin light polypeptide 6 OS=Mus musculus OX=10090 GN=Myl6 PE=1 SV=3 | Q60605 | 0 | 137.295 | 55 | 7 | 274 | 1 | 151 | 16.9 | 4.65 | 604.8 | 0.906 |
| Myosin phosphatase Rho-interacting protein OS=Mus musculus OX=10090 GN=Mprip PE=1 SV=2 | P97434 | 0 | 80.313 | 17 | 12 | 75 | 12 | 1024 | 116.3 | 6.21 | 208.73 | 0.953 |
| Myosin regulatory light chain 12B OS=Mus musculus OX=10090 GN=Myl12b PE=1 SV=2 | Q3THE2 | 0 | 176.4 | 54 | 7 | 338 | 5 | 172 | 19.8 | 4.84 | 869.45 | 1.013 |
| Myosin regulatory light polypeptide 9 OS=Mus musculus OX=10090 GN=Myl9 PE=1 SV=3 | Q9CQ19 | 0 | 98.961 | 50 | 6 | 98 | 4 | 172 | 19.8 | 4.92 | 256.42 | 0.853 |
| Myosin-10 OS=Mus musculus OX=10090 GN=Myh10 PE=1 SV=2 | Q61879 | 0 | 1445.812 | 41 | 77 | 1614 | 66 | 1976 | 228.9 | 5.54 | 4442.49 | 0.952 |
| Myosin-11 OS=Mus musculus OX=10090 GN=Myh11 PE=1 SV=1 | O08638 | 0 | 82.178 | 5 | 8 | 151 | 1 | 1972 | 226.9 | 5.45 | 344.67 | |
| Myosin-14 OS=Mus musculus OX=10090 GN=Myh14 PE=1 SV=1 | Q6URW6 | 0 | 51.076 | 4 | 5 | 112 | 3 | 2000 | 228.4 | 5.55 | 291.1 | 0.943 |
| Myosin-9 OS=Mus musculus OX=10090 GN=Myh9 PE=1 SV=4 | Q8VDD5 | 0 | 2737.276 | 53 | 117 | 3765 | 102 | 1960 | 226.2 | 5.66 | 10489.23 | 1.008 |
| Myotrophin OS=Mus musculus OX=10090 GN=Mtpn PE=1 SV=2 | P62774 | 0 | 123.706 | 33 | 4 | 145 | 4 | 118 | 12.9 | 5.52 | 352.64 | 1.017 |
| Myotubularin-related protein 5 OS=Mus musculus OX=10090 GN=Sbf1 PE=1 SV=2 | Q6ZPE2 | 0 | 51.336 | 3 | 3 | 54 | 3 | 1867 | 208.6 | 7.12 | 131.03 | 1.279 |
| Myristoylated alanine-rich C-kinase substrate OS=Mus musculus OX=10090 GN=Marcks PE=1 SV=2 | P26645 | 0 | 133.726 | 75 | 8 | 279 | 8 | 309 | 29.6 | 4.34 | 510.22 | 1.324 |
| N(4)-(beta-N-acetylglucosaminyl)-L-asparaginase OS=Mus musculus OX=10090 GN=Aga PE=1 SV=1 | Q64191 | 0 | 66.015 | 28 | 4 | 175 | 4 | 346 | 37 | 6.44 | 456.01 | 0.96 |
| N(G),N(G)-dimethylarginine dimethylaminohydrolase 1 OS=Mus musculus OX=10090 GN=Ddah1 PE=1 SV=3 | Q9CWS0 | 0 | 62.783 | 32 | 5 | 51 | 5 | 285 | 31.4 | 5.97 | 133.93 | 1.41 |
| N(G),N(G)-dimethylarginine dimethylaminohydrolase 2 OS=Mus musculus OX=10090 GN=Ddah2 PE=1 SV=1 | Q99LD8 | 0 | 108.415 | 30 | 4 | 180 | 4 | 285 | 29.6 | 6.01 | 537.37 | 0.791 |
| N-acetyl-D-glucosamine kinase OS=Mus musculus OX=10090 GN=Nagk PE=1 SV=3 | Q9QZ08 | 0 | 19.299 | 14 | 3 | 13 | 3 | 343 | 37.2 | 5.73 | 16.81 | 0.936 |
| N-acetylgalactosamine kinase OS=Mus musculus OX=10090 GN=Galk2 PE=1 SV=1 | Q68FH4 | 0 | 69.261 | 23 | 6 | 60 | 6 | 458 | 50.5 | 6.9 | 174.1 | 1.696 |
| N-acetylgalactosamine-6-sulfatase OS=Mus musculus OX=10090 GN=Galns PE=1 SV=2 | Q571E4 | 0 | 11.417 | 5 | 2 | 8 | 2 | 520 | 57.6 | 6.52 | 19.1 | 1.226 |
| N-acetylglucosamine-6-phosphate deacetylase OS=Mus musculus OX=10090 GN=Amdhd2 PE=1 SV=1 | Q8JZV7 | 0 | 15.448 | 8 | 2 | 16 | 2 | 409 | 43.5 | 6.23 | 6.74 | 1.945 |
| N-acetylglucosamine-6-sulfatase OS=Mus musculus OX=10090 GN=Gns PE=1 SV=1 | Q8BFR4 | 0 | 59.663 | 16 | 8 | 73 | 8 | 544 | 61.1 | 8.24 | 166.08 | 0.988 |
| N-acetylneuraminate lyase OS=Mus musculus OX=10090 GN=Npl PE=1 SV=1 | Q9DCJ9 | 0 | 12.983 | 18 | 3 | 9 | 3 | 320 | 35.1 | 8.03 | 19.14 | 1.256 |
| N-acylglucosamine 2-epimerase OS=Mus musculus OX=10090 GN=Renbp PE=1 SV=3 | P82343 | 0 | 23.203 | 9 | 2 | 22 | 2 | 430 | 49.7 | 6.06 | 62.91 | 1.471 |
| N-acylneuraminate cytidylyltransferase OS=Mus musculus OX=10090 GN=Cmas PE=1 SV=2 | Q99KK2 | 0 | 16.814 | 13 | 3 | 9 | 3 | 432 | 48 | 8.1 | 22.55 | 1.026 |
| NAD kinase 2, mitochondrial OS=Mus musculus OX=10090 GN=Nadk2 PE=1 SV=2 | Q8C5H8 | 0 | 17.57 | 7 | 2 | 9 | 2 | 452 | 50.8 | 8.07 | 20.73 | |
| NAD(P) transhydrogenase, mitochondrial OS=Mus musculus OX=10090 GN=Nnt PE=1 SV=2 | Q61941 | 0 | 35.814 | 9 | 6 | 21 | 6 | 1086 | 113.8 | 7.64 | 45.74 | 0.855 |
| NAD(P)H-hydrate epimerase OS=Mus musculus OX=10090 GN=Naxe PE=1 SV=1 | Q8K4Z3 | 0 | 35.47 | 17 | 4 | 48 | 4 | 282 | 31 | 7.69 | 130.77 | 0.936 |
| NAD-dependent malic enzyme, mitochondrial OS=Mus musculus OX=10090 GN=Me2 PE=1 SV=1 | Q99KE1 | 0 | 60.106 | 10 | 4 | 57 | 4 | 589 | 65.8 | 7.61 | 151.33 | 1.102 |
| NAD-dependent protein deacetylase sirtuin-2 OS=Mus musculus OX=10090 GN=Sirt2 PE=1 SV=2 | Q8VDQ8 | 0 | 34.613 | 18 | 4 | 41 | 4 | 389 | 43.2 | 5.35 | 100.76 | 1.399 |
| NADH dehydrogenase [ubiquinone] 1 alpha subcomplex subunit 10, mitochondrial OS=Mus musculus OX=10090 GN=Ndufa10 PE=1 SV=1 | Q99LC3 | 0 | 71.007 | 35 | 9 | 57 | 9 | 355 | 40.6 | 7.78 | 130.2 | 0.885 |
| NADH dehydrogenase [ubiquinone] 1 alpha subcomplex subunit 12 OS=Mus musculus OX=10090 GN=Ndufa12 PE=1 SV=2 | Q7TMF3 | 0 | 27.697 | 26 | 4 | 72 | 4 | 145 | 17.1 | 9.36 | 140.98 | 0.94 |
| NADH dehydrogenase [ubiquinone] 1 alpha subcomplex subunit 13 OS=Mus musculus OX=10090 GN=Ndufa13 PE=1 SV=3 | Q9ERS2 | 0 | 28.726 | 37 | 5 | 20 | 5 | 144 | 16.8 | 9.48 | 31.45 | 1.279 |
| NADH dehydrogenase [ubiquinone] 1 alpha subcomplex subunit 5 OS=Mus musculus OX=10090 GN=Ndufa5 PE=1 SV=3 | Q9CPP6 | 0 | 55.08 | 29 | 3 | 48 | 3 | 116 | 13.4 | 8.1 | 147.1 | 1.127 |
| NADH dehydrogenase [ubiquinone] 1 alpha subcomplex subunit 9, mitochondrial OS=Mus musculus OX=10090 GN=Ndufa9 PE=1 SV=2 | Q9DC69 | 0 | 50.603 | 13 | 3 | 53 | 3 | 377 | 42.5 | 9.74 | 107.22 | 0.777 |
| NADH dehydrogenase [ubiquinone] 1 beta subcomplex subunit 10 OS=Mus musculus OX=10090 GN=Ndufb10 PE=1 SV=3 | Q9DCS9 | 0 | 49.087 | 31 | 5 | 60 | 5 | 176 | 21 | 8.03 | 150.83 | 0.855 |
| NADH dehydrogenase [ubiquinone] 1 beta subcomplex subunit 11, mitochondrial OS=Mus musculus OX=10090 GN=Ndufb11 PE=1 SV=2 | O09111 | 0 | 12.746 | 15 | 2 | 15 | 2 | 151 | 17.4 | 5.22 | 20.72 | 0.667 |
| NADH dehydrogenase [ubiquinone] 1 beta subcomplex subunit 8, mitochondrial OS=Mus musculus OX=10090 GN=Ndufb8 PE=1 SV=1 | Q9D6J5 | 0 | 21.66 | 18 | 2 | 21 | 2 | 186 | 21.9 | 6.64 | 39.39 | 0.811 |
| NADH dehydrogenase [ubiquinone] 1 beta subcomplex subunit 9 OS=Mus musculus OX=10090 GN=Ndufb9 PE=1 SV=3 | Q9CQJ8 | 0 | 17.102 | 26 | 2 | 12 | 2 | 179 | 22 | 7.8 | 26.89 | 0.904 |
| NADH dehydrogenase [ubiquinone] flavoprotein 1, mitochondrial OS=Mus musculus OX=10090 GN=Ndufv1 PE=1 SV=1 | Q91YT0 | 0 | 82.234 | 23 | 6 | 99 | 6 | 464 | 50.8 | 8.21 | 284.15 | 0.947 |
| NADH dehydrogenase [ubiquinone] flavoprotein 2, mitochondrial OS=Mus musculus OX=10090 GN=Ndufv2 PE=1 SV=2 | Q9D6J6 | 0 | 53.321 | 23 | 5 | 76 | 5 | 248 | 27.3 | 7.4 | 193.37 | 0.98 |
| NADH dehydrogenase [ubiquinone] iron-sulfur protein 2, mitochondrial OS=Mus musculus OX=10090 GN=Ndufs2 PE=1 SV=1 | Q91WD5 | 0 | 76.888 | 26 | 8 | 65 | 8 | 463 | 52.6 | 6.99 | 147.56 | 1.018 |
| NADH dehydrogenase [ubiquinone] iron-sulfur protein 3, mitochondrial OS=Mus musculus OX=10090 GN=Ndufs3 PE=1 SV=2 | Q9DCT2 | 0 | 44.459 | 21 | 5 | 77 | 5 | 263 | 30.1 | 7.17 | 186.46 | 1.082 |
| NADH dehydrogenase [ubiquinone] iron-sulfur protein 5 OS=Mus musculus OX=10090 GN=Ndufs5 PE=1 SV=3 | Q99LY9 | 0 | 14.384 | 14 | 2 | 34 | 2 | 106 | 12.6 | 8.92 | 85.79 | 0.863 |
| NADH dehydrogenase [ubiquinone] iron-sulfur protein 6, mitochondrial OS=Mus musculus OX=10090 GN=Ndufs6 PE=1 SV=2 | P52503 | 0 | 11.495 | 21 | 2 | 4 | 2 | 116 | 13 | 8.65 | 16.68 | 0.892 |
| NADH dehydrogenase [ubiquinone] iron-sulfur protein 7, mitochondrial OS=Mus musculus OX=10090 GN=Ndufs7 PE=1 SV=1 | Q9DC70 | 0 | 18.749 | 13 | 3 | 15 | 3 | 224 | 24.7 | 9.92 | 44.47 | 0.872 |
| NADH dehydrogenase [ubiquinone] iron-sulfur protein 8, mitochondrial OS=Mus musculus OX=10090 GN=Ndufs8 PE=1 SV=1 | Q8K3J1 | 0 | 42.789 | 24 | 3 | 43 | 3 | 212 | 24 | 6.21 | 98.64 | 0.91 |
| NADH-cytochrome b5 reductase 1 OS=Mus musculus OX=10090 GN=Cyb5r1 PE=1 SV=1 | Q9DB73 | 0 | 43.113 | 12 | 3 | 35 | 3 | 305 | 34.1 | 8.87 | 91.01 | 1.834 |
| NADH-cytochrome b5 reductase 3 OS=Mus musculus OX=10090 GN=Cyb5r3 PE=1 SV=3 | Q9DCN2 | 0 | 103.191 | 31 | 7 | 85 | 7 | 301 | 34.1 | 8.38 | 251.94 | 0.965 |
| NADH-ubiquinone oxidoreductase 75 kDa subunit, mitochondrial OS=Mus musculus OX=10090 GN=Ndufs1 PE=1 SV=2 | Q91VD9 | 0 | 235.399 | 35 | 17 | 296 | 17 | 727 | 79.7 | 5.72 | 801.5 | 0.889 |
| NADP-dependent malic enzyme OS=Mus musculus OX=10090 GN=Me1 PE=1 SV=2 | P06801 | 0 | 126.992 | 32 | 13 | 160 | 13 | 572 | 63.9 | 7.44 | 373.57 | 0.965 |
| NADPH:adrenodoxin oxidoreductase, mitochondrial OS=Mus musculus OX=10090 GN=Fdxr PE=1 SV=1 | Q61578 | 0 | 20.386 | 5 | 2 | 9 | 2 | 494 | 54.2 | 8.66 | 21.3 | 1.178 |
| NADPH--cytochrome P450 reductase OS=Mus musculus OX=10090 GN=Por PE=1 SV=2 | P37040 | 0 | 160.712 | 25 | 10 | 141 | 10 | 678 | 77 | 5.53 | 266.86 | 1.105 |
| N-alpha-acetyltransferase 15, NatA auxiliary subunit OS=Mus musculus OX=10090 GN=Naa15 PE=1 SV=1 | Q80UM3 | 0 | 101.77 | 21 | 12 | 89 | 11 | 865 | 100.9 | 7.62 | 199.53 | 1.164 |
| N-alpha-acetyltransferase 16, NatA auxiliary subunit OS=Mus musculus OX=10090 GN=Naa16 PE=2 SV=1 | Q9DBB4 | 0 | 16.27 | 6 | 3 | 13 | 2 | 864 | 101.2 | 8.35 | 35.71 | |
| N-alpha-acetyltransferase 25, NatB auxiliary subunit OS=Mus musculus OX=10090 GN=Naa25 PE=1 SV=1 | Q8BWZ3 | 0 | 34.07 | 9 | 5 | 17 | 5 | 972 | 111.6 | 6.52 | 40.59 | 0.86 |
| N-alpha-acetyltransferase 30 OS=Mus musculus OX=10090 GN=Naa30 PE=1 SV=2 | Q8CES0 | 0 | 31.446 | 12 | 3 | 47 | 3 | 364 | 39.4 | 5.64 | 84.56 | 1.116 |
| N-alpha-acetyltransferase 35, NatC auxiliary subunit OS=Mus musculus OX=10090 GN=Naa35 PE=1 SV=1 | Q6PHQ8 | 0 | 27.684 | 7 | 3 | 27 | 3 | 725 | 83.3 | 7.3 | 66.79 | 1.278 |
| N-alpha-acetyltransferase 50 OS=Mus musculus OX=10090 GN=Naa50 PE=1 SV=1 | Q6PGB6 | 0 | 7.211 | 17 | 2 | 2 | 2 | 169 | 19.4 | 8.81 | 2.41 | |
| Nardilysin OS=Mus musculus OX=10090 GN=Nrdc PE=1 SV=1 | Q8BHG1 | 0 | 89.125 | 10 | 7 | 85 | 7 | 1161 | 132.8 | 4.87 | 222.36 | 0.863 |
| Nascent polypeptide-associated complex subunit alpha, muscle-specific form OS=Mus musculus OX=10090 GN=Naca PE=1 SV=2 | P70670 | 0 | 103.312 | 3 | 6 | 187 | 6 | 2187 | 220.4 | 9.35 | 426.58 | 0.884 |
| Nck-associated protein 1 OS=Mus musculus OX=10090 GN=Nckap1 PE=1 SV=2 | P28660 | 0 | 56.567 | 11 | 8 | 63 | 8 | 1128 | 128.7 | 6.62 | 152.83 | 0.852 |
| Nck-associated protein 1-like OS=Mus musculus OX=10090 GN=Nckap1l PE=1 SV=1 | Q8K1X4 | 0 | 34.553 | 3 | 2 | 35 | 2 | 1134 | 128.8 | 6.64 | 103.06 | 1.101 |
| NCK-interacting protein with SH3 domain OS=Mus musculus OX=10090 GN=Nckipsd PE=1 SV=2 | Q9ESJ4 | 0 | 28.104 | 7 | 4 | 16 | 4 | 714 | 78.5 | 6.05 | 50.78 | 1.316 |
| Nectin-1 OS=Mus musculus OX=10090 GN=Nectin1 PE=1 SV=3 | Q9JKF6 | 0 | 9.489 | 10 | 2 | 4 | 2 | 515 | 57 | 6.35 | 2.52 | |
| Nectin-2 OS=Mus musculus OX=10090 GN=Nectin2 PE=1 SV=2 | P32507 | 0 | 14.045 | 11 | 3 | 4 | 3 | 530 | 57.3 | 4.86 | 14.42 | 0.809 |
| Nectin-3 OS=Mus musculus OX=10090 GN=Nectin3 PE=1 SV=1 | Q9JLB9 | 0 | 11.606 | 10 | 4 | 6 | 4 | 549 | 60.5 | 6.54 | 6.8 | 1.54 |
| NEDD4-like E3 ubiquitin-protein ligase WWP1 OS=Mus musculus OX=10090 GN=Wwp1 PE=1 SV=2 | Q8BZZ3 | 0 | 6.716 | 3 | 2 | 4 | 2 | 918 | 104.6 | 6.38 | 7.55 | |
| NEDD8 OS=Mus musculus OX=10090 GN=Nedd8 PE=1 SV=2 | P29595 | 0 | 45.547 | 23 | 2 | 51 | 2 | 81 | 9 | 7.25 | 115.02 | 0.964 |
| NEDD8 ultimate buster 1 OS=Mus musculus OX=10090 GN=Nub1 PE=1 SV=2 | P54729 | 0 | 78.214 | 13 | 5 | 69 | 5 | 614 | 70.3 | 5.88 | 182.3 | 0.939 |
| NEDD8-activating enzyme E1 catalytic subunit OS=Mus musculus OX=10090 GN=Uba3 PE=1 SV=2 | Q8C878 | 0 | 54.946 | 17 | 4 | 57 | 4 | 462 | 51.7 | 5.45 | 166.16 | 1.149 |
| NEDD8-activating enzyme E1 regulatory subunit OS=Mus musculus OX=10090 GN=Nae1 PE=1 SV=1 | Q8VBW6 | 0 | 13.036 | 8 | 3 | 10 | 3 | 534 | 60.2 | 5.52 | 23.44 | 1.245 |
| NEDD8-conjugating enzyme Ubc12 OS=Mus musculus OX=10090 GN=Ube2m PE=1 SV=1 | P61082 | 0 | 29.979 | 21 | 3 | 46 | 3 | 183 | 20.9 | 7.69 | 146.93 | 1.069 |
| Negative elongation factor A OS=Mus musculus OX=10090 GN=Nelfa PE=1 SV=1 | Q8BG30 | 0 | 10.557 | 5 | 2 | 3 | 2 | 530 | 57.5 | 9.11 | 6.81 | 1.083 |
| Negative elongation factor E OS=Mus musculus OX=10090 GN=Nelfe PE=1 SV=2 | P19426 | 0 | 8.314 | 11 | 2 | 3 | 2 | 375 | 42.5 | 9.38 | 5.03 | |
| Nesprin-1 OS=Mus musculus OX=10090 GN=Syne1 PE=1 SV=2 | Q6ZWR6 | 0 | 47.813 | 1 | 5 | 25 | 5 | 8799 | 1009.3 | 5.59 | 80.2 | 0.7 |
| Nesprin-2 OS=Mus musculus OX=10090 GN=Syne2 PE=1 SV=2 | Q6ZWQ0 | 0 | 224.128 | 7 | 32 | 133 | 32 | 6874 | 782.2 | 5.33 | 360.04 | 1.225 |
| Nesprin-3 OS=Mus musculus OX=10090 GN=Syne3 PE=1 SV=1 | Q4FZC9 | 0 | 26.104 | 9 | 6 | 13 | 6 | 975 | 112 | 6.33 | 22.88 | 1.109 |
| Nestin OS=Mus musculus OX=10090 GN=Nes PE=1 SV=1 | Q6P5H2 | 0 | 716.384 | 35 | 42 | 869 | 42 | 1864 | 207 | 4.34 | 2426 | 1.109 |
| Netrin-1 OS=Mus musculus OX=10090 GN=Ntn1 PE=1 SV=3 | O09118 | 0 | 14.71 | 6 | 2 | 5 | 2 | 604 | 67.8 | 8.78 | 7.3 | |
| Neudesin OS=Mus musculus OX=10090 GN=Nenf PE=1 SV=1 | Q9CQ45 | 0 | 45.448 | 36 | 5 | 74 | 5 | 171 | 18.9 | 5.27 | 131.05 | 0.928 |
| Neurabin-2 OS=Mus musculus OX=10090 GN=Ppp1r9b PE=1 SV=1 | Q6R891 | 0 | 51.217 | 10 | 4 | 36 | 4 | 817 | 89.5 | 4.92 | 96.95 | 0.636 |
| Neural cell adhesion molecule 1 OS=Mus musculus OX=10090 GN=Ncam1 PE=1 SV=3 | P13595 | 0 | 238.72 | 23 | 14 | 399 | 14 | 1115 | 119.4 | 4.83 | 670.47 | 0.926 |
| Neural cell adhesion molecule L1 OS=Mus musculus OX=10090 GN=L1cam PE=1 SV=1 | P11627 | 0 | 29.874 | 5 | 3 | 23 | 3 | 1260 | 140.9 | 5.97 | 65.56 | 0.96 |
| Neural cell adhesion molecule L1-like protein OS=Mus musculus OX=10090 GN=Chl1 PE=1 SV=2 | P70232 | 0 | 36.189 | 7 | 5 | 20 | 5 | 1209 | 135 | 5.57 | 55.02 | 2.002 |
| Neural Wiskott-Aldrich syndrome protein OS=Mus musculus OX=10090 GN=Wasl PE=1 SV=1 | Q91YD9 | 0 | 17.662 | 12 | 3 | 6 | 3 | 501 | 54.2 | 7.93 | 13.45 | 0.855 |
| Neurobeachin OS=Mus musculus OX=10090 GN=Nbea PE=1 SV=1 | Q9EPN1 | 0 | 38.207 | 3 | 4 | 38 | 3 | 2936 | 326.5 | 6.2 | 94.07 | 0.75 |
| Neurocalcin-delta OS=Mus musculus OX=10090 GN=Ncald PE=1 SV=4 | Q91X97 | 0 | 14.348 | 10 | 2 | 31 | 2 | 193 | 22.2 | 5.35 | 68.41 | 1.503 |
| Neurocan core protein OS=Mus musculus OX=10090 GN=Ncan PE=1 SV=1 | P55066 | 0 | 8.25 | 3 | 2 | 7 | 2 | 1268 | 137.1 | 5.72 | 2.87 | |
| Neurofascin OS=Mus musculus OX=10090 GN=Nfasc PE=1 SV=1 | Q810U3 | 0 | 11.816 | 7 | 6 | 6 | 6 | 1240 | 137.9 | 6.19 | 2.26 | |
| Neurofibromin OS=Mus musculus OX=10090 GN=Nf1 PE=1 SV=1 | Q04690 | 0 | 5.302 | 1 | 2 | 3 | 2 | 2841 | 319.4 | 7.39 | 6.11 | 1.336 |
| Neurofilament heavy polypeptide OS=Mus musculus OX=10090 GN=Nefh PE=1 SV=3 | P19246 | 0 | 65.767 | 7 | 5 | 35 | 3 | 1090 | 116.9 | 5.81 | 63.8 | 0.564 |
| Neurofilament light polypeptide OS=Mus musculus OX=10090 GN=Nefl PE=1 SV=5 | P08551 | 0 | 190.618 | 38 | 18 | 192 | 16 | 543 | 61.5 | 4.64 | 443.45 | 2.075 |
| Neurofilament medium polypeptide OS=Mus musculus OX=10090 GN=Nefm PE=1 SV=4 | P08553 | 0 | 192.415 | 39 | 21 | 138 | 18 | 848 | 95.9 | 4.77 | 298.06 | 1.455 |
| Neurogenic locus notch homolog protein 1 OS=Mus musculus OX=10090 GN=Notch1 PE=1 SV=3 | Q01705 | 0 | 4.839 | 2 | 2 | 3 | 2 | 2531 | 270.7 | 5.1 | 3.41 | |
| Neurogenic locus notch homolog protein 2 OS=Mus musculus OX=10090 GN=Notch2 PE=1 SV=2 | O35516 | 0 | 5.465 | 2 | 2 | 3 | 2 | 2473 | 265.5 | 5.22 | 2.48 | |
| Neurolysin, mitochondrial OS=Mus musculus OX=10090 GN=Nln PE=1 SV=1 | Q91YP2 | 0 | 33.926 | 11 | 5 | 22 | 5 | 704 | 80.4 | 6.44 | 34 | 0.953 |
| Neuromodulin OS=Mus musculus OX=10090 GN=Gap43 PE=1 SV=1 | P06837 | 0 | 69.256 | 27 | 4 | 124 | 4 | 227 | 23.6 | 4.73 | 239.69 | 0.758 |
| Neuron navigator 1 OS=Mus musculus OX=10090 GN=Nav1 PE=1 SV=2 | Q8CH77 | 0 | 20.615 | 3 | 3 | 11 | 3 | 1875 | 202.2 | 8.06 | 33.79 | |
| Neuronal cell adhesion molecule OS=Mus musculus OX=10090 GN=Nrcam PE=1 SV=2 | Q810U4 | 0 | 9.421 | 3 | 2 | 3 | 2 | 1256 | 138.4 | 5.91 | 5.15 | |
| Neuronal proto-oncogene tyrosine-protein kinase Src OS=Mus musculus OX=10090 GN=Src PE=1 SV=4 | P05480 | 0 | 20.896 | 10 | 4 | 20 | 3 | 541 | 60.6 | 7.84 | 37.43 | 0.993 |
| Neuropilin-1 OS=Mus musculus OX=10090 GN=Nrp1 PE=1 SV=2 | P97333 | 0 | 56.925 | 10 | 6 | 44 | 6 | 923 | 102.9 | 5.9 | 131.22 | 0.765 |
| Neuroplastin OS=Mus musculus OX=10090 GN=Nptn PE=1 SV=3 | P97300 | 0 | 32.738 | 17 | 4 | 32 | 4 | 397 | 44.3 | 7.74 | 49.56 | 1.065 |
| Neurosecretory protein VGF OS=Mus musculus OX=10090 GN=Vgf PE=1 SV=1 | Q0VGU4 | 0 | 8.273 | 7 | 2 | 3 | 2 | 617 | 68.2 | 4.7 | 4.47 | |
| Neutral alpha-glucosidase AB OS=Mus musculus OX=10090 GN=Ganab PE=1 SV=1 | Q8BHN3 | 0 | 176.216 | 21 | 16 | 269 | 16 | 944 | 106.8 | 6.06 | 581.71 | 0.923 |
| Neutral amino acid transporter A OS=Mus musculus OX=10090 GN=Slc1a4 PE=1 SV=1 | O35874 | 0 | 13.823 | 7 | 2 | 10 | 2 | 532 | 56 | 5.87 | 17.6 | 1.053 |
| Neutral cholesterol ester hydrolase 1 OS=Mus musculus OX=10090 GN=Nceh1 PE=1 SV=1 | Q8BLF1 | 0 | 46.583 | 19 | 4 | 35 | 4 | 408 | 45.7 | 7.05 | 61.04 | 1.238 |
| Neutrophil cytosol factor 1 OS=Mus musculus OX=10090 GN=Ncf1 PE=1 SV=3 | Q09014 | 0 | 17.357 | 10 | 3 | 29 | 3 | 390 | 44.6 | 9.01 | 81.89 | 1.318 |
| Neutrophil cytosol factor 4 OS=Mus musculus OX=10090 GN=Ncf4 PE=1 SV=2 | P97369 | 0 | 35.242 | 9 | 3 | 25 | 3 | 339 | 38.7 | 6.21 | 73.87 | 1.659 |
| NF-kappa-B essential modulator OS=Mus musculus OX=10090 GN=Ikbkg PE=1 SV=2 | O88522 | 0 | 17.819 | 12 | 3 | 14 | 3 | 412 | 47.9 | 5.85 | 36.41 | 1.194 |
| NHL repeat-containing protein 3 OS=Mus musculus OX=10090 GN=Nhlrc3 PE=1 SV=1 | Q8CCH2 | 0 | 14.833 | 7 | 2 | 7 | 2 | 347 | 38.2 | 6.23 | 15.24 | 0.913 |

| | | | | | | | | | | | | | | | |
|---|---|---|---|---|---|---|---|---|---|---|---|---|---|---|---|
| Q6IR34-3 | 1.061 | 0.085424656 | 0.962674291 | 71.1 | 94.1 | 105.9 | 34.71 | 25.72 | High | High | High | High | Peak Found | High | High |
| Q60575-3 | 1.241 | 0.311503115 | 0.895647774 | 54.97 | 108.8 | 91.2 | 28.66 | 50.91 | Not Found | Peak Found | Peak Found | High | Not Found | Not Found | Peak Found |
| Q9QXZ0-3 | 0.917 | -0.125006361 | 0.990475871 | 49.74 | 90.7 | 109.3 | 70.37 | 40.11 | High | High | High | High | Peak Found | Peak Found | High |
| Q7TSJ2-3 | | | | | | | | | High | Not Found | Not Found | High | Peak Found | Not Found | Not Found |
| Q8CI51-3 | 0.57 | -0.810966176 | 0.663199077 | 38.15 | 112.7 | 87.3 | 54.03 | 87.59 | Not Found | Peak Found | High | High | Not Found | High | Peak Found |
| Q9D236-1 | 0.951 | -0.072482754 | 0.991386436 | 67.54 | 91.9 | 108.1 | 50.34 | 62.46 | High | Peak Found | High | Peak Found | Peak Found | Peak Found | High |
| Q8BYK6-3 | 0.465 | -1.104697379 | 0.388331066 | 27.96 | 63.5 | 136.5 | 92.96 | | High | High | High | High | High | High | Not Found |
| Q5SSL4-4 | 0.992 | -0.011587974 | 0.999691687 | 61.54 | 102.1 | 97.9 | 55.88 | 41.84 | High | High | High | High | High | High | High |
| Q8C8R3-4 | 0.963 | -0.054392297 | 0.969361739 | 82.63 | 92.7 | 107.3 | 41.02 | 53.99 | High | High | High | High | High | High | High |
| A2CG49-4 | | | | | | | | | High | Not Found | Not Found | Not Found | Not Found | Not Found | Not Found |
| Q6PCP5-4 | | | | | | | | | Not Found | High | Not Found | Not Found | Not Found | Not Found | Not Found |
| Q05860-5 | | | | | | | | | Not Found | High | Not Found | Not Found | Not Found | Not Found | Not Found |
| Q7M6Y3-5 | 1.068 | 0.094911647 | 0.849159812 | 87.05 | 101.9 | 98.1 | 43.22 | 36.71 | High | High | High | High | High | High | Peak Found |
| Q9Z0H4-6 | 1.154 | 0.206643224 | 0.893249611 | 29.73 | 100.1 | 99.9 | 27.96 | 21.51 | Not Found | High | High | High | Not Found | High | High |
| Q9ET54-6 | 0.939 | -0.090802937 | 0.895647774 | 60.94 | 101.8 | 98.2 | 35.57 | 20.13 | High | High | High | High | High | High | High |
| Q9WV92-7 | 1.033 | 0.046840254 | 0.983938563 | 87.95 | 86.8 | 113.2 | 76.58 | 58.66 | High | Peak Found | High | High | High | Peak Found | Peak Found |
| Q61595-9 | 0.893 | -0.16326792 | 0.745941542 | 62.33 | 97.2 | 102.8 | 48.65 | 49.66 | High | High | High | High | High | High | High |
| Q9Z1G4-2 | 1.168 | 0.224040274 | 0.768272619 | 77.99 | 100.4 | 99.6 | 93.71 | 48.07 | High | High | High | High | High | High | High |
| Q61738-5 | 1.795 | 0.843983844 | 0.016400621 | 88.69 | 136.9 | 63.1 | 68.14 | 81.61 | High | High | High | High | High | High | High |
| Q61584-3 | 1.076 | 0.105678078 | 0.941077092 | 83.69 | 105.4 | 94.6 | 39.81 | 38.28 | High | High | High | High | Peak Found | High | High |
| Q99P72-5 | | | | | | | | | Not Found | Peak Found | Not Found | High | Not Found | Not Found | Not Found |
| Q9Z204-2 | 0.991 | -0.013043037 | 0.999000066 | 75.1 | 93.5 | 106.5 | 56.44 | 43.87 | High | High | High | High | High | High | High |
| O88935-1 | | | | | | | | | High | Not Found | Not Found | Not Found | Not Found | Not Found | Not Found |
| Q64487-10 | 1.35 | 0.432959407 | 0.845187331 | 196.05 | 114.9 | 85.1 | 112.8 | | High | Peak Found | High | High | Not Found | High | Not Found |
| P52480-2 | 1.259 | 0.332278283 | 0.136314678 | 54.58 | 111.1 | 88.9 | 36.02 | 28.69 | High | High | High | High | High | High | High |
| P70257-1 | 1.245 | 0.316145742 | 0.893756428 | 97.74 | 108.6 | 91.4 | 68.97 | 89.74 | High | High | High | Not Found | High | High | High |
| Q9QXS1-3 | 1.14 | 0.189033824 | 0.717531582 | 48.01 | 104.6 | 95.4 | 15.21 | 48.91 | High | High | High | High | High | High | High |
| O35226-2 | 1.225 | 0.292781749 | 0.893756428 | 121.81 | 106.1 | 93.9 | 98.59 | 97.68 | Peak Found | Peak Found | High | High | High | Peak Found | High |
| Q60605-2 | 1.259 | 0.332278283 | 0.070247886 | 41.43 | 109.4 | 90.6 | 21.94 | 20.6 | Not Found | High | High | High | Not Found | High | High |
| Q8BU30 | 0.954 | -0.067938829 | 0.95979474 | 73.14 | 91.3 | 108.7 | 79.19 | 83.63 | High | High | High | High | High | High | High |
| Q8BIJ6 | 0.942 | -0.086201035 | 0.984581306 | 58.17 | 91.1 | 108.9 | 72.57 | 41.77 | High | High | High | High | Peak Found | High | High |
| P58044 | 0.762 | -0.392137097 | 0.350356073 | 49.18 | 88.7 | 111.3 | 80.89 | 63.82 | Peak Found | High | High | High | Peak Found | High | High |
| Q9CX00 | 1.095 | 0.13093087 | 0.962674291 | 27.47 | 104.5 | 95.5 | 26.53 | 4.27 | Not Found | High | High | High | Not Found | High | High |
| Q02257 | 0.767 | -0.382701517 | 0.820983453 | 50.07 | 85.9 | 114.1 | 31.14 | 9.99 | Peak Found | High | High | High | High | Peak Found | Peak Found |
| Q5DTX6 | 1.004 | 0.005759269 | 0.995586491 | 15.07 | 100.3 | 99.7 | 84.04 | 6.77 | Not Found | High | High | Peak Found | Not Found | Peak Found | Not Found |
| Q8BNW9 | | | | | | | | | Not Found | Not Found | High | Not Found | Not Found | Not Found | Not Found |
| Q6IFX2 | | | | | | | | | High | Not Found | Peak Found | Not Found | Not Found | Not Found | Not Found |
| Q9Z2T6 | 3.028 | 1.598365205 | 6.4532E-05 | 79.59 | 136.6 | 63.4 | 77.76 | 41.55 | Not Found | High | Peak Found | Peak Found | Peak Found | Peak Found | Peak Found |
| Q922U2 | 0.039 | -4.680382066 | 5.68375E-15 | 0.08 | 7.5 | 192.5 | 66.2 | | High | Peak Found | Not Found | Peak Found | Peak Found | Not Found | Peak Found |
| P50446 | | | | | | | | | High | Not Found | Not Found | Not Found | Not Found | Not Found | Not Found |
| Q9DCV7 | 1.225 | 0.292781749 | 0.692949529 | 83.97 | 115.4 | 84.6 | 60.57 | 83.31 | Peak Found | High | High | High | High | High | High |
| Q8BGZ7 | | | | | | | | | Not Found | Not Found | Not Found | Not Found | Not Found | Not Found | Not Found |
| O54983 | 0.806 | -0.311148256 | 0.221725692 | 51.59 | 97.7 | 102.3 | 47.22 | 64.82 | High | High | High | High | High | High | High |
| Q60749 | 0.908 | -0.139235797 | 0.9583339 | 83.91 | 98.2 | 101.8 | 61.19 | 88.18 | High | High | High | High | Peak Found | High | High |
| Q9R226 | 0.966 | -0.049904906 | 0.999900066 | 97.36 | 98.3 | 101.7 | 123.43 | 72.17 | Peak Found | Peak Found | High | Peak Found | Peak Found | Peak Found | High |
| Q6ZPU9 | 0.999 | -0.001443417 | 0.995586491 | 88.05 | 106.3 | 93.7 | 55.81 | 58.05 | High | High | High | High | High | High | High |
| Q80W68 | 1.162 | 0.216610069 | 0.882906899 | 14.74 | 107.5 | 92.5 | 20.68 | 8.21 | Peak Found | High | High | High | Not Found | Peak Found | High |
| P28738 | | | | | | | | | Peak Found | Not Found | Peak Found | Not Found | High | High | Not Found |
| O88447 | 1.037 | 0.052415894 | 0.947492375 | 84.76 | 107.4 | 92.6 | 32.75 | 42.94 | High | High | High | High | High | High | High |
| Q9DBS5 | 0.429 | -1.220950447 | 0.18034847 | 22.99 | 42.4 | 157.6 | 99.17 | 1.5 | High | Peak Found | High | High | High | Not Found | High |
| Q61768 | 1.083 | 0.115033243 | 0.777814598 | 87.52 | 104 | 96 | 10.38 | 4.4 | High | High | High | High | High | High | High |
| P70188 | 1.388 | 0.473007568 | 0.745269621 | 71.59 | 115.4 | 84.6 | 41.71 | | Peak Found | High | High | High | High | High | Not Found |
| Q9EQW7 | 0.361 | -1.469929258 | 0.026666282 | 4.63 | 58 | 142 | 27.31 | 29.93 | Not Found | Peak Found | Peak Found | Peak Found | Not Found | Peak Found | High |
| B1AVY7 | | | | | | | | | Not Found | High | Peak Found | High | Peak Found | High | Not Found |
| Q60575 | 0.685 | -0.545824107 | 0.782907189 | 39.57 | 114.2 | 85.8 | 57.46 | 101.38 | Not Found | High | Not Found | Peak Found | Not Found | Not Found | Not Found |
| O35071 | 1.068 | 0.094911647 | 0.959299542 | 75.47 | 130.3 | 69.7 | 50.11 | 86.44 | Peak Found | High | High | High | Peak Found | High | Peak Found |
| Q9QXL2 | 1.234 | 0.303342394 | 0.764830559 | 87.82 | 113.9 | 86.1 | 49.21 | 91.3 | High | High | High | High | High | High | Peak Found |
| P28741 | 0.83 | -0.268816758 | 0.791619886 | 54.95 | 90 | 110 | 40.53 | 19.97 | High | High | High | High | High | High | High |
| Q61771 | | | | | | | | | Peak Found | High | High | Peak Found | High | Peak Found | High |
| Q9D083 | | | | | | | | | Not Found | High | Not Found | High | Peak Found | Peak Found | Peak Found |
| Q8BX02 | 1.02 | 0.028569152 | 0.994561598 | 102.55 | 105.1 | 94.9 | 8.5 | 11.8 | High | High | High | High | Peak Found | High | High |
| Q9CPU0 | 0.829 | -0.270555993 | 0.849159812 | 35.13 | 106.4 | 93.6 | 30.23 | 16.09 | Not Found | High | High | High | Peak Found | High | High |
| Q99KP3 | 1.126 | 0.171206827 | 0.957635368 | 100.83 | 115.7 | 84.3 | 29.09 | 79.97 | Peak Found | High | High | High | Peak Found | High | High |
| Q61033 | 0.992 | -0.011587974 | 0.999900066 | 50.29 | 99.2 | 100.8 | 68.74 | 15.69 | High | High | High | High | Peak Found | High | High |
| Q61029 | 1.008 | 0.011495639 | 0.995586491 | 44.34 | 98.1 | 101.9 | 53.15 | 49.5 | High | High | High | High | High | High | High |
| P14733 | 1.017 | 0.024319679 | 0.969674625 | 103.79 | 103.2 | 96.8 | 64.72 | 39.8 | High | High | High | High | High | High | High |
| P21619 | 1.022 | 0.031395196 | 0.962674291 | 71.88 | 100.5 | 99.5 | 12.26 | 29.35 | High | High | High | High | High | High | High |
| P19137 | 0.767 | -0.382701517 | 0.87152606 | 1.65 | 86.8 | 113.2 | 2.67 | | Not Found | High | Not Found | High | Not Found | Peak Found | Peak Found |
| Q60675 | 1.102 | 0.140124224 | 0.792829069 | 82.84 | 108.5 | 91.5 | 24.61 | 59.35 | High | High | High | High | Peak Found | High | High |
| P97927 | 1.279 | 0.355016264 | 0.064070316 | 73.1 | 99.1 | 100.9 | 59.28 | 51.18 | High | High | High | High | High | High | High |
| Q61001 | 1.011 | 0.015782997 | 0.995586491 | 85.33 | 112.9 | 87.1 | 52.93 | 82.8 | Not Found | Peak Found | High | High | Peak Found | Peak Found | Peak Found |
| P02469 | 1.167 | 0.222804561 | 0.350356073 | 77.95 | 116.2 | 83.8 | 54.12 | 51.94 | High | High | High | High | High | High | High |
| Q61292 | 0.96 | -0.058893689 | 0.962674291 | 97.11 | 98.7 | 101.3 | 47.74 | 17.02 | High | High | High | High | High | High | High |
| P02468 | 1.107 | 0.146655222 | 0.67254563 | 90.41 | 106.7 | 93.3 | 24.01 | 26.84 | High | High | High | High | High | High | High |
| Q9JJK2 | 0.931 | -0.103146927 | 0.994561598 | 34.15 | 96.4 | 103.6 | 46.41 | 37.07 | High | High | Not Found | High | High | Peak Found | Peak Found |
| Q8K0C4 | 0.68 | -0.556393349 | 0.633056096 | 49.82 | 75.7 | 124.3 | 107.6 | 49.58 | High | Peak Found | Peak Found | High | High | High | Not Found |
| Q8BLN5 | 0.737 | -0.440263476 | 0.734744981 | 64.02 | 83.9 | 116.1 | 65.81 | 59.33 | High | High | High | High | High | High | High |
| Q6ZQ58 | 1.127 | 0.172487516 | 0.895647774 | 50.64 | 113.3 | 86.7 | 40.34 | 42.64 | Not Found | High | High | High | Not Found | High | High |
| Q6A0A2 | 0.662 | -0.595096878 | 0.340585216 | 45.97 | 64.3 | 135.7 | 38.78 | 48.38 | Not Found | High | High | High | Not Found | High | Peak Found |
| Q9Z1R2 | 0.787 | -0.345564459 | 0.480321306 | 60.48 | 109.6 | 90.4 | 31.29 | 36.18 | High | High | High | High | High | High | High |
| Q8CG19 | | | | | | | | | Peak Found | High | High | High | Not Found | High | High |
| O08999 | 1.163 | 0.217851097 | 0.949775913 | 127.98 | 117.9 | 82.1 | 152.39 | 173.67 | Peak Found | Peak Found | High | Peak Found | High | Peak Found | High |
| Q8K4G1 | 1.061 | 0.085424656 | 0.962674291 | 103.62 | 108.8 | 91.2 | 122.16 | 131.71 | Not Found | High | High | High | High | High | High |
| P70202 | 0.824 | -0.279283757 | 0.764830559 | 29.38 | 90.4 | 109.6 | 47.52 | 26.47 | High | High | High | High | High | High | High |
| O89017 | 1.377 | 0.461528559 | 0.003194653 | 99.13 | 125.7 | 74.3 | 68.43 | 54.79 | High | High | High | High | High | High | High |
| Q6DVA0 | 0.642 | -0.639354798 | 0.724234148 | 26.54 | 78.2 | 121.8 | 99.14 | 101.72 | High | Not Found | High | Not Found | High | Not Found | Peak Found |
| Q80Y17 | | | | | | | | | Not Found | High | High | High | Peak Found | High | High |
| Q8R4U7 | 1.126 | 0.171206827 | 0.911276781 | 103.94 | 90.4 | 109.6 | 39.37 | 35.36 | High | High | High | High | High | High | High |
| Q9JHQ5 | 1.215 | 0.280956314 | 0.559509145 | 111.6 | 130.4 | 69.6 | 49.81 | 47.76 | High | High | High | High | High | High | High |
| Q9JIA1 | | | | | | | | | High | Not Found | Not Found | Not Found | Not Found | Not Found | Not Found |
| Q6PB66 | 0.964 | -0.052894948 | 0.969361739 | 80.99 | 97.7 | 102.3 | 27.94 | 47.87 | High | High | High | High | High | High | High |
| Q8BVU0 | 0.802 | -0.318325858 | 0.911276781 | 38.08 | 106.5 | 93.5 | 15.63 | 40.91 | High | High | High | High | High | High | Peak Found |
| Q921G6 | 0.897 | -0.15682011 | 0.976368403 | 88.16 | 90.9 | 109.1 | 81.06 | 84.39 | Peak Found | High | High | High | Peak Found | High | High |
| Q3UZ39 | 0.716 | -0.481968507 | 0.604385132 | 43.34 | 89.6 | 110.4 | 55.09 | 65.19 | Peak Found | High | High | High | High | High | High |
| Q91WK0 | 1.013 | 0.018634174 | 0.995586491 | 92.23 | 87.7 | 112.3 | 85.8 | 79.26 | High | High | High | High | High | High | High |
| O88520 | 0.696 | -0.522840789 | 0.760047513 | 0 | 82.1 | 117.9 | | | Not Found | High | Peak Found | High | Not Found | High | Not Found |
| Q80VQ1 | | | | | | | | | Not Found | Not Found | Not Found | Not Found | Not Found | Not Found | Not Found |
| Q8K1T1 | 1.38 | 0.464668267 | 0.687904104 | 51.97 | 116 | 84 | 62.25 | 48.9 | Peak Found | High | High | High | Peak Found | Peak Found | High |
| Q9CRC8 | 1.046 | 0.064882852 | 0.976368403 | 55.41 | 107.3 | 92.7 | 49.73 | 64.69 | High | High | High | High | Peak Found | High | Peak Found |
| Q505F5 | 1.087 | 0.12035194 | 0.917028689 | 86.42 | 107.8 | 92.2 | 36.48 | 58.69 | Peak Found | High | High | High | Peak Found | High | High |
| Q3UGP9 | | | | | | | | | Not Found | High | Peak Found | High | Not Found | Peak Found | Not Found |
| Q922Q8 | 0.817 | -0.291592017 | 0.196576563 | 60.38 | 96.6 | 103.4 | 30.47 | 35.85 | High | High | High | High | High | High | High |
| Q8BMJ2 | 0.984 | -0.023269779 | 0.995586491 | 76.25 | 98.8 | 101.2 | 37.92 | 35.19 | High | High | High | High | High | High | High |
| Q8C129 | 1.085 | 0.117695043 | 0.976368403 | 58.78 | 133.2 | 66.8 | 73.47 | 77.11 | High | High | High | High | High | High | High |

| | | | | | | | | | | |
|---|---|---|---|---|---|---|---|---|---|---|
| NHS-like protein 2 OS=Mus musculus OX=10090 GN=Nhsl2 PE=1 SV=2 | B1AXH1 | 0 | 19.269 | 3 | 2 | 10 | 2 | 1219 | 132.3 | [illegible] |
| Niban-like protein 1 OS=Mus musculus OX=10090 GN=Fam129b PE=1 SV=2 | Q8R1F1 | 0 | 166.675 | 29 | 11 | 150 | 11 | 749 | 84.8 | [illegible] |
| Nibrin OS=Mus musculus OX=10090 GN=Nbn PE=1 SV=1 | Q9R207 | 0 | 22.686 | 3 | 2 | 19 | 2 | 751 | 83.7 | [illegible] |
| Nicalin OS=Mus musculus OX=10090 GN=Ncln PE=1 SV=2 | Q8VCM8 | 0 | 17.976 | 5 | 3 | 11 | 3 | 563 | 62.9 | [illegible] |
| Nicastrin OS=Mus musculus OX=10090 GN=Ncstn PE=1 SV=3 | P57716 | 0 | 46.262 | 4 | 2 | 59 | 2 | 708 | 78.4 | [illegible] |
| Nicotinamide phosphoribosyltransferase OS=Mus musculus OX=10090 GN=Nampt PE=1 SV=1 | Q99KQ4 | 0 | 125.149 | 26 | 8 | 138 | 8 | 491 | 55.4 | [illegible] |
| Nicotinate-nucleotide pyrophosphorylase [carboxylating] OS=Mus musculus OX=10090 GN=Qprt PE=1 SV=1 | Q91X91 | 0 | 19.635 | 10 | 2 | 12 | 2 | 299 | 31.5 | [illegible] |
| Nidogen-1 OS=Mus musculus OX=10090 GN=Nid1 PE=1 SV=2 | P10493 | 0 | 132.718 | 17 | 15 | 148 | 15 | 1245 | 136.5 | [illegible] |
| Nidogen-2 OS=Mus musculus OX=10090 GN=Nid2 PE=1 SV=2 | O88322 | 0 | 68.97 | 10 | 9 | 99 | 9 | 1403 | 153.8 | [illegible] |
| Nik-related protein kinase OS=Mus musculus OX=10090 GN=Nrk PE=1 SV=2 | Q9R0G8 | 0 | 16.179 | 2 | 2 | 24 | 2 | 1455 | 163.5 | [illegible] |
| Nischarin OS=Mus musculus OX=10090 GN=Nisch PE=1 SV=2 | Q80TM9 | 0 | 35.649 | 6 | 5 | 18 | 5 | 1593 | 174.9 | [illegible] |
| NLR family member X1 OS=Mus musculus OX=10090 GN=Nlrx1 PE=1 SV=1 | Q3TL44 | 0 | 7.512 | 3 | 2 | 3 | 2 | 975 | 107.8 | [illegible] |
| N-myc-interactor OS=Mus musculus OX=10090 GN=Nmi PE=1 SV=1 | O35309 | 0 | 22.303 | 16 | 4 | 13 | 4 | 314 | 35.2 | [illegible] |
| Nodal modulator 1 OS=Mus musculus OX=10090 GN=Nomo1 PE=1 SV=1 | Q6GQT9 | 0 | 123.691 | 17 | 11 | 98 | 11 | 1214 | 133.3 | [illegible] |
| Non-histone chromosomal protein HMG-14 OS=Mus musculus OX=10090 GN=Hmgn1 PE=1 SV=2 | P18608 | 0 | 24.529 | 39 | 5 | 17 | 5 | 96 | 10.1 | [illegible] |
| Non-POU domain-containing octamer-binding protein OS=Mus musculus OX=10090 GN=Nono PE=1 SV=3 | Q99K48 | 0 | 59.297 | 19 | 5 | 74 | 5 | 473 | 54.5 | [illegible] |
| Non-specific lipid-transfer protein OS=Mus musculus OX=10090 GN=Scp2 PE=1 SV=3 | P32020 | 0 | 91.525 | 18 | 6 | 93 | 6 | 547 | 59.1 | [illegible] |
| NPC intracellular cholesterol transporter 2 OS=Mus musculus OX=10090 GN=Npc2 PE=1 SV=1 | Q9Z0J0 | 0 | 17.316 | 23 | 3 | 13 | 3 | 149 | 16.4 | [illegible] |
| NSFL1 cofactor p47 OS=Mus musculus OX=10090 GN=Nsfl1c PE=1 SV=1 | Q9CZ44 | 0 | 89.599 | 39 | 8 | 139 | 8 | 370 | 40.7 | [illegible] |
| NT-3 growth factor receptor OS=Mus musculus OX=10090 GN=Ntrk3 PE=1 SV=1 | Q6VNS1 | 0 | 5.071 | 4 | 2 | 2 | 2 | 825 | 92.7 | [illegible] |
| N-terminal kinase-like protein OS=Mus musculus OX=10090 GN=Scyl1 PE=1 SV=1 | Q9EQC5 | 0 | 25.493 | 8 | 4 | 19 | 4 | 806 | 89.1 | [illegible] |
| N-terminal Xaa-Pro-Lys N-methyltransferase 1 OS=Mus musculus OX=10090 GN=Ntmt1 PE=1 SV=3 | Q8R2U4 | 0 | 25.999 | 15 | 2 | 31 | 2 | 223 | 25.4 | [illegible] |
| Nuclear autoantigenic sperm protein OS=Mus musculus OX=10090 GN=Nasp PE=1 SV=2 | Q99MD9 | 0 | 90.689 | 25 | 12 | 53 | 12 | 773 | 83.9 | [illegible] |
| Nuclear cap-binding protein subunit 1 OS=Mus musculus OX=10090 GN=Ncbp1 PE=1 SV=2 | Q3UYV9 | 0 | 39.86 | 7 | 4 | 17 | 4 | 790 | 91.9 | [illegible] |
| Nuclear cap-binding protein subunit 2 OS=Mus musculus OX=10090 GN=Ncbp2 PE=1 SV=1 | Q9CQ49 | 0 | 11.052 | 9 | 2 | 6 | 2 | 156 | 18 | [illegible] |
| Nuclear cap-binding protein subunit 3 OS=Mus musculus OX=10090 GN=Ncbp3 PE=1 SV=1 | Q8BZR9 | 0 | 14.885 | 4 | 2 | 11 | 2 | 615 | 70 | [illegible] |
| Nuclear distribution protein nudE homolog 1 OS=Mus musculus OX=10090 GN=Nde1 PE=1 SV=1 | Q9CZA6 | 0 | 8.182 | 9 | 2 | 3 | 1 | 344 | 38.5 | [illegible] |
| Nuclear distribution protein nudE-like 1 OS=Mus musculus OX=10090 GN=Ndel1 PE=1 SV=2 | Q9ERR1 | 0 | 22.707 | 12 | 3 | 10 | 2 | 345 | 38.3 | [illegible] |
| Nuclear factor 1 A-type OS=Mus musculus OX=10090 GN=Nfia PE=1 SV=1 | Q02780 | 0 | 7.82 | 6 | 2 | 6 | 1 | 532 | 58.5 | [illegible] |
| Nuclear factor 1 B-type OS=Mus musculus OX=10090 GN=Nfib PE=1 SV=2 | P97863 | 0 | 8.567 | 4 | 2 | 14 | 1 | 570 | 63.5 | [illegible] |
| Nuclear factor 1 C-type OS=Mus musculus OX=10090 GN=Nfic PE=1 SV=1 | P70255 | 0 | 8.084 | 4 | 2 | 4 | 2 | 439 | 48.7 | [illegible] |
| Nuclear factor NF-kappa-B p100 subunit OS=Mus musculus OX=10090 GN=Nfkb2 PE=1 SV=1 | Q9WTK5 | 0 | 48.482 | 18 | 8 | 37 | 8 | 899 | 96.8 | [illegible] |
| Nuclear factor NF-kappa-B p105 subunit OS=Mus musculus OX=10090 GN=Nfkb1 PE=1 SV=2 | P25799 | 0 | 37.526 | 10 | 5 | 22 | 5 | 971 | 105.5 | [illegible] |
| Nuclear inhibitor of protein phosphatase 1 OS=Mus musculus OX=10090 GN=Ppp1r8 PE=1 SV=1 | Q8R3G1 | 0 | 29.921 | 15 | 3 | 59 | 3 | 351 | 38.5 | [illegible] |
| Nuclear migration protein nudC OS=Mus musculus OX=10090 GN=Nudc PE=1 SV=1 | O35685 | 0 | 44.49 | 25 | 8 | 44 | 8 | 332 | 38.3 | [illegible] |
| [illegible] | [illegible] | [illegible] | [illegible] | [illegible] | [illegible] | [illegible] | [illegible] | [illegible] | [illegible] | [illegible] |

| | | | | | | | | | | | | | | | | | | | | | | | | | | | | |
|---|---|---|---|---|---|---|---|---|---|---|---|---|---|---|---|---|---|---|---|---|---|---|---|---|---|---|---|---|
| Protein farnesyltransferase/geranylgeranyltransferase type-1 subunit alpha OS=Mus musculus OX=10090 GN=Fnta PE=1 SV=1 | Q61239 | 0 | 109.243 | 29 | 7 | 73 | 7 | 377 | 44 | 4.93 | 166.67 | 0.962 | -0.055891201 | 0.995586491 | 76.47 | 111.4 | 88.6 | 97.67 | 106.18 | High | High | High | High | Peak Found | High | High | High | |
| Protein flightless-1 homolog OS=Mus musculus OX=10090 GN=Flii PE=1 SV=1 | Q9JJ28 | 0 | 67.933 | 9 | 9 | 72 | 9 | 1271 | 144.7 | 6.06 | 166.94 | 1.242 | 0.312665174 | 0.445172144 | 91.77 | 110.4 | 89.6 | 16.93 | 42.03 | High | High | High | High | High | High | High | High | Acetyl [N-Term] |
| Protein HEXIM1 OS=Mus musculus OX=10090 GN=Hexim1 PE=1 SV=1 | Q8R409 | 0 | 13.251 | 7 | 2 | 5 | 2 | 356 | 40.2 | 5.38 | 12.42 | 1.479 | 0.564622052 | 0.639342207 | 73.3 | 49.4 | 150.6 | 105.15 | 32.76 | Peak Found | Peak Found | High | High | Not Found | Peak Found | High | Peak Found | |
| Protein Hook homolog 3 OS=Mus musculus OX=10090 GN=Hook3 PE=1 SV=2 | Q8BUK6 | 0 | 147.126 | 28 | 15 | 94 | 15 | 718 | 83.2 | 5.19 | 211.24 | 0.839 | -0.253257284 | 0.636315913 | 53.69 | 97.1 | 102.9 | 45.11 | 47.23 | High | High | High | High | High | High | High | High | Acetyl [N-Term] |
| Protein IMPACT OS=Mus musculus OX=10090 GN=Impact PE=1 SV=2 | O55091 | 0 | 43.405 | 31 | 4 | 34 | 4 | 318 | 36.3 | 5.05 | 62.1 | 1.365 | 0.448900951 | 0.707152641 | 88.13 | 113.3 | 86.7 | 68.74 | 40.12 | High | High | High | High | High | High | Peak Found | High | |
| Protein ITPRID2 OS=Mus musculus OX=10090 GN=Itprid2 PE=1 SV=3 | Q922B9 | 0 | 34.836 | 6 | 5 | 18 | 5 | 1252 | 136.9 | 5.26 | 47.75 | 0.97 | -0.043943348 | 0.995586491 | 97.34 | 112.7 | 87.3 | 38.65 | 42.64 | High | High | High | High | Peak Found | High | High | Peak Found | |
| Protein IWS1 homolog OS=Mus musculus OX=10090 GN=Iws1 PE=1 SV=1 | Q8C1D8 | 0 | 25.507 | 3 | 2 | 36 | 2 | 766 | 85.2 | 4.67 | 78.22 | 0.785 | -0.349235441 | 0.775498881 | 20.27 | 87.3 | 112.7 | 55.66 | 4.08 | Peak Found | High | High | High | Not Found | High | High | High | |
| Protein KHNYN OS=Mus musculus OX=10090 GN=Khnyn PE=2 SV=2 | Q80U38 | 0 | 25.092 | 9 | 4 | 12 | 4 | 671 | 74.5 | 7.87 | 24.62 | 1.999 | 0.999278472 | 0.318093421 | 58 | 133.3 | 66.7 | 54.23 | 8.4 | Not Found | High | High | High | High | Peak Found | Peak Found | High | |
| Protein kinase C alpha type OS=Mus musculus OX=10090 GN=Prkca PE=1 SV=3 | P20444 | 0 | 30.993 | 7 | 4 | 12 | 2 | 672 | 76.8 | 7.14 | 18.34 | 1.488 | 0.573374526 | 0.764830559 | 56.55 | 112.5 | 87.5 | 126.2 | 108.62 | High | Not Found | Peak Found | Not Found | Not Found | Not Found | High | Not Found | |
| Protein kinase C and casein kinase II substrate protein 3 OS=Mus musculus OX=10090 GN=Pacsin3 PE=1 SV=1 | Q99JB8 | 0 | 46.75 | 12 | 3 | 32 | 3 | 424 | 48.6 | 6.1 | 84.93 | 1.426 | 0.511973982 | 0.463710163 | 58.93 | 113.7 | 86.3 | 36.28 | 60.78 | High | High | High | High | High | High | High | High | |
| Protein kinase C and casein kinase substrate in neurons protein 1 OS=Mus musculus OX=10090 GN=Pacsin1 PE=1 SV=1 | Q61644 | 0 | 8.553 | 5 | 2 | 7 | 2 | 441 | 50.5 | 5.24 | 7.62 | | | | | | | | | High | Not Found | Not Found | Not Found | Not Found | High | Not Found | Not Found | |
| Protein kinase C and casein kinase substrate in neurons protein 2 OS=Mus musculus OX=10090 GN=Pacsin2 PE=1 SV=1 | Q9WVE8 | 0 | 135.457 | 22 | 8 | 144 | 8 | 486 | 55.8 | 5.2 | 328.77 | 1.015 | 0.021479727 | 0.991386436 | 72.25 | 100.1 | 99.9 | 18.72 | 26.67 | High | High | High | High | High | High | High | High | |
| Protein kinase C beta type OS=Mus musculus OX=10090 GN=Prkcb PE=1 SV=4 | P68404 | 0 | 27.5 | 5 | 3 | 10 | 1 | 671 | 76.7 | 7.01 | 18.24 | | | | | | | | | High | Not Found | Not Found | Not Found | Not Found | Not Found | Not Found | Not Found | |
| Protein kinase C delta type OS=Mus musculus OX=10090 GN=Prkcd PE=1 SV=3 | P28867 | 0 | 41.018 | 12 | 6 | 28 | 6 | 674 | 77.5 | 7.39 | 59.16 | 1.169 | 0.22527493 | 0.881153299 | 64.46 | 132.1 | 67.9 | 111.91 | 125.07 | Peak Found | High | High | High | Peak Found | High | High | High | |
| Protein kinase C iota type OS=Mus musculus OX=10090 GN=Prkci PE=1 SV=3 | Q62074 | 0 | 21.563 | 5 | 2 | 17 | 2 | 595 | 68.2 | 5.85 | 45.27 | 0.793 | -0.334607229 | 0.791619886 | 40.01 | 88.5 | 111.5 | 33.28 | 83.9 | High | High | Peak Found | High | High | High | Peak Found | High | |
| Protein lin-7 homolog C OS=Mus musculus OX=10090 GN=Lin7c PE=1 SV=2 | O88952 | 0 | 28.182 | 19 | 3 | 29 | 3 | 197 | 21.8 | 8.43 | 72.47 | 1.051 | 0.071762669 | 0.962674291 | 67.33 | 96.1 | 103.9 | 68.18 | 63.41 | Peak Found | High | High | High | Peak Found | High | High | High | |
| Protein LSM14 homolog A OS=Mus musculus OX=10090 GN=Lsm14a PE=1 SV=1 | Q8K2F8 | 0 | 48.235 | 16 | 4 | 41 | 4 | 462 | 50.5 | 9.52 | 119.02 | 0.997 | -0.00433459 | 0.995586491 | 68.22 | 113.8 | 86.2 | 56.59 | 59.24 | High | High | High | High | High | High | High | High | |
| Protein LYRIC OS=Mus musculus OX=10090 GN=Mtdh PE=1 SV=1 | Q80WJ7 | 0 | 70.508 | 13 | 4 | 87 | 4 | 579 | 63.8 | 9.33 | 218.68 | 1.023 | 0.032806145 | 0.99502699 | 51.68 | 104.2 | 95.8 | 45.38 | 51.5 | High | High | High | High | High | High | High | High | |
| Protein LZIC OS=Mus musculus OX=10090 GN=Lzic PE=1 SV=1 | Q8K3C3 | 0 | 69.267 | 29 | 4 | 76 | 4 | 190 | 21.5 | 4.96 | 212.91 | 0.833 | -0.263611599 | 0.546899677 | 69.35 | 95.3 | 104.7 | 66.35 | 75 | High | High | High | High | High | High | High | High | |
| Protein mago nashi homolog OS=Mus musculus OX=10090 GN=Magoh PE=2 SV=1 | P61327 | 0 | 12.307 | 19 | 2 | 38 | 2 | 146 | 17.2 | 6.11 | 77.5 | 1.105 | 0.14404637 | 0.791619886 | 67.18 | 108.3 | 91.7 | 38.41 | 46.57 | High | High | High | High | High | High | High | High | Acetyl [N-Term] |
| Protein MAK16 homolog OS=Mus musculus OX=10090 GN=Mak16 PE=1 SV=1 | Q8BGS0 | 0 | 15.653 | 10 | 2 | 14 | 2 | 296 | 35.2 | 5.31 | 21.51 | | | | | | | | | High | High | Not Found | High | Not Found | High | Peak Found | High | Acetyl [N-Term] |
| Protein MEMO1 OS=Mus musculus OX=10090 GN=Memo1 PE=1 SV=1 | Q91VH6 | 0 | 16.276 | 10 | 2 | 13 | 2 | 297 | 33.7 | 7.14 | 21.17 | 0.884 | -0.177881725 | 0.987145434 | 109.43 | 109.6 | 90.4 | 125.86 | 118.53 | High | Peak Found | High | High | High | Peak Found | High | High | |
| Protein misato homolog 1 OS=Mus musculus OX=10090 GN=Msto1 PE=1 SV=1 | Q2YDW2 | 0 | 17.117 | 8 | 3 | 9 | 3 | 556 | 61.2 | 6.49 | 18.3 | | | | | | | | | High | Not Found | High | High | High | Not Found | Not Found | High | |
| Protein MON2 homolog OS=Mus musculus OX=10090 GN=Mon2 PE=1 SV=2 | Q80TL7 | 0 | 25.748 | 3 | 4 | 16 | 4 | 1715 | 189 | 6.13 | 31.95 | 1.021 | 0.029982866 | 0.991414075 | 19.67 | 56.2 | 143.8 | 100.05 | 8.92 | Peak Found | High | Peak Found | High | Not Found | High | Peak Found | High | |
| Protein mono-ADP-ribosyltransferase PARP14 OS=Mus musculus OX=10090 GN=Parp14 PE=1 SV=3 | Q2EMV9 | 0 | 32.355 | 3 | 3 | 16 | 3 | 1817 | 203.7 | 6.79 | 45.44 | | | | | | | | | Not Found | High | High | High | Not Found | High | Not Found | High | |
| Protein mono-ADP-ribosyltransferase PARP3 OS=Mus musculus OX=10090 GN=Parp3 PE=2 SV=1 | Q3ULW8 | 0 | 44.276 | 18 | 6 | 43 | 6 | 533 | 59.9 | 7.12 | 90.78 | 0.62 | -0.689659879 | 0.207390151 | 40.38 | 97.8 | 102.2 | 40.63 | 48.03 | High | High | High | High | High | High | High | High | |
| Protein mono-ADP-ribosyltransferase PARP4 OS=Mus musculus OX=10090 GN=Parp4 PE=1 SV=1 | E9PYK3 | 0 | 19.563 | 3 | 4 | 13 | 4 | 1969 | 216 | 6.07 | 19.29 | 0.713 | -0.488026018 | 0.828469429 | 50.93 | 71.7 | 128.3 | 138.45 | 44.92 | High | High | High | Peak Found | High | High | Peak Found | High | |
| Protein NDRG1 OS=Mus musculus OX=10090 GN=Ndrg1 PE=1 SV=1 | Q62433 | 0 | 55.063 | 16 | 4 | 38 | 4 | 394 | 43 | 6.1 | 96.2 | 0.964 | -0.052894948 | 0.991386436 | 101.66 | 115.9 | 84.1 | 34.55 | 14.16 | High | High | High | High | Peak Found | High | High | High | |
| Protein NDRG2 OS=Mus musculus OX=10090 GN=Ndrg2 PE=1 SV=1 | Q9QYG0 | 0 | 23.319 | 10 | 2 | 29 | 2 | 371 | 40.8 | 5.4 | 66.06 | 0.774 | -0.369594529 | 0.782186853 | 43.3 | 87.3 | 112.7 | 50.85 | 63.5 | High | High | High | High | Not Found | High | High | High | |
| Protein NDRG3 OS=Mus musculus OX=10090 GN=Ndrg3 PE=1 SV=1 | Q9QYF9 | 0 | 30.607 | 10 | 3 | 24 | 3 | 375 | 41.5 | 5.25 | 71.9 | 0.859 | -0.219269964 | 0.911276781 | 95.84 | 96.7 | 103.3 | 62.35 | 55.33 | Not Found | Peak Found | High | Peak Found | High | High | High | High | Acetyl [N-Term] |
| Protein Niban OS=Mus musculus OX=10090 GN=Fam129a PE=1 SV=2 | Q3UW53 | 0 | 182.051 | 26 | 14 | 240 | 14 | 926 | 102.6 | 4.78 | 568.56 | 0.913 | -0.131313235 | 0.800693956 | 63.47 | 107.8 | 92.2 | 16.9 | 25.4 | High | High | High | High | High | High | High | High | |
| Protein Njmu-R1 OS=Mus musculus OX=10090 PE=1 SV=2 | Q9CYI0 | 0 | 13.445 | 18 | 3 | 11 | 3 | 393 | 44.4 | 5.05 | 30.12 | 1.132 | 0.178873958 | 0.897667358 | 62.45 | 96.2 | 103.8 | 61.38 | 27.58 | Peak Found | High | Peak Found | High | Not Found | High | High | Peak Found | Acetyl [N-Term] |
| Protein Noxp20 OS=Mus musculus OX=10090 GN=Fam114a1 PE=1 SV=1 | Q9D281 | 0 | 157.226 | 45 | 13 | 165 | 13 | 569 | 61 | 4.56 | 339.4 | 0.938 | -0.092340172 | 0.898432344 | 63.16 | 98 | 102 | 11.56 | 18.13 | High | High | High | High | High | High | High | High | |
| Protein odr-4 homolog OS=Mus musculus OX=10090 GN=Odr4 PE=1 SV=2 | Q4PJX1 | 0 | 29.515 | 8 | 2 | 14 | 2 | 447 | 50 | 6.14 | 26.65 | 0.615 | -0.701341684 | 0.663199077 | 44.57 | 71.3 | 128.7 | 91.5 | 72.73 | Not Found | High | High | High | Not Found | Peak Found | Peak Found | High | |
| Protein O-GlcNAcase OS=Mus musculus OX=10090 GN=Oga PE=1 SV=2 | Q9EQQ9 | 0 | 23.136 | 7 | 3 | 15 | 3 | 916 | 103.1 | 4.92 | 16.36 | 0.794 | -0.332789088 | 0.946558434 | 84.84 | 88.5 | 111.5 | 119.6 | 77.34 | Not Found | High | Peak Found | High | Not Found | High | Peak Found | High | |

[illegible]

| | | | | | | | | | | | | | | | | | | | | | | | | | | | | |
|---|---|---|---|---|---|---|---|---|---|---|---|---|---|---|---|---|---|---|---|---|---|---|---|---|---|---|---|---|
| Serine/threonine-protein phosphatase 5 OS=Mus musculus OX=10090 GN=Ppp5c PE=1 SV=3 | Q60676 | 0 | 47.511 | 14 | 4 | 59 | 4 | 499 | 56.8 | 6.2 | 165.2 | 1.059 | 0.082702589 | 0.89151663 | 81.63 | 105.4 | 94.6 | 43.66 | 59.68 | High | High | High | High | High | High | High | High | |
| Serine/threonine-protein phosphatase 6 regulatory subunit 1 OS=Mus musculus OX=10090 GN=Ppp6r1 PE=1 SV=1 | Q7TSI3 | 0 | 45.365 | 13 | 5 | 29 | 5 | 856 | 94.5 | 4.64 | 73.72 | 1.835 | 0.875780063 | 0.019369027 | 80.7 | 112.4 | 87.6 | 89.64 | 15.15 | High | High | High | High | Not Found | High | Peak Found | High | |
| Serine/threonine-protein phosphatase 6 regulatory subunit 3 OS=Mus musculus OX=10090 GN=Ppp6r3 PE=1 SV=1 | Q922D4 | 0 | 32.07 | 6 | 4 | 42 | 4 | 844 | 94.6 | 4.58 | 86.51 | 0.772 | -0.373327247 | 0.346229416 | 46.75 | 94.7 | 105.3 | 23.19 | 14.66 | High | High | High | High | High | High | High | High | |
| Serine/threonine-protein phosphatase CPPED1 OS=Mus musculus OX=10090 GN=Cpped1 PE=1 SV=1 | Q8BFS6 | 0 | 26.068 | 12 | 2 | 21 | 2 | 312 | 35.2 | 5.34 | 60.53 | | | | | | | | | Not Found | High | High | High | Not Found | High | High | High | |
| Serine/threonine-protein phosphatase PP1-alpha catalytic subunit OS=Mus musculus OX=10090 GN=Ppp1ca PE=1 SV=1 | P62137 | 0 | 105.014 | 28 | 8 | 97 | 2 | 330 | 37.5 | 6.33 | 227.79 | | | | | | | | | Peak Found | High | High | High | Not Found | High | Not Found | High | |
| Serine/threonine-protein phosphatase PP1-beta catalytic subunit OS=Mus musculus OX=10090 GN=Ppp1cb PE=1 SV=3 | P62141 | 0 | 102.144 | 33 | 9 | 93 | 3 | 327 | 37.2 | 6.19 | 228.28 | 0.936 | -0.095419565 | 0.947103884 | 74.24 | 102.3 | 97.7 | 66.99 | 81.48 | Not Found | High | Not Found | High | Not Found | High | Not Found | High | |
| Serine-threonine kinase receptor-associated protein OS=Mus musculus OX=10090 GN=Strap PE=1 SV=2 | Q9Z1Z2 | 0 | 132.885 | 41 | 11 | 257 | 11 | 350 | 38.4 | 5.12 | 565.22 | 1.028 | 0.039840265 | 0.95979474 | 74.46 | 105.1 | 94.9 | 24.15 | 20.93 | High | High | High | High | High | High | High | High | |
| Serine--tRNA ligase, cytoplasmic OS=Mus musculus OX=10090 GN=Sars PE=1 SV=3 | P26638 | 0 | 248.765 | 30 | 14 | 352 | 14 | 512 | 58.4 | 6.3 | 886.12 | 1.115 | 0.15704371 | 0.639342207 | 77.53 | 102.4 | 97.6 | 42.07 | 25.15 | High | High | High | High | High | High | High | High | Acetyl [N-Term] |
| Serpin B6 OS=Mus musculus OX=10090 GN=Serpinb6 PE=1 SV=1 | Q60854 | 0 | 341.122 | 42 | 12 | 677 | 12 | 378 | 42.6 | 5.74 | 1809.85 | 0.913 | -0.131313235 | 0.801444012 | 67.2 | 97.5 | 102.5 | 24.79 | 27.32 | High | High | High | High | High | High | High | High | Acetyl [N-Term] |
| Serpin B8 OS=Mus musculus OX=10090 GN=Serpinb8 PE=1 SV=2 | O08800 | 0 | 29.856 | 12 | 3 | 30 | 3 | 374 | 42.1 | 6.46 | 75.96 | 1.288 | 0.365132593 | 0.805818532 | 65.7 | 123.7 | 76.3 | 11.71 | 25.15 | Peak Found | High | Peak Found | High | Not Found | High | High | High | Acetyl [N-Term] |
| Serpin H1 OS=Mus musculus OX=10090 GN=Serpinh1 PE=1 SV=3 | P19324 | 0 | 439.942 | 56 | 22 | 671 | 22 | 417 | 46.5 | 8.82 | 1627.84 | 0.971 | -0.042456799 | 0.982111074 | 78.89 | 98.6 | 101.4 | 18.88 | 21.83 | High | High | High | High | High | High | High | High | |
| Serrate RNA effector molecule homolog OS=Mus musculus OX=10090 GN=Srrt PE=1 SV=1 | Q99MR6 | 0 | 39.24 | 8 | 6 | 62 | 6 | 875 | 100.4 | 5.97 | 123.22 | 0.734 | -0.446148032 | 0.602986469 | 44.72 | 124.5 | 75.5 | 42.96 | 80.86 | Peak Found | High | High | High | High | High | High | High | |
| Serum albumin OS=Mus musculus OX=10090 GN=Alb PE=1 SV=3 | P07724 | 0 | 27.26 | 6 | 3 | 190 | 3 | 608 | 68.6 | 6.07 | 558.65 | | | | | | | | | High | Not Found | High | Peak Found | Peak Found | High | Not Found | High | |
| S-formylglutathione hydrolase OS=Mus musculus OX=10090 GN=Esd PE=1 SV=1 | Q9R0P3 | 0 | 105.639 | 58 | 8 | 94 | 8 | 282 | 31.3 | 7.12 | 185.58 | 0.963 | -0.054392297 | 0.969361739 | 48.25 | 100.2 | 99.8 | 6.7 | 2.66 | High | High | High | High | Peak Found | High | High | High | |
| SH2 domain-containing protein 7 OS=Mus musculus OX=10090 GN=Sh2d7 PE=2 SV=2 | Q8BI17 | 0 | 5.95 | 7 | 2 | 7 | 2 | 458 | 50.5 | 7.61 | 9.81 | 1.315 | 0.3950628 | 0.814249469 | 111.39 | 113.6 | 86.4 | 76.61 | 90.09 | High | High | Peak Found | High | Not Found | Peak Found | Peak Found | Peak Found | |
| SH3 and PX domain-containing protein 2A OS=Mus musculus OX=10090 GN=Sh3pxd2a PE=1 SV=2 | O89032 | 0 | 33.354 | 5 | 3 | 44 | 3 | 1124 | 124.1 | 8.75 | 91.57 | 1.042 | 0.059355278 | 0.969361739 | 56.84 | 98 | 102 | 63.91 | 63.33 | High | High | High | High | Peak Found | High | High | High | |
| SH3 and PX domain-containing protein 2B OS=Mus musculus OX=10090 GN=Sh3pxd2b PE=1 SV=1 | A2AAY5 | 0 | 19.798 | 5 | 2 | 8 | 2 | 908 | 101.5 | 8.66 | 27.33 | 3.462 | 1.791605725 | 0.025810784 | 101.75 | 155.4 | 44.6 | 56.31 | 139.13 | Peak Found | High | High | Not Found | Not Found | High | Peak Found | Peak Found | |
| SH3 domain-binding glutamic acid-rich-like protein 3 OS=Mus musculus OX=10090 GN=Sh3bgrl3 PE=1 SV=1 | Q91VW3 | 0 | 65.945 | 56 | 3 | 57 | 3 | 93 | 10.5 | 5.14 | 145.47 | 1.143 | 0.192825404 | 0.473402226 | 101.73 | 115 | 85 | 71.46 | 89.51 | High | High | High | High | High | High | High | High | |
| SH3 domain-binding glutamic acid-rich-like protein OS=Mus musculus OX=10090 GN=Sh3bgrl PE=1 SV=1 | Q9JJU8 | 0 | 73.083 | 84 | 8 | 100 | 8 | 114 | 12.8 | 4.92 | 217.55 | 0.744 | -0.426625474 | 0.01607704 | 64.24 | 102.9 | 97.1 | 36.66 | 46.19 | High | High | High | High | High | High | High | High | |
| SH3 domain-binding protein 1 OS=Mus musculus OX=10090 GN=Sh3bp1 PE=1 SV=3 | P55194 | 0 | 98.75 | 13 | 6 | 113 | 6 | 680 | 74.1 | 6.6 | 333.7 | 1.181 | 0.240008965 | 0.634308281 | 60.8 | 113.3 | 86.7 | 36.04 | 51.8 | High | High | High | High | High | High | High | High | |
| SH3 domain-containing kinase-binding protein 1 OS=Mus musculus OX=10090 GN=Sh3kbp1 PE=1 SV=1 | Q8R550 | 0 | 115.304 | 14 | 7 | 111 | 7 | 709 | 78.1 | 7.55 | 289.57 | 1.066 | 0.092207438 | 0.895647774 | 55.51 | 109.6 | 90.4 | 22.19 | 33.96 | High | High | High | High | High | High | High | High | |
| SH3 domain-containing protein 19 OS=Mus musculus OX=10090 GN=Sh3d19 PE=1 SV=2 | Q91X43 | 0 | 30.243 | 5 | 3 | 19 | 3 | 789 | 86 | 8.53 | 35.46 | 0.906 | -0.142417045 | 0.959299542 | 33.49 | 104.6 | 95.4 | 56.12 | 47.43 | Peak Found | High | High | High | Not Found | High | High | High | |
| SHC-transforming protein 1 OS=Mus musculus OX=10090 GN=Shc1 PE=1 SV=3 | P98083 | 0 | 41.62 | 11 | 4 | 52 | 4 | 579 | 62.6 | 6.54 | 95.37 | 0.954 | -0.067938829 | 0.968806194 | 79.72 | 94.2 | 105.8 | 42.97 | 45.44 | High | High | High | High | High | High | High | High | |
| Short/branched chain specific acyl-CoA dehydrogenase, mitochondrial OS=Mus musculus OX=10090 GN=Acadsb PE=1 SV=1 | Q9DBL1 | 0 | 65.657 | 33 | 7 | 73 | 7 | 432 | 47.8 | 7.87 | 185.99 | 0.926 | -0.110915901 | 0.928358602 | 96.24 | 102.7 | 97.3 | 41.78 | 74.72 | High | High | High | High | High | High | High | High | |
| Short-chain specific acyl-CoA dehydrogenase, mitochondrial OS=Mus musculus OX=10090 GN=Acads PE=1 SV=2 | Q07417 | 0 | 30.815 | 19 | 6 | 20 | 6 | 412 | 44.9 | 8.47 | 40.71 | 0.486 | -1.040971781 | 0.200268904 | 28.41 | 65.4 | 134.6 | 159.76 | 133.61 | High | High | High | High | High | High | High | High | |
| Sialate O-acetylesterase OS=Mus musculus OX=10090 GN=Siae PE=1 SV=3 | P70665 | 0 | 14.018 | 5 | 2 | 11 | 2 | 541 | 60.7 | 6.79 | 19.17 | 1.072 | 0.100304906 | 0.976368403 | 95.86 | 103.5 | 96.5 | 74.91 | 107.35 | Peak Found | High | Peak Found | High | Peak Found | High | Peak Found | Peak Found | |
| Sialic acid synthase OS=Mus musculus OX=10090 GN=Nans PE=1 SV=1 | Q99J77 | 0 | 70.909 | 20 | 6 | 66 | 6 | 359 | 40 | 7.06 | 155.69 | 1.017 | 0.024319679 | 0.976368403 | 58.31 | 109.3 | 90.7 | 44.9 | 53.37 | High | High | High | High | High | High | High | High | |
| Sialoadhesin OS=Mus musculus OX=10090 GN=Siglec1 PE=1 SV=2 | Q62230 | 0 | 120.572 | 9 | 9 | 68 | 9 | 1695 | 182.9 | 6.67 | 197.85 | 0.868 | -0.204233052 | 0.911396012 | 63.34 | 127 | 73 | 120.65 | 166.51 | Peak Found | High | High | High | Peak Found | High | High | Peak Found | |
| Sickle tail protein OS=Mus musculus OX=10090 GN=Skt PE=1 SV=1 | A2AQ25 | 0 | 25.682 | 3 | 4 | 21 | 4 | 1946 | 212.9 | 7.77 | 26.48 | 0.585 | -0.77349147 | 0.125158808 | 18.3 | 87.5 | 112.5 | 37.44 | 59.3 | Peak Found | High | High | High | Not Found | High | High | High | |
| Sideroflexin-1 OS=Mus musculus OX=10090 GN=Sfxn1 PE=1 SV=3 | Q99JR1 | 0 | 82.801 | 28 | 5 | 102 | 5 | 322 | 35.6 | 9.23 | 242.28 | 1.035 | 0.049630768 | 0.954982698 | 75.85 | 102 | 98 | 54.55 | 47.97 | High | High | High | High | High | High | High | High | |
| Sideroflexin-2 OS=Mus musculus OX=10090 GN=Sfxn2 PE=1 SV=1 | Q925N2 | 0 | 11.71 | 8 | 2 | 12 | 2 | 322 | 36.1 | 8.82 | 26.49 | 0.36 | -1.473931188 | 0.004706589 | 11.56 | 68.5 | 131.5 | 57.31 | 77.56 | High | High | High | High | High | High | Peak Found | High | Acetyl [N-Term] |
| Sideroflexin-3 OS=Mus musculus OX=10090 GN=Sfxn3 PE=1 SV=1 | Q91V61 | 0 | 86.481 | 29 | 6 | 105 | 6 | 321 | 35.4 | 9.51 | 240.56 | 1.012 | 0.01720929 | 0.995586491 | 106.49 | 109 | 91 | 87.36 | 92.24 | High | High | High | High | High | High | High | High | |
| Sigma non-opioid intracellular receptor 1 OS=Mus musculus OX=10090 GN=Sigmar1 PE=1 SV=1 | O55242 | 0 | 7.213 | 11 | 2 | 5 | 2 | 223 | 25.2 | 5.91 | 12.38 | | | | | | | | | Not Found | Not Found | High | Not Found | Not Found | Not Found | Not Found | Not Found | |
| Signal peptidase complex catalytic subunit SEC11A OS=Mus musculus OX=10090 GN=Sec11a PE=1 SV=1 | Q9R0P6 | 0 | 33.316 | 15 | 3 | 40 | 3 | 179 | 20.6 | 9.33 | 88.99 | 0.947 | -0.078563669 | 0.962674291 | 58.06 | 107.6 | 92.4 | 29.65 | 22.76 | High | High | High | High | Not Found | High | High | High | Acetyl [N-Term] |
| Signal peptidase complex subunit 2 OS=Mus musculus OX=10090 GN=Spcs2 PE=1 SV=1 | Q9CYN2 | 0 | 37.92 | 31 | 3 | 37 | 3 | 226 | 25 | 8.57 | 103.82 | 1.04 | 0.056583528 | 0.968507988 | 50.42 | 124.5 | 75.5 | 35.79 | 63.82 | Not Found | High | High | High | Peak Found | High | Peak Found | High | |
| Signal peptide peptidase-like 2A OS=Mus musculus OX=10090 GN=Sppl2a PE=1 SV=2 | Q9JJF9 | 0 | 7.773 | 9 | 2 | 6 | 2 | 523 | 58.1 | 7.06 | 7.29 | 0.979 | -0.030619235 | 0.995586491 | 89.31 | 118.4 | 81.6 | 105.82 | 127.32 | Not Found | High | Peak Found | Peak Found | Not Found | Peak Found | Peak Found | Peak Found | |
| Signal recognition particle 14 kDa protein OS=Mus musculus OX=10090 GN=Srp14 PE=1 SV=1 | P16254 | 0 | 12.724 | 22 | 3 | 13 | 3 | 110 | 12.5 | 10.17 | 24.84 | 0.869 | -0.202571918 | 0.95979474 | 77.61 | 105.6 | 94.4 | 103.84 | 96.18 | High | High | High | High | Not Found | High | Peak Found | High | |
| Signal recognition particle 19 kDa protein OS=Mus musculus OX=10090 GN=Srp19 PE=1 SV=1 | Q9D7A6 | 0 | 47.408 | 26 | 2 | 43 | 2 | 144 | 16.2 | 9.96 | 131.25 | 1.142 | 0.191562651 | 0.837113595 | 66.5 | 104.8 | 95.2 | 65.92 | 37.93 | Peak Found | High | High | High | Not Found | High | High | High | |
| Signal recognition particle 54 kDa protein OS=Mus musculus OX=10090 GN=Srp54 PE=1 SV=2 | P14576 | 0 | 113.012 | 25 | 10 | 148 | 10 | 504 | 55.7 | 8.75 | 401.58 | 1.026 | 0.037030731 | 0.962674291 | 98.28 | 99.9 | 100.1 | 67.92 | 54.86 | High | High | High | High | High | High | High | High | |
| Signal recognition particle receptor subunit alpha OS=Mus musculus OX=10090 GN=Srpra PE=1 SV=1 | Q9DBG7 | 0 | 94.936 | 20 | 8 | 100 | 8 | 636 | 69.6 | 8.95 | 276.07 | 1.054 | 0.075874867 | 0.932720092 | 84.28 | 117.2 | 82.8 | 45.77 | 21.7 | High | High | High | High | High | High | High | High | |
| Signal recognition particle receptor subunit beta OS=Mus musculus OX=10090 GN=Srprb PE=1 SV=1 | P47758 | 0 | 16.059 | 16 | 3 | 7 | 3 | 269 | 29.6 | 9.28 | 11.18 | 0.517 | -0.951763814 | 0.319447605 | 34.35 | 98.5 | 101.5 | 128.62 | 113.57 | Peak Found | High | Peak Found | High | Not Found | High | Peak Found | Peak Found | |
| Signal recognition particle subunit SRP68 OS=Mus musculus OX=10090 GN=Srp68 PE=1 SV=2 | Q8BMA6 | 0 | 165.249 | 32 | 12 | 166 | 12 | 625 | 70.5 | 8.57 | 492.49 | 0.976 | -0.035046947 | 0.992199867 | 84.86 | 104 | 96 | 36.13 | 40.73 | High | High | High | High | High | High | High | High | |
| Signal transducer and activator of transcription 1 OS=Mus musculus OX=10090 GN=Stat1 PE=1 SV=1 | P42225 | 0 | 138.546 | 26 | 15 | 185 | 15 | 749 | 87.1 | 5.58 | 428.7 | 0.98 | -0.029146346 | 0.995586491 | 95.9 | 93.1 | 106.9 | 104.73 | 76.75 | High | High | High | High | High | High | High | High | |
| Signal transducer and activator of transcription 2 OS=Mus musculus OX=10090 GN=Stat2 PE=1 SV=1 | Q9WVL2 | 0 | 27.507 | 8 | 5 | 23 | 5 | 923 | 105.4 | 5.27 | 48.57 | 0.827 | -0.274040765 | 0.805818532 | 34.98 | 97.3 | 102.7 | 54.97 | 67.01 | High | High | High | High | Not Found | High | High | High | |
| Signal transducer and activator of transcription 3 OS=Mus musculus OX=10090 GN=Stat3 PE=1 SV=2 | P42227 | 0 | 197.82 | 23 | 12 | 176 | 12 | 770 | 88 | 6.3 | 452.49 | 0.907 | -0.140825544 | 0.777814598 | 69.84 | 97.9 | 102.1 | 17.39 | 54.67 | High | High | High | High | High | High | High | High | |
| Signal transducer and activator of transcription 5A OS=Mus musculus OX=10090 GN=Stat5a PE=1 SV=1 | P42230 | 0 | 20.68 | 5 | 3 | 7 | 1 | 793 | 90.8 | 6.39 | 21.18 | 0.951 | -0.072482754 | 0.995586491 | 0 | 97.5 | 102.5 | | | Not Found | High | Not Found | High | Not Found | High | Not Found | Not Found | |
| Signal transducer and activator of transcription 5B OS=Mus musculus OX=10090 GN=Stat5b PE=1 SV=1 | P42232 | 0 | 28.4 | 7 | 4 | 24 | 2 | 786 | 89.9 | 6.05 | 73.78 | 0.466 | -1.10159814 | 0.00775693 | 29.1 | 110.2 | 89.8 | 26.03 | 51.49 | Not Found | High | Peak Found | High | Not Found | High | High | High | |
| Signal transducing adapter molecule 1 OS=Mus musculus OX=10090 GN=Stam PE=1 SV=3 | P70297 | 0 | 55.944 | 10 | 4 | 53 | 4 | 548 | 59.7 | 4.84 | 129.64 | 0.651 | -0.619270551 | 0.235208083 | 51.21 | 102.6 | 97.4 | 83.21 | 77.26 | High | High | High | High | Peak Found | High | High | High | |
| [illegible] | | | | | | | | | | | | | | | | | | | | | | | | | | | | |

| | | | | | | | | | | | | | | |
|---|---|---|---|---|---|---|---|---|---|---|---|---|---|---|
| Tubulin gamma-1 chain OS=Mus musculus OX=10090 GN=Tubg1 PE=1 SV=1 | P83887 | 0 | 30.039 | 16 | 4 | 20 | 4 | 451 | 51.1 | 6.02 | 39.34 | 1.077 | 0.10701825 | 0.962674291 |
| Tubulin polymerization-promoting protein family member 3 OS=Mus musculus OX=10090 GN=Tppp3 PE=1 SV=1 | Q9CRB6 | 0 | 65.704 | 40 | 6 | 77 | 6 | 176 | 19 | 9.11 | 183.93 | 1.243 | 0.313826296 | 0.43743225 |
| Tubulin polymerization-promoting protein OS=Mus musculus OX=10090 GN=Tppp PE=1 SV=1 | Q7TQD2 | 0 | 9.788 | 13 | 2 | 9 | 2 | 218 | 23.6 | 9.42 | 11.96 | | | |
| Tubulin-folding cofactor B OS=Mus musculus OX=10090 GN=Tbcb PE=1 SV=2 | Q9D1E6 | 0 | 28.754 | 27 | 3 | 24 | 3 | 244 | 27.4 | 5.24 | 64.04 | 0.992 | -0.011587974 | 0.995586491 |
| Tubulin-specific chaperone A OS=Mus musculus OX=10090 GN=Tbca PE=1 SV=3 | P48428 | 0 | 66.738 | 36 | 5 | 34 | 5 | 108 | 12.8 | 5.27 | 100.19 | 1.045 | 0.063502942 | 0.917028689 |
| Tubulin-specific chaperone C OS=Mus musculus OX=10090 GN=Tbcc PE=1 SV=1 | Q8VCN9 | 0 | 52.265 | 23 | 5 | 63 | 5 | 341 | 38.1 | 5.3 | 152.06 | 1.055 | 0.077242999 | 0.95979474 |
| Tubulin-specific chaperone cofactor E-like protein OS=Mus musculus OX=10090 GN=Tbcel PE=1 SV=1 | Q8C5W3 | 0 | 8.976 | 7 | 2 | 9 | 2 | 424 | 48 | 5.53 | 22.48 | 1.015 | 0.021479727 | 0.995586491 |
| Tubulin-specific chaperone D OS=Mus musculus OX=10090 GN=Tbcd PE=1 SV=1 | Q8BYA0 | 0 | 77.34 | 9 | 7 | 60 | 7 | 1196 | 133.2 | 6.51 | 155.59 | 0.623 | -0.682695932 | 0.125158808 |
| Tubulin--tyrosine ligase-like protein 12 OS=Mus musculus OX=10090 GN=Ttll12 PE=1 SV=1 | Q3UDE2 | 0 | 52.913 | 21 | 9 | 28 | 9 | 639 | 74 | 5.63 | 57.95 | 0.895 | -0.160040413 | 0.976368403 |
| Tuftelin-interacting protein 11 OS=Mus musculus OX=10090 GN=Tfip11 PE=1 SV=1 | Q9ERA6 | 0 | 5.261 | 3 | 2 | 2 | 2 | 838 | 96.2 | 5.9 | 2.49 | | | |
| Tumor necrosis factor alpha-induced protein 8 OS=Mus musculus OX=10090 GN=Tnfaip8 PE=1 SV=1 | Q921Z5 | 0 | 30.554 | 27 | 3 | 31 | 3 | 198 | 22.9 | 7.93 | 59.25 | 0.731 | -0.452056689 | 0.505452934 |
| Tumor necrosis factor receptor superfamily member 10B OS=Mus musculus OX=10090 GN=Tnfrsf10b PE=2 SV=3 | Q9QZM4 | 0.002 | 4.37 | 7 | 2 | 3 | 2 | 381 | 42.1 | 7.11 | 1.63 | 0.456 | -1.13289427 | 0.323636666 |
| Tumor necrosis factor receptor superfamily member 16 OS=Mus musculus OX=10090 GN=Ngfr PE=1 SV=1 | Q9Z0W1 | 0 | 76.299 | 38 | 8 | 80 | 8 | 417 | 44.7 | 4.64 | 163.27 | 0.946 | -0.080087911 | 0.990475871 |
| Tumor necrosis factor receptor type 1-associated DEATH domain protein OS=Mus musculus OX=10090 GN=Tradd PE=1 SV=1 | Q3U0V2 | 0 | 102.282 | 34 | 7 | 93 | 7 | 310 | 34.6 | 5.2 | 272.24 | 0.944 | -0.083141235 | 0.932385213 |
| Tumor protein D52 OS=Mus musculus OX=10090 GN=Tpd52 PE=1 SV=2 | Q62393 | 0 | 104.002 | 33 | 6 | 122 | 6 | 224 | 24.3 | 4.72 | 291.66 | 1.079 | 0.109694865 | 0.78846753 |
| Tumor protein D54 OS=Mus musculus OX=10090 GN=Tpd52l2 PE=1 SV=1 | Q9CYZ2 | 0 | 72.594 | 36 | 8 | 62 | 8 | 220 | 24 | 6.15 | 146.02 | 0.967 | -0.048412205 | 0.976368403 |
| Tumor susceptibility gene 101 protein OS=Mus musculus OX=10090 GN=Tsg101 PE=1 SV=2 | Q61187 | 0 | 31.76 | 17 | 4 | 34 | 4 | 391 | 44.1 | 6.71 | 54.67 | 1 | 0 | 0.995717381 |
| Twinfilin-1 OS=Mus musculus OX=10090 GN=Twf1 PE=1 SV=2 | Q91YR1 | 0 | 97.719 | 15 | 4 | 96 | 4 | 350 | 40.1 | 6.67 | 251.38 | 1.047 | 0.066261442 | 0.932720092 |
| Twinfilin-2 OS=Mus musculus OX=10090 GN=Twf2 PE=1 SV=1 | Q9Z0P5 | 0 | 43.087 | 25 | 5 | 28 | 5 | 349 | 39.4 | 6.8 | 68.07 | 0.806 | -0.311148256 | 0.660475093 |
| Tyrosine-protein kinase BAZ1B OS=Mus musculus OX=10090 GN=Baz1b PE=1 SV=2 | Q9Z277 | 0 | 59.53 | 6 | 6 | 43 | 6 | 1479 | 170.5 | 8.6 | 108.83 | 0.8 | -0.321928095 | 0.680546304 |
| Tyrosine-protein kinase CSK OS=Mus musculus OX=10090 GN=Csk PE=1 SV=2 | P41241 | 0 | 27.431 | 13 | 4 | 33 | 4 | 450 | 50.7 | 7.06 | 71.62 | 0.804 | -0.314732593 | 0.841253962 |
| Tyrosine-protein kinase Fyn OS=Mus musculus OX=10090 GN=Fyn PE=1 SV=4 | P39688 | 0 | 10.501 | 7 | 3 | 16 | 2 | 537 | 60.6 | 6.67 | 26.63 | 0.242 | -2.046921047 | 0.002256857 |
| Tyrosine-protein kinase HCK OS=Mus musculus OX=10090 GN=Hck PE=1 SV=4 | P08103 | 0 | 26.395 | 7 | 3 | 38 | 2 | 524 | 59.1 | 7.24 | 80.56 | 1.222 | 0.289244285 | 0.768272619 |
| Tyrosine-protein kinase Lyn OS=Mus musculus OX=10090 GN=Lyn PE=1 SV=4 | P25911 | 0 | 41.288 | 13 | 5 | 45 | 4 | 512 | 58.8 | 7.15 | 94.23 | 1.007 | 0.010063683 | 0.995586491 |
| Tyrosine-protein kinase SYK OS=Mus musculus OX=10090 GN=Syk PE=1 SV=2 | P48025 | 0 | 25.37 | 7 | 2 | 18 | 2 | 629 | 71.3 | 7.88 | 49.94 | 1.636 | 0.710172748 | 0.222448638 |
| Tyrosine-protein kinase transmembrane receptor ROR2 OS=Mus musculus OX=10090 GN=Ror2 PE=1 SV=2 | Q9Z138 | 0 | 46.458 | 7 | 5 | 33 | 5 | 944 | 104.9 | 6.47 | 77.15 | 0.986 | -0.020340448 | 0.999900066 |
| Tyrosine-protein kinase Yes OS=Mus musculus OX=10090 GN=Yes1 PE=1 SV=3 | Q04736 | 0 | 8.589 | 6 | 2 | 16 | 1 | 541 | 60.6 | 6.64 | 25.56 | | | |
| Tyrosine-protein phosphatase non-receptor type 1 OS=Mus musculus OX=10090 GN=Ptpn1 PE=1 SV=2 | P35821 | 0 | 51.387 | 20 | 7 | 46 | 7 | 432 | 49.6 | 6.16 | 103.47 | 0.834 | -0.261880711 | 0.727041832 |
| Tyrosine-protein phosphatase non-receptor type 11 OS=Mus musculus OX=10090 GN=Ptpn11 PE=1 SV=2 | P35235 | 0 | 54.104 | 21 | 7 | 51 | 7 | 597 | 68.4 | 7.3 | 127.57 | 0.896 | -0.158429363 | 0.939815692 |
| Tyrosine-protein phosphatase non-receptor type 12 OS=Mus musculus OX=10090 GN=Ptpn12 PE=1 SV=3 | P35831 | 0 | 15.556 | 5 | 3 | 13 | 3 | 775 | 86.5 | 5.99 | 17.26 | 0.446 | -1.164884385 | 0.189088814 |
| Tyrosine-protein phosphatase non-receptor type 2 OS=Mus musculus OX=10090 GN=Ptpn2 PE=1 SV=2 | Q06180 | 0 | 33.146 | 8 | 3 | 24 | 3 | 406 | 47.3 | 8.37 | 64.09 | 0.794 | -0.332789088 | 0.849159812 |
| Tyrosine-protein phosphatase non-receptor type 23 OS=Mus musculus OX=10090 GN=Ptpn23 PE=1 SV=2 | Q6PB44 | 0 | 71.999 | 10 | 9 | 51 | 9 | 1692 | 185.1 | 6.8 | 145.53 | 1.147 | 0.197865391 | 0.806428765 |
| Tyrosine-protein phosphatase non-receptor type 6 OS=Mus musculus OX=10090 GN=Ptpn6 PE=1 SV=2 | P29351 | 0 | 91.043 | 24 | 10 | 127 | 10 | 595 | 67.5 | 7.81 | 277.15 | 1.394 | 0.479230561 | 0.088531344 |
| Tyrosine-protein phosphatase non-receptor type substrate 1 OS=Mus musculus OX=10090 GN=Sirpa PE=1 SV=2 | P97797 | 0 | 33.964 | 17 | 4 | 22 | 1 | 513 | 56.4 | 8.28 | 30.52 | | | |
| Tyrosine--tRNA ligase, cytoplasmic OS=Mus musculus OX=10090 GN=Yars PE=1 SV=3 | Q91WQ3 | 0 | 76.318 | 28 | 11 | 60 | 11 | 528 | 59.1 | 7.01 | 159.17 | 0.914 | -0.12973393 | 0.882906899 |
| U1 small nuclear ribonucleoprotein 70 kDa OS=Mus musculus OX=10090 GN=Snrnp70 PE=1 SV=2 | Q62376 | 0 | 11.691 | 19 | 3 | 8 | 3 | 448 | 52 | 9.94 | 6.73 | 1.273 | 0.348232419 | 0.855420743 |
| U1 small nuclear ribonucleoprotein C OS=Mus musculus OX=10090 GN=Snrpc PE=1 SV=1 | Q62241 | 0 | 34.159 | 25 | 3 | 34 | 3 | 159 | 17.4 | 9.67 | 80.96 | 1.033 | 0.046840254 | 0.983938563 |
| U2 small nuclear ribonucleoprotein A' OS=Mus musculus OX=10090 GN=Snrpa1 PE=1 SV=2 | P57784 | 0 | 49.508 | 17 | 3 | 38 | 3 | 255 | 28.3 | 8.62 | 126.4 | 0.893 | -0.16326792 | 0.868660687 |
| U2 snRNP-associated SURP motif-containing protein OS=Mus musculus OX=10090 GN=U2surp PE=1 SV=3 | Q6NV83 | 0 | 47.584 | 6 | 5 | 37 | 5 | 1029 | 118.2 | 8.47 | 79.31 | 0.945 | -0.081613766 | 0.990475871 |
| U3 small nucleolar RNA-associated protein 14 homolog A OS=Mus musculus OX=10090 GN=Utp14a PE=1 SV=1 | Q640M1 | 0 | 10.776 | 5 | 3 | 3 | 3 | 767 | 87.2 | 9.2 | 8.98 | | | |
| U4/U6 small nuclear ribonucleoprotein Prp31 OS=Mus musculus OX=10090 GN=Prpf31 PE=1 SV=3 | Q8CCF0 | 0 | 24.507 | 13 | 4 | 21 | 4 | 499 | 55.4 | 5.67 | 39.75 | 1.099 | 0.136191386 | 0.894782103 |
| U4/U6.U5 tri-snRNP-associated protein 1 OS=Mus musculus OX=10090 GN=Sart1 PE=1 SV=1 | Q9Z315 | 0 | 61.492 | 16 | 8 | 46 | 8 | 806 | 90.8 | 5.82 | 124.71 | 0.838 | -0.254977851 | 0.851237473 |
| U4/U6.U5 tri-snRNP-associated protein 2 OS=Mus musculus OX=10090 GN=Usp39 PE=1 SV=2 | Q3TIX9 | 0 | 12.645 | 5 | 2 | 8 | 2 | 564 | 65.1 | 8.9 | 23.01 | 0.957 | -0.06340917 | 0.995586491 |
| U5 small nuclear ribonucleoprotein 200 kDa helicase OS=Mus musculus OX=10090 GN=Snrnp200 PE=1 SV=1 | Q6P4T2 | 0 | 315.417 | 18 | 29 | 301 | 29 | 2136 | 244.4 | 6.06 | 772.89 | 0.985 | -0.02180437 | 0.995586491 |
| U6 snRNA-associated Sm-like protein LSm4 OS=Mus musculus OX=10090 GN=Lsm4 PE=1 SV=1 | Q9QXA5 | 0 | 29.306 | 34 | 5 | 32 | 5 | 137 | 15.1 | 10.05 | 64.22 | 1.112 | 0.153156788 | 0.849159812 |
| U6 snRNA-associated Sm-like protein LSm8 OS=Mus musculus OX=10090 GN=Lsm8 PE=1 SV=3 | Q6ZWM4 | 0 | 53.302 | 42 | 2 | 48 | 2 | 96 | 10.4 | 4.48 | 137.15 | 1.132 | 0.178873958 | 0.633056096 |
| U8 snoRNA-decapping enzyme OS=Mus musculus OX=10090 GN=Nudt16 PE=1 SV=1 | Q6P3D0 | 0 | 16.719 | 23 | 4 | 22 | 4 | 195 | 21.8 | 7.12 | 40.5 | 1.204 | 0.267835392 | 0.745941542 |
| UAP56-interacting factor OS=Mus musculus OX=10090 GN=Fyttd1 PE=1 SV=1 | Q91Z49 | 0 | 17.623 | 11 | 2 | 11 | 2 | 317 | 35.9 | 11.84 | 25.33 | 0.876 | -0.190997225 | 0.907852237 |
| Ubiquilin-1 OS=Mus musculus OX=10090 GN=Ubqln1 PE=1 SV=1 | Q8R317 | 0 | 50.333 | 20 | 6 | 32 | 1 | 582 | 61.9 | 4.94 | 82.29 | 0.994 | -0.008682243 | 0.995586491 |
| Ubiquilin-2 OS=Mus musculus OX=10090 GN=Ubqln2 PE=1 SV=2 | Q9QZM0 | 0 | 91.128 | 22 | 9 | 68 | 7 | 638 | 67.3 | 5.22 | 163.65 | 1.095 | 0.13093087 | 0.895647774 |
| Ubiquilin-4 OS=Mus musculus OX=10090 GN=Ubqln4 PE=1 SV=1 | Q99NB8 | 0 | 27.883 | 13 | 5 | 24 | 4 | 596 | 63.5 | 5.03 | 36.13 | 0.897 | -0.15682011 | 0.962674291 |
| Ubiquinol-cytochrome-c reductase complex assembly factor 2 OS=Mus musculus OX=10090 GN=Uqcc2 PE=1 SV=1 | Q9CQY6 | 0 | 37.275 | 38 | 3 | 24 | 3 | 136 | 16.3 | 9.19 | 76.32 | 0.586 | -0.77102743 | 0.458531266 |
| Ubiquinone biosynthesis protein COQ9, mitochondrial OS=Mus musculus OX=10090 GN=Coq9 PE=1 SV=1 | Q8K1Z0 | 0 | 36.899 | 12 | 2 | 25 | 2 | 313 | 35.1 | 5.92 | 69.26 | 0.921 | -0.118726939 | 0.992199867 |
| Ubiquitin carboxyl-terminal hydrolase 10 OS=Mus musculus OX=10090 GN=Usp10 PE=1 SV=3 | P52479 | 0 | 23.776 | 6 | 3 | 32 | 3 | 792 | 87 | 5.17 | 86.65 | 0.711 | -0.492078535 | 0.406969725 |
| Ubiquitin carboxyl-terminal hydrolase 11 OS=Mus musculus OX=10090 GN=Usp11 PE=1 SV=4 | Q99K46 | 0 | 14.162 | 4 | 3 | 7 | 2 | 921 | 105.3 | 4.96 | 12.09 | 1.25 | 0.321928095 | 0.893756428 |
| Ubiquitin carboxyl-terminal hydrolase 14 OS=Mus musculus OX=10090 GN=Usp14 PE=1 SV=3 | Q9JMA1 | 0 | 71.136 | 29 | 9 | 56 | 9 | 493 | 56 | 5.24 | 133.31 | 1.071 | 0.09895848 | 0.962674291 |
| Ubiquitin carboxyl-terminal hydrolase 15 OS=Mus musculus OX=10090 GN=Usp15 PE=1 SV=1 | Q8R5H1 | 0 | 32.622 | 6 | 4 | 21 | 4 | 981 | 112.3 | 5.17 | 50.71 | 1.387 | 0.471967788 | 0.745941542 |
| Ubiquitin carboxyl-terminal hydrolase 16 OS=Mus musculus OX=10090 GN=Usp16 PE=1 SV=2 | Q99LG0 | 0 | 6.226 | 3 | 2 | 2 | 2 | 825 | 93.4 | 6.18 | 1.82 | | | |
| Ubiquitin carboxyl-terminal hydrolase 19 OS=Mus musculus OX=10090 GN=Usp19 PE=1 SV=1 | Q3UJD6 | 0 | 32.495 | 6 | 4 | 20 | 4 | 1360 | 150.5 | 6.38 | 44.22 | 0.905 | -0.144010303 | 0.983930052 |
| Ubiquitin carboxyl-terminal hydrolase 24 OS=Mus musculus OX=10090 GN=Usp24 PE=1 SV=1 | B1AY13 | 0 | 16.602 | 1 | 2 | 9 | 2 | 2617 | 293.8 | 6.19 | 24.35 | 1.369 | 0.453122447 | 0.704043501 |
| Ubiquitin carboxyl-terminal hydrolase 34 OS=Mus musculus OX=10090 GN=Usp34 PE=1 SV=3 | Q6ZQ93 | 0 | 5.932 | 1 | 2 | 9 | 2 | 3582 | 408 | 5.68 | 19.19 | 0.659 | -0.60164963 | 0.805818532 |
| Ubiquitin carboxyl-terminal hydrolase 4 OS=Mus musculus OX=10090 GN=Usp4 PE=1 SV=3 | P35123 | 0 | 41.918 | 14 | 7 | 43 | 6 | 962 | 108.3 | 5.64 | 117.62 | 0.999 | -0.001443417 | 0.999900066 |
| Ubiquitin carboxyl-terminal hydrolase 40 OS=Mus musculus OX=10090 GN=Usp40 PE=1 SV=2 | Q8BWR4 | 0 | 35.172 | 4 | 3 | 31 | 3 | 1235 | 139.9 | 5.63 | 62.92 | | | |
| Ubiquitin carboxyl-terminal hydrolase 47 OS=Mus musculus OX=10090 GN=Usp47 PE=1 SV=2 | Q8BY87 | 0 | 108.619 | 11 | 10 | 105 | 10 | 1376 | 157.4 | 5.11 | 253.67 | 1.007 | 0.010063683 | 0.995717381 |
| Ubiquitin carboxyl-terminal hydrolase 5 OS=Mus musculus OX=10090 GN=Usp5 PE=1 SV=1 | P56399 | 0 | 193.47 | 30 | 15 | 232 | 15 | 858 | 95.8 | 5.01 | 559.11 | 1.078 | 0.108357178 | 0.791619886 |
| Ubiquitin carboxyl-terminal hydrolase 7 OS=Mus musculus OX=10090 GN=Usp7 PE=1 SV=1 | Q6A4J8 | 0 | 22.784 | 6 | 4 | 12 | 4 | 1103 | 128.4 | 5.5 | 30.86 | 1.051 | 0.071762669 | 0.969361739 |
| Ubiquitin carboxyl-terminal hydrolase isozyme L1 OS=Mus musculus OX=10090 GN=Uchl1 PE=1 SV=1 | Q9R0P9 | 0 | 83.339 | 47 | 5 | 60 | 5 | 223 | 24.8 | 5.24 | 137.76 | 0.893 | -0.16326792 | 0.893249611 |
| Ubiquitin carboxyl-terminal hydrolase isozyme L3 OS=Mus musculus OX=10090 GN=Uchl3 PE=1 SV=2 | Q9JKB1 | 0 | 49.674 | 26 | 5 | 47 | 5 | 230 | 26.1 | 5.05 | 133.25 | 1.025 | 0.03562391 | 0.962674291 |
| Ubiquitin carboxyl-terminal hydrolase isozyme L5 OS=Mus musculus OX=10090 GN=Uchl5 PE=1 SV=2 | Q9WUP7 | 0 | 41.7 | 16 | 3 | 40 | 1 | 329 | 37.6 | 5.33 | 119.18 | 1.137 | 0.185232254 | 0.895647774 |
| Ubiquitin carboxyl-terminal hydrolase MINDY-3 OS=Mus musculus OX=10090 GN=MINDY3 PE=1 SV=2 | Q9CV28 | 0 | 17.154 | 9 | 2 | 13 | 2 | 444 | 49.6 | 4.78 | 28.36 | 1.04 | 0.056583528 | 0.976368403 |
| Ubiquitin conjugation factor E4 B OS=Mus musculus OX=10090 GN=Ube4b PE=1 SV=3 | Q9ES00 | 0 | 49.353 | 11 | 7 | 28 | 7 | 1173 | 133.2 | 6.07 | 38.62 | 1.347 | 0.429749851 | 0.417919889 |
| Ubiquitin thioesterase OTUB1 OS=Mus musculus OX=10090 GN=Otub1 PE=1 SV=2 | Q7TQI3 | 0 | 95.925 | 30 | 6 | 93 | 6 | 271 | 31.3 | 4.94 | 219.23 | 0.821 | -0.284545873 | 0.411153887 |
| Ubiquitin-40S ribosomal protein S27a OS=Mus musculus OX=10090 GN=Rps27a PE=1 SV=2 | P62983 | 0 | 131.4 | 47 | 7 | 267 | 7 | 156 | 17.9 | 9.64 | 649.34 | 1.186 | 0.24610401 | 0.255206714 |
| Ubiquitin-associated and SH3 domain-containing protein B OS=Mus musculus OX=10090 GN=Ubash3b PE=1 SV=1 | Q8BGG7 | 0 | 11.361 | 4 | 2 | 4 | 2 | 638 | 71.4 | 6.81 | 6.21 | | | |
| Ubiquitin-associated domain-containing protein 2 OS=Mus musculus OX=10090 GN=Ubac2 PE=1 SV=1 | Q8R1K1 | 0 | 9.677 | 8 | 2 | 7 | 2 | 345 | 39 | 9.44 | 17.87 | | | |
| Ubiquitin-associated protein 1 OS=Mus musculus OX=10090 GN=Ubap1 PE=1 SV=1 | Q8BH48 | 0 | 26.413 | 10 | 3 | 29 | 3 | 502 | 55 | 5.07 | 61.79 | 0.542 | -0.883635243 | 0.43743225 |
| Ubiquitin-associated protein 2 OS=Mus musculus OX=10090 GN=Ubap2 PE=1 SV=1 | Q91VX2 | 0 | 29.102 | 3 | 2 | 26 | 2 | 1132 | 117.9 | 7.72 | 66.57 | 1.086 | 0.119024103 | 0.953707805 |
| Ubiquitin-associated protein 2-like OS=Mus musculus OX=10090 GN=Ubap2l PE=1 SV=1 | Q80X50 | 0 | 118.949 | 17 | 9 | 149 | 9 | 1107 | 116.7 | 7.11 | 404.47 | 0.897 | -0.15682011 | 0.764830559 |
| [illegible] | | | | | | | | | | | | | | |

| Description | Accession | Exp. q-value | Sum PEP Score | Coverage [%] | # Peptides | # PSMs | # Unique Peptides | # AAs | MW [kDa] | calc. pI | Score Sequest HT: Sequest HT | Abundance Ratio: (shGAN) / (Scramble) | Log Fold Change: (shGAN) / (Scramble) | Abundance Ratio Adj. P-Value: (shGAN) / (Scramble) | Abundance Ratio Variability [%]: (shGAN) / (Scramble) | Abundances (Grouped): shGAN | Abundances (Grouped): Scramble | Abundances (Grouped) CV [%]: shGAN | Abundances (Grouped) CV [%]: Scramble | shGAN DRG1124 (Light) | ShGAN DRG1207 (Heavy) | shGAN DRG1204 (Light) | ShGAN DRG1211 (Heavy) | Scramble DRG1124 (Heavy) | Scramble DRG1207 (Light) | Scramble DRG1204 (Heavy) | Scramble DRG1211 (Light) | Modifications |
|---|---|---|---|---|---|---|---|---|---|---|---|---|---|---|---|---|---|---|---|---|---|---|---|---|---|---|---|---|
| 2',3'-cyclic-nucleotide 3'-phosphodiesterase OS=Mus musculus OX=10090 GN=Cnp PE=1 SV=3 | P16330 | 0 | 35.904 | 8 | 3 | 53 | 3 | 420 | 47.1 | 8.97 | 58.15 | 2.909 | 1.540523296 | 0.03788856 | 99.42 | 56.9 | 143.1 | 138.22 | | High | Not Found | High | Peak Found | Peak Found | Not Found | Peak Found | Peak Found | |
| ADP-ribose pyrophosphatase, mitochondrial OS=Mus musculus OX=10090 GN=Nudt9 PE=1 SV=1 | Q8BVU5 | 0 | 14.188 | 14 | 2 | 5 | 2 | 350 | 38.6 | 6.76 | 10.42 | 0.278 | -1.846843212 | 0.034578825 | 23.77 | 43.5 | 156.5 | 127.36 | | Peak Found | Peak Found | High | Peak Found | High | High | High | Peak Found | |
| ADP-ribosylation factor-binding protein GGA2 OS=Mus musculus OX=10090 GN=Gga2 PE=1 SV=1 | Q6P5E6 | 0 | 52.107 | 7 | 2 | 58 | 2 | 603 | 66 | 7.3 | 120.73 | 1.638 | 0.711935357 | 0.01978112 | 128.38 | 105.7 | 94.3 | 100.06 | 188.05 | High | High | High | High | Peak Found | High | High | High | |
| ADP-ribosylation factor-binding protein GGA3 OS=Mus musculus OX=10090 GN=Gga3 PE=1 SV=2 | Q8BMI3 | 0 | 37.932 | 5 | 2 | 18 | 2 | 718 | 77.9 | 5.66 | 56.57 | 0.268 | -1.899695094 | 1.89371E-07 | 14.1 | 42.2 | 157.8 | 112.96 | | High | High | High | Peak Found | Not Found | High | High | High | |
| A-kinase anchor protein 12 OS=Mus musculus OX=10090 GN=Akap12 PE=1 SV=1 | Q9WTQ5 | 0 | 1416.672 | 72 | 76 | 2467 | 76 | 1684 | 180.6 | 4.44 | 5764.98 | 0.738 | -0.438307279 | 0.01231637 | 53.5 | 94.8 | 105.2 | 28.48 | 18.05 | High | High | High | High | High | High | High | High | |
| Alpha-actinin-3 OS=Mus musculus OX=10090 GN=Actn3 PE=2 SV=1 | O88990 | 0 | 102.639 | 10 | 10 | 97 | 3 | 900 | 103 | 5.45 | 246.91 | 1.374 | 0.458382004 | 0.003516143 | 94.22 | 122.9 | 77.1 | 48.82 | 84.21 | High | High | High | High | Peak Found | High | High | High | |
| ATP-binding cassette sub-family B member 6, mitochondrial OS=Mus musculus OX=10090 GN=Abcb6 PE=1 SV=1 | Q9DC29 | 0 | 21.276 | 6 | 3 | 9 | 3 | 842 | 93.7 | 8.05 | 21.01 | 0.346 | -1.531156057 | 0.019270889 | 12.74 | 87.5 | 112.5 | 45.15 | 37.8 | High | High | Peak Found | High | Peak Found | High | Not Found | High | |
| ATP-dependent zinc metalloprotease YME1L1 OS=Mus musculus OX=10090 GN=Yme1l1 PE=1 SV=1 | O88967 | 0 | 63.087 | 8 | 3 | 50 | 3 | 715 | 80 | 8.97 | 162.71 | 0.54 | -0.888968688 | 0.020444886 | 20.32 | 56.1 | 143.9 | 66.69 | 77.72 | High | High | High | High | High | High | High | High | |
| AT-rich interactive domain-containing protein 1A OS=Mus musculus OX=10090 GN=Arid1a PE=1 SV=1 | A2BH40 | 0 | 22.807 | 1 | 2 | 21 | 2 | 2283 | 241.9 | 6.68 | 46.79 | 4.997 | 2.321062218 | 4.30071E-06 | 115.34 | 169.1 | 30.9 | 5.49 | 112.92 | High | High | High | High | High | High | High | High | |
| BRCA1-associated protein OS=Mus musculus OX=10090 GN=Brap PE=1 SV=1 | Q99MP8 | 0 | 18.603 | 7 | 3 | 12 | 3 | 591 | 66.9 | 5.64 | 28.34 | 0.291 | -1.780908942 | 0.000927906 | 2.54 | 45.1 | 154.9 | 3.81 | 28.69 | Not Found | High | High | High | Peak Found | Peak Found | High | High | |
| BTB/POZ domain-containing protein KCTD12 OS=Mus musculus OX=10090 GN=Kctd12 PE=1 SV=1 | Q6WVG3 | 0 | 56.305 | 31 | 7 | 132 | 7 | 327 | 35.9 | 5.81 | 305.64 | 0.575 | -0.798366139 | 1.72773E-08 | 40.23 | 77.8 | 122.2 | 105.78 | 52.56 | High | High | High | High | High | High | High | High | |
| Calcium-activated chloride channel regulator 3A-1 OS=Mus musculus OX=10090 GN=Clca3a1 PE=1 SV=1 | Q9QX15 | 0 | 12.143 | 3 | 2 | 10 | 2 | 902 | 100 | 6.86 | 21.34 | 0.021 | -5.573466862 | 5.68375E-15 | 0.05 | 4.2 | 195.8 | 188.93 | | High | Peak Found | Peak Found | Peak Found | High | High | Not Found | Not Found | |
| Cathepsin B OS=Mus musculus OX=10090 GN=Ctsb PE=1 SV=2 | P10605 | 0 | 235.999 | 40 | 11 | 510 | 11 | 339 | 37.3 | 5.91 | 1149.23 | 1.453 | 0.539034703 | 0.000294893 | 74.81 | 114.6 | 85.4 | 58.77 | 37.84 | High | High | High | High | High | High | High | High | |
| CD2-associated protein OS=Mus musculus OX=10090 GN=Cd2ap PE=1 SV=3 | Q9JLQ0 | 0 | 24.629 | 10 | 5 | 19 | 5 | 637 | 70.4 | 6.38 | 42.52 | 0.367 | -1.446148032 | 0.010671883 | 19.69 | 85.1 | 114.9 | 134.38 | 111.61 | High | High | High | High | High | High | High | Peak Found | |
| Cellular retinoic acid-binding protein 2 OS=Mus musculus OX=10090 GN=Crabp2 PE=2 SV=2 | P22935 | 0 | 82.199 | 47 | 5 | 152 | 5 | 138 | 15.7 | 5.71 | 399.74 | 1.394 | 0.479230561 | 0.005319816 | 80.71 | 114.2 | 85.8 | 69.09 | 68 | Peak Found | High | High | High | High | High | High | High | |
| Cis-aconitate decarboxylase OS=Mus musculus OX=10090 GN=Acod1 PE=1 SV=2 | P54987 | 0 | 18.362 | 5 | 3 | 25 | 3 | 488 | 53.7 | 7.55 | 63.25 | 2.418 | 1.273814245 | 4.06082E-07 | 102.34 | 137.2 | 62.8 | 50.63 | 93.45 | High | High | High | High | Peak Found | High | High | High | |
| Clathrin interactor 1 OS=Mus musculus OX=10090 GN=Clint1 PE=1 SV=2 | Q99KN9 | 0 | 72.493 | 16 | 7 | 92 | 7 | 631 | 68.5 | 6.25 | 237.15 | 0.686 | -0.543719518 | 0.047740226 | 41.85 | 86.7 | 113.3 | 69.47 | 65.25 | High | High | High | High | High | High | High | High | |
| Cleft lip and palate transmembrane protein 1 homolog OS=Mus musculus OX=10090 GN=Clptm1 PE=1 SV=1 | Q8VBZ3 | 0 | 36.476 | 10 | 3 | 27 | 3 | 664 | 75.2 | 6.3 | 64.88 | 2.395 | 1.260025656 | 0.035868102 | 111.13 | 103.3 | 96.7 | 63.01 | 79.27 | Peak Found | High | High | High | Not Found | High | High | High | |
| Clusterin OS=Mus musculus OX=10090 GN=Clu PE=1 SV=1 | Q06890 | 0 | 11.89 | 6 | 2 | 17 | 2 | 448 | 51.6 | 5.67 | 38.68 | 2.912 | 1.542010356 | 0.000687267 | 128.72 | 134.2 | 65.8 | 163.47 | 119.73 | High | High | High | High | High | High | Not Found | High | |
| CMRF35-like molecule 3 OS=Mus musculus OX=10090 GN=Cd300ld3 PE=1 SV=1 | Q6SJQ5 | 0 | 5.401 | 12 | 2 | 3 | 2 | 245 | 27.3 | 4.7 | 4.27 | 0.355 | -1.49410907 | 0.001073733 | 9.95 | 52.4 | 147.6 | 17.38 | 64.37 | Peak Found | High | Peak Found | Not Found | Not Found | Peak Found | Peak Found | Peak Found | |
| Coactosin-like protein OS=Mus musculus OX=10090 GN=Cotl1 PE=1 SV=3 | Q9CQI6 | 0 | 57.487 | 55 | 7 | 32 | 7 | 142 | 15.9 | 5.4 | 45.55 | 1.528 | 0.611644543 | 0.002267345 | 84.75 | 120.7 | 79.3 | 77.83 | 84.27 | Peak Found | High | High | High | Peak Found | High | High | High | |
| Coagulation factor XIII A chain OS=Mus musculus OX=10090 GN=F13a1 PE=1 SV=3 | Q8BH61 | 0 | 114.633 | 18 | 11 | 145 | 11 | 732 | 83.2 | 5.92 | 389.86 | 1.98 | 0.98550043 | 1.53281E-11 | 118.53 | 134.9 | 65.1 | 52.05 | 98.03 | High | High | High | High | High | High | High | High | |
| Coiled-coil domain-containing protein 115 OS=Mus musculus OX=10090 GN=Ccdc115 PE=1 SV=1 | Q8VE99 | 0 | 22.112 | 17 | 2 | 26 | 2 | 180 | 19.7 | 8.81 | 71.01 | 2.014 | 1.010063683 | 0.027225658 | 81.91 | 133.7 | 66.3 | 40.96 | 35.35 | High | High | High | High | High | Peak Found | High | High | |
| Collagen alpha-1(I) chain OS=Mus musculus OX=10090 GN=Col1a1 PE=1 SV=4 | P11087 | 0 | 182.693 | 12 | 13 | 284 | 13 | 1453 | 137.9 | 5.85 | 625.99 | 0.705 | -0.504304837 | 0.001813678 | 52.52 | 89.5 | 110.5 | 61.95 | 68.1 | High | High | High | High | High | High | High | High | |
| Collagen alpha-1(IV) chain OS=Mus musculus OX=10090 GN=Col4a1 PE=1 SV=4 | P02463 | 0 | 21.166 | 2 | 2 | 12 | 2 | 1669 | 160.6 | 8.24 | 25.51 | 2.216 | 1.147957881 | 0.000237716 | 78.38 | 140.9 | 59.1 | 82.55 | 78.05 | Not Found | High | High | High | Not Found | High | Peak Found | High | |
| Collagen alpha-2(I) chain OS=Mus musculus OX=10090 GN=Col1a2 PE=1 SV=2 | Q01149 | 0 | 297.917 | 14 | 14 | 521 | 14 | 1372 | 129.5 | 9.19 | 1263.91 | 0.767 | -0.382701517 | 0.045147397 | 35.41 | 87.6 | 112.4 | 9.88 | 9.81 | High | High | High | High | High | High | High | High | |
| COMM domain-containing protein 10 OS=Mus musculus OX=10090 GN=Commd10 PE=1 SV=1 | Q8JZY2 | 0 | 15.679 | 16 | 2 | 8 | 2 | 202 | 22.8 | 6.65 | 14.68 | 0.412 | -1.279283757 | 0.000413304 | 23.94 | 58.3 | 141.7 | 98.16 | 53.48 | Peak Found | High | Peak Found | High | Not Found | Peak Found | Not Found | High | |
| Creatine kinase U-type, mitochondrial OS=Mus musculus OX=10090 GN=Ckmt1 PE=1 SV=1 | P30275 | 0 | 50.218 | 11 | 2 | 393 | 2 | 418 | 47 | 8.16 | 450.74 | 100 | 6.64385619 | 5.68375E-15 | 242.28 | 198.7 | 1.3 | | 120.63 | High | Not Found | Peak Found | Not Found | Peak Found | High | Not Found | High | |
| CUGBP Elav-like family member 1 OS=Mus musculus OX=10090 GN=Celf1 PE=1 SV=2 | P28659 | 0 | 12.579 | 7 | 2 | 11 | 2 | 486 | 52.1 | 8.46 | 19.52 | 6.033 | 2.592875582 | 0.000294893 | 270.4 | 171.6 | 28.4 | | 124.01 | High | Not Found | High | High | High | Not Found | Not Found | High | Acetyl [N-Term] |
| Cytosolic 10-formyltetrahydrofolate dehydrogenase OS=Mus musculus OX=10090 GN=Aldh1l1 PE=1 SV=1 | Q8R0Y6 | 0 | 72.643 | 11 | 7 | 94 | 6 | 902 | 98.6 | 5.91 | 291.15 | 1.443 | 0.5290713 | 0.012216491 | 94.1 | 115.8 | 84.2 | 99.53 | 71.79 | High | High | High | High | High | High | High | High | |
| Deubiquitinating protein VCIP135 OS=Mus musculus OX=10090 GN=Vcpip1 PE=1 SV=1 | Q8CDG3 | 0 | 24.753 | 5 | 4 | 14 | 4 | 1220 | 134.4 | 7.17 | 39.51 | 0.479 | -1.061902439 | 0.039919274 | 26.36 | 95 | 105 | 72.39 | 74.38 | Peak Found | High | High | High | Peak Found | Peak Found | High | High | |
| Developmentally-regulated GTP-binding protein 2 OS=Mus musculus OX=10090 GN=Drg2 PE=1 SV=1 | Q9QXB9 | 0 | 42.775 | 18 | 5 | 21 | 5 | 364 | 40.7 | 8.88 | 48.76 | 0.459 | -1.123433941 | 0.048877231 | 30.6 | 63 | 137 | 124.22 | 8.52 | Peak Found | High | High | High | Peak Found | High | Peak Found | High | |
| Dihydropyrimidinase-related protein 1 OS=Mus musculus OX=10090 GN=Crmp1 PE=1 SV=1 | P97427 | 0 | 55.425 | 17 | 7 | 61 | 2 | 572 | 62.1 | 7.12 | 118.7 | 0.383 | -1.384583703 | 0.000782935 | 11.23 | 54.3 | 145.7 | 40.1 | 65.13 | High | High | Not Found | Not Found | Not Found | Not Found | Not Found | Not Found | |
| Dipeptidyl peptidase 3 OS=Mus musculus OX=10090 GN=Dpp3 PE=1 SV=2 | Q99KK7 | 0 | 211.362 | 24 | 13 | 265 | 13 | 738 | 82.8 | 5.38 | 722.5 | 0.769 | -0.378944497 | 0.047455484 | 58.08 | 91.7 | 108.3 | 36.37 | 42.85 | High | High | High | High | High | High | High | High | |
| DNA ligase 1 OS=Mus musculus OX=10090 GN=Lig1 PE=1 SV=2 | P37913 | 0 | 13.875 | 3 | 2 | 9 | 2 | 916 | 102.2 | 6.81 | 25.74 | 0.265 | -1.915935735 | 4.69416E-06 | 9.47 | 41.9 | 158.1 | 95.3 | | Not Found | High | High | High | Not Found | High | High | Not Found | |
| DNA replication licensing factor MCM5 OS=Mus musculus OX=10090 GN=Mcm5 PE=1 SV=2 | P49718 | 0 | 22.771 | 8 | 3 | 16 | 3 | 734 | 82.4 | 8.29 | 41.03 | 0.501 | -0.997117491 | 0.040911075 | 24.45 | 69.2 | 130.8 | 106.87 | 9.52 | Peak Found | High | Peak Found | High | High | High | Not Found | High | |
| Drebrin OS=Mus musculus OX=10090 GN=Dbn1 PE=1 SV=4 | Q9QXS6 | 0 | 119.192 | 24 | 10 | 169 | 10 | 706 | 77.2 | 4.49 | 407.71 | 0.77 | -0.377069649 | 0.048877231 | 38.23 | 84.4 | 115.6 | 15.6 | 17.68 | High | High | High | High | High | High | High | High | |
| E3 ubiquitin-protein ligase RNF25 OS=Mus musculus OX=10090 GN=Rnf25 PE=1 SV=2 | Q9QZR0 | 0 | 17.127 | 9 | 3 | 11 | 3 | 456 | 51.2 | 6.29 | 16.9 | 0.4 | -1.321928095 | 0.047682917 | 16.93 | 74.2 | 125.8 | 97.24 | 87.49 | High | High | Peak Found | High | Peak Found | Peak Found | High | High | |
| E3 ubiquitin-protein ligase UBR1 OS=Mus musculus OX=10090 GN=Ubr1 PE=1 SV=2 | O70481 | 0 | 29.473 | 2 | 2 | 35 | 2 | 1757 | 200.1 | 6.05 | 84.85 | 0.415 | -1.268816758 | 0.020444886 | 15.73 | 71.6 | 128.4 | 61.62 | 26.11 | Peak Found | High | High | High | Not Found | High | High | High | |
| Ectonucleoside triphosphate diphosphohydrolase 1 OS=Mus musculus OX=10090 GN=Entpd1 PE=1 SV=1 | P55772 | 0 | 46.021 | 8 | 2 | 30 | 2 | 510 | 57.2 | 5.86 | 78.84 | 1.741 | 0.799916203 | 0.016744742 | 81.63 | 140.8 | 59.2 | 35.87 | 57.78 | Peak Found | High | High | High | Not Found | High | High | High | |
| EH domain-containing protein 2 OS=Mus musculus OX=10090 GN=Ehd2 PE=1 SV=1 | Q8BH64 | 0 | 107.959 | 19 | 8 | 125 | 8 | 543 | 61.1 | 6.51 | 364.53 | 0.708 | -0.498178735 | 0.014529882 | 38.43 | 88.4 | 111.6 | 45.3 | 49.68 | High | High | High | High | High | High | High | High | |
| EH domain-containing protein 3 OS=Mus musculus OX=10090 GN=Ehd3 PE=1 SV=2 | Q9QXY6 | 0 | 48.937 | 15 | 7 | 32 | 2 | 535 | 60.8 | 6.46 | 75.11 | 0.437 | -1.194294815 | 0.017798276 | 16.06 | 27.8 | 172.2 | 138.35 | 13.55 | High | High | High | High | Peak Found | High | Not Found | High | |
| Electrogenic sodium bicarbonate cotransporter 1 OS=Mus musculus OX=10090 GN=Slc4a4 PE=1 SV=2 | O88343 | 0 | 38.594 | 7 | 3 | 20 | 3 | 1079 | 121.4 | 6.84 | 51.9 | 0.598 | -0.74178261 | 0.000128847 | 26.41 | 70.2 | 129.8 | 36.2 | 56.28 | High | High | High | High | Not Found | Peak Found | High | High | |
| Ena/VASP-like protein OS=Mus musculus OX=10090 GN=Evl PE=1 SV=2 | P70429 | 0 | 11.834 | 11 | 2 | 8 | 2 | 414 | 44.3 | 8.85 | 16.93 | 4.213 | 2.074847916 | 0.001127308 | 128.92 | 152 | 48 | 118.69 | 105.66 | Peak Found | High | High | High | Peak Found | Peak Found | Peak Found | High | |
| Ephrin-B1 OS=Mus musculus OX=10090 GN=Efnb1 PE=1 SV=1 | P52795 | 0 | 14.78 | 10 | 3 | 11 | 3 | 345 | 37.8 | 9.03 | 16.06 | 0.146 | -2.775959726 | 9.10755E-08 | 7.44 | 24 | 176 | 131.84 | 9.06 | High | High | High | Peak Found | High | High | Not Found | High | |
| ERO1-like protein alpha OS=Mus musculus OX=10090 GN=Ero1a PE=1 SV=2 | Q8R180 | 0 | 105.799 | 28 | 9 | 159 | 9 | 464 | 54.1 | 6.54 | 422.86 | 1.555 | 0.63691458 | 0.000270773 | 123.74 | 129.8 | 70.2 | 70.68 | 60.37 | Peak Found | High | High | High | Not Found | High | High | High | |
| Excitatory amino acid transporter 2 OS=Mus musculus OX=10090 GN=Slc1a2 PE=1 SV=1 | P43006 | 0 | 33.094 | 10 | 4 | 53 | 4 | 572 | 62 | 6.68 | 72.53 | 6.388 | 2.675364313 | 0.000120061 | 119.06 | 172.9 | 27.1 | 31.52 | 100.36 | High | Not Found | High | Not Found | Not Found | High | High | Not Found | |
| Fatty acid-binding protein 5 OS=Mus musculus OX=10090 GN=Fabp5 PE=1 SV=3 | Q05816 | 0 | 310.079 | 82 | 13 | 318 | 13 | 135 | 15.1 | 6.54 | 744.83 | 1.688 | 0.755314904 | 3.89691E-08 | 115.03 | 142.1 | 57.9 | 96.1 | 158.2 | Peak Found | High | High | High | Peak Found | High | High | High | |
| Fatty acid-binding protein, adipocyte OS=Mus musculus OX=10090 GN=Fabp4 PE=1 SV=3 | P04117 | 0 | 63.46 | 38 | 5 | 75 | 5 | 132 | 14.6 | 8.4 | 159.24 | 1.715 | 0.778208576 | 0.000738611 | 134.79 | 130.6 | 69.4 | 119.58 | 50.9 | High | High | High | High | High | High | High | High | |
| F-box/WD repeat-containing protein 8 OS=Mus musculus OX=10090 GN=Fbxw8 PE=1 SV=2 | Q8BIA4 | 0 | 43.478 | 13 | 5 | 33 | 5 | 598 | 67.9 | 6.16 | 67.71 | 0.394 | -1.343732465 | 0.000782935 | 12.17 | 60.3 | 139.7 | 133.33 | 34.88 | High | High | High | High | High | High | Peak Found | High | |
| Fibulin-1 OS=Mus musculus OX=10090 GN=Fbln1 PE=1 SV=2 | Q08879 | 0 | 159.252 | 25 | 10 | 263 | 10 | 705 | 78 | 5.16 | 606.09 | 0.764 | -0.388355457 | 0.040323389 | 58.58 | 86 | 114 | 75.02 | 43.89 | High | High | High | High | High | High | High | High | |
| Formin-like protein 1 OS=Mus musculus OX=10090 GN=Fmnl1 PE=1 SV=1 | Q9JL26 | 0 | 40.896 | 7 | 5 | 40 | 5 | 1094 | 122 | 5.82 | 98.99 | 1.821 | 0.864730922 | 0.014529882 | 66.26 | 130.4 | 69.6 | 17.82 | 100.81 | High | High | High | High | High | High | High | High | |
| Glutamyl aminopeptidase OS=Mus musculus OX=10090 GN=Enpep PE=1 SV=1 | P16406 | 0 | 50.966 | 10 | 6 | 57 | 6 | 945 | 107.9 | 5.44 | 116.17 | 0.59 | -0.76121314 | 0.01970772 | 36.46 | 84.7 | 115.3 | 84.11 | 64.44 | High | High | Peak Found | High | High | High | Peak Found | High | |
| Glutaredoxin-1 OS=Mus musculus OX=10090 GN=Glrx PE=1 SV=3 | Q9QUH0 | 0 | 44.681 | 31 | 3 | 47 | 3 | 107 | 11.9 | 8.37 | 153.4 | 1.746 | 0.804053559 | 0.025810784 | 137.46 | 155.4 | 44.6 | 55.44 | 107.65 | High | High | High | High | Peak Found | High | High | High | |
| Glutathione S-transferase A4 OS=Mus musculus OX=10090 GN=Gsta4 PE=1 SV=3 | P24472 | 0.001 | 4.648 | 5 | 2 | 2 | 2 | 222 | 25.5 | 7.39 | 1.88 | 7.244 | 2.856786546 | 2.04912E-05 | 401.11 | 175.7 | 24.3 | | 132.65 | Peak Found | Not Found | High | Not Found | High | Peak Found | Not Found | Not Found | |
| Glycerol-3-phosphate dehydrogenase [NAD(+)], cytoplasmic OS=Mus musculus OX=10090 GN=Gpd1 PE=1 SV=3 | P13707 | 0 | 10.463 | 14 | 3 | 6 | 3 | 349 | 37.5 | 7.17 | 11.94 | 0.103 | -3.279283757 | 5.68375E-15 | 11 | 18.6 | 181.4 | 140.01 | | High | High | Not Found | High | Not Found | Peak Found | Not Found | Not Found | |
| Glycogen phosphorylase, liver form OS=Mus musculus OX=10090 GN=Pygl PE=1 SV=4 | Q9ET01 | 0 | 122.773 | 19 | 14 | 126 | 10 | 850 | 97.4 | 7.09 | 269.31 | 1.501 | 0.585923977 | 0.004020027 | 89.62 | 133 | 67 | 41.24 | 52.42 | High | High | High | High | High | High | High | High | |
| GTPase-activating protein and VPS9 domain-containing protein 1 OS=Mus musculus OX=10090 GN=Gapvd1 PE=1 SV=2 | Q6PAR5 | 0 | 45.858 | 6 | 4 | 26 | 4 | 1458 | 162.3 | 5.19 | 80.85 | 0.619 | -0.691988685 | 0.047740226 | 44.03 | 75.9 | 124.1 | 90.48 | 42.6 | Peak Found | High | High | Peak Found | Not Found | Peak Found | High | Peak Found | |
| GTP-binding nuclear protein Ran OS=Mus musculus OX=10090 GN=Ran PE=1 SV=3 | P62827 | 0 | 37.923 | 36 | 5 | 21 | 5 | 216 | 24.4 | 7.49 | 41.04 | 1.436 | 0.522055749 | 0.027234364 | 83.4 | 118.4 | 81.6 | 84.17 | 82.31 | High | High | Peak Found | High | Not Found | High | Peak Found | High | |
| Histone H2B type 3-A OS=Mus musculus OX=10090 GN=Hist3h2ba PE=1 SV=3 | Q9D2U9 | 0 | 127.669 | 40 | 4 | 446 | 1 | 126 | 14 | 10.37 | 1352.67 | 0.147 | -2.76611194 | 3.05671E-06 | 11.88 | 25.6 | 174.4 | 136.59 | | Not Found | Peak Found | Peak Found | Not Found | Not Found | Not Found | High | Not Found | |
| Hydroxymethylglutaryl-CoA synthase, cytoplasmic OS=Mus musculus OX=10090 GN=Hmgcs1 PE=1 SV=1 | Q8JZK9 | 0 | 98.224 | 21 | 9 | 76 | 9 | 520 | 57.5 | 5.99 | 198.66 | 0.546 | -0.873027144 | 0.000599129 | 27.44 | 87.7 | 112.3 | 51.95 | 76.71 | High | High | High | High | High | High | Peak Found | High | |
| Immunoglobulin superfamily member 8 OS=Mus musculus OX=10090 GN=Igsf8 PE=1 SV=2 | Q8R366 | 0 | 39.109 | 11 | 4 | 36 | 4 | 611 | 65 | 7.99 | 82.92 | 0.528 | -0.921390165 | 0.036769274 | 35.03 | 71.9 | 128.1 | 109.41 | 39.89 | High | High | High | High | High | High | High | High | |
| Inactive glycosyltransferase 25 family member 3 OS=Mus musculus OX=10090 GN=Cercam PE=3 SV=1 | A3KGW5 | 0 | 9.916 | 4 | 2 | 4 | 2 | 592 | 67.6 | 6.76 | 9.79 | 0.377 | -1.407363571 | 0.004829798 | 7.55 | 54.8 | 145.2 | 49.29 | 5.57 | Not Found | High | Peak Found | High | Not Found | High | Not Found | Peak Found | |
| Interferon-induced helicase C domain-containing protein 1 OS=Mus musculus OX=10090 GN=Ifih1 PE=1 SV=1 | Q8R5F7 | 0 | 13.783 | 3 | 2 | 7 | 2 | 1025 | 115.9 | 6.25 | 13.94 | 0.365 | -1.454031631 | 0.00240035 | 12.66 | 42.6 | 157.4 | 56.82 | 78.47 | Peak Found | High | High | Peak Found | Not Found | High | Peak Found | Peak Found | |
| Isoform 2 of Tyrosine-protein phosphatase non-receptor type substrate 1 OS=Mus musculus OX=10090 GN=Sirpa | P97797-2 | 0 | 70.482 | 16 | 4 | 69 | 1 | 509 | 56 | 8.28 | 180.73 | 1.799 | 0.847195187 | 0.008978805 | 115.44 | 121.7 | 78.3 | 100.8 | 110.45 | High | High | High | High | Not Found | High | High | High | |
| Isoform Alpha-7X2B of Integrin alpha-7 OS=Mus musculus OX=10090 GN=Itga7 | Q61738-5 | 0 | 32.349 | 4 | 3 | 28 | 3 | 1135 | 124.6 | 5.91 | 78.96 | 1.795 | 0.843983844 | 0.016400621 | 88.69 | 136.9 | 63.1 | 68.14 | 81.61 | High | High | High | High | High | High | High | High | |
| Keratin, type II cuticular Hb5 OS=Mus musculus OX=10090 GN=Krt85 PE=1 SV=2 | Q9Z2T6 | 0 | 6.73 | 6 | 2 | 9 | 1 | 507 | 55.7 | 6.42 | 14.16 | 3.028 | 1.598365205 | 6.4532E-05 | 79.59 | 136.6 | 63.4 | 77.76 | 41.55 | Not Found | High | Peak Found | Peak Found | Peak Found | Peak Found | High | Peak Found | |
| Keratin, type II cytoskeletal 5 OS=Mus musculus OX=10090 GN=Krt5 PE=1 SV=1 | Q922U2 | 0 | 12.843 | 4 | 2 | 32 | 1 | 580 | 61.7 | 7.75 | 74.1 | 0.039 | -4.680382066 | 5.68375E-15 | 0.08 | 7.5 | 192.5 | 66.2 | | High | Peak Found | Not Found | Peak Found | Peak Found | Not Found | Peak Found | Not Found | |
| Kinesin-like protein KIF13A OS=Mus musculus OX=10090 GN=Kif13a PE=1 SV=1 | Q9EQW7 | 0 | 9.586 | 1 | 2 | 13 | 1 | 1749 | 195.7 | 5.41 | 31.19 | 0.361 | -1.469929258 | 0.026666282 | 4.63 | 58 | 142 | 27.31 | 29.93 | Not Found | Peak Found | Peak Found | Peak Found | Not Found | Peak Found | High | Not Found | |
| Legumain OS=Mus musculus OX=10090 GN=Lgmn PE=1 SV=1 | O89017 | 0 | 111.901 | 20 | 8 | 154 | 8 | 435 | 49.3 | 6.39 | 380.25 | 1.377 | 0.461528559 | 0.003194653 | 99.13 | 125.7 | 74.3 | 68.43 | 54.79 | High | High | High | High | High | High | High | High | |
| Lysozyme C-2 OS=Mus musculus OX=10090 GN=Lyz2 PE=1 SV=2 | P08905 | 0 | 104.657 | 50 | 5 | 155 | 5 | 148 | 16.7 | 8.81 | 437.64 | 1.628 | 0.7031007 | 4.06082E-07 | 100.88 | 113.7 | 86.3 | 35.49 | 39.37 | High | High | High | High | High | High | High | High | |
| Macrophage-expressed gene 1 protein OS=Mus musculus OX=10090 GN=Mpeg1 PE=1 SV=1 | A1L314 | 0 | 19.217 | 6 | 3 | 24 | 3 | 713 | 78.3 | 5.92 | 56.67 | 0.427 | -1.227692025 | 0.019886736 | 19.56 | 67.4 | 132.6 | 84.24 | 99.27 | High | High | High | High | High | High | Peak Found | Peak Found | |
| Mannosyl-oligosaccharide 1,2-alpha-mannosidase IA OS=Mus musculus OX=10090 GN=Man1a1 PE=1 SV=1 | P45700 | 0 | 22.276 | 5 | 2 | 12 | 2 | 655 | 73.2 | 6.81 | 25.49 | 2.926 | 1.548929769 | 3.16161E-05 | 105.97 | 149.1 | 50.9 | | 100.37 | Not Found | High | Peak Found | Peak Found | Not Found | High | Peak Found | High | |
| Metalloreductase STEAP4 OS=Mus musculus OX=10090 GN=Steap4 PE=1 SV=1 | Q923B6 | 0 | 34.493 | 12 | 3 | 27 | 3 | 470 | 53 | 9.17 | 51.47 | 0.477 | -1.067938829 | 0.001457049 | 21.06 | 60.6 | 139.4 | 107.28 | 47.6 | Peak Found | High | High | High | High | High | Peak Found | High | |
| Methylcrotonoyl-CoA carboxylase subunit alpha, mitochondrial OS=Mus musculus OX=10090 GN=Mccc1 PE=1 SV=2 | Q99MR8 | 0 | 15.286 | 5 | 2 | 9 | 2 | 717 | 79.3 | 7.83 | 21.53 | 3.097 | 1.630871383 | 1.50448E-06 | 140.15 | 156.2 | 43.8 | 100.27 | 87.67 | High | High | High | High | Not Found | High | Peak Found | High | |
| Multidrug resistance protein 1A OS=Mus musculus OX=10090 GN=Abcb1a PE=1 SV=3 | P21447 | 0 | 110.019 | 11 | 9 | 121 | 9 | 1276 | 140.6 | 8.85 | 291.28 | 0.638 | -0.648371671 | 0.009375864 | 34.22 | 80.3 | 119.7 | 15.27 | 56.04 | High | High | High | High | High | High | High | High | Acetyl [N-Term] |
| Myristoylated alanine-rich C-kinase substrate OS=Mus musculus OX=10090 GN=Marcks PE=1 SV=2 | P26645 | 0 | 133.726 | 75 | 8 | 279 | 8 | 309 | 29.6 | 4.34 | 510.22 | 1.324 | 0.404903122 | 0.014529882 | 122.14 | 109.2 | 90.8 | 86.75 | 105.69 | High | High | High | High | High | High | High | High | |
| N-acetylgalactosamine kinase OS=Mus musculus OX=10090 GN=Galk2 PE=1 SV=1 | Q68FH4 | 0 | 69.261 | 23 | 6 | 60 | 6 | 458 | 50.5 | 6.9 | 174.1 | 1.696 | 0.76213617 | 0.034935407 | 78.34 | 126.9 | 73.1 | 46.05 | 91.74 | High | High | High | High | Peak Found | High | High | High | |
| N-acetylglucosamine-6-phosphate deacetylase OS=Mus musculus OX=10090 GN=Amdhd2 PE=1 SV=1 | Q8JZV7 | 0 | 15.448 | 8 | 2 | 16 | 2 | 409 | 43.5 | 6.23 | 6.74 | 1.945 | 0.959770155 | 0.034578825 | 61.13 | 138.8 | 61.2 | 66.2 | 24.16 | Not Found | High | High | High | Not Found | High | Not Found | High | |
| NADH-cytochrome b5 reductase 1 OS=Mus musculus OX=10090 GN=Cyb5r1 PE=1 SV=1 | Q9DB73 | 0 | 43.113 | 12 | 3 | 35 | 3 | 305 | 34.1 | 8.87 | 91.01 | 1.834 | 0.874993639 | 0.000240007 | 66.11 | 128.1 | 71.9 | 3.21 | 73.09 | Not Found | High | High | High | Not Found | High | High | High | |
| Neurofilament heavy polypeptide OS=Mus musculus OX=10090 GN=Nefh PE=1 SV=3 | P19246 | 0 | 65.767 | 7 | 5 | 35 | 3 | 1090 | 116.9 | 5.81 | 63.8 | 0.564 | -0.826232932 | 0.00729236 | 18.58 | 64.6 | 135.4 | 54.02 | 20.62 | Not Found | High | High | High | Not Found | Peak Found | Peak Found | High | |
| Neurofilament light polypeptide OS=Mus musculus OX=10090 GN=Nefl PE=1 SV=5 | P08551 | 0 | 190.618 | 38 | 18 | 192 | 16 | 543 | 61.5 | 4.64 | 443.45 | 2.075 | 1.053111336 | 5.68375E-15 | 92.25 | 135.7 | 64.3 | 64.76 | 80.2 | High | High | High | High | High | High | High | High | |
| Neurofilament medium polypeptide OS=Mus musculus OX=10090 GN=Nefm PE=1 SV=4 | P08553 | 0 | 192.415 | 39 | 21 | 138 | 18 | 848 | 95.9 | 4.77 | 298.06 | 1.455 | 0.541019153 | 0.009928181 | 87.06 | 135.9 | 64.1 | 49.73 | 49.1 | High | High | High | High | High | High | High | High | |
| N-myc-interactor OS=Mus musculus OX=10090 GN=Nmi PE=1 SV=1 | O35309 | 0 | 22.303 | 16 | 4 | 13 | 4 | 314 | 35.2 | 5.05 | 35.23 | 0.502 | -0.994240731 | 0.043579339 | 27.8 | 52.9 | 147.1 | 100.5 | 92.02 | Peak Found | High | Peak Found | High | Peak Found | High | Not Found | High | |
| Nuclear distribution protein nudE homolog 1 OS=Mus musculus OX=10090 GN=Nde1 PE=1 SV=1 | Q9CZA6 | 0 | 8.182 | 9 | 2 | 3 | 1 | 344 | 38.5 | 5.34 | 10.91 | 0.405 | -1.304006187 | 0.005598168 | 18.18 | 57.6 | 142.4 | | 109.31 | Not Found | Not Found | High | Not Found | Not Found | Peak Found | Peak Found | Peak Found | |
| Nuclear protein localization protein 4 homolog OS=Mus musculus OX=10090 GN=Nploc4 PE=1 SV=3 | P60670 | 0 | 82.907 | 12 | 5 | 102 | 5 | 608 | 68 | 6.46 | 298.14 | 0.695 | -0.524915117 | 0.031582936 | 50.87 | 100.4 | 99.6 | 39.93 | 22.31 | High | High | High | High | High | High | High | High | |
| Nuclear receptor corepressor 2 OS=Mus musculus OX=10090 GN=Ncor2 PE=1 SV=3 | Q9WU42 | 0 | 18.267 | 3 | 3 | 8 | 3 | 2472 | 269.6 | 7.44 | 17.21 | 0.311 | -1.685013515 | 0.044397852 | 10.58 | 47.4 | 152.6 | 140.04 | 92.42 | High | Peak Found | High | High | Peak Found | Peak Found | High | High | |
| Nucleolar protein 16 OS=Mus musculus OX=10090 GN=Nop16 PE=1 SV=1 | Q9CPT5 | 0 | 12.711 | 19 | 2 | 5 | 2 | 178 | 21.1 | 9.91 | 9.06 | 0.208 | -2.265344567 | 4.05862E-06 | 3.93 | 40.2 | 159.8 | 67.77 | 0.33 | Not Found | High | Peak Found | High | Not Found | Peak Found | Not Found | High | |
| Nucleolar transcription factor 1 OS=Mus musculus OX=10090 GN=Ubtf PE=1 SV=1 | P25976 | 0 | 36.772 | 7 | 4 | 21 | 4 | 765 | 89.5 | 5.76 | 50.28 | 0.409 | -1.289827252 | 0.043579339 | 19.93 | 46.8 | 153.2 | 54.97 | 116.83 | High | High | High | High | Peak Found | High | Peak Found | High | |
| Peripherin OS=Mus musculus OX=10090 GN=Prph PE=1 SV=2 | P15331 | 0 | 321.744 | 37 | 20 | 466 | 16 | 475 | 54.2 | 5.52 | 1171.7 | 1.647 | 0.719840555 | 1.83689E-07 | 103.79 | 119.2 | 80.8 | 76.46 | 70.62 | High | High | High | High | High | High | High | High | |
| Phosphatidylinositol 3,4,5-trisphosphate 5-phosphatase 1 OS=Mus musculus OX=10090 GN=Inpp5d PE=1 SV=2 | Q9ES52 | 0 | 13.455 | 4 | 3 | 4 | 3 | 1191 | 133.5 | 7.8 | 10.97 | 0.259 | -1.948975997 | 0.000167847 | 4.27 | 41.1 | 158.9 | 88.73 | | Not Found | Peak Found | High | Peak Found | Not Found | Not Found | High | Not Found | |
| Phosphatidylinositol 4-phosphate 3-kinase C2 domain-containing subunit alpha OS=Mus musculus OX=10090 GN=Pik3c2a PE=1 SV=2 | Q61194 | 0 | 17.97 | 4 | 4 | 9 | 4 | 1686 | 190.6 | 8.02 | 15.12 | 0.483 | -1.049904906 | 0.040323389 | 18.11 | 60.6 | 139.4 | 102.66 | 47.3 | High | High | High | High | Peak Found | Peak Found | Peak Found | Peak Found | |
| Phosphofurin acidic cluster sorting protein 1 OS=Mus musculus OX=10090 GN=Pacs1 PE=1 SV=2 | Q8K212 | 0 | 13.635 | 4 | 3 | 5 | 3 | 961 | 104.8 | 7.74 | 12.12 | 0.174 | -2.522840789 | 5.90417E-07 | 3.58 | 96.8 | 103.2 | 109.8 | 83.08 | High | Peak Found | High | Peak Found | Peak Found | High | Not Found | Peak Found | |
| Phosphoglucomutase-1 OS=Mus musculus OX=10090 GN=Pgm1 PE=1 SV=4 | Q9D0F9 | 0 | 125.168 | 31 | 16 | 128 | 16 | 562 | 61.4 | 6.57 | 274.08 | 1.363 | 0.446785562 | 0.005177686 | 96.35 | 108.4 | 91.6 | 39.59 | 59.18 | High | High | High | High | High | High | High | High | |
| Phosphoglucomutase-2 OS=Mus musculus OX=10090 GN=Pgm2 PE=1 SV=1 | Q7TSV4 | 0 | 72.738 | 15 | 6 | 55 | 6 | 620 | 68.7 | 6.14 | 97.28 | 1.502 | 0.586884813 | 0.036678803 | 100.65 | 144.7 | 55.3 | 67.36 | 82.73 | Peak Found | High | High | High | Peak Found | High | High | High | |
| Phosphoglycerate mutase 1 OS=Mus musculus OX=10090 GN=Pgam1 PE=1 SV=3 | Q9DBJ1 | 0 | 202.722 | 61 | 13 | 296 | 13 | 254 | 28.8 | 7.18 | 680.52 | 1.413 | 0.498761466 | 0.000967667 | 104.37 | 108.7 | 91.3 | 74.58 | 82.38 | High | High | High | High | High | High | High | High | |
| Phosphoinositide 3-kinase adapter protein 1 OS=Mus musculus OX=10090 GN=Pik3ap1 PE=1 SV=1 | Q9EQ32 | 0 | 24.92 | 7 | 3 | 23 | 3 | 811 | 90.9 | 5.17 | 56.01 | 1.886 | 0.915329676 | 0.006417946 | 53.92 | 130.7 | 69.3 | 21.56 | 70.22 | High | High | High | High | Peak Found | High | High | High | |
| Plexin-A1 OS=Mus musculus OX=10090 GN=Plxna1 PE=1 SV=1 | P70206 | 0 | 7.685 | 3 | 3 | 5 | 2 | 1894 | 211 | 6.9 | 3.81 | 0.216 | -2.210896782 | 0.005598168 | 2.74 | 35.5 | 164.5 | 102.22 | | High | Peak Found | Not Found | Peak Found | Peak Found | Not Found | Not Found | Not Found | |
| Pre-rRNA-processing protein TSR1 homolog OS=Mus musculus OX=10090 GN=Tsr1 PE=1 SV=1 | Q5SWD9 | 0 | 33.881 | 8 | 5 | 21 | 5 | 803 | 92 | 6.99 | 49.24 | 2.683 | 1.423847055 | 0.009307628 | 73.67 | 135.6 | 64.4 | 61.54 | 56.76 | High | High | High | High | High | High | High | Peak Found | |
| PRKC apoptosis WT1 regulator protein OS=Mus musculus OX=10090 GN=Pawr PE=1 SV=2 | Q925B0 | 0 | 21.196 | 12 | 2 | 29 | 2 | 333 | 35.9 | 5.92 | 65.03 | 0.507 | -0.979942348 | 0.001203097 | 28.32 | 60.5 | 139.5 | 113.61 | 60.34 | High | High | High | High | High | High | High | High | |
| Probable leucine--tRNA ligase, mitochondrial OS=Mus musculus OX=10090 GN=Lars2 PE=1 SV=1 | Q8VDC0 | 0 | 24.629 | 6 | 4 | 20 | 4 | 902 | 101.4 | 8.19 | 38.86 | 2.417 | 1.273217473 | 0.026772883 | 121.12 | 112 | 88 | 81.42 | 76.94 | Peak Found | High | Not Found | High | Not Found | High | Peak Found | High | |
| Pro-cathepsin H OS=Mus musculus OX=10090 GN=Ctsh PE=1 SV=2 | P49935 | 0 | 75.898 | 33 | 6 | 76 | 6 | 333 | 37.1 | 8.4 | 185.44 | 0.735 | -0.444183845 | 0.018795489 | 50.21 | 94 | 106 | 35.14 | 48.63 | High | High | High | High | High | High | High | High | |
| Protein arginine N-methyltransferase 5 OS=Mus musculus OX=10090 GN=Prmt5 PE=1 SV=3 | Q8CIG8 | 0 | 27.042 | 5 | 2 | 23 | 2 | 637 | 72.6 | 6.42 | 64.92 | 0.576 | -0.795859283 | 0.01231637 | 27.98 | 82.8 | 117.2 | 51.99 | 57.05 | Not Found | High | High | High | Not Found | High | Peak Found | High | |
| Protein ELYS OS=Mus musculus OX=10090 GN=Ahctf1 PE=1 SV=1 | Q8CJF7 | 0 | 20.748 | 1 | 2 | 15 | 2 | 2243 | 247.5 | 6.55 | 41.25 | 0.405 | -1.304006187 | 0.014970986 | 1.88 | 57.6 | 142.4 | 50.26 | | High | High | High | High | High | High | High | High | |
| Protein phosphatase 1 regulatory subunit 21 OS=Mus musculus OX=10090 GN=Ppp1r21 PE=1 SV=2 | Q3TDD9 | 0 | 67.909 | 20 | 9 | 29 | 9 | 780 | 88.3 | 6.9 | 64.85 | 0.483 | -1.049904906 | 0.012216491 | 28.06 | 78.4 | 121.6 | 75.55 | 65.12 | High | High | High | High | High | High | High | High | |
| Protein scribble homolog OS=Mus musculus OX=10090 GN=Scrib PE=1 SV=2 | Q80U72 | 0 | 56.201 | 6 | 5 | 46 | 4 | 1612 | 174 | 5.12 | 137.95 | 0.652 | -0.61705613 | 0.014970986 | 32.52 | 96.5 | 103.5 | 61.27 | 65.46 | High | Peak Found | High | High | Peak Found | High | High | High | |
| PSME3-interacting protein OS=Mus musculus OX=10090 GN=Fam192a PE=1 SV=1 | Q91WE2 | 0 | 20.182 | 11 | 2 | 23 | 2 | 254 | 28.7 | 5.14 | 47.82 | 1.769 | 0.822934048 | 0.01917508 | 103.64 | 122.9 | 77.1 | 59.67 | 105.89 | High | High | High | High | High | Peak Found | High | High | Acetyl [N-Term] |
| Pumilio homolog 3 OS=Mus musculus OX=10090 GN=Pum3 PE=1 SV=2 | Q8BKS9 | 0 | 35.251 | 7 | 3 | 14 | 3 | 647 | 72.8 | 9.66 | 40.95 | 0.238 | -2.070966521 | 1.26604E-06 | 5.56 | 23.6 | 176.4 | 150.36 | 64.78 | Peak Found | High | Peak Found | High | Not Found | High | Peak Found | High | |
| Pyruvate dehydrogenase E1 component subunit alpha, somatic form, mitochondrial OS=Mus musculus OX=10090 GN=Pdha1 PE=1 SV=1 | P35486 | 0 | 43.644 | 14 | 4 | 24 | 4 | 390 | 43.2 | 8.19 | 82.7 | 0.442 | -1.177881725 | 0.000442534 | 27.08 | 46.9 | 153.1 | 125.76 | 74.32 | Peak Found | High | High | Peak Found | Not Found | Peak Found | High | Peak Found | |
| Ragulator complex protein LAMTOR2 OS=Mus musculus OX=10090 GN=Lamtor2 PE=1 SV=1 | Q9JHS3 | 0 | 16.142 | 26 | 2 | 11 | 2 | 125 | 13.5 | 5.4 | 23.32 | 0.326 | -1.61705613 | 0.000276851 | 7.78 | 49.2 | 150.8 | 88.42 | 14.98 | Not Found | Peak Found | Peak Found | High | Not Found | Peak Found | Not Found | Peak Found | |
| Ras-related protein Rab-3D OS=Mus musculus OX=10090 GN=Rab3d PE=1 SV=1 | P35276 | 0 | 33.393 | 34 | 6 | 28 | 3 | 219 | 24.4 | 4.93 | 50.74 | 2.475 | 1.307428525 | 1.30156E-05 | 88.45 | 142.4 | 57.6 | 36.17 | 76.98 | High | High | High | High | Not Found | High | High | High | |
| Ras-related protein Rab-4A OS=Mus musculus OX=10090 GN=Rab4a PE=1 SV=2 | P56371 | 0 | 32.953 | 38 | 5 | 22 | 3 | 218 | 24.4 | 6.07 | 47.81 | 0.351 | -1.510457064 | 0.001457049 | 10.72 | 57.9 | 142.1 | 56.55 | 56.22 | High | High | Peak Found | High | Peak Found | High | Peak Found | High | |
| Ras-related protein Rab-8A OS=Mus musculus OX=10090 GN=Rab8a PE=1 SV=2 | P55258 | 0 | 25.694 | 13 | 3 | 20 | 1 | 207 | 23.7 | 9.07 | 51.22 | 0.65 | -0.621488377 | 0.045466815 | 68.28 | 94.8 | 105.2 | 29.3 | 98.75 | Peak Found | High | High | High | High | High | High | High | |
| Rho GTPase-activating protein 5 OS=Mus musculus OX=10090 GN=Arhgap5 PE=1 SV=2 | P97393 | 0 | 15.943 | 2 | 2 | 22 | 2 | 1501 | 172 | 6.34 | 48.47 | 0.516 | -0.954557029 | 0.010432585 | 28.71 | 45.1 | 154.9 | 124.19 | 67.18 | High | High | High | High | High | High | High | High | |
| Ribosomal protein S6 kinase alpha-1 OS=Mus musculus OX=10090 GN=Rps6ka1 PE=1 SV=1 | P18653 | 0 | 17.088 | 6 | 3 | 6 | 2 | 724 | 81.5 | 8.1 | 12.68 | 0.318 | -1.652901329 | 0.012129266 | 44.82 | 48.3 | 151.7 | | 134.08 | Not Found | High | High | Not Found | Not Found | Peak Found | Not Found | Peak Found | |
| Ribulose-phosphate 3-epimerase OS=Mus musculus OX=10090 GN=Rpe PE=1 SV=1 | Q8VEE0 | 0 | 41.234 | 25 | 3 | 19 | 3 | 228 | 24.9 | 5.41 | 59.42 | 0.244 | -2.035046947 | 1.31495E-07 | 8.31 | 43.7 | 156.3 | 94.69 | 74.73 | Not Found | High | High | High | Peak Found | High | High | High | |
| RNA-binding protein with serine-rich domain 1 OS=Mus musculus OX=10090 GN=Rnps1 PE=1 SV=1 | Q99M28 | 0 | 38.924 | 11 | 4 | 57 | 4 | 305 | 34.2 | 11.84 | 123.88 | 1.651 | 0.72334012 | 0.049952191 | 107.89 | 113.7 | 86.3 | 98.87 | 94.52 | High | High | High | High | High | High | High | High | |
| Septin-5 OS=Mus musculus OX=10090 GN=Sept5 PE=1 SV=2 | Q9Z2Q6 | 0 | 25.213 | 15 | 4 | 22 | 4 | 369 | 42.7 | 6.67 | 45.66 | 0.26 | -1.943416472 | 0.000848147 | 4.4 | 41.2 | 158.8 | 154.83 | | High | High | High | High | High | High | Peak Found | High | |
| Serine/threonine-protein phosphatase 6 regulatory subunit 1 OS=Mus musculus OX=10090 GN=Ppp6r1 PE=1 SV=1 | Q7TSI3 | 0 | 45.365 | 13 | 5 | 29 | 5 | 856 | 94.5 | 4.64 | 73.72 | 1.835 | 0.875780063 | 0.019369027 | 80.7 | 112.4 | 87.6 | 89.64 | 15.15 | High | High | High | High | Not Found | High | Peak Found | High | |
| SH3 and PX domain-containing protein 2B OS=Mus musculus OX=10090 GN=Sh3pxd2b PE=1 SV=1 | A2AAY5 | 0 | 19.798 | 5 | 2 | 8 | 2 | 908 | 101.5 | 8.66 | 27.33 | 3.462 | 1.791605725 | 0.025810784 | 101.75 | 155.4 | 44.6 | 56.31 | 139.13 | Peak Found | High | High | Not Found | Not Found | High | Peak Found | Peak Found | |
| SH3 domain-binding glutamic acid-rich-like protein OS=Mus musculus OX=10090 GN=Sh3bgrl PE=1 SV=1 | Q9JJU8 | 0 | 73.083 | 84 | 8 | 100 | 8 | 114 | 12.8 | 4.92 | 217.55 | 0.744 | -0.426625474 | 0.01607704 | 64.24 | 102.9 | 97.1 | 36.66 | 46.19 | High | High | High | High | High | High | High | High | |
| Sideroflexin-2 OS=Mus musculus OX=10090 GN=Sfxn2 PE=1 SV=1 | Q925N2 | 0 | 11.71 | 8 | 2 | 12 | 2 | 322 | 36.1 | 8.82 | 26.49 | 0.36 | -1.473931188 | 0.004706589 | 11.56 | 68.5 | 131.5 | 57.31 | 77.56 | High | High | High | High | High | High | Peak Found | High | Acetyl [N-Term] |
| Signal transducer and activator of transcription 5B OS=Mus musculus OX=10090 GN=Stat5b PE=1 SV=1 | P42232 | 0 | 28.4 | 7 | 4 | 24 | 2 | 786 | 89.9 | 6.05 | 73.78 | 0.466 | -1.10159814 | 0.00775693 | 29.1 | 110.2 | 89.8 | 26.03 | 51.49 | Not Found | High | Peak Found | High | Not Found | High | High | High | |
| Slit homolog 2 protein OS=Mus musculus OX=10090 GN=Slit2 PE=2 SV=2 | Q9R1B9 | 0 | 35.35 | 6 | 5 | 22 | 5 | 1521 | 168.7 | 7.11 | 62.4 | 0.073 | -3.775959726 | 6.73091E-13 | 1.32 | 13.6 | 186.4 | 132.15 | 59.4 | Not Found | High | High | High | Not Found | High | High | High | |
| Sodium-coupled neutral amino acid transporter 2 OS=Mus musculus OX=10090 GN=Slc38a2 PE=1 SV=1 | Q8CFE6 | 0 | 11.202 | 9 | 2 | 6 | 2 | 504 | 55.5 | 7.94 | 2.13 | 0.388 | -1.365871442 | 0.045362278 | 12.34 | 55.9 | 144.1 | 102.03 | 20.3 | High | Peak Found | Peak Found | Peak Found | Peak Found | High | Not Found | High | |
| Son of sevenless homolog 1 OS=Mus musculus OX=10090 GN=Sos1 PE=1 SV=2 | Q62245 | 0 | 15.389 | 3 | 3 | 8 | 3 | 1319 | 150.8 | 6.89 | 18.43 | 0.333 | -1.586405918 | 0.004888218 | 11.75 | 39.5 | 160.5 | 126.32 | 89.29 | High | High | High | High | High | Not Found | Peak Found | Peak Found | |
| Sorting nexin-17 OS=Mus musculus OX=10090 GN=Snx17 PE=1 SV=2 | Q8BVL3 | 0 | 20.527 | 7 | 2 | 11 | 2 | 470 | 52.8 | 7.49 | 26.56 | 0.378 | -1.40354186 | 0.000120061 | 13.68 | 65.3 | 134.7 | 41.65 | 107.06 | Peak Found | High | Peak Found | High | Peak Found | High | Peak Found | Peak Found | |
| Sterile alpha motif domain-containing protein 9-like OS=Mus musculus OX=10090 GN=Samd9l PE=1 SV=2 | Q69Z37 | 0 | 54.364 | 6 | 5 | 34 | 5 | 1561 | 180.1 | 6.99 | 94.45 | 0.47 | -1.089267338 | 0.001026559 | 16.62 | 114.8 | 85.2 | 51.64 | 103.32 | High | High | High | High | High | High | Peak Found | High | |
| Superoxide dismutase [Mn], mitochondrial OS=Mus musculus OX=10090 GN=Sod2 PE=1 SV=3 | P09671 | 0 | 21.646 | 16 | 2 | 20 | 2 | 222 | 24.6 | 8.62 | 63.2 | 1.727 | 0.788268083 | 0.032137929 | 126.56 | 124.4 | 75.6 | 81.67 | 72.65 | Not Found | High | High | High | Not Found | High | High | High | |
| Symplekin OS=Mus musculus OX=10090 GN=Sympk PE=1 SV=1 | Q80X82 | 0 | 12.61 | 3 | 2 | 8 | 2 | 1284 | 142.2 | 6.05 | 17.48 | 2.43 | 1.280956314 | 0.040323389 | 52.77 | 141.7 | 58.3 | 20.32 | 108.17 | Not Found | High | Peak Found | High | Not Found | High | Peak Found | High | |
| Syntaxin-1A OS=Mus musculus OX=10090 GN=Stx1a PE=1 SV=3 | O35526 | 0 | 100.904 | 45 | 9 | 126 | 9 | 288 | 33 | 5.24 | 117.17 | 2.996 | 1.583037624 | 0.001457049 | 127.34 | 124.9 | 75.1 | 196.55 | 171.58 | High | High | High | High | Peak Found | High | Peak Found | Peak Found | |
| TBC1 domain family member 8B OS=Mus musculus OX=10090 GN=Tbc1d8b PE=1 SV=1 | A3KGB4 | 0 | 25.503 | 5 | 3 | 20 | 3 | 1114 | 127.8 | 5.49 | 51.62 | 2.749 | 1.458906907 | 0.000186724 | 105.24 | 146.7 | 53.3 | 53.27 | 107.03 | Peak Found | High | Peak Found | High | High | High | High | High | |
| Transcription elongation factor SPT6 OS=Mus musculus OX=10090 GN=Supt6h PE=1 SV=2 | Q62383 | 0 | 102.705 | 13 | 13 | 78 | 13 | 1726 | 199 | 4.93 | 188.71 | 0.613 | -0.706041021 | 0.022163659 | 31.43 | 93.4 | 106.6 | 51.05 | 65.95 | High | High | High | High | High | High | High | High | |
| Trans-L-3-hydroxyproline dehydratase OS=Mus musculus OX=10090 GN=L3hypdh PE=1 SV=1 | Q9CXA2 | 0 | 24.307 | 7 | 3 | 51 | 3 | 354 | 37.8 | 6.7 | 145.81 | 1.757 | 0.813114191 | 0.000294893 | 86.55 | 100.7 | 99.3 | 41.26 | 58.15 | High | High | High | High | High | High | High | High | Acetyl [N-Term] |
| Translin-associated protein X OS=Mus musculus OX=10090 GN=Tsnax PE=1 SV=1 | Q9QZE7 | 0 | 17.029 | 25 | 4 | 9 | 4 | 290 | 32.9 | 6.55 | 23.31 | 0.328 | -1.60823228 | 9.10755E-08 | 12.34 | 86.6 | 113.4 | 53.18 | 40.64 | High | High | High | Peak Found | High | High | Peak Found | Peak Found | |
| Transmembrane glycoprotein NMB OS=Mus musculus OX=10090 GN=Gpnmb PE=1 SV=2 | Q99P91 | 0 | 65.146 | 3 | 2 | 47 | 2 | 574 | 63.6 | 7.61 | 132.67 | 1.443 | 0.5290713 | 0.017764595 | 73.82 | 128.3 | 71.7 | 54.24 | 42.95 | Not Found | High | High | High | Not Found | High | High | High | |
| Transmembrane protein 119 OS=Mus musculus OX=10090 GN=Tmem119 PE=1 SV=1 | Q8R138 | 0 | 53.011 | 23 | 3 | 43 | 3 | 280 | 29.4 | 4.51 | 93.04 | 0.628 | -0.671163536 | 0.005364244 | 37.15 | 78.9 | 121.1 | 21.16 | 41.42 | Peak Found | High | High | High | Not Found | High | High | High | |
| Triosephosphate isomerase OS=Mus musculus OX=10090 GN=Tpi1 PE=1 SV=4 | P17751 | 0 | 190.401 | 61 | 13 | 311 | 13 | 299 | 32.2 | 5.74 | 888.61 | 1.285 | 0.361768359 | 0.040323389 | 62.22 | 118.2 | 81.8 | 26.48 | 34.23 | High | High | High | High | High | High | High | High | |
| Tripartite motif-containing protein 26 OS=Mus musculus OX=10090 GN=Trim26 PE=2 SV=3 | Q99PN3 | 0 | 10.726 | 5 | 2 | 9 | 2 | 545 | 62.8 | 5.05 | 12.98 | 0.295 | -1.76121314 | 0.000857752 | 2.54 | 45.6 | 154.4 | 84.47 | | High | Peak Found | Not Found | Peak Found | High | Not Found | Not Found | High | |
| tRNA (guanine(37)-N1)-methyltransferase OS=Mus musculus OX=10090 GN=Trmt5 PE=1 SV=1 | Q9D0C4 | 0 | 15.177 | 5 | 2 | 9 | 2 | 501 | 56.8 | 7.91 | 21.8 | 0.161 | -2.634867407 | 5.68375E-15 | 2.14 | 30.9 | 169.1 | 81.37 | 30.61 | Not Found | High | Peak Found | High | Not Found | Peak Found | Peak Found | High | |
| tRNA methyltransferase 10 homolog C OS=Mus musculus OX=10090 GN=Trmt10c PE=1 SV=2 | Q3UFY8 | 0 | 9.112 | 9 | 2 | 4 | 2 | 414 | 48.4 | 9.38 | 11.47 | 0.045 | -4.473931188 | 9.08673E-13 | 0.15 | 8.6 | 191.4 | 156.01 | 12.26 | Not Found | Peak Found | High | Peak Found | Peak Found | Not Found | High | High | |
| Tyrosine-protein kinase Fyn OS=Mus musculus OX=10090 GN=Fyn PE=1 SV=4 | P39688 | 0 | 10.501 | 7 | 3 | 16 | 2 | 537 | 60.6 | 6.67 | 26.63 | 0.242 | -2.046921047 | 0.002256857 | 12.99 | 39 | 161 | 133.92 | 83.5 | Not Found | High | Peak Found | High | Not Found | Not Found | Peak Found | Peak Found | |
| UBX domain-containing protein 6 OS=Mus musculus OX=10090 GN=Ubxn6 PE=1 SV=1 | Q99PL6 | 0 | 34.417 | 15 | 4 | 20 | 4 | 442 | 49.8 | 8.54 | 42.42 | 0.384 | -1.380821784 | 0.007154989 | 20.38 | 58.5 | 141.5 | 54.38 | 74.61 | High | High | High | High | High | Peak Found | Peak Found | High | |
| Vacuolar protein sorting-associated protein 33B OS=Mus musculus OX=10090 GN=Vps33b PE=1 SV=1 | P59016 | 0 | 20.447 | 8 | 3 | 21 | 3 | 617 | 70.5 | 6.86 | 47.27 | 0.302 | -1.727379545 | 0.001577803 | 18.69 | 110.5 | 89.5 | 99.8 | 118.16 | High | High | High | High | High | Peak Found | Not Found | High | |
| Vacuolar protein sorting-associated protein 37C OS=Mus musculus OX=10090 GN=Vps37c PE=1 SV=1 | Q8R105 | 0 | 44.588 | 21 | 5 | 35 | 5 | 352 | 38.4 | 5.31 | 85.02 | 4.629 | 2.210700563 | 1.89812E-10 | 120.25 | 176.3 | 23.7 | 79.73 | 136.06 | High | High | High | High | High | High | Peak Found | High | Acetyl [N-Term] |
| Versican core protein OS=Mus musculus OX=10090 GN=Vcan PE=1 SV=2 | Q62059 | 0 | 65.315 | 3 | 6 | 77 | 6 | 3357 | 366.6 | 4.64 | 191.5 | 0.639 | -0.646112164 | 0.032582412 | 39.15 | 63.1 | 136.9 | 75.88 | 49.02 | Peak Found | High | High | High | Peak Found | High | High | High | |
| Vesicle-associated membrane protein-associated protein B OS=Mus musculus OX=10090 GN=Vapb PE=1 SV=3 | Q9QY76 | 0 | 21.79 | 19 | 3 | 8 | 2 | 243 | 26.9 | 7.78 | 20.66 | 6.206 | 2.633663697 | 1.0045E-08 | 118.36 | 172.2 | 27.8 | | 124.11 | Not Found | High | High | Not Found | Not Found | Peak Found | High | Peak Found | |
| Wiskott-Aldrich syndrome protein family member 2 OS=Mus musculus OX=10090 GN=Wasf2 PE=1 SV=1 | Q8BH43 | 0 | 54.393 | 13 | 4 | 70 | 4 | 497 | 54 | 5.53 | 169.33 | 1.389 | 0.474046599 | 0.017306261 | 87.48 | 122 | 78 | 63.32 | 70.92 | High | High | High | High | High | High | High | High | |
| Zyxin OS=Mus musculus OX=10090 GN=Zyx PE=1 SV=2 | Q62523 | 0 | 51.169 | 8 | 2 | 42 | 2 | 564 | 60.5 | 6.4 | 100.15 | 0.525 | -0.929610672 | 0.000442534 | 23.21 | 77 | 123 | 79.58 | 66.68 | High | High | High | High | Peak Found | High | High | High | |
