## Supplementary material for "Neurofilament accumulation disrupts autophagy in giant axonal neuropathy": Table S2

| **Ingenuity Canonical Pathways** | **-Log(p-value)** |
| --- | --- |
| PPARα/RXRα Activation | 1.38 |
| Actin Cytoskeleton Signaling | 1.68 |
| Estrogen Receptor Signaling | 0.483 |
| Glycogen Degradation II | 2.83 |
| Glycogen Degradation III | 2.66 |
| Semaphorin Neuronal Repulsive Signaling Pathway | 3.22 |
| Phagosome Formation | 0.613 |
| FAK Signaling | 0.561 |
| Paxillin Signaling | 2.26 |
| Thrombopoietin Signaling | 2.12 |
| CNTF Signaling | 2.12 |
| Growth Hormone Signaling | 2.08 |
| IL-2 Signaling | 2.08 |
| Role of Tissue Factor in Cancer | 2.08 |
| ERB2-ERBB3 Signaling | 2.05 |
| Amyotrophic Lateral Sclerosis Signaling | 1.99 |
| LPS/IL-1 Mediated Inhibition of RXR Function | 1.93 |
| Semaphorin Signaling in Neurons | 1.89 |
| Angiopoietin Signaling | 1.86 |
| Integrin Signaling | 2.34 |
| Intrinsic Prothrombin Activation Pathway | 1.85 |
| GM-CSF Signaling | 1.8 |
| IL-9 Signaling | 1.79 |
| Glycolysis I | 1.79 |
| Role of p14/p19ARF in Tumor Suppression | 1.79 |
| IL-3 Signaling | 1.75 |
| GP6 Signaling Pathway | 2.4 |
| JAK/STAT Signaling | 1.73 |
| Role of JAK2 in Hormone-like Cytokine Signaling | 1.69 |
| ERK/MAPK Signaling | 1.85 |
| Melanocyte Development and Pigmentation Signaling | 1.68 |
| PEDF Signaling | 1.65 |
| Neurotrophin/TRK Signaling | 1.65 |
| Glucose and Glucose-1-phosphate Degradation | 1.61 |
| Ascorbate Recycling (Cytosolic) | 1.61 |
| Glycerol-3-phosphate Shuttle | 1.61 |
| GDP-glucose Biosynthesis | 1.61 |
| Oncostatin M Signaling | 1.61 |
| Phagosome Maturation | 1.56 |
| TEC Kinase Signaling | 1.53 |
| Ephrin A Signaling | 1.53 |
| Acute Myeloid Leukemia Signaling | 1.52 |
| VEGF Signaling | 1.5 |
| Glycerol Degradation I | 1.44 |
| Acetyl-CoA Biosynthesis I (Pyruvate Dehydrogenase Complex) | 1.44 |

| | |
|---|---|
| Glutathione Redox Reactions II | 1.44 |
| Rapoport-Luebering Glycolytic Shunt | 1.44 |
| N-acetylglucosamine Degradation I | 1.44 |
| Fc Epsilon RI Signaling | 1.41 |
| PPAR Signaling | 1.41 |
| Role of IL-17A in Arthritis | 1.39 |
| Neuregulin Signaling | 1.39 |
| PAK Signaling | 1.39 |
| Chronic Myeloid Leukemia Signaling | 1.37 |
| Synaptogenesis Signaling Pathway | 0.701 |
| Creatine-phosphate Biosynthesis | 1.32 |
| Glucocorticoid Biosynthesis | 1.32 |
| Mineralocorticoid Biosynthesis | 1.32 |
| N-acetylglucosamine Degradation II | 1.32 |
| Clathrin-mediated Endocytosis Signaling | 1.31 |
| NGF Signaling | 1.29 |
| Sertoli Cell-Sertoli Cell Junction Signaling | 1.28 |
| Role of JAK1 and JAK3 in γc Cytokine Signaling | 1.26 |
| Assembly of RNA Polymerase I Complex | 1.22 |
| Sucrose Degradation V (Mammalian) | 1.22 |
| HGF Signaling | 1.2 |
| IL-6 Signaling | 1.2 |
| EGF Signaling | 1.18 |
| Erythropoietin Signaling Pathway | 1.17 |
| PKCθ Signaling in T Lymphocytes | 1.15 |
| Extrinsic Prothrombin Activation Pathway | 1.14 |
| Superoxide Radicals Degradation | 1.14 |
| ERBB4 Signaling | 1.14 |
| Adipogenesis pathway | 1.13 |
| HER-2 Signaling in Breast Cancer | 1.11 |
| Endometrial Cancer Signaling | 1.1 |
| Xenobiotic Metabolism PXR Signaling Pathway | 1.08 |
| Androgen Biosynthesis | 1.08 |
| Ketogenesis | 1.08 |
| VEGF Family Ligand-Receptor Interactions | 1.06 |
| Xenobiotic Metabolism Signaling | 1.05 |
| Insulin Receptor Signaling | 1.05 |
| Caveolar-mediated Endocytosis Signaling | 1.04 |
| IL-15 Signaling | 1.04 |
| Axonal Guidance Signaling | 1.04 |
| FcγRIIB Signaling in B Lymphocytes | 1.03 |
| IL-4 Signaling | 1.03 |
| DNA Double-Strand Break Repair by Homologous Recombination | 1.02 |
| Leukocyte Extravasation Signaling | 1.01 |
| CTLA4 Signaling in Cytotoxic T Lymphocytes | 1.01 |
| FGF Signaling | 1.01 |

| | |
|---|---|
| T Cell Receptor Signaling | 0.983 |
| Mevalonate Pathway I | 0.975 |
| Regulation of Cellular Mechanics by Calpain Protease | 0.975 |
| Inflammasome pathway | 0.975 |
| PD-1, PD-L1 cancer immunotherapy pathway | 0.975 |
| Renal Cell Carcinoma Signaling | 0.959 |
| GDNF Family Ligand-Receptor Interactions | 0.959 |
| Estrogen-Dependent Breast Cancer Signaling | 0.943 |
| MSP-RON Signaling In Macrophages Pathway | 0.943 |
| Glutathione Redox Reactions I | 0.932 |
| Superpathway of Geranylgeranyldiphosphate Biosynthesis I (via Mevalonate) | 0.932 |
| Pulmonary Healing Signaling Pathway | 0.921 |
| TR/RXR Activation | 0.914 |
| PDGF Signaling | 0.914 |
| Regulation of eIF4 and p70S6K Signaling | 0.91 |
| Production of Nitric Oxide and Reactive Oxygen Species in Macrophages | 0.9 |
| ERBB Signaling | 0.9 |
| Antioxidant Action of Vitamin C | 0.9 |
| Glutathione-mediated Detoxification | 0.893 |
| Vitamin-C Transport | 0.893 |
| IL-13 Signaling Pathway | 0.886 |
| Atherosclerosis Signaling | 0.886 |
| Th2 Pathway | 0.886 |
| Role of NFAT in Regulation of the Immune Response | 0.873 |
| Regulation of IL-2 Expression in Activated and Anergic T Lymphocytes | 0.873 |
| RAN Signaling | 0.86 |
| Glutaryl-CoA Degradation | 0.86 |
| Tryptophan Degradation III (Eukaryotic) | 0.86 |
| Methionine Degradation I (to Homocysteine) | 0.86 |
| Cysteine Biosynthesis III (mammalia) | 0.86 |
| Wound Healing Signaling Pathway | 0.857 |
| LXR/RXR Activation | 0.857 |
| Actin Nucleation by ARP-WASP Complex | 0.857 |
| Germ Cell-Sertoli Cell Junction Signaling | 0.848 |
| Role of NANOG in Mammalian Embryonic Stem Cell Pluripotency | 0.833 |
| UVA-Induced MAPK Signaling | 0.833 |
| Coagulation System | 0.827 |
| Non-Small Cell Lung Cancer Signaling | 0.807 |
| Mouse Embryonic Stem Cell Pluripotency | 0.807 |
| Virus Entry via Endocytic Pathways | 0.796 |
| Ephrin Receptor Signaling | 0.796 |
| Prolactin Signaling | 3.45 |
| Gap Junction Signaling | 0.78 |
| Systemic Lupus Erythematosus In B Cell Signaling Pathway | 0.78 |
| IL-22 Signaling | 0.77 |
| Role of JAK family kinases in IL-6-type Cytokine Signaling | 0.77 |

| | |
|---|---|
| SPINK1 Pancreatic Cancer Pathway | 0.77 |
| Systemic Lupus Erythematosus In T Cell Signaling Pathway | 0.75 |
| Gluconeogenesis I | 0.745 |
| RAR Activation | 0.742 |
| 3-phosphoinositide Biosynthesis | 0.742 |
| Telomerase Signaling | 0.74 |
| IL-12 Signaling and Production in Macrophages | 0.73 |
| CD28 Signaling in T Helper Cells | 0.73 |
| SAPK/JNK Signaling | 0.73 |
| Antiproliferative Role of TOB in T Cell Signaling | 0.721 |
| Renin-Angiotensin Signaling | 0.719 |
| Th1 and Th2 Activation Pathway | 0.719 |
| RHOGDI Signaling | 0.714 |
| IL-7 Signaling Pathway | 3.12 |
| Huntington's Disease Signaling | 0.68 |
| Apelin Liver Signaling Pathway | 0.678 |
| Glucocorticoid Receptor Signaling | 0.664 |
| Phospholipase C Signaling | 0.66 |
| Role of IL-17F in Allergic Inflammatory Airway Diseases | 0.658 |
| Superpathway of Methionine Degradation | 0.658 |
| Prostate Cancer Signaling | 0.654 |
| HOTAIR Regulatory Pathway | 0.654 |
| Superpathway of Inositol Phosphate Compounds | 0.648 |
| MSP-RON Signaling Pathway | 0.638 |
| IGF-1 Signaling | 0.635 |
| Dendritic Cell Maturation | 0.627 |
| RHOA Signaling | 0.627 |
| Inhibition of Angiogenesis by TSP1 | 0.622 |
| Tumor Microenvironment Pathway | 0.618 |
| FLT3 Signaling in Hematopoietic Progenitor Cells | 2.62 |
| Superpathway of Cholesterol Biosynthesis | 0.606 |
| Xenobiotic Metabolism CAR Signaling Pathway | 0.602 |
| Aryl Hydrocarbon Receptor Signaling | 0.595 |
| p70S6K Signaling | 0.595 |
| MSP-RON Signaling In Cancer Cells Pathway | 0.595 |
| Docosahexaenoic Acid (DHA) Signaling | 0.588 |
| IL-23 Signaling Pathway | 0.588 |
| NAD Signaling Pathway | 0.587 |
| 14-3-3-mediated Signaling | 0.587 |
| Glioma Signaling | 0.587 |
| Hereditary Breast Cancer Signaling | 0.587 |
| tRNA Charging | 0.573 |
| RAC Signaling | 0.572 |
| Insulin Secretion Signaling Pathway | 1.36 |
| TREM1 Signaling | 0.559 |
| BER (Base Excision Repair) Pathway | 0.559 |

| | |
|---|---|
| Systemic Lupus Erythematosus Signaling | 0.558 |
| Cardiac Hypertrophy Signaling | 0.545 |
| PTEN Signaling | 0.536 |
| Cell Cycle Control of Chromosomal Replication | 0.532 |
| SPINK1 General Cancer Pathway | 0.532 |
| Endothelin-1 Signaling | 0.529 |
| Apelin Pancreas Signaling Pathway | 0.519 |
| Role of PI3K/AKT Signaling in the Pathogenesis of Influenza | 0.507 |
| Acute Phase Response Signaling | 0.503 |
| Reelin Signaling in Neurons | 0.503 |
| Glioblastoma Multiforme Signaling | 0.503 |
| Regulation of the Epithelial-Mesenchymal Transition Pathway | 0.503 |
| Regulation Of The Epithelial Mesenchymal Transition By Growth Factors Pathway | 0.503 |
| Hepatic Fibrosis / Hepatic Stellate Cell Activation | 0.498 |
| Aldosterone Signaling in Epithelial Cells | 0.498 |
| Cancer Drug Resistance By Drug Efflux | 0.495 |
| PXR/RXR Activation | 0.484 |
| Lymphotoxin β Receptor Signaling | 0.484 |
| UVB-Induced MAPK Signaling | 0.484 |
| Cell Cycle: G2/M DNA Damage Checkpoint Regulation | 0.484 |
| Melanoma Signaling | 0.474 |
| Opioid Signaling Pathway | 0.474 |
| CD40 Signaling | 0.462 |
| IL-17A Signaling in Airway Cells | 0.462 |
| Molecular Mechanisms of Cancer | 0.461 |
| B Cell Receptor Signaling | 0.456 |
| Sirtuin Signaling Pathway | 0.44 |
| Adrenomedullin signaling pathway | 0.44 |
| Senescence Pathway | 0.44 |
| D-myo-inositol (1,4,5,6)-Tetrakisphosphate Biosynthesis | 0.434 |
| D-myo-inositol (3,4,5,6)-tetrakisphosphate Biosynthesis | 0.434 |
| Leptin Signaling in Obesity | 0.433 |
| Xenobiotic Metabolism AHR Signaling Pathway | 0.433 |
| ID1 Signaling Pathway | 0.419 |
| VDR/RXR Activation | 0.416 |
| Macropinocytosis Signaling | 0.416 |
| PI3K/AKT Signaling | 0.413 |
| Role of BRCA1 in DNA Damage Response | 0.399 |
| Glioma Invasiveness Signaling | 0.399 |
| D-myo-inositol-5-phosphate Metabolism | 0.399 |
| HIF1α Signaling | 0.395 |
| Tight Junction Signaling | 0.39 |
| ILK Signaling | 0.39 |
| 3-phosphoinositide Degradation | 0.385 |
| NF-κB Activation by Viruses | 0.383 |
| Th1 Pathway | 0.383 |

| | |
|---|---|
| FXR/RXR Activation | 0.376 |
| Mitochondrial Dysfunction | 0.368 |
| ERK5 Signaling | 0.368 |
| EIF2 Signaling | 0.36 |
| Antiproliferative Role of Somatostatin Receptor 2 | 0.354 |
| TGF-β Signaling | 0.354 |
| Apelin Adipocyte Signaling Pathway | 0.347 |
| NRF2-mediated Oxidative Stress Response | 0.343 |
| Thrombin Signaling | 0.343 |
| BMP signaling pathway | 0.341 |
| mTOR Signaling | 0.335 |
| Role of NFAT in Cardiac Hypertrophy | 0.335 |
| LPS-stimulated MAPK Signaling | 0.334 |
| RANK Signaling in Osteoclasts | 0.334 |
| cAMP-mediated signaling | 0.332 |
| Remodeling of Epithelial Adherens Junctions | 0.328 |
| Apelin Cardiomyocyte Signaling Pathway | 0.322 |
| Protein Kinase A Signaling | 0.318 |
| Role of Pattern Recognition Receptors in Recognition of Bacteria and Viruses | 0.316 |
| IL-15 Production | 0.316 |
| ICOS-ICOSL Signaling in T Helper Cells | 0.316 |
| Thyroid Cancer Signaling | 0.316 |
| Fcγ Receptor-mediated Phagocytosis in Macrophages and Monocytes | 0.299 |
| Small Cell Lung Cancer Signaling | 0.294 |
| NER (Nucleotide Excision Repair, Enhanced Pathway) | 0.294 |
| Ceramide Signaling | 0.289 |
| Neuropathic Pain Signaling In Dorsal Horn Neurons | 0.284 |
| p53 Signaling | 0.279 |
| Sphingosine-1-phosphate Signaling | 0.279 |
| p38 MAPK Signaling | 0.279 |
| Colorectal Cancer Metastasis Signaling | 0.277 |
| BAG2 Signaling Pathway | 0.274 |
| α-Adrenergic Signaling | 0.269 |
| Regulation of Actin-based Motility by Rho | 0.26 |
| Apoptosis Signaling | 0.26 |
| Xenobiotic Metabolism General Signaling Pathway | 0.26 |
| Autophagy | 0.258 |
| Neuroinflammation Signaling Pathway | 0.258 |
| G-Protein Coupled Receptor Signaling | 0.252 |
| Oxytocin Signaling Pathway | 0.25 |
| IL-17 Signaling | 0.247 |
| Cholecystokinin/Gastrin-mediated Signaling | 0.247 |
| Nitric Oxide Signaling in the Cardiovascular System | 0.247 |
| Sumoylation Pathway | 0.243 |
| Signaling by Rho Family GTPases | 0.242 |
| CDC42 Signaling | 0.235 |

| | |
|---|---|
| AMPK Signaling | 0.228 |
| Pancreatic Adenocarcinoma Signaling | 0.224 |
| CCR3 Signaling in Eosinophils | 0.22 |
| G Beta Gamma Signaling | 0.22 |
| Iron homeostasis signaling pathway | 0.22 |
| Synaptic Long Term Potentiation | 0.217 |
| P2Y Purigenic Receptor Signaling Pathway | 0.213 |
| T Cell Exhaustion Signaling Pathway | 0.213 |
| eNOS Signaling | 0.21 |
| Hepatic Cholestasis | 0.206 |
| Ferroptosis Signaling Pathway | 0.206 |
| fMLP Signaling in Neutrophils | 0.203 |
| Gα12/13 Signaling | 0.203 |
| HMGB1 Signaling | 0.197 |
| Gαi Signaling | 0.197 |
| Oxytocin In Brain Signaling Pathway | 0 |
| CLEAR Signaling Pathway | 0 |
| SNARE Signaling Pathway | 0 |
| Circadian Rhythm Signaling | 0 |
| IL-8 Signaling | 0 |
| CXCR4 Signaling | 0 |
| Relaxin Signaling | 0 |
| GNRH Signaling | 0 |
| Human Embryonic Stem Cell Pluripotency | 0 |
| CREB Signaling in Neurons | 0 |
| Type II Diabetes Mellitus Signaling | 0 |
| Role of Osteoblasts, Osteoclasts and Chondrocytes in Rheumatoid Arthritis | 0 |
| Ovarian Cancer Signaling | 0 |
| Role of Macrophages, Fibroblasts and Endothelial Cells in Rheumatoid Arthritis | 0 |
| Breast Cancer Regulation by Stathmin1 | 0 |
| PI3K Signaling in B Lymphocytes | 0 |
| Epithelial Adherens Junction Signaling | 0 |
| Gαq Signaling | 0 |
| Sperm Motility | 0 |
| Cardiac β-adrenergic Signaling | 0 |
| Protein Ubiquitination Pathway | 0 |
| NF-κB Signaling | 0 |
| Osteoarthritis Pathway | 0 |
| Endocannabinoid Developing Neuron Pathway | 0 |
| Endocannabinoid Cancer Inhibition Pathway | 0 |
| Apelin Endothelial Signaling Pathway | 0 |
| Cardiac Hypertrophy Signaling (Enhanced) | 0 |
| Necroptosis Signaling Pathway | 0 |

| Ratio | z-score | Proteins |
|---|---|---|
| 0.0377 | 2 | GPD1,NCOR2,SOS1,STAT5B |
| 0.0391 | 0 | Actn3,ITGA7,PIK3C2A,SOS1,WASF2 |
| 0.017 | 0 | NCOR2,PIK3C2A,SOD2,SOS1 |
| 0.4 | | PGM1,PYGL |
| 0.333 | | PGM1,PYGL |
| 0.0706 | -0.447 | CRMP1,FYN,ITGA7,PIK3C2A,PLXNA1,VCAN |
| 0.0184 | -0.447 | FYN,ITGA7,PIK3C2A,SOS1,WASF2 |
| 0.0175 | -0.447 | COL1A2,FYN,ITGA7,PIK3C2A,SOS1 |
| 0.069 | | Actn3,ITGA7,PIK3C2A,SOS1 |
| 0.0909 | | PIK3C2A,SOS1,STAT5B |
| 0.0909 | | PIK3C2A,RPS6KA1,SOS1 |
| 0.0882 | | PIK3C2A,RPS6KA1,STAT5B |
| 0.0882 | | PIK3C2A,SOS1,STAT5B |
| 0.0615 | | FYN,PIK3C2A,RPS6KA1,STAT5B |
| 0.0857 | | PIK3C2A,SOS1,STAT5B |
| 0.058 | | NEFH,NEFL,PIK3C2A,PRPH |
| 0.045 | | ABCB1,ALDH1L1,FABP4,FABP5,HMGCS1 |
| 0.075 | | CRMP1,FYN,PLXNA1 |
| 0.0732 | | PIK3C2A,SOS1,STAT5B |
| 0.0476 | -0.816 | Actn3,ARHGAP5,FYN,ITGA7,PIK3C2A,SOS1 |
| 0.133 | | COL1A2,F13A1 |
| 0.0698 | | PIK3C2A,SOS1,STAT5B |
| 0.125 | | PIK3C2A,STAT5B |
| 0.125 | | PGAM1,TPI1 |
| 0.125 | | PIK3C2A,UBTF |
| 0.0667 | | PIK3C2A,SOS1,STAT5B |
| 0.0755 | -1 | COL1A2,COL4A1,FYN,PIK3C2A |
| 0.0652 | | PIK3C2A,SOS1,STAT5B |
| 0.111 | | SIRPA,STAT5B |
| 0.0431 | -1 | FYN,ITGA7,PIK3C2A,RPS6KA1,SOS1 |
| 0.0625 | | PIK3C2A,RPS6KA1,SOS1 |
| 0.0612 | | PIK3C2A,SOD2,WASF2 |
| 0.0612 | | PIK3C2A,RPS6KA1,SOS1 |
| 0.5 | | PGM1 |
| 0.5 | | GLRX |
| 0.5 | | GPD1 |
| 0.5 | | PGM1 |
| 0.1 | | SOS1,STAT5B |
| 0.043 | | CTSB,CTSH,STX1A,VPS33B |
| 0.0421 | | FYN,ITGA7,PIK3C2A,STAT5B |
| 0.0909 | | FYN,PIK3C2A |
| 0.0545 | | PIK3C2A,SOS1,STAT5B |
| 0.0536 | | Actn3,PIK3C2A,SOS1 |
| 0.333 | | GPD1 |
| 0.333 | | PDHA1 |

| | | |
|---|---|---|
| 0.333 | | GLRX |
| 0.333 | | PGAM1 |
| 0.333 | | AMDHD2 |
| 0.0492 | | FYN,PIK3C2A,SOS1 |
| 0.0492 | | NCOR2,SOS1,STAT5B |
| 0.0769 | | PIK3C2A,RPS6KA1 |
| 0.0484 | | ITGA7,SOS1,STAT5B |
| 0.0484 | | ITGA7,PIK3C2A,SOS1 |
| 0.0476 | | PIK3C2A,SOS1,STAT5B |
| 0.0214 | -1 | FYN,PIK3C2A,SOS1,STX1A |
| 0.25 | | CKMT1A/CKMT1B |
| 0.25 | | Gsta4 |
| 0.25 | | Gsta4 |
| 0.25 | | AMDHD2 |
| 0.0357 | | CD2AP,CLU,PIK3C2A,RAB4A |
| 0.0441 | | PIK3C2A,RPS6KA1,SOS1 |
| 0.0351 | | Actn3,CLINT1,ITGA7,SYMPK |
| 0.0645 | | PIK3C2A,STAT5B |
| 0.2 | | UBTF |
| 0.2 | | TPI1 |
| 0.0405 | | ITGA7,PIK3C2A,SOS1 |
| 0.0405 | | ABCB1,PIK3C2A,SOS1 |
| 0.0588 | | PIK3C2A,SOS1 |
| 0.0395 | | PIK3C2A,SOS1,STAT5B |
| 0.0385 | | FYN,PIK3C2A,SOS1 |
| 0.167 | | F13A1 |
| 0.167 | | SOD2 |
| 0.0556 | | PIK3C2A,SOS1 |
| 0.038 | | FABP4,RPS6KA1,STAT5B |
| 0.0305 | | AHCTF1,FYN,PIK3C2A,SOS1 |
| 0.0526 | | PIK3C2A,SOS1 |
| 0.0361 | | ABCB1,ALDH1L1,NCOR2 |
| 0.143 | | Gsta4 |
| 0.143 | | HMGCS1 |
| 0.05 | | PIK3C2A,SOS1 |
| 0.0292 | | ABCB1,ALDH1L1,NCOR2,PIK3C2A |
| 0.0349 | | FYN,PIK3C2A,SOS1 |
| 0.0488 | | FYN,ITGA7 |
| 0.0488 | | PIK3C2A,STAT5B |
| 0.0237 | | FYN,ITGA7,PIK3C2A,PLXNA1,SLIT2,SOS1 |
| 0.0476 | | PIK3C2A,SOS1 |
| 0.0476 | | PIK3C2A,SOS1 |
| 0.125 | | LIG1 |
| 0.0337 | | Actn3,ARHGAP5,PIK3C2A |
| 0.0465 | | FYN,PIK3C2A |
| 0.0465 | | PIK3C2A,SOS1 |

| | | |
|---|---|---|
| 0.0326 | | FYN,PIK3C2A,SOS1 |
| 0.111 | | HMGCS1 |
| 0.0444 | | Actn3,ITGA7 |
| 0.111 | | CTSB |
| 0.0444 | | PIK3C2A,STAT5B |
| 0.0435 | | PIK3C2A,SOS1 |
| 0.0435 | | PIK3C2A,SOS1 |
| 0.0426 | | PIK3C2A,STAT5B |
| 0.0426 | | PIK3C2A,SOS1 |
| 0.1 | | Gsta4 |
| 0.1 | | HMGCS1 |
| 0.0306 | | FYN,SOS1,STAT5B |
| 0.0408 | | NCOR2,PIK3C2A |
| 0.0408 | | PIK3C2A,SOS1 |
| 0.0303 | | ITGA7,PIK3C2A,SOS1 |
| 0.03 | | CLU,PIK3C2A,SIRPA |
| 0.04 | | PIK3C2A,SOS1 |
| 0.04 | | GLRX,STAT5B |
| 0.0909 | | Gsta4 |
| 0.0909 | | GLRX |
| 0.0392 | | FYN,PIK3C2A |
| 0.0392 | | CLU,COL1A2 |
| 0.0392 | | PIK3C2A,STAT5B |
| 0.0291 | | FYN,PIK3C2A,SOS1 |
| 0.0385 | | FYN,SOS1 |
| 0.0833 | | RAN |
| 0.0833 | | L3HYPDH |
| 0.0833 | | L3HYPDH |
| 0.0833 | | PRMT5 |
| 0.0833 | | PRMT5 |
| 0.0286 | | COL1A2,COL4A1,SOS1 |
| 0.0377 | | CLU,NCOR2 |
| 0.0377 | | ITGA7,SOS1 |
| 0.0283 | | Actn3,CLINT1,PIK3C2A |
| 0.0364 | | PIK3C2A,SOS1 |
| 0.0364 | | PIK3C2A,RPS6KA1 |
| 0.0769 | | F13A1 |
| 0.0351 | | PIK3C2A,SOS1 |
| 0.0351 | | PIK3C2A,SOS1 |
| 0.0345 | | FYN,PIK3C2A |
| 0.0268 | | FYN,ITGA7,SOS1 |
| 0.1 | -2 | FYN,NMI,PIK3C2A,SOS1,STAT5B |
| 0.0263 | | DBN1,PIK3C2A,SOS1 |
| 0.0263 | | FYN,PIK3C2A,SOS1 |
| 0.0667 | | STAT5B |
| 0.0667 | | STAT5B |

| | | |
|---|---|---|
| 0.0667 | | CTSB |
| 0.0254 | | PIK3C2A,RAB4A,SOS1 |
| 0.0625 | | PGAM1 |
| 0.0252 | | ARID1A,NCOR2,STAT5B |
| 0.0252 | | NUDT9,PIK3C2A,SIRPA |
| 0.0317 | | PIK3C2A,SOS1 |
| 0.0312 | | CLU,PIK3C2A |
| 0.0312 | | FYN,PIK3C2A |
| 0.0312 | | PIK3C2A,SOS1 |
| 0.0588 | | RPS6KA1 |
| 0.0308 | | PIK3C2A,SOS1 |
| 0.0308 | | PIK3C2A,STAT5B |
| 0.0244 | | ARHGAP5,ITGA7,WASF2 |
| 0.118 | -2 | FYN,PIK3C2A,SOS1,STAT5B |
| 0.0209 | | NCOR2,PIK3C2A,SOS1,STX1A |
| 0.0526 | | COL1A2 |
| 0.0192 | | ARID1A,NCOR2,PIK3C2A,SOS1,STAT5B |
| 0.0229 | | FYN,ITGA7,SOS1 |
| 0.05 | | RPS6KA1 |
| 0.05 | | PRMT5 |
| 0.0278 | | PIK3C2A,SOS1 |
| 0.0278 | | COL1A2,PIK3C2A |
| 0.0226 | | NUDT9,PIK3C2A,SIRPA |
| 0.0476 | | PIK3C2A |
| 0.027 | | PIK3C2A,SOS1 |
| 0.0267 | | COL1A2,PIK3C2A |
| 0.0267 | | ARHGAP5,PLXNA1 |
| 0.0455 | | FYN |
| 0.0263 | | COL1A2,PIK3C2A |
| 0.087 | -2 | PIK3C2A,RPS6KA1,SOS1,STAT5B |
| 0.0435 | | HMGCS1 |
| 0.0256 | | ABCB1,ALDH1L1 |
| 0.0253 | | ALDH1L1,NCOR2 |
| 0.0253 | | PIK3C2A,SOS1 |
| 0.0253 | | PIK3C2A,SOS1 |
| 0.0417 | | PIK3C2A |
| 0.0417 | | PIK3C2A |
| 0.025 | | PIK3C2A,SOD2 |
| 0.025 | | PIK3C2A,RPS6KA1 |
| 0.025 | | PIK3C2A,SOS1 |
| 0.025 | | ARID1A,PIK3C2A |
| 0.04 | | LARS2 |
| 0.0244 | | ITGA7,PIK3C2A |
| 0.0321 | -2 | FYN,PDHA1,PIK3C2A,STAT5B,STX1A |
| 0.0385 | | STAT5B |
| 0.0385 | | LIG1 |

| | |
|---|---|
| 0.0238 | PIK3C2A,SOS1 |
| 0.0199 | PIK3C2A,RPS6KA1,SOS1 |
| 0.023 | ITGA7,SOS1 |
| 0.0357 | LIG1 |
| 0.0357 | PIK3C2A |
| 0.0227 | PIK3C2A,SOS1 |
| 0.0345 | PIK3C2A |
| 0.0333 | PIK3C2A |
| 0.0217 | SOD2,SOS1 |
| 0.0217 | FYN,PIK3C2A |
| 0.0217 | PIK3C2A,SOS1 |
| 0.0217 | PIK3C2A,SOS1 |
| 0.0217 | PIK3C2A,SOS1 |
| 0.0215 | COL1A2,COL4A1 |
| 0.0215 | PIK3C2A,SOS1 |
| 0.0323 | ABCB1 |
| 0.0312 | ABCB1 |
| 0.0312 | PIK3C2A |
| 0.0312 | PIK3C2A |
| 0.0312 | RPS6KA1 |
| 0.0303 | PIK3C2A |
| 0.0181 | FYN,RPS6KA1,SOS1 |
| 0.0294 | PIK3C2A |
| 0.0294 | PIK3C2A |
| 0.0166 | FYN,ITGA7,PIK3C2A,SOS1 |
| 0.02 | PIK3C2A,SOS1 |
| 0.0172 | PDHA1,PGAM1,SOD2 |
| 0.0194 | PIK3C2A,SOS1 |
| 0.0172 | PDHA1,PIK3C2A,SOD2 |
| 0.0192 | NUDT9,SIRPA |
| 0.0192 | NUDT9,SIRPA |
| 0.027 | PIK3C2A |
| 0.027 | ALDH1L1 |
| 0.0187 | FYN,PIK3C2A |
| 0.0256 | NCOR2 |
| 0.0256 | PIK3C2A |
| 0.0185 | ITGA7,SOS1 |
| 0.0244 | ARID1A |
| 0.0244 | PIK3C2A |
| 0.018 | NUDT9,SIRPA |
| 0.0179 | PIK3C2A,RAN |
| 0.0177 | STX1A,SYMPK |
| 0.0177 | Actn3,PIK3C2A |
| 0.0175 | NUDT9,SIRPA |
| 0.0233 | PIK3C2A |
| 0.0233 | PIK3C2A |

| | | |
|---|---|---|
| 0.0227 | | CLU |
| 0.0169 | | PDHA1,SOD2 |
| 0.0222 | | RPS6KA1 |
| 0.0167 | | PIK3C2A,SOS1 |
| 0.0213 | | PIK3C2A |
| 0.0213 | | SOS1 |
| 0.0208 | | Gsta4 |
| 0.0161 | | PIK3C2A,SOD2 |
| 0.0161 | | PIK3C2A,SOS1 |
| 0.0204 | | SOS1 |
| 0.0159 | | PIK3C2A,RPS6KA1 |
| 0.0159 | | PIK3C2A,SOS1 |
| 0.02 | | PIK3C2A |
| 0.02 | | PIK3C2A |
| 0.0157 | | AKAP12,RPS6KA1 |
| 0.0196 | | Actn3 |
| 0.0192 | | PIK3C2A |
| 0.0143 | | AKAP12,PYGL,SIRPA |
| 0.0189 | | PIK3C2A |
| 0.0189 | | FYN |
| 0.0189 | | PIK3C2A |
| 0.0189 | | PIK3C2A |
| 0.0179 | | FYN |
| 0.0175 | | PIK3C2A |
| 0.0175 | | LIG1 |
| 0.0172 | | PIK3C2A |
| 0.0169 | | PIK3C2A |
| 0.0167 | | PIK3C2A |
| 0.0167 | | PIK3C2A |
| 0.0167 | | RPS6KA1 |
| 0.014 | | PIK3C2A,SOS1 |
| 0.0164 | | CTSB |
| 0.0161 | | PYGL |
| 0.0156 | | ITGA7 |
| 0.0156 | | RPS6KA1 |
| 0.0156 | | PIK3C2A |
| 0.0134 | | PIK3C2A,VPS33B |
| 0.0134 | | PIK3C2A,SOD2 |
| 0.0125 | -2 | FYN,PIK3C2A,RPS6KA1,SOS1 |
| 0.0132 | | PIK3C2A,SOS1 |
| 0.0149 | | PIK3C2A |
| 0.0149 | | SOS1 |
| 0.0149 | | PIK3C2A |
| 0.0147 | | RAN |
| 0.0129 | | ITGA7,PIK3C2A |
| 0.0143 | | ITGA7 |

| | |
|---|---|
| 0.0125 | ARID1A,PIK3C2A |
| 0.0137 | PIK3C2A |
| 0.0135 | PIK3C2A |
| 0.0135 | SOS1 |
| 0.0135 | STAT5B |
| 0.0133 | RPS6KA1 |
| 0.0132 | PIK3C2A |
| 0.0132 | PIK3C2A |
| 0.013 | PIK3C2A |
| 0.0128 | ABCB1 |
| 0.0128 | CTSB |
| 0.0127 | PIK3C2A |
| 0.0127 | PIK3C2A |
| 0.0123 | PIK3C2A |
| 0.0123 | SOS1 |
| 0.0098 | PIK3C2A |
| 0.011 | CTSB,LAMTOR2 |
| 0.0114 | STX1A |
| 0.0067 | FYN |
| 0.0083 | PIK3C2A |
| 0.0103 | PIK3C2A |
| 0.0118 | PIK3C2A |
| 0.0098 | SOS1 |
| 0.0116 | PIK3C2A |
| 0.0111 | PIK3C2A,RPS6KA1,SOS1 |
| 0.0111 | PIK3C2A |
| 0.0086 | PIK3C2A |
| 0.0114 | PIK3C2A |
| 0.0064 | PIK3C2A |
| 0.0076 | PIK3C2A,SOS1 |
| 0.012 | FYN |
| 0.01 | FYN |
| 0.0098 | PIK3C2A |
| 0.0083 | FYN |
| 0.0098 | AKAP12 |
| 0.0058 | UBR1 |
| 0.0109 | PIK3C2A |
| 0.0088 | ITGA7 |
| 0.012 | PIK3C2A |
| 0.0115 | PIK3C2A |
| 0.0116 | PIK3C2A |
| 0.0075 | ITGA7,PIK3C2A |
| 0.0119 | PYGL |
